# Probabilistic classification of gene-by-treatment interactions on molecular count phenotypes

**DOI:** 10.1101/2024.08.03.605142

**Authors:** Yuriko Harigaya, Nana Matoba, Brandon D. Le, Jordan M. Valone, Jason L. Stein, Michael I. Love, William Valdar

## Abstract

Genetic variation can modulate response to treatment (G×T) or environmental stimuli (G×E), both of which may be highly consequential in biomedicine. An effective approach to identifying G×T signals and gaining insight into molecular mechanisms is mapping quantitative trait loci (QTL) of molecular count phenotypes, such as gene expression and chromatin accessibility, under multiple treatment conditions, which is termed response molecular QTL mapping. Although standard approaches evaluate the interaction between genetics and treatment conditions, they do not distinguish between meaningful interpretations such as whether a genetic effect is only observed in the treated condition or whether a genetic effect is observed but accentuated in the treated condition. To address this gap, we have developed a downstream method for classifying response molecular QTLs into subclasses with meaningful genetic interpretations. Our method uses Bayesian model selection and assigns posterior probabilities to different types of G×T interactions for a given feature-SNP pair. We compare linear and nonlinear regression of log-scale counts, noting that the latter accounts for an expected biological relationship between the genotype and the molecular count phenotype. Through simulation and application to existing datasets of molecular response QTLs, we show that our method provides an intuitive and well-powered framework to report and interpret G×T interactions. We provide a software package, ClassifyGxT, which is available at https://github.com/yharigaya/classifygxt.

**Author summary:** Responses to treatment, such as drug, therapeutic intervention, and infection, can vary across individuals at least in part due to their genetic backgrounds. This phenomenon can be conceptualized as a manifestation of gene-by-treatment (G×T) or gene-by-environment (G×E) interactions, which refer to non-additive effects of genotype and treatment on traits and phenotypes. An understading of G×T or G×E interactions can potentially improve strategies for prevention and treatment of diseases, for example by selecting treatments for which a patient is most likely to respond given their genetic information, or enhanced screening for individuals most susceptible to environmental exposures. An effective approach to G×T discovery is response quantitative trait loci (QTL) mapping, where the effect of the treatment on the association between the genotype and phenotype is examined using a linear regression model including the genotype, treatment, and G×T interaction terms. Despite its effectiveness in identifying a large number of associations, the response QTL mapping relies on hypothesis testing, which does not provide classification of different types of G×T interactions. Herein, we propose a use of Bayesian model selection to classify the G×T types of response QTLs and provide a software package for this method. In addition to standard linear regression, our package provides an option to use nonlinear regression that is suited for molecular count phenotypes, such as gene expression and chromatin accessibility, measured by sequencing-based techniques. It also provides an option to use mixed effect models to accommodate replicate measurements per donor, which are common in data generated in *in-vitro* cell systems.

## Introduction

Gene-by-treatment (G×T) interactions describe associations between genotype and phenotype that are modulated by treatment or, equivalently, phenotypic responses to treatment that are modulated by genotype. An understanding of G×T interactions is crucial for interpreting disease-associated genetic variants and, eventually, for clinical decision-making. These phenomena may also be called gene-by-environment interactions in a broader context [1].

An effective approach to G×T discovery is quantitative trait loci (QTL) mapping, which examines statistical associations between genotype and phenotype. For G×T discovery, there exist at least three types of methods, all of which utilize phenotype data in control and treated conditions. In the first approach, which we call the “stratified” approach, the association between genotype and phenotype is examined separately in each of the conditions. Then, genetic variants involved in G×T interactions are identified as those that exhibit significant association with the genotype only in the treated condition [2, 3]. In the second approach, which requires paired data and which we call the “delta” approach, the difference in the phenotypes between control and treated conditions is used as an outcome variable [2, 4–6]. In the third approach, which we term the “interaction” approach but is also called response QTL mapping, phenotype data in the control and treated conditions is jointly modeled using a linear model including the genotype, treatment, and G×T interaction terms [7–11]. Multiple studies undertaking these approaches have been effective in identifying a large number of G×T interactions, which have, in turn, led to valuable biological insights about genetic control of cellular and tissue responses to treatment.

Despite their effectiveness in G×T detection, however, the aforementioned approaches are not well suited to classifying the type of G×T interaction that is present. Specifically, the way in which genetics differ under alternative treatments can fall into starkly different classes. To illustrate this point, we consider graphical representation of G×T interactions through a linear regression framework, where the phenotype is regressed on the genotype at the site of a single nucleotide polymorphism (SNP) in the control and treated conditions separately (Fig 1). Without loss of generality, we will refer to genetic variants as SNPs. Whereas parallel regression lines indicate the absence of G×T interactions (“no-G×T”), non-parallel regression lines represent the presence thereof. In some cases, association between the phenotype and genotype is present in both conditions but to different extents, which we call the “altered” genotype effect pattern. In other cases, association between the phenotype and genotype is present only in the treated condition, which we call the “induced” genotype effect pattern, where the SNP would be discovered as a putative *cis*-regulatory variant only upon treatment. The induced pattern may occur, for example, when the transcription factor whose binding is modulated by the genetic variant is only activated after treatment, or when the transcriptional response necessitates co-factors that are only active after treatment [9, 12]. Furthermore, G×T interaction can result in a situation where the treatment has the opposite effects (i.e., effects with different signs) depending on the genotype, which is often called a “crossover” interaction effect [13]. Formally distinguishing between these different types of G×T interactions can have biological and biomedical implications. In the current practice, a typical mapping procedure returns a list of feature-SNP pairs with significant associations from *p*-value based hypothesis testing and does not provide probabilistic classification of these different types of G×T interactions without requiring *ad hoc* post-processing (see Results for details). This limitation can potentially hamper meaningful interpretation and prioritization for downstream analysis. Some studies have undertaken probabilistic approaches to detecting G×T or G×E interactions, though these proposed methods focused specifically on paired designs [14, 15].

**Fig 1.**
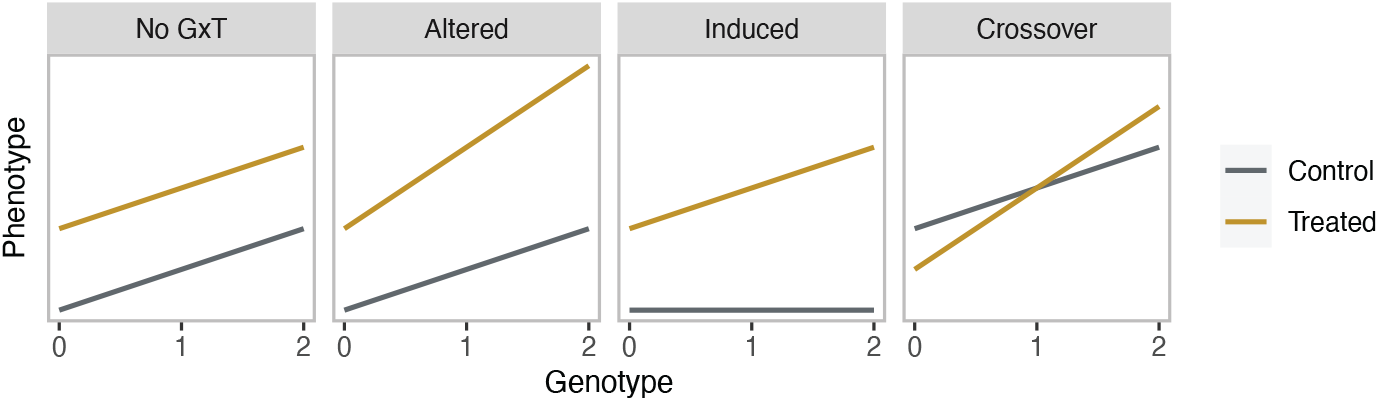
Illustration of the presence and absence of G×T interactions. Schematics of hypothetical linear regression where the phenotype is regressed on the genotype, coded as 0, 1, and 2, separately in the control (gray) and treated (brown) conditions. Shown are examples of lack of G×T interaction (no-G×T), the altered genotype effect pattern, where the genotype effect is present in both conditions but to different strengths, the induced genotype effect pattern, where the genotype effect emerges only upon treatment, and the crossover interaction, where the treatment has the opposite effect depending on the genotype.

Another potential limitation of the current practice in G×T analysis is specifically pertinent to molecular QTL mapping, where the phenotype is molecular count data, such as gene expression or chromatin accessibility. That is, in a typical procedure of molecular QTL mapping, the molecular count data is transformed, commonly using the log function, and a linear relationship is assumed between the genotype and the transformed molecular phenotype. However, previous studies have shown that, for the majority of the cases, the linearity holds in the original count scale, not in the transformed scale and that the linearity assumption can result in inaccurate estimation of the genotype effect [16–18]. Such inaccuracy can potentially lead to a pitfall in G×T analysis, which infers the *difference* in genotype effects between treatment conditions, while it may not substantially impact single-condition molecular QTL mapping, which infers the *presence* of genotype effects. To our knowledge, the issue of allelic additivity and modeling of molecular counts for response QTL classification has not been addressed or evaluated in the published literature.

In this study, we developed a method for probabilistic classification of types of G×T interactions on molecular count phenotypes using Bayesian model selection (BMS). Our method takes a set of SNPs identified by standard mapping procedures, such as the “stratified”, “delta”, and “interaction” approaches, as input and generates posterior probabilities for candidate models representing different types of G×T interactions.

Within this framework, we examined three modeling approaches: 1) applying a linear model to the log transformation of the molecular phenotype (log-LM); 2) applying a linear model to the rank-based inverse Normal transformation (RINT) [19] of the molecular phenotype (RINT-LM); and 3) applying a nonlinear model to the log transformation of the molecular phenotype that explicitly accounts for allelic additivity (log-NL). In our simulation experiments, in which we generated data according to the known nonlinear relationship between the genotype and transformed molecular phenotype, we observed that nonlinear regression (log-NL) exhibited moderately but consistently higher accuracy than linear regression (log-LM and RINT-LM). We also observed that empirical Bayes approaches to elicit priors can improve the accuracy as assessed by the posterior probabilities of the correct model as well as those of the incorrect models. We then illustrate the utility of our method through reanalysis of previously published gene expression and chromatin accessibility data in human primary neural progenitor cells (hNPCs) as examples.

## Results

### A Bayesian model selection (BMS) framework for classifying G×T interactions with molecular count phenotypes

#### Overview of the framework

We consider effects of treatment on associations between genotypes and molecular phenotypes, such as gene expression and chromatin accessibility measured by RNA-sequencing (RNA-seq) [20] and assay of transposase-accessible chromatin sequencing (ATAC-seq) [21] techniques. In what follows, we refer to the measurement unit for a given molecular phenotype as a feature. For example, genes and accessible chromatin regions, the latter of which are often called candidate *cis*-regulatory elements (cCREs), are considered as features for gene expression and chromatin accessibility, respectively.

The goal of our framework is to provide a principled approach to interpreting and prioritizing feature-SNP pairs that have been identified by standard methods, such as the “stratified”, “delta”, and “interaction” approaches (Introduction), for further experimental or bioinformatic interrogation. The types of feature-SNP pairs include expression QTLs (eQTLs) and chromatin accessibility QTLs (caQTLs). The framework takes the genotype and molecular phenotype data across subjects in control and treated conditions for a pre-selected set of feature-SNP pairs as input. The final output is posterior probabilities of eight possible models, representing whether the regression coefficients for the genotype, treatment, and G×T interaction terms are non-zero. We refer to the eight model categories using a vector of indicator variables **m** = (*m*_*g*_, *m*_*t*_, *m*_*g×t*_) ∈ *{*0, 1*}*^3^, where *m*_*g*_, *m*_*t*_, and *m*_*g×t*_ denote the inclusion and exclusion of the genotype, treatment, and G×T interaction terms in the model, respectively (Fig 2). In the simplest form using linear regression, the model can be cast as

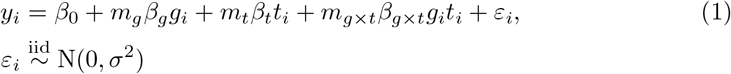

where *i* indexes samples, *y*_*i*_ is the log-transformed molecular count data, *g*_*i*_ is the genotype coded as *{*0, 1, 2*}* or the imputation-based allelic dosage *g*_*i*_ ∈ [0, 2], *t*_*i*_ is an indicator variable for a treatment, ***β*** = (*β*_0_, *β*_*g*_, *β*_*t*_, *β*_*g×t*_)^*T*^ denotes the regression coefficients, and *ε*_*i*_ ∼ N(0, *σ*^2^) is the residual error with error variance *σ*^2^. Each of the eight models can be obtained by substituting elements of the corresponding **m** vector for *m*_*g*_, *m*_*t*_, and *m*_*g×t*_ in, Eq (1). For example, **m** = (0, 0, 0) corresponds to the intercept only model,

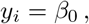

**m** = (1, 1, 0) corresponds to

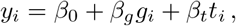

and **m** = (1, 1, 1) corresponds to

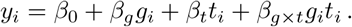

Thus, models corresponding to (0, 0, 1), (1, 0, 1), (0, 1, 1), and (1, 1, 1) include the G×T interaction term, whereas those corresponding to (0, 0, 0), (1, 0, 0), (0, 1, 0), and (1, 1, 0) do not. These model categories are also applicable to nonlinear regression that we propose in this work (Fig 2). It is useful to consider the biological meaning of the eight model categories through the taxonomy and corresponding nomenclature in Figure 2 and Table 1. Nonetheless, we will primarily refer to the models using the **m** vectors in the rest of the manuscript for simplicity.

**Fig 2.**
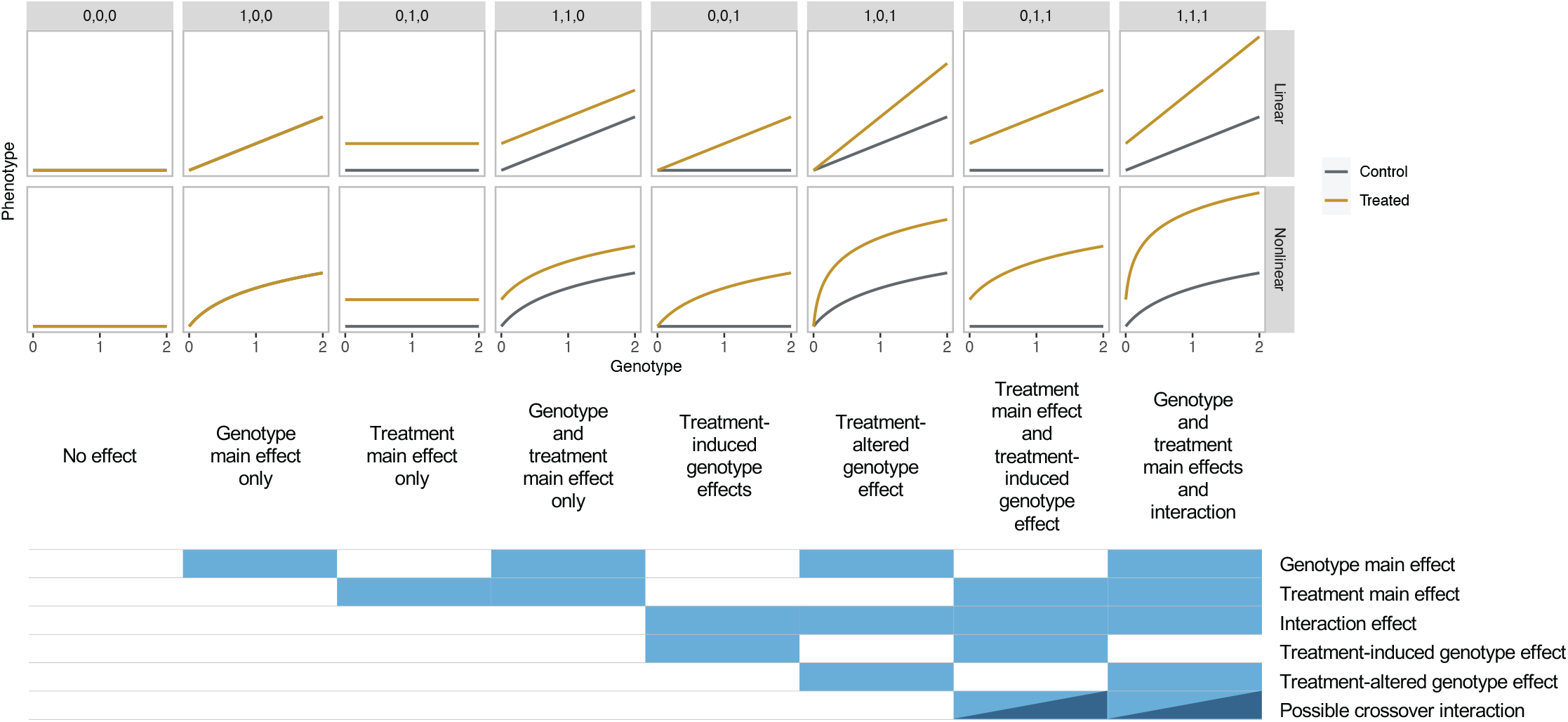
Illustration of types of G×T interactions. Schematics of hypothetical linear regression where the phenotype is regressed on the genotype, coded as 0, 1, and 2, separately in the control (gray) and treated (brown) conditions. Shown on the top are the **m** vectors. Top: Linear models. Bottom: Nonlinear models. Shown below the schematics is the taxonomy of G×T interactions. The dark blue color indicates that a crossover interaction can possibly appear in (0, 1, 1) and (1, 1, 1). See text for details.

**Table 1.** Types of G×T interactions.

| **m** vector | Taxonomic nomenclature | Aggregate category |
| --- | --- | --- |
| (0, 0, 0) | No effect | No $G \times T$ |
| (1, 0, 0) | Genotype main effect only | No $G \times T$ |
| (0, 1, 0) | Treatment main effect only | No $G \times T$ |
| (1, 1, 0) | Genotype and treatment main effects only | No $G \times T$ |
| (0, 0, 1) | Treatment-induced genotype effect | Induced |
| (1, 0, 1) | Treatment-altered genotype effect | Altered |
| (0, 1, 1) | Treatment main effect and treatment-induced genotype effect | Induced |
| (1, 1, 1) | Genotype and treatment main effects and interaction | Altered |

We propose a BMS approach, which consists of four steps, as in Bayesian approaches more generally. First, we specify models that consist of likelihood and prior probabilities (Methods). Second, we compute the posterior probability. Third, we summarize the posterior probability of interest, which is the probability corresponding to different types of G×T interactions in our method. Fourth, we optionally make a decision based on individual or aggregated posterior probabilities. For this decision, the posterior mode is taken based on the 0-1 loss function, ie, we select the model with the highest posterior probability.

We note that three of our models, (1, 0, 1), (0, 1, 1), and (0, 0, 1), contain the interaction term without one or both of the main effect terms, which could be seen as counter to the principle of marginality [22], which forbids such formulations, and in tension with the related principle of heredity [23], which gives them diminished or zero prior probabilities. Our reading of these principles, however, is that they primarily apply to model selection that proceeds sequentially (ie, fitting main effects first, then interactions, not the other way around) or large-scale model search (where ruling out models *a priori* is computationally advantageous), and to modeled data where a zero main effect has no meaningful interpretation. By contrast, our model selection is not sequential, since we compare the full repertoire of possible models simultaneously, we are examining a small space of models, and the models with zero main effects are tied to interpretations that are specific and meaningful in this biological context.

The framework also provides three types of post-processing, which correspond to the third step of BMS described above. First, we compute posterior probabilities of aggregated categories, which can be more relevant than each of the eight models in practical settings. This is straightforward since the eight models are mutually exclusive and the posterior probabilities of aggregated categories can be obtained by summing posterior probabilities of the corresponding models. For example, the posterior probability of models where the G×T interaction is non-zero is computed as the sum of the posterior probabilities of (0, 0, 1), (1, 0, 1), (0, 1, 1), and (1, 1, 1); we refer to this aggregated category as “interaction” and denote it by (∗, ∗, 1). It is also possible and useful to compute the probability of the induced genotype effect, corresponding to (0, ∗, 1), where the association between the genotype and phenotype emerges only upon treatment, as well as the probability of the altered genotype effect, corresponding to (1, ∗, 1), where the genotype-phenotype association exists in the absence of treatment but is the strength of the association is altered upon treatment (Fig 1). Other possible aggregated categories of interest include that of the “restricted” treatment effect, corresponding to (∗, 0, 1), where the treatment only affects individuals with genotype levels 1 and 2, as well as the “varying” treatment effect thereof, corresponding to (∗, 1, 1), where the treatment affects all individuals but to different extents, while we do not specifically consider these further in this manuscript.

The second type of post-processing computes the posterior probability of a crossover interaction, where the treatment has the opposite effects (ie, effects with different signs) depending on the genotype (Fig 1, Methods). The third classifies feature-SNP pairs into 27 groups that correspond to **m** ∈ *{*−1, 0, 1*}*^3^, taking into account the signs of the regression coefficients (Methods).

We are primarily motivated by analyzing datasets generated in *in-vitro* cell systems with and without treatment. In this type of data, samples from both control and treatment conditions are available for the same donor; there may be replicate samples per combination of donor and condition; or replication may be consistently or sporadically absent. To account for a possible error correlation structure within a group of samples derived from the same donor, our framework can include effects of donors and/or genetic relatedness (kinship) between them as a random effect (Methods).

#### Non-BMS approaches to classifying G×T

In this section, we discuss the advantage of BMS over alternative approaches. We first consider an alternative approach that consists of multiple steps and has been used in practice. This first approach performs (molecular) QTL mapping in the control and treated conditions separately as well as response (molecular) QTL mapping. These tests identify a set of feature-SNP pairs for which the null hypothesis is rejected and a set for which the test fails to reject the null hypothesis after multiple testing correction. Subsequently, feature-SNP pairs are classified by taking set differences. For example, feature-SNP pairs corresponding to **m** = (0, 0, 1) and **m** = (0, 1, 1), which represent situations where genetic effects become detectable only in the presence of treatment, can be obtained by taking a set of response (molecular) QTLs for which the genotype-phenotype association is significant in the treated condition but not in the control condition. Those corresponding to **m** = (0, 0, 1) and those corresponding to **m** = (0, 1, 1) can be distinguished by additional hypothesis testing to examine whether the feature is differentially expressed in the control condition. Although conceptually simple, this approach yeilds a single, high variance estimate that lacks any kind of uncertainty quantification—that is, it provides no information about the expected rate of false positive classifications. Moreover, classification based on hypothesis testing is inherently unsatisfactory due to the asymmetry between the null and alternative hypotheses — that is, it only rejects or fails to reject and cannot select the null model. Other possible approaches include significance-guided model selection, such as forward selection and backward elimination [24], which involve sequential hypothesis testing. However, the same caveat applies to these hypothesis testing-based approaches.

An approach more akin to BMS is to fit all the eight models and choose a model using a model selection criterion, such as Akaike’s An Information Criterion (AIC) [25] or the Bayesian information criterion (BIC) [26]. Though this approach is entirely feasible for our purpose due to the small number of models being compared, it also lacks uncertainty quantification, at least without additional enclosing procedures such as resampling [27–29]. For example, consider a scenario where BMS assigns posterior probability of 1.0 to (0, 1, 1) and another scenario where posterior probabilities 0.4 and 0.6 are assigned to (0, 0, 1) and (0, 1, 1), respectively. In either scenario, criterion-based selection would choose (0, 1, 1) with no indication of the fact that the latter scenario implies substantially higher uncertainty.

### Modeling the relationship between molecular phenotype and genotype: transformations and allelic additivity

In the current practice of response molecular QTL mapping, molecular count phenotypes are routinely processed using a variance-stabilizing transformation, such as the log transformation, and the relationship between the genotype and the transformed phenotype is assumed to be linear. Previous studies that have examined this, however, have found that, for the majority of gene-SNP pairs, linearity holds on the original count scale, not on the transformed scale: specifically, that observed data is consistent with contributions from the two alleles combining in an additive manner to the molecular count [17, 18]. This assumption is called *allelic additivity*.

To illustrate the allelic additivity assumption, we consider the effect of a genetic variant in a *cis*-regulatory region on the expression of a target gene. A common assumption is that gene regulatory feedback mechanisms are rare and that gene expression from different alleles is independent. This leads to the assumption that, in a diploid cell, the total gene expression count is the sum of gene expression count from the two alleles and that the gene expression count is linear with respect to the genotype (Fig. 3A). Fig. 3B shows an example of previously published gene eQTL data in hNPCs in the original count scale, which is consistent with this idea. However, after variance-stabilizing transformation to achieve homoscedastic error, the mean expression values are no longer linear with respect to the genotype (Fig. 3C). Similarly, RINT-transformation does not preserve the linear relationship unless there are approximately equal numbers of major and minor allele homozygous donors (Fig 3D). Moreover, with RINT transformation, adjustment for covariates, such as sex, cannot be handled properly, and large effects can be greatly underestimated. A similar argument also applies to molecular count data other than gene expression. Regardless, in practice, linearity is commonly assumed, which can lead to inaccurate inference. Although some previous studies have addressed this issue by accounting for the nonlinearity between transformed molecular count data and genotype [17, 18] for eQTL mapping under a single condition, to our knowledge, this type of nonlinear model has not been used in response QTL mapping. As we shall see, failing to account for this nonlinear relationship can lead to reduced accuracy, and reduced separation of posterior probabilities of correct models from incorrect ones.

**Fig 3.**
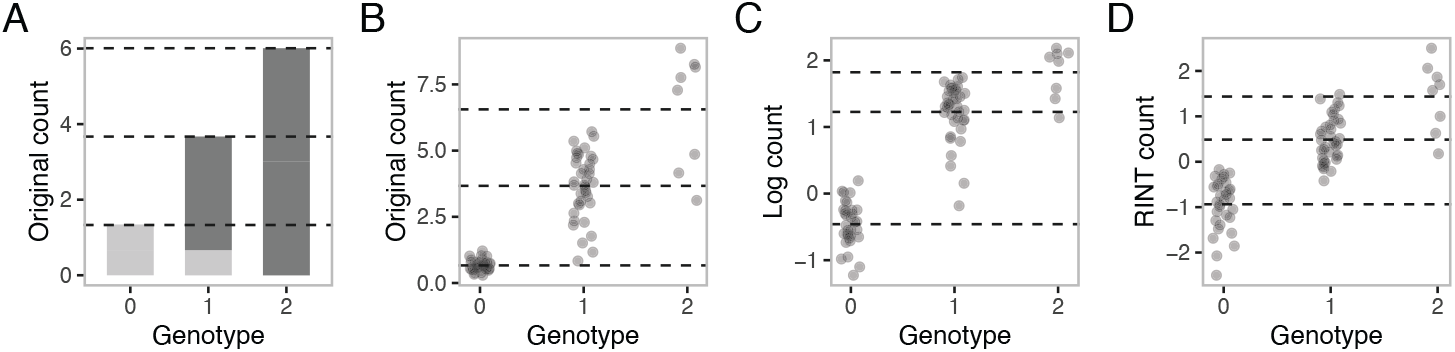
The assumption of allelic additivity. **A**. Barplots illustrating that allelic additivity results in the linear relationship between the phenotype and the genotype. The dark and pale gray boxes represent the hypothetical molecular count signals from the two alleles. **B**. An example of molecular count data on the original scale. Circles represent samples; dashed horizontal lines correspond to the mean expression values conditional on the genotype. **C**. The same as molecular count data in **B** but on the log scale. **D**. The same as the molecular count data in **B** but with RINT transformation.

To account for the nonlinear relationship between the genotype and the transformed molecular count data, we adapted the previously developed methods for single-condition molecular QTL mapping to G×T analysis (Eq (11)). The method, which we call log-NL, explicitly models the nonlinear (NL) relationship between the genotype and the log-transformed molecular count phenotype. W call the commonly used methods, which assume a linear relationship between the genotype and the log-or RINT-transformed phenotype, as log-LM and RINT-LM, respectively, referring to the types of transformation. Throughout the manuscript, we consider comparisons between these three approaches.

### Assessing the allelic additivity in hNPCs with growth stimulation

Previous studies have analyzed gene expression data from genetically diverse donors in the postmortem human tissues and found little evidence to suggest deviations from the allelic additivity [17, 18]. To assess whether the conclusion also holds in *in-vitro* cell systems, we reanalyzed previously published eQTL data in hNPCs, a system in which numerous associations between regulatory features and SNPs have been identified [10, 30–32]. In this analysis, we focused on 3073 feature-SNP pairs with significant G×T pairs (Methods) and fit the log-transformed data to the nonlinear and linear models by maximum likelihood estimation (MLE) for each condition separately (Methods). For comparison, we also fit the ANOVA model with three genotype levels, which is a larger, more flexible model that can describe both nonlinear and linear relationships.

The analysis led to the following observations. First, across the feature-SNP pairs, the maximized likelihood values are higher for the nonlinear model than for the linear model or similar between the models (S1 Fig). Since the number of fitted parameters is identical, this suggests that the nonlinear allelic additivity model is more adequate than the linear model. Second, the values are similar between the nonlinear and ANOVA models, suggesting that the nonlinear model sufficiently captures the complexity of the data with fewer parameters (S1 Fig). Third, the values are lower for the linear model than for the ANOVA model, or they are similar between the models (S1 Fig). Overall, the results show little evidence for deviations from the allelic additivity, consistent with previous studies [17, 18].

We next examined whether the allelic additivity holds for chromatin accessibility, focused on 42576 feature-SNP pairs with significant G×T pairs (Methods), and observed essentially the same trends as for the gene expression data (S1 Fig). To our knowledge, this is the first evidence to suggest that chromatin accessibility signals with respect to the genotype are linear in the original count scale in the majority of cases.

### Simulation: classifying G×T interactions using BMS

We evaluated our G×T classification procedure using simulation. Since we observed little evidence of deviations from allelic additivity in experimental data, we assumed this mechanism to simulate data from the eight models specified in Eq (11) (Methods). The true parameter values were set according to the nonlinear regression results for the response eQTLs in hNPCs (Methods). The data comprised one observation per condition for 80 donors (total sample size *n* = 160). We simulated four scenarios, by generating data with and without a donor random effect, and performing analyses with or without a donor random effect term in the model. For computation, we used both Markov Chain Monte Carlo (MCMC) followed by bridge sampling and maximum a posteriori (MAP) estimation followed by Laplace approximation (Methods and S1 Text). In addition to assessing each of the eight models, we computed posterior probabilities of the aggregated model categories no-G×T, induced, and altered. Within the BMS framework, we compared the performance of the three modeling approaches, log-NL, log-LM, and RINT-LM as follows. First, we examined the distributions of posterior probabilities of the correct models (ie, the data-generating model) as well as those of the incorrect models. Second, we stratified the data based on the correct model and the posterior mode model and inspected the distribution of posterior probabilities in each stratum. Third, we constructed receiver operating characteristics (ROC) curves assessing classification error at different posterior probability thresholds. Fourth, we assessed calibration of the procedure, that is, whether the posterior probabilities inferred for each model match the empirical frequencies with which those models were used to generate data [33].

The analysis omitting the random effect led to the following points. Overall, the three modeling approaches show comparable performance as assessed by ROC curves (Fig 4A, S4 Fig, and S5 Fig) as well as by calibration (S2 Fig and S3 Fig). In the stratified analysis, the posterior mode models matched the correct models for a large fraction of instances (S6 Fig, S7 Fig, S8 Fig, S9 Fig, S10 Fig, and S11 Fig) across all methods. Nonetheless, there was a consistent tendency for log-NL to outperform log-LM and RINT-LM, as assessed by the distribution of the posterior probability of the correct (ie, data-generating) model and incorrect (ie, not data-generating) models thereof (Fig 4B, S16 Fig, and S17 Fig). Specifically, for the aggregated model categories, no-G×T, induced, and altered, log-NL gave higher median posterior probabilities of the correct model as well as lower median posterior probabilities for the incorrect models than other approaches. This was also the case for the individual model categories (S12 Fig, S13 Fig, S14 Fig, and S14 Fig) except for (1, 0, 0) and (1, 1, 0), where the median posterior probability was higher for log-LM than for log-NL. The posterior probability of the correct models may be low for a fraction of the simulation instances, due to the level of error variance chosen for the simulation; still, there exist clear distributional differences between the correct and incorrect models. The performance of BMS for selecting correct models can rather be evaluated by the calibration and ROC curves (Fig 4A, S2 Fig, S3 Fig, S4 Fig, and S5 Fig). Similar results were obtained in other scenarios. Last, these trends did not significantly differ across computational strategies.

**Fig 4.**
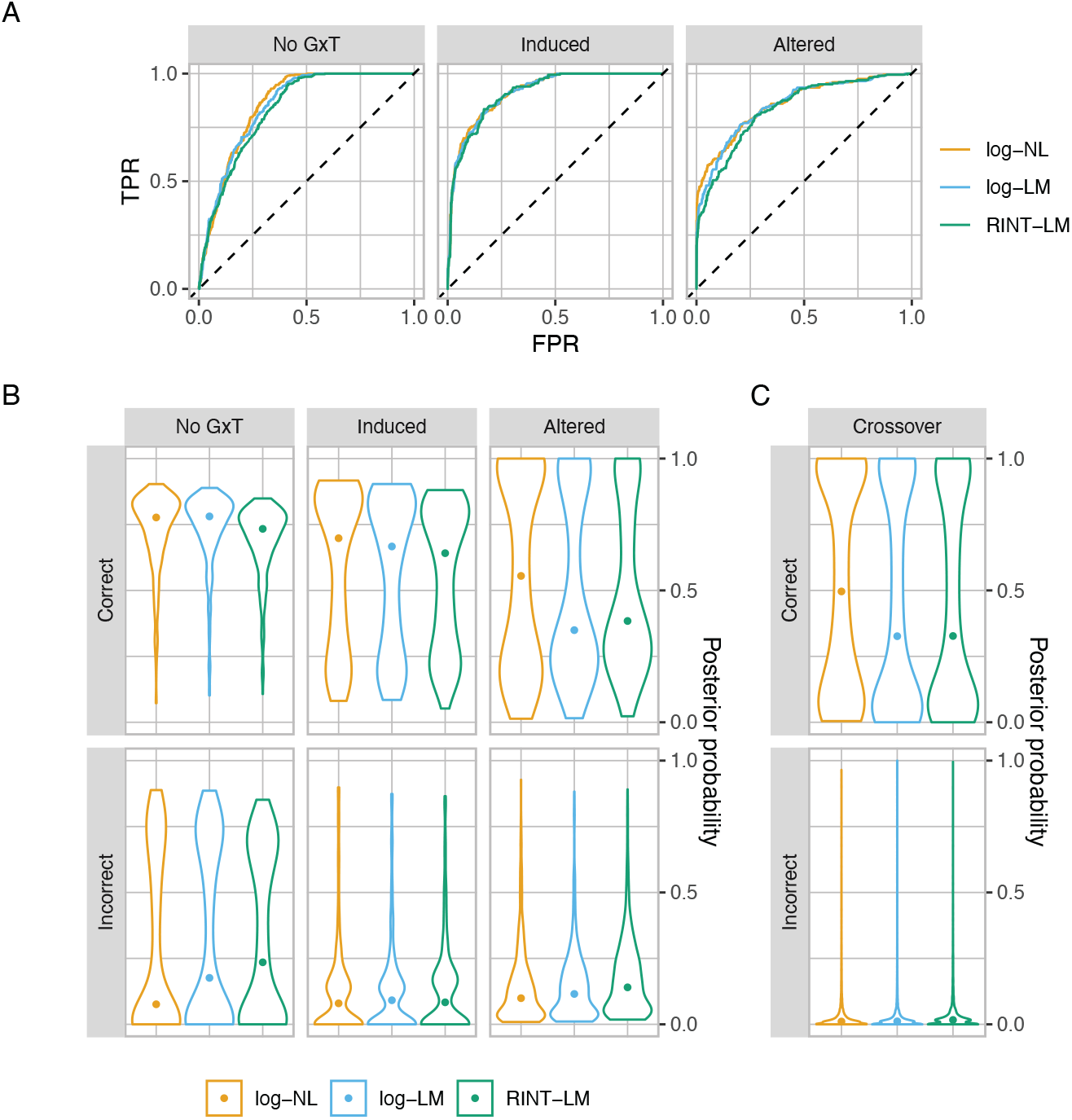
BMS on simulated data. **A**. ROC curves comparing log-NL, log-LM, and RINT-LM for the no-G×T, induced, and altered model categories. **B**. Violin plots showing distributions of the posterior probability of the correct (top) and incorrect (bottom) models for log-NL, log-LM, and RINT-LM. Shown in the top of each panel is the model category. **C**. The same as **A** but for crossover interactions.

### Identifying crossover interactions through post-processing

In practice, one situation of interest is where the G×T interaction results in the treatment effect acting in the opposite direction depending on the genotype (crossover interaction). Probabilistic characterization of such an event is only possible and straightforward with BMS. To identify feature-SNP pairs of this type, we performed post-processing to compute the posterior probability of crossover interaction for the 8000 datasets generated from the nonlinear allelic additivity model (Methods). The ground truth was obtained by examining the simulated effects of the homozygous genotype (*g* = 0 or *g* = 2) under alternate treatments and prior to the addition of noise. The posterior probabilities of crossover interaction were then obtained from log-NL, log-LM, and RINT-LM models for both truly crossover and non-crossover cases. We observed that log-NL gave higher median posterior probability than log-LM and RINT-LM in cases where there was a true crossover interaction, and lower median posterior probability when there was not (Fig 4C).

Overall, the results from the simulation analysis suggest strong performance of our BMS framework as well as advantages of log-NL over log-LM and RINT-LM.

### Classifying G×T interactions for response eQTLs in hNPCs with growth stimulation

As an example application of our BMS framework, we computed posterior probabilities for the types of G×T interactions for a set of response eQTLs identified in hNPCs by Matoba *et al*. [10]. In particular, we focused on 98 response eQTLs on autosomes with the CHIR treatment (Methods). The response eQTLs represent pairs of genes and index SNPs for which the G×T interaction term was significantly non-zero based on hypothesis testing. CHIR, also known as CHIR99021, is an activator of the canonical Wnt pathway, which has been implicated in proliferation of hNPCs, cortical patterning, and complex brain traits [34–39]. As in the simulation experiments, we compared the three modeling approaches (log-NL, log-LM, RINT-LM), but using mixed effect models to account for the genetic relatedness between the donors (Methods).

Overall, the analysis led to the following two points. First, the BMS results were largely concordant with the previous response QTL mapping results at the level of calling significant interactions of genotype and treatment (S28 Fig). Specifically, of the 98 previously identified response eQTLs, G×T models achieved the highest posterior probabilities in 94 eQTL analyzed by log-NL, with 95 for log-LM, and 92 for RINT-LM. The small discrepancy is likely due to the difference in the data preprocessing (Methods). Second, the three approaches gave comparable but not identical classification and inference. Specifically, for the 90 out of 98 feature-SNP pairs, the posterior mode models were identical between approaches. Examples of feature-SNP pairs with varying degrees of concordance from the three modeling approaches are shown in Fig 5. For the long intergenic non-protein coding RNA LINC02073 and its neighboring SNP, rs7212610, the highest posterior probability was given to (1, 1, 1) by both log-NL and log-LM, but not by RINT-LM (Fig 5A and B). Furthermore, log-NL captures the reduction of the genotype effect size upon treatment with higher certainty than log-LM, likely due to its ability to account for the nonlinear relationship between the genotype and phenotype. The result is consistent with the idea that accuracy in effect size estimation can have a nonnegligible impact on the detection of G×T interactions. For the *SLC35F3* gene encoding a thiamine transporter and its neighboring SNP, rs650866, all three modeling approaches assigned the highest and second highest posterior probabilities to (1, 0, 1) and (1, 1, 1), respectively. However, log-NL gave larger certainty on the posterior mode model than log-LM and RINT-LM did. A possible explanation for this observation is that the inadequate linear constraint of log-LM as well as the nonparametric transformation in the RINT-LM approach led to an increased probability of an incorrect inference of the treatment effect for the major allele homozygote (Fig 5C and D).

**Fig 5.**
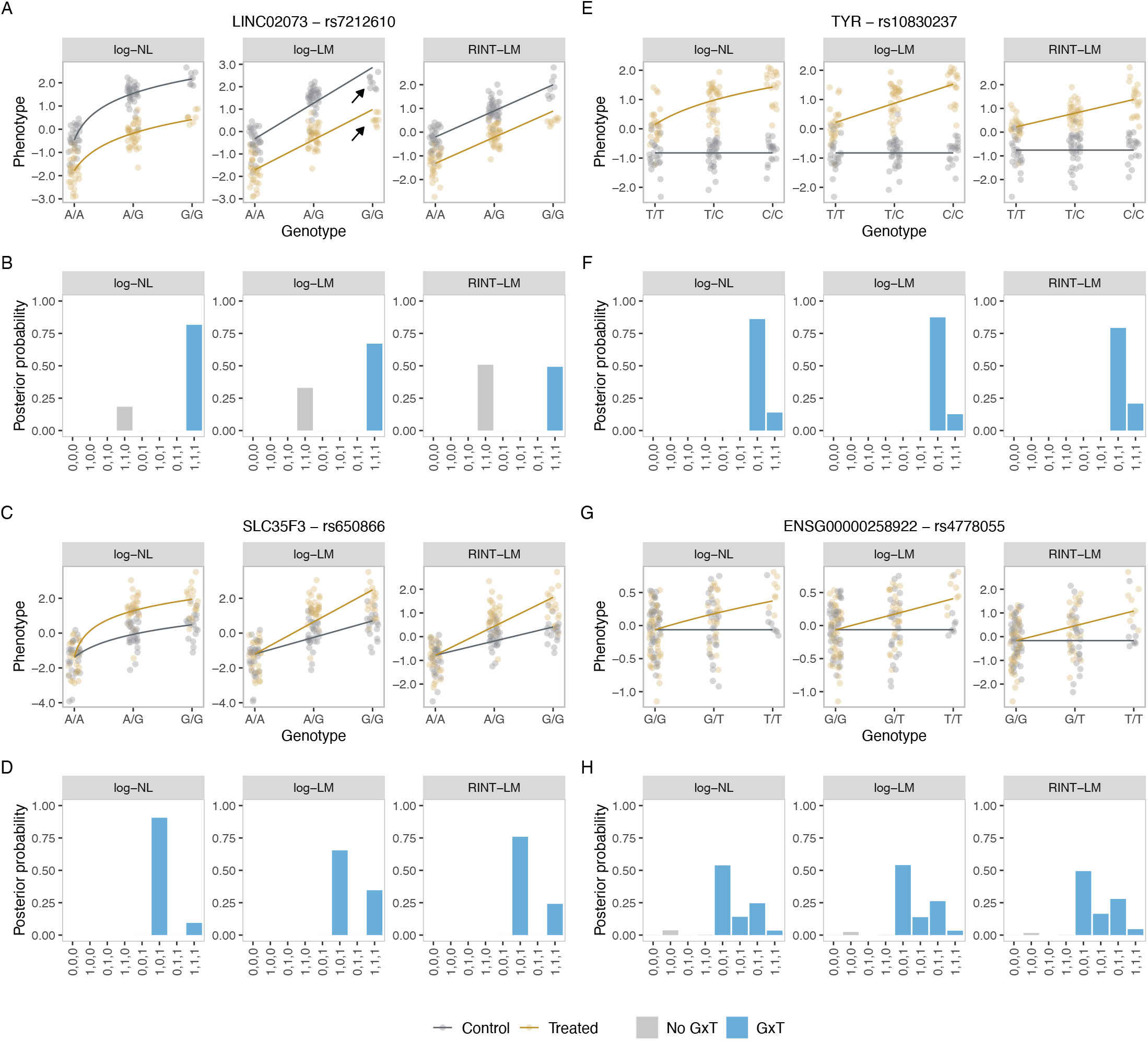
Representative BMS results for the response eQTL in hNPCs. The upper panels show fitted functional relationships between the genotype and the transformed molecular count phenotype based on the posterior mode model. The circles represent the data. The gray and brown colors represent the control and treated conditions, respectively. The arrows indicate deviations from regression lines (**A, C, E, G**). The lower panels show posterior probability of the different types of G×T interactions. The gray and blue colors represent the probability of the no-G×T and “interaction” categories, respectively. (**B, D, F, H**). Within each subfigure, the left, middle, and right panels show results from log-NL, log-LM, and RINT-LM, respectively.

Indeed, the mean phenotype values for the major allele homozygote were -1.45 and -1.39 with the standard errors of 0.97 and 0.87 for the control and treated conditions, respectively, which did not differ significantly (*P* = 0.68 from a paired *t*-test).

We also computed aggregated probabilities of interest. For 16 and 75 feature-SNP pairs, the results from the three modeling approaches agreed in assigning the highest posterior probability to the induced and altered genotype effect patterns, respectively, among the three aggregated categories (no-G×T, induced, and altered). The TYR gene encoding a tyrosinase and its neighboring SNP, rs10830237, as well as the novel long non-coding RNA gene ENSG00000258922 and its neighboring SNP, rs4778055, provide examples of the induced category (Fig 5E, F, G and H).

Overall, the analysis illustrates the utility of our framework in that it provides interpretable posterior probabilities rather than hard classifications without uncertainty quantification. It also suggests potential advantages in employing the log-NL approach for response QTL. We note that similar results were obtained using mixed effect models to account for possible error correlation structures between samples derived from the same donor (donor random effect) rather than polygenic (kinship) effect (Methods and S32 Fig).

### Identifying crossover interactions in the hNPC data

To identify gene-SNP pairs for which the treatment has the opposite effect depending on the genotype, we performed post-processing of the log-NL results to compute the posterior probability of crossover interaction for the previously identified 98 response eQTLs (Methods). Examples with high posterior probability of crossover interactions are shown in Fig 6. Interestingly, for the *ZNHIT3* gene encoding the zinc finger HIT domain-containing protein 3, which is known to be defective in a severe encephalopathy, and a neighboring SNP, rs4796224, the posterior probability of crossover interaction was close to one (Fig 6A and B). In this case, the model category (1, 1, 1) solely received non-zero probability. By contrast, for the ENSG00000287315 gene encoding a novel antisense transcript and a neighboring SNP, rs10157612, the model categories (1, 1, 0) and (1, 1, 1) both received non-zero posterior probability (Fig 6C and D). For the *TLCD4* gene encoding the TLC domain-containing protein 4 and a neighboring SNP, rs7556223, log-NL, log-LM, the posterior probability of a crossover interaction was lower than for other examples, representing the uncertainty (Fig 6E and F). We emphasize that this utility is uniquely provided by BMS and presents its additional advantage.

**Fig 6.**
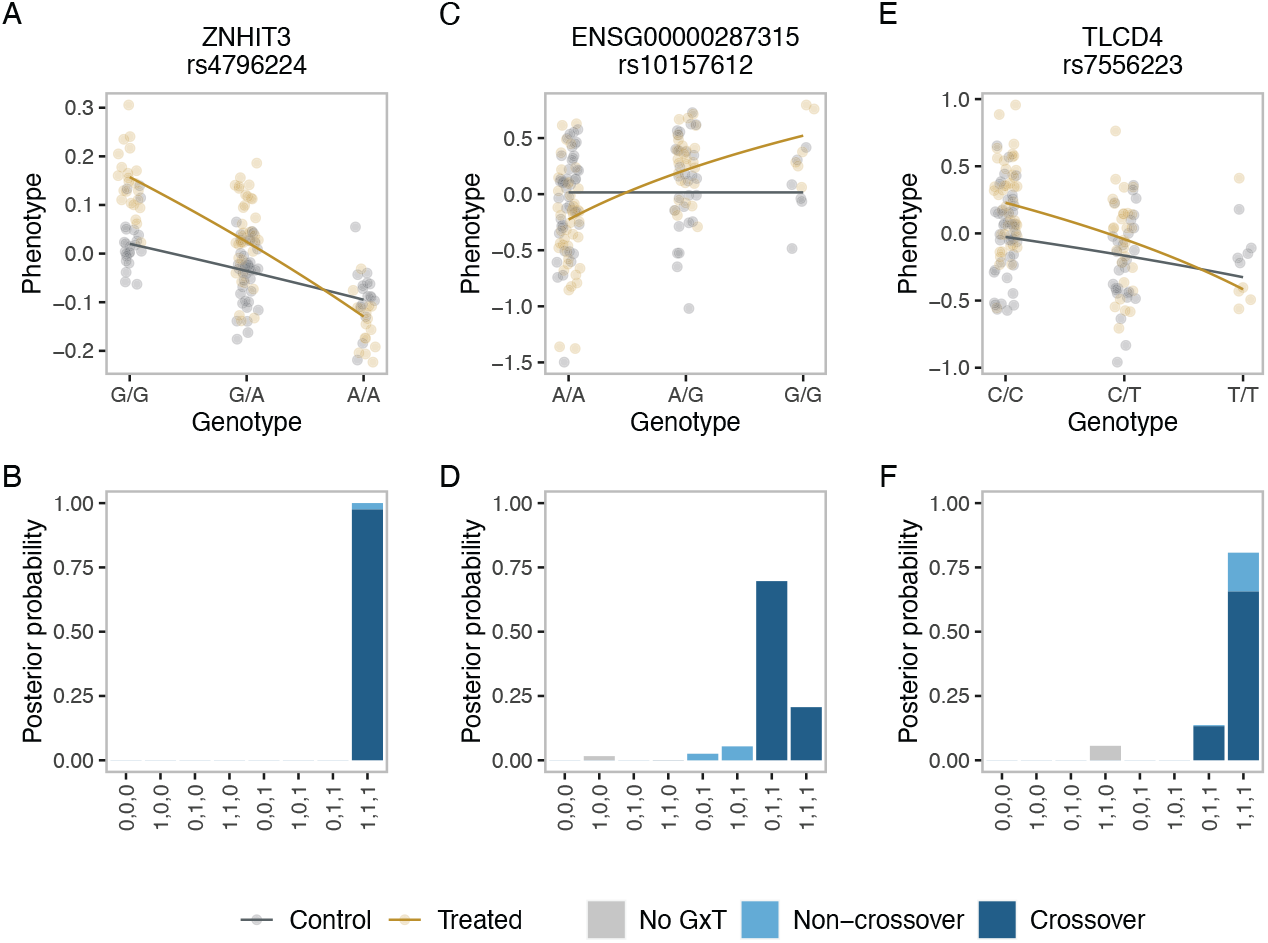
Examples of response eQTLs with the crossover interaction in hNPCs. The upper panels show fitted functional relationships between the genotype and the transformed molecular count phenotype based on the posterior mode model obtained by BMS with log-NL. The circles represent the data. The gray and brown colors represent the control and treated conditions, respectively (**A, C, E**). The lower panels show posterior probability of the different types of G×T interactions obtained by BMS with log-NL. The gray color represents the probability of the no-G×T category. The pale and dark blue colors represent the probability of the “non-crossover” and crossover interactions, respectively (**B, D, F**).

In practice, the distinction between positive and negative effects can be crucial for the interpretation and prioritization of response molecular QTLs. Post-processing of the output from our BMS framework can generate posterior probability of 27 categories that correspond to **m** ∈ *{*−1, 0, 1*}*^3^, accounting for the sign of the effect sizes. The results of such analysis are summarized as heatmaps in S28 Fig.

### Classifying G×T interactions with respect to chromatin accessibility

To illustrate the usability of our framework on a molecular phenotype other than gene expression, we reanalyzed 1775 autosomal response caQTLs identified previously with the CHIR treatment [10]. Analogously to the analysis of the response eQTLs, this analysis led to the following observations. First, the BMS results are reasonably concordant with the response caQTL results at the level of calling significant interactions of genotype and treatment (S29 Fig). Specifically, for 1588, 1588, and 1523 of the 1775 feature-SNP pairs, the log-NL, log-LM, and RINT-LM approaches, respectively, assigned the highest posterior probability to one of the G×T interaction models. The three approaches gave comparable but not identical classification and inference. For the 1516 feature-SNP pars, the posterior mode models were identical between them. However, we observed a number of examples where the posterior probability patterns substantially differed between the approaches yet gave the same highest posterior probability model (S30 Fig). Among the three aggregated categories, no-G×T, induced, and altered, for 538 and 848 feature-SNP pairs, the results from the three modeling approaches agreed in assigning the highest posterior probability to the induced and altered effect patterns, respectively. An example of the induced effect pattern is shown in S30 Fig. We also observed a number of feature-SNP pairs with the crossover interaction. Examples are shown in S31 Fig. The classification results accounting for the sign of the effect size are summarized as heatmaps in S29 Fig.

## Discussion

We have developed a method to help interpret and prioritize response molecular QTLs. The method uses BMS to assign posterior probability to different types of G×T interactions. Within this framework, we compared three different modeling approaches. The first approach, log-NL, assumes that molecular signals are additive with respect to allelic counts and explicitly models the nonlinear relationship between the genotype and the molecular counts after the log transformation. This approach is justified based on previous studies as well as our analysis examining the adequacy of the model using experimental data [16–18]. The second approach, log-LM, assumes a linear relationship between the genotype and the log-transformed molecular phenotype. Though commonly applied, this approach can lead to nontrivial model misspecification. The third approach, RINT-LM, assumes a linear relationship between the genotype and the molecular phenotype that has been transformed using a rank-based method. In our simulation experiments with realistic effect and sample sizes for *in-vitro* cell systems, log-NL outperformed other approaches moderately but consistently. Our analysis of previously published experimental data illustrates the utility of our framework in extracting practically relevant information from response molecular QTL mapping data. Individual inspections of the experimental data and fitted models revealed cases where the failure of linear regression of log-transformed counts in capturing the data characteristics led to posterior probability patterns that differ from those obtained by nonlinear regression. These observations collectively suggest the benefit of nonlinear regression over the commonly used linear regression.

A variety of statistical methods have been developed to identify G×T and G×E interactions [1]. Earlier studies used the “stratified” approach, which focuses on the set difference in significant associations between the control and treated conditions [2, 3]. Despite being methodologically simple, this approach is limited in that an uncertainty measure is not readily available for a set difference and that it cannot detect subtle interactions where the association between the phenotype and the genotype is significant in both conditions but to different extents. Moreover, there is no sharing of information between the conditions even if the data is collected from identical or overlapping sets of individuals. Another method is the “delta” approach, where the outcome variable is the difference in the phenotype between the control and treated conditions [2, 4–6]. Though this approach can be better powered than other approaches in some situations, it removes the information regarding the genotype-phenotype association in individual conditions. Moreover, for a given genotype, it requires phenotypic measurements in both control and treated conditions and, thus, is not applicable to a setting, such as that of clinical trials, where an individual of a given genotype either receives or does not receive treatment. One of the most commonly used methods is the “interaction” approach, which is often called response QTL mapping, where phenotype data in the control and treated conditions is jointly modeled using a linear model including the genotype, treatment, and G×T interaction terms [7–11]. This approach does not require paired data and potentially exhibits increased power, the latter of which may result from joint modeling. Our method is built on this type of approach.

The proposed BMS approach is distinct from the large majority of existing methods of G×T and G×E analyses in that the goal is to classify different types of interactions, whereas existing methods focus on detection with some notable exceptions, including those developed by Barber *et al*. [14] and Maranville *et al*. [15]. The pattern of posterior probabilities of G×T interaction resulting from our approach provides different strengths of evidence for different G×T interactions, which are intuitive and interpretable. It is straightforward to extract practically relevant information through post-processing. This includes the posterior probability of an event where the genotype effect is induced or altered by treatment, an event where the treatment effect is restricted to a specific genotype or varies depending on the genotype, and an event where the treatment has the opposite effect (i.e., effects with different signs) depending on the genotype. Moreover, it is possible to obtain probabilities of association patterns with sign constraints of interest. The feature-SNP pairs can then be prioritized based on the probability of specific association patterns of interest, for example, cases of enhancer priming where activation of co-factors is required to induce the transcriptional effect of an enhancer on a gene promoter [9]. This can facilitate designing downstream experimental and bioinformatics analyses, such as feature enrichment analyses to gain further biological insights (e.g., gene set enrichment).

Our method differs from the existing probabilistic approach developed by Maranville *et al*. [15], which was an extension and improvement of earlier work by Barber *et al*. [14], at least in the following aspects. First, in our model specification, the control and treated conditions are asymmetric in that the former is considered basal, unlike in the bivariate outcome model used in the existing method [15]. This specification is based on the assumption that treatment with stimuli tends to induce a genetically regulated response rather than suppress it, as exemplified by a previous study in immune cells [9]. Nonetheless, in a situation where generic variation is expected to be suppressed, the coding of the treatment indicator variable can be reversed. Second, our method does not require paired data and, thus, can accommodate a wider range of settings. Third, our model assumes equal variances for residual errors on the log scale between the two conditions, whereas the existing method allows heterogeneous variances. That said, in principle, our model could be trivially modified to accommodate such additional heterogeneity. Fourth, we account for the possible error correlation structure between samples derived from the same donor by including random effects, whereas the existing method [15] accommodates the correlation structure by assuming that the errors follow a bivariate Normal distribution with a non-diagonal covariance matrix. Last and importantly, our method accounts for the inherent relationship between the genotype and molecular count phenotype using nonlinear regression while accounting for heteroskedasticity.

Prior specification is a key consideration in Bayesian analysis. BMS requires prior on the candidate models (model prior) as well as on the model parameters (effect prior) (Methods). In BMS, as in other Bayesian analyses that involve computation of the marginal likelihood, the effect prior requires proper calibration. Our extensive simulation analysis (S1 Text) shows moderate prior sensitivity (S18 Fig to S25 Fig), suggesting that data-driven prior elicitation, such as our use of empirical Bayes, can be desirable. Although our current implementation uses a grid search, this process can be made more efficient by employing other optimization techniques, such as a coordinate ascent algorithm [40]. For our model prior, we place a uniform prior as in previous studies using BMS for similar purposes [41], and we consider this an easily interpretable default for our setting. Nonetheless, other lines of research using BMS suggest advantages of model priors that are data-driven [42, 43]. Determining how best model prior is elicited for our BMS framework is an important future direction.

Although our focus here was on molecular count phenotypes, our method can be applied to a broader range of phenotypes. In particular, certain continuous phenotypes can be adequately modeled using linear regression as an approximation (Eq (5), (12)).

For this class of phenotypes, G×T interactions, or more broadly, G×E interactions, can be efficiently classified if random effects need not be modeled. BMS can then take advantage of the exact analytical form of the marginal likelihood, which is available for the linear models with a particular prior specification [44]. Unlike the method we proposed in the current work, this method does not require sampling or optimization and, thus, provides superior computational efficiency albeit with less flexibility.

## Conclusion

We have developed a statistical framework for classifying G×T interactions for molecular phenotypes that facilitates the interpretation and prioritization of response molecular QTLs. Our method takes a set of response molecular QTLs identified by a standard method and assigns posterior probabilities to different types of G×T interactions using a BMS approach. In our simulation experiments, we compared linear and non-linear regression of log-scale counts and observed moderate but consistent performance advantage of the latter over the former. We then applied our framework to experimental data generated in *in-vitro* cell system derived from genetically diverse donors with and without growth stimulation. Although both linear and non-linear regression approaches were successful in recovering the G×T signals, we observed individual examples where the latter captured the data more adequately than the former and the two approaches resulted in different posterior probability patterns. Our method revealed different strengths of evidence for different types of G×T interactions across feature-SNP pairs. This type of information is not provided by existing methods for analyzing response molecular QTLs and can be effective for the interpretation and prioritization of genetic variants underlying the diversity in treatment response among individuals.

## Methods

### Datasets and preprocessing

Primary human neural progenitor cell (hNPC) genotypes, RNA-seq, and ATAC-seq data were obtained from a previous study [10] and were based on the GRCh38 genome assembly. The genotype data was coded as *{*0, 1, 2*}* to represent the number of minor (alternative) alleles and contained 78 and 72 donors for the RNA- and ATAC-seq data, respectively. Among the previously generated data with multiple treatments, we focused on the vehicle and CHIR treatments as control and treated conditions, respectively. The dataset contained one sample per combination of donor and treatment condition. That is, the donors were shared in both control and treated conditions, and there was no missingness.

For assessing the allelic additivity assumption, we considered a set of feature-SNP pairs for which a significant association between the genotype and the phenotype was identified at least in one of the control and treated conditions by the previous study [10]. We excluded feature-SNP pairs for which the number of donors at any genotype level is zero since such data is not suited to the ANOVA that assumes three genotype levels (Eq (7)). For eQTL and caQTL data, the filtering resulted in 3073 and 42576 feature-SNP pairs, respectively. For Bayesian model selection (BMS), we considered 98 response eQTLs as well as 1775 response caQTLs, identified by the previous study [10]. For both types of analysis, we only included SNPs on autosomes.

In the previous study, the count data was transformed using a variance stabilizing transformation other than the log transformation [10]. To achieve compatibility with the nonlinear model that assumes the log transformation (Eq (11)), we reprocessed the raw count data according to previous studies on the allelic additivity [17, 18]. Briefly, we first scaled the original count data as

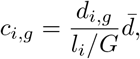

where *i* = 1, …, *n* indexes the samples, *g* = 1, …, *G* indexes the features, such as genes and ATAC-seq peak regions, *d*_*i,g*_ denote elements of the original count matrix, 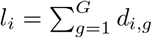be the library size for sample *i*, and 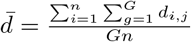 is the overall mean. We note that the experiments have similar sequencing depth and, thus, that the variance of the scaled counts is similar to that of the original counts. For a given gene *g*, we transformed *c*_*i*_ as

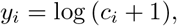

where *y*_*i*_ is the scaled, log-transformed count data. Note that the subscript *g* is omitted for simplicity. The total numbers of genes and ATAC peak regions were 22354 and 172887, respectively. To aid comparisons between competing methods, we additionally transform count data 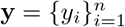 using the rank inverse normal transformation (RINT),

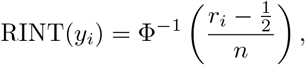

where *r*_*i*_ is the rank order of *y*_*i*_ in **y**, and Φ is the cumulative density function of the standard Normal distribution. For response molecular QTL mapping and BMS, we applied the transformation to the phenotype data residualized with respect to confounding variables (see **Including covariates** in **Methods**).

### Assessing the allelic additivity assumption in a given condition

For assessing the allelic additivity assumption in a condition-specific manner, we computed the maximum likelihood estimates (MLE) of the model parameters based on the previously developed nonlinear model [17, 18] (Eq (2), (3), (4)) and compared the values with those obtained from a standard linear model (Eq (5)) as well as those from an ANOVA model (Eq (7)). In our notation, for a given condition and the *i*-th donor (*i* = 1, …, *n*), the nonlinear model representing the allelic additivity can be written as

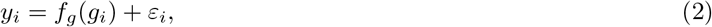

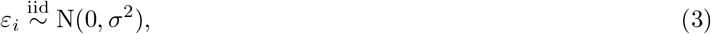

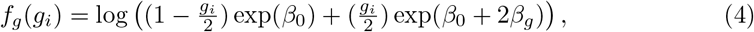

where *n* is the number of donors, *y*_*i*_ is the log-transformed molecular count data, *g*_*i*_ is the genotype coded as *{*0, 1, 2*}* or the imputation-based allelic dosage *g*_*i*_ ∈ [0, 2], *ε*_*i*_ is the residual error, and *σ*^2^ is the residual error variance, and *β*_0_ is the intercept. The parameter *β*_*g*_ represents the half the difference in the expected values of *y*_*i*_ for the major allele homozygote vs the minor allele homozygote. That is, the quantities can be defined as

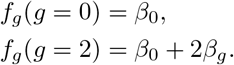

Our definitions and interpretations of the coefficients differ from those in the previous studies to make the scales comparable to those in the standard linear model for QTL mapping, which can be cast as

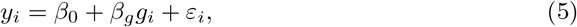

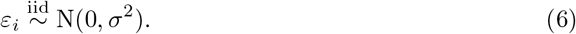

The ANOVA model is cast as

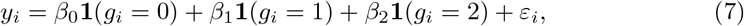

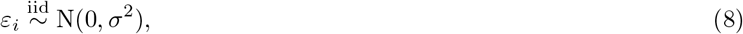

where **1**(*·*) is an indicator function, and parameters *β*_0_, *β*_1_, and *β*_2_ are, respectively, the expected values of *y*_*i*_ for donors with the homozygous major allele, the heterozygote, and the homozygous minor allele. Note that, for a given condition, the hNPC data contained one measurement per donor and that, for the ease of exposition, confounding factors are omitted (for model formulation with confounding factors, see **Including covariates**).

### Extending the allelic additivity model for G×T analysis

The nonlinear allelic additivity model above can be modified for G×T analysis as follows. For the *i*-th sample (*i* = 1, …, *n*), we model

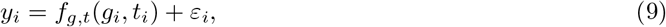

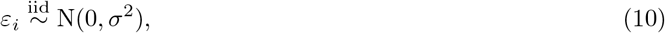

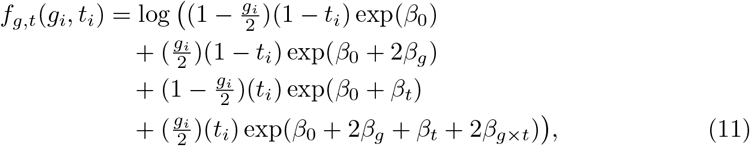

where *y*_*i*_, *g*_*i*_, *ε*_*i*_, *σ*^2^, and *β*_0_ are defined as earlier, *t*_*i*_ denotes an indicator variable for a treatment, *β*_*g*_ now represents the half the difference in the expected values of *y*_*i*_ for the major allele homozygote vs the minor allele homozygote *under the control condition, β*_*t*_ is the difference in the expected values of *y*_*i*_ in the control condition vs the treated condition for the minor allele homozygote, and *β*_*g×t*_ allows the effect of genotype to vary between treatment groups. That is, the quantities can be defined as

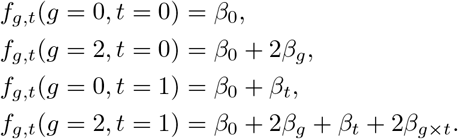

Our definitions and interpretations of the coefficients differ from those in the previous studies [17, 18] to make the scales comparable to those in the standard linear model for G×T analysis, which can be cast as

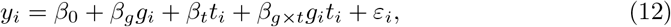

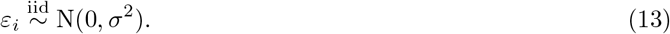

Note that, for the ease of exposition, confounding factors are omitted (for model formulation with confounding factors, see **Including covariates**).

### Bayesian model selection

To classify the types of G×T interactions, we propose a Bayesian model selection approach considering eight models where the *β*_*g*_, *β*_*t*_, and *β*_*g×t*_ parameters are either zero or non-zero in Eq (11). The approach consists of four steps as in Bayesian approaches more generally. First, we specify the likelihood and prior probabilities. Second, we compute the posterior probabilities. Third, we summarize the posterior probabilities of interest. Fourth, optionally, we make a decision based on a loss function.

Using a vector of indicators, **m** = (*m*_*g*_, *m*_*t*_, *m*_*g×t*_) ∈ *ℳ* = *{*0, 1*}*^3^, that specifies one of the eight candidate models in the set *ℳ*, we cast the model as

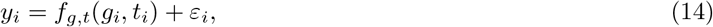

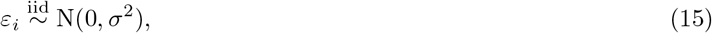

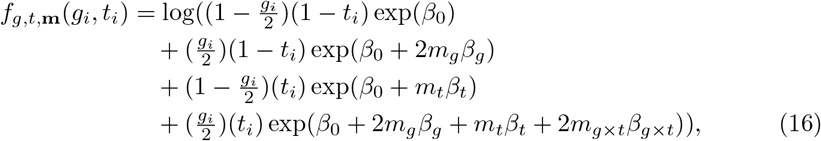

where *y*_*i*_, *g*_*i*_, *t*_*i*_, *ε*_*i*_, and *σ*^2^ are defined as earlier and ***β*** = (*β*_0_, *β*_*g*_, *β*_*t*_, *β*_*g×t*_)^*T*^ denotes the regression coefficients. For concreteness, we write

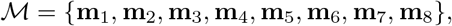

Where

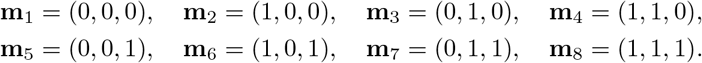

Using *j* to index the models, we have

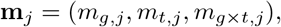

for *j* = 1, …, 8. The conditional joint likelihood is

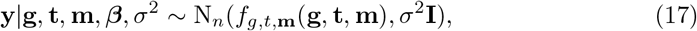

where **y** ∈ ℝ^*n*^ denotes a vector of log-transformed molecular count data, **g** ∈ ℝ^*n*^ denotes a vector of genotypes, **t** ∈ ℝ^*n*^ denotes a vector of treatment indicators, ***β*** = (*β*_0_, *β*_*g*_, *β*_*t*_, *β*_*g×t*_)^*T*^ and *f*_*g,t*,**m**_(**g, t, m**) = (*f*_*g,t*,**m**_(*g*_1_, *t*_1_), …, *f*_*g,t*,**m**_(*g*_*n*_, *t*_*n*_))^*T*^ ∈ ℝ^*n*^ denotes a vector of mean values in the untransformed scale. In an exact mathematical form, Eq (17) corresponds to

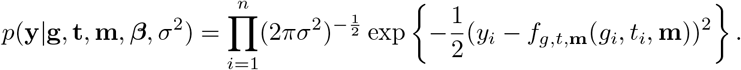

A BMS with linear regression can be formulated by replacing Eq (14) with

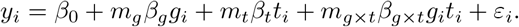

In BMS, it is necessary to specify a prior probability that each of the model is correct, Pr(**m** = **m**_*j*_), and prior distributions for the non-zero parameters in the models, *β*_*g*_, *β*_*t*_, and *β*_*g×t*_. We call the former and latter model and effect priors, respectively. For the model prior, by default, we place the uniform prior probability across the eight models, which corresponds to

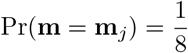

for *j* = 1, …, 8, where *j* indexes the models. This choice is reasonably justified since we apply BMS to a pre-selected set of feature-SNP pairs that are likely to have significant associations. For the effect prior, we use the Normal-Gamma prior, which is commonly used due to its conjugacy for linear models. That is, we set

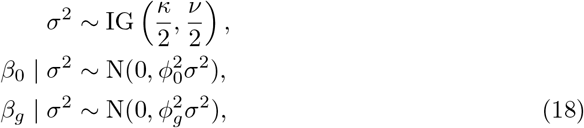

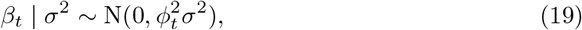

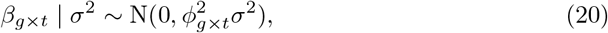

where *κ*, 𝒱, *ϕ*_0_ and ***ϕ*** = (*ϕ*_*g*_, *ϕ*_*t*_, *ϕ*_*g×t*_)^*T*^ are the prior hyperparameters. For the residual variance *σ*^2^ and the intercept *β*_0_, nearly non-informative priors are used by setting 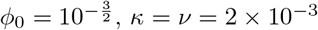The hyperparameters ***ϕ*** = (*ϕ, ϕ, ϕ*) control the effect sizes relative to the residual error standard deviation and need to be properly calibrated for the computation of the marginal likelihood. We employ an empirical Bayes approach where the hyperparamer values maximizing the sum of the marginal likelihood are sought by a grid search.

In our BMS procedure, the second step corresponds to computing the posterior probabilities of the model, Pr(**m** = **m**_*j*_ | **y**) ∝ *p*(**y** | **m** = **m**_*j*_) Pr(**m** = **m**_*j*_), which is proportional to the product of the marginal likelihood conditional on the model, *p*(**y** | **m**), and the prior for the models, Pr(**m** = **m**_*j*_). The marginal likelihood is defined

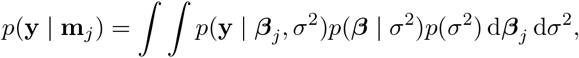

for *j* = 1, …, 8. ***β***^*j*^ is a vector consisting of the non-zero coefficients. For example, ***β***^4^ = (*β*_0_, *β*_*g*_, *β*_*t*_)^*T*^ and ***β***^8^ = (*β*_0_, *β*_*g*_, *β*_*t*_, *β*_*g×t*_)^*T*^. The posterior probability of the *j*th model is computed as

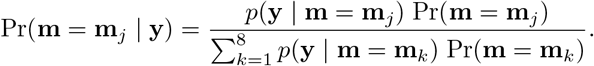

This involves fitting each of the eight models separately. Since the posterior distributions of the parameters and the marginal likelihood, *p*(**y** | **m** = **m**_*j*_), are intractable, we obtain approximate values by either of two methods. In the first method, we fit the models using the Markov Chain Monte Carlo (MCMC) method as implemented in the R package rstan [45] and compute the marginal likelihood using bridge sampling [46]. In the second method, we obtain a maximum *a posteriori* (MAP) estimate using optimization and compute the marginal likelihood using Laplace’s method [44, 47] (see **Computing the marginal likelihood by Laplace approximation** for details). After computing the posterior probabilities of the eight models, in the third step of our BMS procedure, we summarize them in a way that is practically informative. We describe this step in detail in **Refining posterior inference through post-processing**. For the optional fourth step in BMS, we make a decision using a 0-1 loss function, which corresponds to selecting the model with the highest posterior probability Pr(**m** = **m**_*j*_ | **y**) (a maximum *a posteriori*, MAP, estimation).

In principle, this procedure is equivalent to fitting a Bayesian variable selection regression (BVSR) model where spike-and-slab priors are placed on the regression coefficients and all possible models are simultaneously considered in a sampling process [48]. For this type of model, sampling can be trapped in one model, resulting in poor mixing. By contrast, in our approach, all models are separately fit, which is possible since only a small number of models are considered.

### Conditional and model averaged effect posteriors

It is straightforward to obtain model-averaged posterior distributions of the effect sizes from the corresponding conditional distributions [49] as

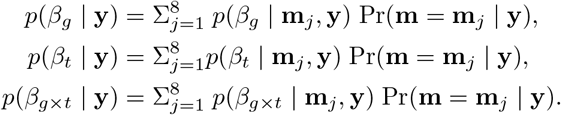

We do not, however, present results from this type of analysis in this manuscript since it is not a main feature of our framework.

### Refining posterior inference through post-processing

In the third step of our BMS procedure, we summarize the posterior probability of interest.

#### Aggregating model categories

One approach is to compute posterior probability of an aggregated category, which can be of practical relevance. This is possible for either of the approaches to compute posterior probabilities of the models, namely, MCMC followed by bridge sampling and MAP estimation followed by Laplace approximation (see **Bayesian model selection** above). Since the eight models are mutually exclusive, posterior probabilities of aggregated categories can be obtained by summing posterior probabilities of the corresponding models. For example, the posterior probability of models where the genotype coefficient is non-zero is computed as the sum of the probability of (1, 0, 0), (1, 1, 0), (1, 0, 1), and (1, 1, 1). We denote this aggregated category by (1, ∗, ∗). Likewise, a category of models with non-zero G×T interaction can be written as (∗, ∗, 1), which we refer to the “interaction” effect category. Additionally, the following aggregated categories can be considered. First, the induced genotype effect category, corresponding to (0, ∗, 1), represents an event where the association between the genotype and phenotype emerges only upon treatment. Second, the altered genotype effect, corresponding to (1, ∗, 1), represents an event where the genotype-phenotype association exists in the absence of treatment but is the strength of the association is altered upon treatment. Third, the “restricted” treatment effect, corresponding to (∗, 0, 1), represents an event where the treatment only affects individuals with genotype levels 1 and 2. Fourth, the “varying” treatment effect, corresponding to (∗, 1, 1), represents an event where the treatment affects all individuals but to different extents. In practical settings, the “interaction”, induced, altered, “restricted”, and “varying” effect categories may be more relevant than each of the eight models considered individually.

#### Posterior probability of crossover interaction

From MCMC samples, it is possible to compute posterior probability of the crossover interaction as follows. Let *A* be the event of a crossover interaction. Then, for *j* = 1, …, 8, we have

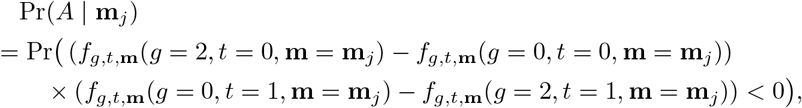

where *f*_*g,t*,**m**_(*·, ·*) is defined as in Eq (16). Since Pr(*A*|**m**_*j*_) = 0 for *j* = 1, …, 6, by the law of total probability, we have

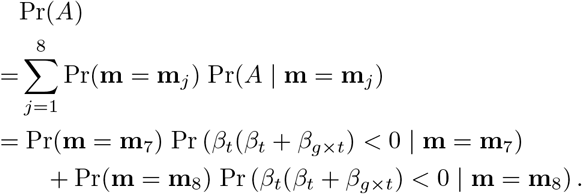

For *j* = 7, 8, we approximated the conditional posterior probability as

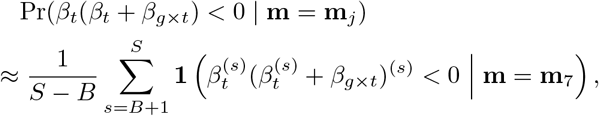

where *S* is the number of MCMC samples, *B* is the number of burn-in iterations, 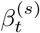 and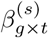 denote samples at the *s*-th iteration. We set *S* = 2000 and *B* = 1000.

#### Subcategories of models based on the sign of effects

To account for the sign of the effect sizes for a given model, we examined the sign of the posterior means or MAP estimates of non-zero coefficients and assigned the same amount of probability to the model with corresponding sign combination and zero probability to others. For example, if **m** = **m**_7_ = (0, 1, 1), we examined the sign of the estimates 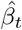 and 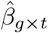 given that the seventh model **m**_7_ was correct. If 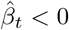 and 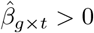, we set

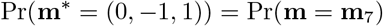

and

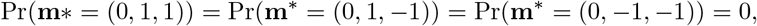

where **m**^∗^ = *{*−1, 0, 1*}*^3^ is a vector specifying the 27 model categories.

#### Computing the marginal likelihood by Laplace approximation

To approximate the marginal likelihood, we obtain a MAP estimate as well as an approximate Hessian matrix evaluated at the MAP value using the BFGS algorithm as implemented in the optim function in the R package stats for each of the eight models. We then compute the value as

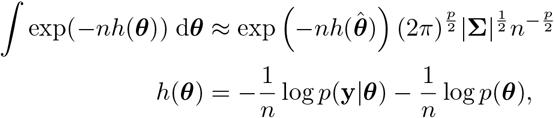

where *n* denotes the sample size, **y** ∈ ℝ^*n*^ denotes a vector of outcome variables, ***θ*** = (*β*_0_, *β*_*g*_, *β*_*t*_, *β*_*g×t*_, *σ*^2^)^*T*^ ∈ ℝ^*p*^ denotes a vector of parameters 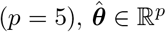 is the mode of −*h*(***θ***), **Σ** ∈ ℝ^*p×p*^ denotes the inverse negative of the Hessian matrix evaluated at the mode. Note that the marginal likelihood takes the form of

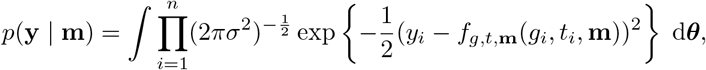

where *f*_*g,t*,**m**_ is defined as in Eq (16).

### Hyperparameter optimization using empirical Bayes

In BMS, properly optimizing the prior hyperparameters can improve inference. For the optimization, we undertook an empirical Bayes approach, where we used a grid search to obtain the values of ***ϕ*** that maximize the sum of the log of marginal likelihood across all feature-SNP pairs under consideration. Note that ***ϕ*** = (*ϕ*_*g*_, *ϕ*_*t*_, *ϕ*_*g×t*_) control the effect sizes relative to the residual error standard deviation. In this approach, the sum of the log of marginal likelihood is maximized over a grid of candidate hyperparameter values. For simulation analysis, optimal values were searched over a grid spanning from 0.25 to 5.0 by 0.25. For analysis of the experimental data, searches were performed over a grid spanning from 0.25 to 3.0 by 0.25. For computational efficiency, we use MAP estimation via optimization followed by Laplace approximation rather than HMC. In principle, the sum of the maximum likelihood across all candidate feature-SNP pairs in the data needs to be maximized. For computational efficiency, however, we estimate hyperparameter values only from the pre-selected set of response molecular QTLs that are being categorized. This choice can be justified as follows. Assume that, for a given feature-SNP pair, the “correct” model is **m** = (1, 1, 0). Then, consider the marginal likelihood

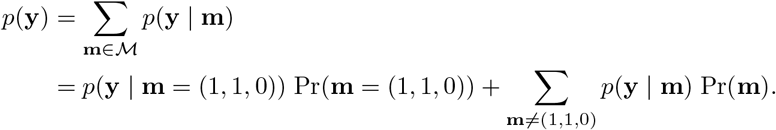

In the last line, the first term has a large *p*(**y** | **m**) value and does not contain *ϕ*_*g×t*_. The second term is small since all other *p*(**y** | **m**) values are small. Thus, *p*(**y**) does not substantially depend on *ϕ*_*g×t*_. That is, this pair does not substantially affect the functional relationship between the marginal likelihood and *ϕ*_*g×t*_. If the “correct” model is **m** = (0, 0, 0), none of the ***ϕ*** elements substantially impact the marginal likelihood. Thus, it is reasonable to focus on feature-SNP pairs with significant associations to estimate optimal values of ***ϕ***. We also note that, for the accuracy of model selection, hyperparameter values only need to be in an appropriate scale but do not need to be exactly maximizing the likelihood of the data. The hyperparameter optimization results are described in S1 Text. The optimal values we obtained are summarized in S1 Table and S2 Table.

### Including covariates

In analysis of experimental data, it is crucial to control for confounding factors to avoid spurious inference. To specify the model that includes confounding factors as fixed or random effects, we let *k* denote the number of donors and *n*_*j*_ denote the number of samples in the *j*-th donor group. That is, 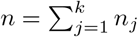. Then, for the *i*-th sample in the *j*-th donor group (*i* = 1, …, *n*_*j*_, *j* = 1, …, *k*), a G×T interaction model can be cast as

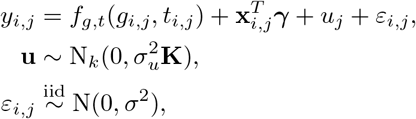

where *y*_*i,j*_ denotes the phenotype, *f*_*g,t*_(*·, ·*) is defined as in Eq (11), **x**_*i,j*_ ∈ R^*q*^ denotes a vector of fixed-effect factors, ***γ*** ∈ ℝ^*q*^ denotes a vector of the corresponding coefficient, **u** ∈ ℝ^*k*^ denotes a vector of random effects, *σ*_*u*_ denotes the random effect standard deviation, *ε*_*i,j*_ denotes the residual error, and *σ*^2^ denotes the residual error variance.

The matrix **K** denotes a kernel matrix and can be a known kinship matrix, which represents genetic relatedness between donors. Alternatively, when the genetic relatedness is included as a fixed effect, **K** can be set to the *k*-dimensional identity matrix **I**_*k*_. In the matrix form, we have

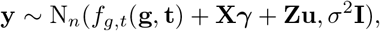

where **y** ∈ ℝ^*n*^ denotes the outcome vector,

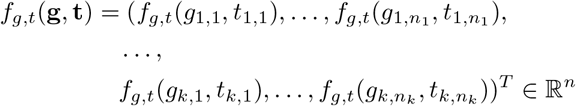

denotes the mean values, **X** ∈ ℝ^*n×q*^ denotes a matrix of fixed-effect factors, and **Z** ∈ ℝ^*n×k*^ denotes an incidence matrix indicating the group membership of the samples, Since the goal is to estimate the parameters of the *f*_*g,t*_(*·*) function, *β*, we marginalize out the random effects **u** and obtain

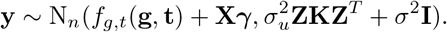

Then, the parameters *β, γ, σ*_*u*_, and *σ* can be estimated simultaneously by a frequentists or a Bayesian approach.

In our G×T analysis of the gene expression and chromatin accessibility data from hNPCs, we included top 10 principal components (PCs) of the molecular count data as fixed effects and the genetic relatedness (kinship) among the donors as a random effect as in the previous study [10]. The kinship matrix was obtained using the R package GENESIS [50]. The molecular count PCs were identified in two steps. First, for each feature, we combined the library size-scaled, log-transformed molecular count data in the control and treated conditions and residualized the combined data with respect to the treatment indicator variable. We did not regress out the donor or kinship random effect since the effect is unlikely to be captured in the top PCs. Second, we obtained PCs by performing principal component analysis (PCA) on the residuals. Then residualized phenotype data, *y*_*i*_, was obtained by regressing out the top 10 PCs from the library size-scaled, log-transformed data before the first step. We included 22354 genes and 172887 ATAC-peak regions as features. Although it is ideal to control for the factors simultaneously with examining the associations of the phenotype with the genotype and treatment, such model formulation can pose computational challenges due to a large number of parameters to be estimated. Therefore, we regressed out the library-size scaled, log-transformed count data with respect to the fixed-effect confounding factors and fit models including the random effect only. Note that we did not consider fixed-effect confounding factors in the simulation experiments.

For the single-condition analysis for assessing the allelic additivity, we modeled

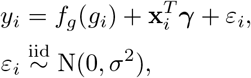

where *y*_*i*_, *g*_*i*_, *σ*^2^, and ∈_*i*_ are defined as in Eq (2) and (3), *f*_*g*_(*·*) is defined as in Eq (4), **x**_*i*_ ∈ ℝ^*q*^ is a vector of fixed-effect factors for the *i*-th sample, and ***γ*** ∈ ℝ^*q*^ is a vector of the corresponding coefficient. As fixed-effect factors, we included top 10 PCs, which we obtained by performing PCA separately for each condition.

### Generating data for simulations

For simulation experiments, we generated 1000 sets of feature-SNP pairs for each of the eight models specified in Eq (16), summing to 8000 feature-SNP pairs in total. Each simulated feature-SNP pair comprised the genotypes for 80 individuals, with these drawn from a Binomial distribution, Binom(*n* = 2, *p* = *π*), where *π* ∼ Uniform(0.05, 0.5), corresponding to minor allele frequency (MAF) ranging from 0.05 to 0.5, and simulated normalized count data 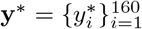(ie, the phenotype vector). In generating a simulated phenotype vector from a given model, we fixed the intercept and residual error standard deviation to *b*_0_ = 0 and *σ* = 1, set the donor random effect variance 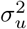 to 0.2, and set the kernel matrix **K** to the identity. The regression coefficients were drawn from Normal distributions as in Eq (18, 19, 20) with ***ϕ*** = (*ϕ*_*g*_, *ϕ*_*t*_, *ϕ*_*g×t*_)^*T*^ = (1.5, 2.0, 1.0)^*T*^, with these values based on the results of hyperparameter calibration using the hNPC eQTL data. This generated 80 phenotype values for each of the control and treated conditions (160 in total) based on Eq (14, 16). For evaluating relative performance of log-NL, log-LM, and RINT-LM, we analyzed the entire set of 8000 simulated feature-SNP pairs using both MAP estimation followed by Laplace approximation to get posterior model probabilities, and MCMC followed by bridge sampling to compute the posterior probability of crossover interaction. In the other MCMC analyses comparing the three approaches, we used 100 feature-SNP pairs, summing to 800 in total, due to the computational cost. In our comparisons of estimating the marginal likelihoods via Laplace approximation vs bridge sampling, we used 10 feature-SNP pairs, summing to 80 in total, albeit for 125 combinations of hyperparameter values.

## Supporting information

**S1 Fig. Assessing the allelic additivity assumption in hNPCs. A**.

Scatterplots comparing the maximized likelihood between nonlinear and linear regression for 3073 gene-SNP pairs under the control condition. **B**. The same as in **A** but between the ANOVA model and nonlinear regression. **C**. The same as in **A** but between the ANOVA model and linear regression. **D**. The same as in **A** but under the treated condition. **E**. The same as in **B** but under the treated condition. **F**. The same as in **C** but under the treated condition. **G**. Scatterplots comparing the maximized likelihood between nonlinear and linear regression for 42576 cCRE-SNP pairs under the control condition. **H**. The same as in **G** but between the ANOVA model and nonlinear regression. **I**. The same as in **G** but between the ANOVA model and linear regression. **J**. The same as in **G** but under the treated condition. **K**. The same as in **H** but under the treated condition. **L**. The same as in **I** but under the treated condition.

**S2 Fig. Calibration of BMS with** log**-NL**, log**-LM, and RINT-LM for the no-G**×**T, induced, and altered categories using MCMC and bridge sampling**.

The *x*- and *y*-axis represent the posterior probability and the fraction of the corresponding events, respectively. The results from 800 simulations are grouped into ten equally-spaced bins. The vertical bars represent the standard errors assuming a binomial distribution. Shown are results from analysis without random effect, which we call scenario 1 (**A**), those with donor random effect in model fitting but not in data generation (scenario 2) (**B**), those with donor random effect in data generation but not in model fitting (scenario 3) (**C**), and those with donor effect in both data generation and model fitting (scenario 4) (**D**).

**S3 Fig. Calibration of BMS with** log**-NL**, log**-LM, and RINT-LM for the no-G**×**T, induced, and altered categories using MAP and Laplace approximation**. The same as in S2 Fig but from 8000 simulations using MAP estimation and Laplace approximation.

**S4 Fig. ROC curves assessing the performance of BMS with** log**-NL**, log**-LM, and RINT-LM for the no-G**×**T, induced, and altered categories using MCMC and bridge sampling**. The panels **A** to **D** show the results in scenarios 1 to 4, which are described in the legend to S2 Fig.

**S5 Fig. ROC curves assessing the performance of BMS with** log**-NL**, log**-LM, and RINT-LM for the no-G**×**T, induced, and altered categories using MAP and Laplace approximation**. The panels **A** to **D** show the results in scenarios 1 to 4, which are described in the legend to S2 Fig.

**S6 Fig. Stratified histograms of posterior probability of the eight models obtained by BMS with** log**-NL using MCMC and bridge sampling**. In each panel, the rows and columns represent the data-generating and posterior mode models, respectively. The panels **A** to **D** show the results in scenarios 1 to 4, which are described in the legend to S2 Fig.

**S7 Fig. Stratified histograms of posterior probability of the eight models obtained by BMS with** log**-LM using MCMC and bridge sampling**. The same as in S6 Fig but for log-LM.

**S8 Fig. Stratified histograms of posterior probability of the eight models obtained by BMS with RINT-LM using MCMC and bridge sampling**. The same as in S6 Fig but for RINT-LM.

**S9 Fig. Stratified histograms of posterior probability of the eight models obtained by BMS with** log**-NL using MAP estimation and Laplace approximation**. The same as in S6 Fig but for MAP estimation and Laplace approximation.

**S10 Fig. Stratified histograms of posterior probability of the eight models obtained by BMS with** log**-LM using MAP estimation and Laplace approximation**. The same as in S9 Fig but for log-LM.

**S11 Fig. Stratified histograms of posterior probability of the eight models obtained by BMS wth RINT-LM using MAP estimation and Laplace approximation**. The same as in S9 Fig but for RINT-LM.

**S12 Fig. Posterior probability of the correct and incorrect models for the eight categories obtained by BMS using MCMC and bridge sampling on data generated without random effect**. Violin plots for comparing the performance of BMS with log-NL, log-LM, and RINT-LM based on the distribution of posterior probability of the correct and incorrect models for each of the eight model categories. The closed circles represent median values. The panels **A** and **B** respectively show the results in scenarios 1 and 2, which are described in the legend to S2 Fig.

**S13 Fig. Posterior probability of the correct and incorrect models for the eight categories obtained by BMS using MCMC and bridge sampling on data generated with donor random effect**.. The same as S12 Fig but on data generated with donor random effect. The panels **A** and **B** respectively show the results in scenarios 3 and 4, which are described in the legend to S2 Fig.

**S14 Fig. Posterior probability of the correct and incorrect models for the eight categories obtained by BMS using MAP estimation and Laplace approximation on data generated without random effect**. The same as in S12 Fig but for MAP estimation and Laplace approximation.

**S15 Fig. Posterior probability of the correct and incorrect models for the eight categories obtained by BMS using MAP estimation and Laplace approximation on data generated with donor random effect**. The same as in S13 Fig but for MAP estimation and Laplace approximation.

**S16 Fig. Posterior probability of the correct and incorrect models for aggregated categories obtained by BMS using MCMC and bridge sampling**. Violin plots for comparing the performance of BMS with log-NL, log-LM, and RINT-LM based on the distribution of posterior probability of the correct and incorrect models for the no-G×T, induced, and altered model categories. The closed circles represent median values. The panels **A** to **D** show the results in scenarios 1 to 4, which are described in the legend to S2 Fig.

**S17 Fig. Posterior probability of the correct and incorrect models for aggregated categories obtained by BMS using MAP estimation and Laplace approximation**. The same as in S16 Fig but for MAP estimation and Laplace approximation.

**S18 Fig. Assessing the impact of the effect prior on the posterior probability of the correct and incorrect models from analyses without random effect using MCMC and bridge sampling**. Violin plots showing the distribution of posterior probability of the correct (**A**) and incorrect (**B**) models for each of the eight model categories with varying hyper parameter values (see S1 Text). The closed circles represent median values. Shown is the results in scenario 1, which is described in the legend to S2 Fig.

**S19 Fig. Assessing the impact of the effect prior on the posterior probability of the correct and incorrect models from analyses with donor random effect in model fitting but not in data generation using MCMC and bridge sampling**. The same as in S18 Fig but in scenario 2, which is described in the legend to S2 Fig.

**S20 Fig. Assessing the impact of the effect prior on the posterior probability of the correct and incorrect models from analyses with donor random effect in data generation but not in model fitting using MCMC and bridge sampling**. The same as in S18 Fig but in scenario 3, which is described in the legend to S2 Fig.

**S21 Fig. Assessing the impact of the effect prior on the posterior probability of the correct and incorrect models from analyses with donor random effect in both data generation and model fitting using MCMC and bridge sampling**. The same as in S18 Fig but in scenario 4, which is described in the legend to S2 Fig.

**S22 Fig. Assessing the impact of the effect prior on the posterior probability of the correct and incorrect models from analyses without random effect using MAP estimation and Laplace approximation**. The same as in S18 Fig but for MAP estimation and Laplace approximation.

**S23 Fig. Assessing the impact of the effect prior on the posterior probability of the correct and incorrect models from analyses with donor random effect in model fitting but not in data generation using MAP estimation and Laplace approximation**. The same as in S19 Fig but for MAP estimation and Laplace approximation.

**S24 Fig. Assessing the impact of the effect prior on the posterior probability of the correct and incorrect models from analyses with donor random effect in data generation but not in model fitting using MAP estimation and Laplace approximation**. The same as in S20 Fig but for MAP estimation and Laplace approximation.

**S25 Fig. Assessing the impact of the effect prior on the posterior probability of the correct and incorrect models from analyses with donor random effect in both data generation and model fitting using MAP estimation and Laplace approximation**. The same as in S21 Fig but for MAP estimation and Laplace approximation.

**S26 Fig. Comparison of the sum of the** log **of marginal likelihood across combinations of hyperparameter values between two computational approaches for** log**-NL**, log**-LM, and RINT-LM with and without donor random effect**. Scatter plots comparing results obtained by MCMC followed by bridge sampling and those obtained by MAP estimation followed by Laplace approximation. Each point represents the log of the marginal likelihood summed over 80 feature-SNP pairs. The values are compared across 125 combinations of the hyperparameter values (see S1 Text for details).

**S27 Fig. Comparison of posterior probability at the optimal hyperparameter values between two computational approaches for** log**-NL**, log**-LM, and RINT-LM with and without donor random effect**. Scatter plots comparing results obtained by MCMC followed by bridge sampling and those obtained by MAP estimation followed by Laplace approximation. Each point represents the posterior probability of a mode for a feature-SNP pair. The values are compared across eight models and 800 feature-SNP pairs (ie, 6400 combinations).

**S28 Fig. Posterior probability of the models with and without accounting for the sign of effect size for the response eQTL data in hNPCs**. The heatmaps show the posterior probability of the eight models for the 98 response eQTLs, which represent gene-SNP pairs with significant G×T interactions (**A**), as well as that of the 27 models accounting for the sign of effect size (**B**). The rows and columns represent the models and gene-SNP pairs, respectively. The gene-SNP pairs are ordered by *P* values for significant G×T interactions from a previous study [10]. The leftmost column corresponds to the smallest *P* value.

**S29 Fig. Posterior probability of the models with and without accounting for the sign of effect size for the response caQTL data in hNPCs**. The same as in S28 Fig but for 1775 response caQTLs.

**S30 Fig. Representative BMS results for the response caQTL data in hNPCs**. The same as in Fig. 5 but for response caQTLs.

**S31 Fig. Examples of response caQTLs with the crossover interaction in hNPCs**. The same as in Fig. 6 but for response caQTLs.

**S32 Fig. Comparison of posterior probability between results with donor random effect and those with polygenic random effect for response eQTLs**. Scatter plots comparing results obtained by BMS with polygenic (kinship) random effect and those with donor random effect. Each point represents the posterior probability of a mode for a feature-SNP pair. The values are compared across eight models and 98 feature-SNP pairs (ie, 784 combinations).

**S1 Text Supplementary methods and results**.

**S1 Table Optimal hyperparameter values for simulation**. Shown are values of the hyperparameters ***ϕ*** that maximize the sum of the log-marginal likelihood across 8000 feature-SNP pairs. The **Method** column indicates modeling approaches used in the analysis. The plus and minus signs in the **Generation** and **BMS** columns respectively indicate the presence and absence of random effect in data generation and model fitting. The **Genotype, Treatment**, and **Interaction** columns correspond to *ϕ*_*g*_, *ϕ*_*t*_, and *ϕ*_*g×t*_, respectively.

**S2 Table Optimal hyperparameter values for the hNPC data**. Shown are values of the hyperparameters ***ϕ*** that maximize the sum of the log-marginal likelihood across feature-SNP pairs. The **Method, Genotype, Treatment**, and **Interaction** columns are as in Table S1 Table. The **Data** and **Random effect** columns indicate the type of data and random effect, respectively.

**S1 File A frozen version (0.0.1) of classifygxt, the R package implementation**.

**S2 File A frozen version of code for the analyses**.

## Acknowledgments

We thank Samir Kelada and Yun Li of the University of North Carolina at Chapel Hill for helpful discussion and suggestions on this work.

