## Supplemental Text for "Probabilistic classification of gene-by-treatment interactions on molecular count phenotypes"

### S1 Text

July 31, 2024

#### 1 The log-NL model of G×T analysis

Consider the association of a molecular phenotype, measured for a given feature, such as a gene or accessible chromatin region, with a putative *cis*-regulatory variant that has two alleles in the population, A and B, under two mutually exclusive treatment conditions. We model

$$y_i = \log(\mu_{g_i,t_i}) + \varepsilon_i, \tag{1}$$

$$\varepsilon_i \overset{\text{iid}}{\sim} \mathrm{N}(0, \sigma^2), \tag{2}$$

$$\begin{aligned}\mu_{g,t}(g_i, t_i) = &(1 - \tfrac{g_i}{2})(1 - t_i)\exp(\beta_0) + (\tfrac{g_i}{2})(1 - t_i)\exp(\beta_0 + 2\beta_g) \\ &+ (1 - \tfrac{g_i}{2})(t_i)\exp(\beta_0 + \beta_t) + (\tfrac{g_i}{2})(t_i)\exp(\beta_0 + 2\beta_g + \beta_t + 2\beta_{g\times t}),\end{aligned} \tag{3}$$

where $y_i$ denotes log-transformed molecular phenotype for the $i$-th individual ($i = 1, \cdots, n$), $g_i$ denotes the genotype coded as $\{0, 1, 2\}$ or the imputation-based allelic dosage as $g_i \in [0, 2]$, $t_i$ denotes an indicator variable for a treatment, $\beta_0$, $\beta_g$, $\beta_t$, and $\beta_{g\times t}$ denote regression coefficients, $\varepsilon_i$ denotes the residual error, and $\sigma^2$ denotes the residual error variance. The $\mu_{g,t}(g, t)$ function returns a value corresponding to the phenotype in the original count scale except that, according to previous studies ([1, 2]), we do not model a pseudocount of one, which is typically added to empirical count data. Note that, in the main text, we define $f_{g,t}(g, t) = \log(\mu_{g,t}(g, t))$. For simplicity, confounding factors are omitted.

##### 1.1 Derivation

The functional form of $\mu_{g,t}(g_i, t_i)$ in Eq (3) can be derived as follows. In what follows, the subscript $i$ is omitted for simplicity. Denote the major and minor alleles of a given SNP as A and B, respectively. Let $\mu_{AA,c}$ and $\mu_{AA,t}$ denote the phenotype values in the control and treated conditions, for the major allele homozygous respectively. Let $\mu_{BB,c}$ and $\mu_{BB,t}$ denote the phenotype values in the control and treated conditions, for the minor allele homozygous respectively. The quantities can be defined as

$$\begin{aligned}\log(\mu_{AA,c}) &= \log(\mu_{g,t}(g = 0, t = 0)) = \beta_0, \\ \log(\mu_{BB,c}) &= \log(\mu_{g,t}(g = 2, t = 0)) = \beta_0 + 2\beta_g, \\ \log(\mu_{AA,t}) &= \log(\mu_{g,t}(g = 0, t = 1)) = \beta_0 + \beta_t, \\ \log(\mu_{BB,t}) &= \log(\mu_{g,t}(g = 2, t = 1)) = \beta_0 + 2\beta_g + \beta_t + 2\beta_{g\times t}.\end{aligned}$$

By exponentiating sides, we have

$$\mu_{AA,c} = \exp(\beta_0), \tag{4}$$
$$\mu_{BB,c} = \exp(\beta_0 + 2\beta_g), \tag{5}$$
$$\mu_{AA,t} = \exp(\beta_0 + \beta_t), \tag{6}$$
$$\mu_{BB,t} = \exp(\beta_0 + 2\beta_g + \beta_t + 2\beta_{g\times t}). \tag{7}$$

In this scale, the expression values are linear with respect to the genotype. Thus, by considering the interpretation of the coefficients in a linear regression model, we have

$$\mu_{g,t}(g,t) = \mu_{AA,c} + \tfrac{\mu_{BB,c}-\mu_{AA,c}}{2} g + (\mu_{AA,t} - \mu_{AA,c})t + (\tfrac{\mu_{BB,t}-\mu_{AA,t}}{2} - \tfrac{\mu_{BB,c}-\mu_{AA,c}}{2})gt \tag{8}$$
$$= (1 - \tfrac{g}{2})(1-t)\mu_{AA,c} + (\tfrac{g}{2})(1-t)\mu_{BB,c} + (1-\tfrac{g}{2})(t)\mu_{AA,t} + (\tfrac{g}{2})(t)\mu_{BB,t} \tag{9}$$
$$= (1 - \tfrac{g}{2})(1-t)\exp(\beta_0) + (\tfrac{g}{2})(1-t)\exp(\beta_0 + 2\beta_g)$$
$$+ (1-\tfrac{g}{2})(t)\exp(\beta_0+\beta_t) + (\tfrac{g}{2})(t)\exp(\beta_0 + 2\beta_g + \beta_t + 2\beta_{g\times t}), \tag{10}$$

for any $g \in [0, 2]$ and $t \in \{0, 1\}$.

#### 1.2 An alternative notation

Similarly to a previous study by Mohammadi *et al.* [1], we let $\mu_{A,c} = \alpha$ denote the contribution to the phenotype value in the original count scale, $\exp(y)$, from one copy of the reference allele A in the control condition. We denote the fraction of an increase in the phenotype value per one copy of the alternative allele in the control condition by $\delta_g$. Likewise, we denote the fraction of an increase in the phenotype value by treatment for one copy of the reference allele by $\delta_t$. We then denote the fraction of an increase in the phenotype value per one copy of the alternative allele in the treatment condition relative to that in the control condition by $\delta_{g\times t}$. Then, the contribution from one copy of a given allele in a given condition is given by

$$\mu_{B,c} = \alpha\delta_g,$$
$$\mu_{A,t} = \alpha\delta_t,$$
$$\mu_{B,t} = \alpha\delta_g\delta_t\delta_{g\times t}.$$

Thus, the phenotype values can be written as

$$\mu_{AA,c} = \mu_{A,c} + \mu_{A,c} = 2\alpha, \tag{11}$$
$$\mu_{AB,c} = \mu_{BA,c} = \mu_{A,c} + \mu_{B,c} = (1+\delta_g)\alpha, \tag{12}$$
$$\mu_{BB,c} = \mu_{B,c} + \mu_{B,c} = 2\alpha\delta_g, \tag{13}$$
$$\mu_{AA,t} = \mu_{A,t} + \mu_{A,t} = 2\alpha\delta_t, \tag{14}$$
$$\mu_{AB,t} = \mu_{BA,t} = \mu_{A,t} + \mu_{B,t} = (1+\delta_g\delta_{g\times t})\alpha\delta_t, \tag{15}$$
$$\mu_{BB,t} = \mu_{B,t} + \mu_{B,t} = 2\alpha\delta_g\delta_t\delta_{g\times t}, \tag{16}$$

based on the assumption of allelic additivity. Substituting Eq (11), (13), (14), and (16) for $\mu_{A,c}$, $\mu_{A,t}$, $\mu_{B,c}$, and $\mu_{B,t}$ in Eq (8), we have

$$\mu_{g,t} = 2\alpha + \alpha(\delta_g - 1)g + 2\alpha(\delta_t - 1)t + \alpha(\delta_g\delta_t\delta_{g\times t} - \delta_g - \delta_t + 1)gt, \tag{17}$$

for any $g \in [0,2]$ and $t \in \{0,1\}$. From a comparison of Eq (8), Eq (4), and Eq (17), we have

$$\begin{aligned}
\alpha &= \frac{1}{2}\exp(\beta_0), \\
\delta_g &= \exp(2\beta_g), \\
\delta_t &= \exp(\beta_t), \\
\delta_{g\times t} &= \exp(2\beta_{g\times t}).
\end{aligned}$$

The eight models can be obtained as

$$\mu_{g,t} = \begin{cases}
2\alpha, & \text{if } \delta_g = 1,\ \delta_t = 1,\ \delta_{g\times t} = 1 \\
2\alpha + \alpha(\delta_g - 1)g, & \text{if } \delta_g \neq 1,\ \delta_t = 1,\ \delta_{g\times t} = 1 \\
2\alpha + 2\alpha(\delta_t - 1)t, & \text{if } \delta_g = 1,\ \delta_t \neq 1,\ \delta_{g\times t} = 1 \\
2\alpha + 2\alpha(\delta_g - 1)g + 2\alpha(\delta_t - 1)t + \alpha(\delta_g\delta_t - \delta_g - \delta_t + 1)gt, & \text{if } \delta_g \neq 1,\ \delta_t \neq 1,\ \delta_{g\times t} = 1 \\
2\alpha + \alpha(\delta_{g\times t} - 1)gt, & \text{if } \delta_g = 1,\ \delta_t = 1,\ \delta_{g\times t} \neq 1 \\
2\alpha + \alpha(\delta_g - 1)g + \alpha\delta_g(\delta_{g\times t} - 1)gt, & \text{if } \delta_g \neq 1,\ \delta_t = 1,\ \delta_{g\times t} \neq 1 \\
2\alpha + 2\alpha(\delta_t - 1)t + \alpha\delta_t(\delta_{g\times t} - 1)gt, & \text{if } \delta_g = 1,\ \delta_t \neq 1,\ \delta_{g\times t} \neq 1 \\
2\alpha + \alpha(\delta_g - 1)g + 2\alpha(\delta_t - 1)t + \alpha(\delta_g\delta_t\delta_{g\times t} - \delta_g - \delta_t + 1)gt, & \text{if } \delta_g \neq 1,\ \delta_t \neq 1,\ \delta_{g\times t} \neq 1.
\end{cases}$$

Moreover, in the control and treated conditions ($t = 0$ and $t = 1$), we respectively have

$$\mu_g = 2\alpha + \alpha(\delta_g - 1)g$$

and

$$\mu_g = 2\alpha\delta_t + \alpha\delta_t(\delta_g\delta_{g\times t} - 1)g.$$

Here, $\alpha$ and $\alpha\delta_t$ correspond to the phenotype values for the major allele homozygous donors in the control and treated conditions, respectively. $\delta_g$ and $\delta_g\delta_{g\times t}$ correspond to the allelic fold changes (the fraction of an increase in the phenotype value per one copy of the alternative allele) in the control and treated conditions, respectively. Thus, $\delta_{g\times t} = 1$ ($\beta_{g\times t} = 0$) corresponds to a situation where the allelic fold changes are identical between the two conditions. Note that, if $\delta_t = \delta_{g\times t} = 1$, this model reduces to the nonlinear model for estimating the aFC value (2.1) and the ACME model (2.2). The $\alpha$ parameter corresponds to $e_0$ and $\frac{1}{2}\beta_0$ in the aFC and ACME models, respectively. The $\delta_g$ parameter corresponds to $\exp(\log_2(\delta_{0,1}))$ and $2\eta + 1$ in the aFC and ACME models, respectively.

### 2 Review of previously developed methods

In this section, we review previously identified methods for single-condition molecular QTL mapping based on the allelic additivity assumption. Note that we use notations by the authors, which are not consistent with those in other sections or the main text.

#### 2.1 The aFC method

Mohammadi *et al.* [1] proposed a use of allelic fold change (aFC), $\delta_{0,1}$, for quantifying the effect size of *cis*-eQTLs. In the study, they described a model to estimate aFC from eQTL data with multiplicative noise, which is closely related to our approach. The model is cast as

$$y_i = \{(2 - t_n) - t_n\delta_{1,0}\}e_0\,\varepsilon_n, \tag{18}$$

where $y_n$ denotes the gene expression in the original count scale for the $n$-th individual, $t_n$ denotes the number of alternative alleles coded as $\{0, 1, 2\}$, $e_0$ is the expression per copy of the reference allele, and $\varepsilon_n$ denotes a noise such that $\log_2(\varepsilon_n)$ is normally distributed with null mean and unknown variance. Taking the logarithm of the both-hand sides of Eq (18) yields

$$\log_2(y_n) = \log_2\{(2 - t_n) + t_n\delta_{1,0}\} + \log_2 e_0 + \log_2 \varepsilon_n.$$

Note that confounding factors are omitted for the ease of exposition.

#### 2.2 The ACME model

Palowitch *et al.* [2] proposed the Additive Contribution on the original expression scale with Multiplicative Error (ACME) model. The model is cast as

$$y_i = \log(\beta_0 + \beta_1 s_i) + \mathbf{Z}_i^T \boldsymbol{\gamma} + \varepsilon_i, \tag{19}$$

$$\varepsilon_i \overset{\text{iid}}{\sim} \mathrm{N}(0, \sigma^2), \tag{20}$$

where $y_i$ denotes the log-transformed read count for the $i$-th individual ($i = 1, \cdots, n$), $s_i$ denotes the genotype (the minor allele count) coded as $\{0, 1, 2\}$, $\beta_0$ is the baseline mean expression, $\beta_1$ is the additive contribution of each allele, $\mathbf{Z}_i \in \mathbb{R}^{p\times 1}$ denotes a vector of covariates, $\boldsymbol{\gamma} \in \mathbb{R}^{p\times 1}$ denotes corresponding coefficients, $\varepsilon_i$ denotes independent residual errors, and $\sigma^2$ denotes the residual error variance. Eq (19) can be rewritten as

$$\begin{aligned} y_i &= \log(\beta_0) + \log(1 + \frac{\beta_1}{\beta_0}) + \mathbf{Z}_i \boldsymbol{\gamma} + \varepsilon_i, \\ &= \log(\beta_0) + \log(1 + \eta) + \mathbf{Z}_i \boldsymbol{\gamma} + \varepsilon_i, \end{aligned}$$

where $\beta$ and $\eta$ determine the baseline expression and the effect of genotype, respectively.

### 3 Evaluating the Laplace approximation

Although MCMC followed by bridge sampling [3] provides accurate approximations with sufficient sampling [4], this approach is computationally intensive and is prohibitive for optimizing hyperparameters. MAP estimation followed by Laplace approximation [5, 6] is magnitudes of order faster and suited for a large number of model fitting routines. Since error bounds for Laplace approximation are not easily obtained [5], we empirically assessed its performance by comparing simulation results obtained by MAP estimation followed by Laplace approximation with those obtained by MCMC followed by bridge sampling. Specifically, we compared the sum of log-marginal likelihood over 80 feature-SNP pairs (i.e., 10 instances per data-generating model) across 125 combinations of hyperparameter values, corresponding to a grid spanning from 0.5 to 2.5 by 0.5. We also compared the posterior probability across the eight models for 800 feature-SNP pairs (i.e., 100 instances per data-generating model).

The analyses led to the following points. First, the sum of log-marginal likelihood $p(\mathbf{y})$ was nearly identical up to normalization constants across the hyperparameter combinations (**S26 Fig.**) In particular, this result can justify the use of MAP estimation and Laplace approximation for optimizing the hyperparameter values. We note that our implementation of Laplace approximation only gives relative log-marginal likelihood values, which suffices for our purposes. Second, the posterior probability of the eight models obtained by the two approaches were nearly identical (**S27 Fig.**) These results collectively suggest the adequacy of Laplace approximation at least in our settings.

#### 4 Hyperparameter optimization

We optimized the hyperparameters of the effect prior for the simulated and experimental data (see **Methods**). The optimal values for the simulation are summarized in **S1 Table**. As expected, log-NL recovered the standard deviations of the non-zero coefficients relative to the residual error standard deviation ($(1.50, 2.00, 1.00)^T$) used for generating the data except for the scenario where the data was generated with random effect but BMS was performed without random effect. The optimal values for the eQTL and caQTL data in hNPCs are summarized in **S2 Table**.

#### 5 Sensitivity of the BMS to the effect prior specification

Prior specification of the model parameters can affect the accuracy of BMS. To assess the sensitivity of our BMS to the effect prior specification, we performed BMS with varying values of the hyper-parameters, $\boldsymbol{\phi} = (\phi_g, \phi_t, \phi_{g\times t})$. Specifically, we examine a half and twice of the optimal values, which are restrictive and permissive, respectively. Overall, we observed moderate but consistent impacts of the effect prior specification. With the restrictive effect priors, more complex models were favored. By contrast, with the permissive effect priors, simpler models were favored (**S18-25 Fig.**).

[5] Jon Wakefield. *Bayesian and Frequentist Regression Methods.* Springer Series in Statistics. Springer New York, New York, NY, 2013.

[6] Jeff Gill. *Bayesian methods: a social and behavioral sciences approach.* Chapman & Hall/CRC statistics in the social and behavioral sciences series. CRC Press, Taylor & Francis Group, Boca Raton, third edition edition, 2015.
