## Supplemental File 1 for "Probabilistic classification of gene-by-treatment interactions on molecular count phenotypes": 404.html

Page not found (404) • classifygxt
 


  

 


Toggle navigation


classifygxt
0.0.1

- Get started
- Reference
- Changelog

### Page not found (404)

Content not found. Please use links in the navbar.

#### Contents

Developed by Yuriko Harigaya, Michael Love, William Valdar.

Site built with pkgdown 2.0.9.
