## Supplemental File 1 for "Probabilistic classification of gene-by-treatment interactions on molecular count phenotypes": classifygxt.html

ClassifyGxT - classifying gene-by-treatment interactions • classifygxt
 


  


 


Toggle navigation


classifygxt
0.0.1

- Get started
- Reference
- Changelog

### ClassifyGxT - classifying gene-by-treatment interactions

###### Yuriko Harigaya, Michael Love, William Valdar

#### 2024-07-10

`classifygxt.Rmd`

#### Introduction

The ClassifyGxT software is an implementation of a Bayesian model
selection (BMS) framework for classifying GxT interactions. The method
was developed primarily for molecular count phenotypes, such as RNA-seq
and ATAC-seq data, although it can be used for other types of
phenotypes.

- The input is a list of feature-SNP pairs for which significant GxT
  interactions have been identified by the standard response molecular QTL
  mapping procedure (see HIPSCI Consortium et al.
  (2018) and Matoba et al. (2023) for
  example). For each of the feature-SNP pairs, individual genotype and
  phenotype data are required.
- The main output is the posterior probability for different GxT
  types. See **Overview** on the main page for the model
  categories representing the types of GxT interactions.
- The package provides functions to process a single feature-SNP pair.
  In practice, we recommend running parallel jobs on the computer cluster
  with appropriate grouping.

#### Input data

For each feature-SNP pair, a list object containing the following
elements needs to be generated:

- `y`: A vector of length \(n\) containing preprocessed molecular count
  phenotypes, where \(n\) is the number
  of samples. See **Data processing** below for
  preprocessing.
- `g`: A vector of length \(n\) containing genotypes coded as \({0, 1, 2}\) to represent the number of
  minor (alternative) alleles or the imputation-based allelic dosage in
  \([0, 2]\).
- `t`: A vector of indicator variables for the
  treatment.
- `subject`: An optional character or numeric vector
  corresponding to the subjects. This is necessary when the model includes
  donor or polygenic (kinship) random effects.
- `feat.id`: An optional character string representing the
  feature.
- `snp.id`: An optional character string representing the
  SNP.

Note that the samples must be in the same order in the
`y`, `g`, `t`, and `subject`
elements and that the list can contain additional elements.

#### Data preprocessing

For molecular count phenotypes, the raw count data from sequencing
experiments can be processed as follows:

- Step 1: make a matrix of feature counts with rows and columns
  representing features and samples, respectively
- Step 2: scale the count data by the library size according to Palowitch et al. (2018)
- Step 3: optionally filter features based on the scaled counts
  according to Matoba et al. (2023)
- Step 4: transform using the function \(\log(x + 1)\)
- Step 5: regress out the treatment indicator (and optionally other
  nuisance factors, such as sex and age of the subjects)
- Step 6: perform principal component analysis (PCA)
- Step 7: regress out an appropriate number of PCs from the original,
  transformed data (from Step 4)

#### Loading packages

```
library(ggplot2)
library(classifygxt)
```

#### Generating data

In this section, we simulate data using `make_data()`. The
simulated data will have the same format as discussed in the previous
section and will be used for demonstrating how to run BMS in the next
sections. For simplicity, we omit random effects. See the function
documentation (`?make_data`) for how to include random
effects.

We first specify the number of feature-SNP pairs for each of the
eight models.

```
(model.name <- get_model_names())
#> [1] "0,0,0" "1,0,0" "0,1,0" "1,1,0" "0,0,1" "1,0,1" "0,1,1" "1,1,1"
```

For this demonstration, we only make data for 10 pairs for each
model.

```
num <- rep(10, 8)
names(num) <- model.name
num
#> 0,0,0 1,0,0 0,1,0 1,1,0 0,0,1 1,0,1 0,1,1 1,1,1 
#>    10    10    10    10    10    10    10    10
```

With this specification, the output will be a list of 80 lists. Each
of the lists corresponds to each feature-SNP pair. The first ten lists
are based on `"0,0,0"`, the next ten are based on
`"1,0,0"`, and so forth.

The genotype, treatment, and interaction effects will be drawn from
Normal distributions with a zero mean and user-specified standard
deviations. These values represent “typical” magnitudes of the effects.
We specify standard deviations of the effects as follows.

```
sd.g <- 1.5 # genotype
sd.t <- 2.0 # treatment
sd.gxt <- 1.0 # interaction
sd <- c(sd.g, sd.t, sd.gxt)
```

Note that the residual error standard deviation \(\sigma\), which represents a typical
magnitude of noise, is set to 1 by default.

The following code generates a data frame specifying the mapping
between samples, subjects, and treatment conditions.

```
n.sub <- 80 # number of subjects
anno <- data.frame(
    subject=rep(seq_len(n.sub), each=2),
    condition=rep(c(0, 1), times=n.sub))
head(anno)
#>   subject condition
#> 1       1         0
#> 2       1         1
#> 3       2         0
#> 4       2         1
#> 5       3         0
#> 6       3         1
```

Now we can generate fake data using `make_data()`.

```
data.list <- make_data(
    anno=anno, fn="nonlinear",
    num=num, sd=sd)
```

#### Performing BMS

In this section, we perform BMS using `do_bms()`. For
simplicity, we do not model random effects. See **Including donor
random effects** below for how to include random effects.

We first specify the “model prior.” Note that this is an optional
argument and that, by default, a uniform prior is used. This
specification reflects a prior belief that all models are equally
likely. Here, we explicitly specify the default model prior for
demonstration purposes.

```
p.m <- rep(1/8, 8)
```

We also specify the “effect prior” by choosing the values of the
\(\phi\_g\), \(\phi\_t\), and \(\phi\_{g \times t}\) hyperparameters, which
correspond to the genotype, treatment, and GxT interaction effects,
respectively. The hyperparameters represent our prior beliefs about the
effects relative to the residual error standard deviation (noise).
Specifically, we place priors, \[\begin{equation}
\beta\_g \sim \mathrm{N}(0, \phi\_g^2 \sigma^2), \quad
\beta\_t \sim \mathrm{N}(0, \phi\_t^2 \sigma^2), \quad
\beta\_{g \times t} \sim \mathrm{N}(0, \phi\_{g \times t}^2 \sigma^2).
\end{equation}\] In practice, we recommend optimizing the values
via an empirical Bayes approach (see **Optimizing the effect prior
hyperparameters** below). Here, we set them to the same values as
the effect standard deviations in the data-generating models. This is a
reasonable choice since we set the residual error standard deviation to
1 when generating the data.

```
phi.g <- 1.5 # genotype
phi.t <- 2.0 # treatment
phi.gxt <- 1.0 # interaction
phi <- c(phi.g, phi.t, phi.gxt)
```

The following code performs BMS for the 71st data using
`do_bms()` with nonlinear regression
(`fn="nonlinear"`). Recall that the 71-80th data were
generated based on the eighth model, `"1,1,1"` (see
**Generating data** above). The `method` input
argument must be set to either `"mcmc.bs"`, which represents
Markov Chain Monte Carlo (MCMC) followed by bridge sampling, or
`"map.lap"`, which represents MAP estimation followed by
Laplace approximation. Although the latter method can be orders of
magnitude faster, we recommend using `"mcmc.bs"` for the
final results, if possible, since it is not straightforward to obtain
error bounds for Laplace approximation.

```
k <- 71
data <- data.list[[k]]
res <- do_bms(
    data=data, p.m=p.m, phi=phi,
    fn="nonlinear", method="map.lap")
```

This returns a list object containing the following elements. See the
function documentation (`?do_bms`) for details.

```
names(res)
#> [1] "fn"             "ranef"          "rint"           "p.m"           
#> [5] "seed"           "ln.p.y.given.m" "ln.p.y"         "p.m.given.y"   
#> [9] "optim.list"
```

For multiple feature-SNP pairs, we recommend processing the data in
batches using a workflow such as Snakemake. Within
a batch, BMS can be performed in serial or parallel processes.

For a moderate number of feature-SNP pairs, the following code can be
used.

```
res.list <- lapply(
    X=data.list, FUN=do_bms, p.m=p.m, phi=phi,
    fn="nonlinear", method="map.lap")
```

##### Extracting the posterior probability of the models

We can extract the posterior probability of the model as follows.

```
(pp <- get_pp(res))
#>        0,0,0        1,0,0        0,1,0        1,1,0        0,0,1        1,0,1 
#> 3.848884e-31 3.786639e-13 2.710346e-25 6.229057e-01 8.811563e-19 6.447401e-12 
#>        0,1,1        1,1,1 
#> 4.816906e-14 3.770943e-01
```

Note that, in this particular example, the highest posterior
probability is assigned to the fourth model `"1,1,0"`, even
though the data-generating model is “`1,1,1`”. However, we
will see that BMS tends to choose the correct models across multiple
feature-SNP pairs (**Heatmap of the posterior probability of the
models** below).

The following code can be used to extract the posterior probability
from a list object storing results for multiple feature-SNP pairs.

```
pp.mat <- sapply(X=res.list, FUN=get_pp)
```

In this case, the output is a matrix that contains rows and columns
corresponding to the models and the feature-SNP pairs, respectively.

#### Including random effects

In analyses of experimental data, it is ofen desirable to include
random effects in the model. To include donor random effects, we first
need to run `get_tu_lambda()`.

```
tu.lambda <- get_tu_lambda(data)
```

We then use the output as input when running
`do_bms()`.

```
res.ranef <- do_bms(
    data=data, p.m=p.m, phi=phi,
    fn="nonlinear", method="map.lap",
    tu.lambda=tu.lambda)
```

We can also include polygenic random effects rather than donor random
effects to accout for the genetic relatedness and popuplation structure.
See the function documentation (`?get_tu_lambda`) for
details.

#### Visualizing the results

##### Generating a genotype-phenotype (GP) plot

We recommend visually inspecting the model fit by generating scatter
plots, which we call a “GP” plot, for each feature-SNP pair (or SNP). To
make a GP plot, we first create a list of data frames using
`format_gp()`.

```
gp.plot <- format_gp(data=data, fit=res)
```

We then create a `ggplot2` object using
`make_gp_plot()`. The appearance of the plot can be modified
as usual.

```
p1 <- make_gp_plot(gp=gp.plot)
p1
```

##### Generating a posterior probability (PP) plot

It is also useful to generate a barplot of posterior probability,
which we call a “PP” plot. To make a PP plot, we first create a data
frame using `format_pp()`.

```
pp.plot <- format_pp(fit=res)
```

We then create a `ggplot2` object using
`make_pp_plot()`. The appearance of the plot can be modified
as usual.

```
p2 <- make_pp_plot(pp=pp.plot) 
p2
```

##### Heatmap of the posterior probability of the models

We can generate a heatmap to visualize posterior probability across
multiple feature-SNP pairs using `make_heatmap()`. Note that
we transpose the matrix using `t()` in the following
code.

```
p3 <- make_heatmap(t(pp.mat)) 
p3 + theme(legend.position="bottom",
           legend.title=element_blank())
```

As expected, we see that the highest probability tends to be assigned
to the correct (i.e., data-generating) model. That is, the MAP model
tends to be `"0,0,0"` for the first ten feature-SNP pairs in
the left-most columns, `"1,0,0"` for the 11-20th pairs, and
so forth.

#### Optimizing the effect prior hyperparameters

We recommend optimizing the effect prior hyperparmeters by an
empirical Bayes approach. In this approach, we obtain the \(\phi\_g\), \(\phi\_t\), and \(\phi\_{g \times t}\) hyperparameter values
that maximize the sum of \(\log\)-transformed marginal likelihood
across all feature-SNP pairs that are being analyzed. The most
conceptually straightforward method is to perform a grid search (see our
manuscript for details). We recommend using a workflow such as Snakemake. For
faster computation, we use MAP estimation and Laplace approximation,
setting `method="map.lap"` (see **Performing
BMS** above). From the BMS result for a feature-SNP pair, the
\(\log\) of the marginal likelihood can
be extracted as follows.

```
(ln.p.y <- res$ln.p.y)
#> [1] -252.8906
```

Since error bounds for Laplace approximation are not easily obtained,
we recommend rerunning MCMC and bridge sampling with optimal
hyperparameter values for the final result.

#### Session information

```
sessionInfo()
#> R version 4.1.2 (2021-11-01)
#> Platform: x86_64-apple-darwin17.0 (64-bit)
#> Running under: macOS Big Sur 10.16
#> 
#> Matrix products: default
#> BLAS:   /Library/Frameworks/R.framework/Versions/4.1/Resources/lib/libRblas.0.dylib
#> LAPACK: /Library/Frameworks/R.framework/Versions/4.1/Resources/lib/libRlapack.dylib
#> 
#> locale:
#> [1] en_US.UTF-8/en_US.UTF-8/en_US.UTF-8/C/en_US.UTF-8/en_US.UTF-8
#> 
#> attached base packages:
#> [1] stats     graphics  grDevices utils     datasets  methods   base     
#> 
#> other attached packages:
#> [1] classifygxt_0.0.1 ggplot2_3.4.4    
#> 
#> loaded via a namespace (and not attached):
#>  [1] matrixStats_0.61.0      fs_1.5.0                fontquiver_0.2.1       
#>  [4] rstan_2.21.5            tools_4.1.2             bslib_0.3.0            
#>  [7] utf8_1.2.2              R6_2.5.1                DBI_1.1.1              
#> [10] colorspace_2.0-2        withr_3.0.0             tidyselect_1.1.1       
#> [13] gridExtra_2.3           prettyunits_1.1.1       processx_3.8.1         
#> [16] Brobdingnag_1.2-7       curl_4.3.2              compiler_4.1.2         
#> [19] extrafontdb_1.0         textshaping_0.3.6       cli_3.6.1              
#> [22] desc_1.4.3              fontBitstreamVera_0.1.1 labeling_0.4.2         
#> [25] sass_0.4.6              scales_1.3.0            mvtnorm_1.1-3          
#> [28] callr_3.7.6             pkgdown_2.0.9           systemfonts_1.0.4      
#> [31] stringr_1.4.0           digest_0.6.27           StanHeaders_2.21.0-7   
#> [34] rmarkdown_2.11          gfonts_0.2.0            pkgconfig_2.0.3        
#> [37] htmltools_0.5.5         extrafont_0.19          highr_0.9              
#> [40] fastmap_1.1.1           htmlwidgets_1.5.4       rlang_1.1.1            
#> [43] rstudioapi_0.13         httpcode_0.3.0          shiny_1.7.1            
#> [46] farver_2.1.0            jquerylib_0.1.4         generics_0.1.2         
#> [49] jsonlite_1.7.2          dplyr_1.0.10            inline_0.3.19          
#> [52] magrittr_2.0.1          loo_2.5.1               Matrix_1.3-4           
#> [55] Rcpp_1.0.7              munsell_0.5.0           fansi_0.5.0            
#> [58] gdtools_0.3.3           viridis_0.6.1           lifecycle_1.0.4        
#> [61] stringi_1.7.4           yaml_2.2.1              pkgbuild_1.2.0         
#> [64] grid_4.1.2              hrbrthemes_0.8.0        parallel_4.1.2         
#> [67] promises_1.2.0.1        crayon_1.5.1            lattice_0.20-45        
#> [70] knitr_1.34              ps_1.6.0                pillar_1.8.1           
#> [73] codetools_0.2-18        stats4_4.1.2            rstantools_2.4.0       
#> [76] crul_1.2.0              glue_1.4.2              evaluate_0.14          
#> [79] fontLiberation_0.1.0    RcppParallel_5.1.5      vctrs_0.6.2            
#> [82] httpuv_1.6.3            Rttf2pt1_1.3.12         gtable_0.3.0           
#> [85] purrr_1.0.1             assertthat_0.2.1        cachem_1.0.6           
#> [88] xfun_0.26               mime_0.11               xtable_1.8-4           
#> [91] coda_0.19-4             later_1.3.0             ragg_1.2.5             
#> [94] viridisLite_0.4.0       tibble_3.1.8            memoise_2.0.1          
#> [97] ellipsis_0.3.2          bridgesampling_1.1-2
```

#### Contents

Developed by Yuriko Harigaya, Michael Love, William Valdar.

Site built with pkgdown 2.0.9.
