## Supplemental File 1 for "Probabilistic classification of gene-by-treatment interactions on molecular count phenotypes": index.html

Articles • classifygxt       

Toggle navigation


classifygxt
0.0.1

- Get started
- Reference
- Changelog

### Articles

##### All vignettes

ClassifyGxT - classifying gene-by-treatment interactions

Developed by Yuriko Harigaya, Michael Love, William Valdar.

Site built with pkgdown 2.0.9.
