## Supplemental File 1 for "Probabilistic classification of gene-by-treatment interactions on molecular count phenotypes": authors.html

Authors and Citation • classifygxt       

Toggle navigation


classifygxt
0.0.1

- Get started
- Reference
- Changelog

### Authors and Citation

- **Yuriko Harigaya**. Author, maintainer.
- **Michael Love**. Author.
- **William Valdar**. Author.

### Citation

Harigaya Y, Love M, Valdar W (2024).
*classifygxt: ClassifyGxT - classifying gene-by-treatment interactions*.
R package version 0.0.1.

```
@Manual{,
  title = {classifygxt: ClassifyGxT - classifying gene-by-treatment interactions},
  author = {Yuriko Harigaya and Michael Love and William Valdar},
  year = {2024},
  note = {R package version 0.0.1},
}
```

Developed by Yuriko Harigaya, Michael Love, William Valdar.

Site built with pkgdown 2.0.9.
