## Supplemental File 1 for "Probabilistic classification of gene-by-treatment interactions on molecular count phenotypes": bootstrap-toc.js

```
/*!
 * Bootstrap Table of Contents v0.4.1 (http://afeld.github.io/bootstrap-toc/)
 * Copyright 2015 Aidan Feldman
 * Licensed under MIT (https://github.com/afeld/bootstrap-toc/blob/gh-pages/LICENSE.md) */
(function() {
  'use strict';

  window.Toc = {
    helpers: {
      // return all matching elements in the set, or their descendants
      findOrFilter: function($el, selector) {
        // http://danielnouri.org/notes/2011/03/14/a-jquery-find-that-also-finds-the-root-element/
        // http://stackoverflow.com/a/12731439/358804
        var $descendants = $el.find(selector);
        return $el.filter(selector).add($descendants).filter(':not([data-toc-skip])');
      },

      generateUniqueIdBase: function(el) {
        var text = $(el).text();
        var anchor = text.trim().toLowerCase().replace(/[^A-Za-z0-9]+/g, '-');
        return anchor || el.tagName.toLowerCase();
      },

      generateUniqueId: function(el) {
        var anchorBase = this.generateUniqueIdBase(el);
        for (var i = 0; ; i++) {
          var anchor = anchorBase;
          if (i > 0) {
            // add suffix
            anchor += '-' + i;
          }
          // check if ID already exists
          if (!document.getElementById(anchor)) {
            return anchor;
          }
        }
      },

      generateAnchor: function(el) {
        if (el.id) {
          return el.id;
        } else {
          var anchor = this.generateUniqueId(el);
          el.id = anchor;
          return anchor;
        }
      },

      createNavList: function() {
        return $('<ul class="nav"></ul>');
      },

      createChildNavList: function($parent) {
        var $childList = this.createNavList();
        $parent.append($childList);
        return $childList;
      },

      generateNavEl: function(anchor, text) {
        var $a = $('<a></a>');
        $a.attr('href', '#' + anchor);
        $a.text(text);
        var $li = $('<li></li>');
        $li.append($a);
        return $li;
      },

      generateNavItem: function(headingEl) {
        var anchor = this.generateAnchor(headingEl);
        var $heading = $(headingEl);
        var text = $heading.data('toc-text') || $heading.text();
        return this.generateNavEl(anchor, text);
      },

      // Find the first heading level (`<h1>`, then `<h2>`, etc.) that has more than one element. Defaults to 1 (for `<h1>`).
      getTopLevel: function($scope) {
        for (var i = 1; i <= 6; i++) {
          var $headings = this.findOrFilter($scope, 'h' + i);
          if ($headings.length > 1) {
            return i;
          }
        }

        return 1;
      },

      // returns the elements for the top level, and the next below it
      getHeadings: function($scope, topLevel) {
        var topSelector = 'h' + topLevel;

        var secondaryLevel = topLevel + 1;
        var secondarySelector = 'h' + secondaryLevel;

        return this.findOrFilter($scope, topSelector + ',' + secondarySelector);
      },

      getNavLevel: function(el) {
        return parseInt(el.tagName.charAt(1), 10);
      },

      populateNav: function($topContext, topLevel, $headings) {
        var $context = $topContext;
        var $prevNav;

        var helpers = this;
        $headings.each(function(i, el) {
          var $newNav = helpers.generateNavItem(el);
          var navLevel = helpers.getNavLevel(el);

          // determine the proper $context
          if (navLevel === topLevel) {
            // use top level
            $context = $topContext;
          } else if ($prevNav && $context === $topContext) {
            // create a new level of the tree and switch to it
            $context = helpers.createChildNavList($prevNav);
          } // else use the current $context

          $context.append($newNav);

          $prevNav = $newNav;
        });
      },

      parseOps: function(arg) {
        var opts;
        if (arg.jquery) {
          opts = {
            $nav: arg
          };
        } else {
          opts = arg;
        }
        opts.$scope = opts.$scope || $(document.body);
        return opts;
      }
    },

    // accepts a jQuery object, or an options object
    init: function(opts) {
      opts = this.helpers.parseOps(opts);

      // ensure that the data attribute is in place for styling
      opts.$nav.attr('data-toggle', 'toc');

      var $topContext = this.helpers.createChildNavList(opts.$nav);
      var topLevel = this.helpers.getTopLevel(opts.$scope);
      var $headings = this.helpers.getHeadings(opts.$scope, topLevel);
      this.helpers.populateNav($topContext, topLevel, $headings);
    }
  };

  $(function() {
    $('nav[data-toggle="toc"]').each(function(i, el) {
      var $nav = $(el);
      Toc.init($nav);
    });
  });
})();
```
