## Supplemental File 1 for "Probabilistic classification of gene-by-treatment interactions on molecular count phenotypes": docsearch.js

```
$(function() {

  // register a handler to move the focus to the search bar
  // upon pressing shift + "/" (i.e. "?")
  $(document).on('keydown', function(e) {
    if (e.shiftKey && e.keyCode == 191) {
      e.preventDefault();
      $("#search-input").focus();
    }
  });

  $(document).ready(function() {
    // do keyword highlighting
    /* modified from https://jsfiddle.net/julmot/bL6bb5oo/ */
    var mark = function() {

      var referrer = document.URL ;
      var paramKey = "q" ;

      if (referrer.indexOf("?") !== -1) {
        var qs = referrer.substr(referrer.indexOf('?') + 1);
        var qs_noanchor = qs.split('#')[0];
        var qsa = qs_noanchor.split('&');
        var keyword = "";

        for (var i = 0; i < qsa.length; i++) {
          var currentParam = qsa[i].split('=');

          if (currentParam.length !== 2) {
            continue;
          }

          if (currentParam[0] == paramKey) {
            keyword = decodeURIComponent(currentParam[1].replace(/\+/g, "%20"));
          }
        }

        if (keyword !== "") {
          $(".contents").unmark({
            done: function() {
              $(".contents").mark(keyword);
            }
          });
        }
      }
    };

    mark();
  });
});

/* Search term highlighting ------------------------------*/

function matchedWords(hit) {
  var words = [];

  var hierarchy = hit._highlightResult.hierarchy;
  // loop to fetch from lvl0, lvl1, etc.
  for (var idx in hierarchy) {
    words = words.concat(hierarchy[idx].matchedWords);
  }

  var content = hit._highlightResult.content;
  if (content) {
    words = words.concat(content.matchedWords);
  }

  // return unique words
  var words_uniq = [...new Set(words)];
  return words_uniq;
}

function updateHitURL(hit) {

  var words = matchedWords(hit);
  var url = "";

  if (hit.anchor) {
    url = hit.url_without_anchor + '?q=' + escape(words.join(" ")) + '#' + hit.anchor;
  } else {
    url = hit.url + '?q=' + escape(words.join(" "));
  }

  return url;
}
```
