## Supplemental File 1 for "Probabilistic classification of gene-by-treatment interactions on molecular count phenotypes": index.html

ClassifyGxT - classifying gene-by-treatment interactions • classifygxt
 


  


 


Toggle navigation


classifygxt
0.0.1

- Get started
- Reference
- Changelog

### ClassifyGxT

#### About

ClassifyGxT is a method for classifying gene-by-treatment (GxT) interactions using Bayesian model selection (BMS). The method is primarily designed for molecular count phenotypes, such as gene expression and chromatin accessibility, although it can be used for other types of phenotypes. It takes as input a list of feature-SNP pairs (or SNPs) with significant GxT interactions that have already been identified by a standard method and assigns posterior probability of different types of GxT interactions to each of the feature-SNP pairs (or SNPs). See **Overview** below for the types of GxT interactions.

See `Get started` and `Reference` for usage instructions.

#### Publication

Yuriko Harigaya, Nana Matoba, Brandon D. Le Jordan M. Valone Jason L. Stein Michael I. Love\* William Valdar\*. “Bayesian model selection for classifying gene-by-treatment interactions on molecular count phenotypes.” (\* These authors contributed equally to this work.)

#### Overview

In the simplest form, our BMS framework can be described as follows. For *n* samples indexed by *i*, we consider a linear model shown below. *y*_*i*_ is the phenotype, *g*_*i*_ is the genotype, and *t*_*i*_ is the indicator variable as to whether the sample *i* was treated. *ε*_*i*_ is the residual error (noise) and is assumed to follow a Normal distributiion with a zero mean and variance *σ*^2^. *β*_0_ is the intercept. *β*_*g*_, *β*_*t*_, and *β*_*g* × *t*_ are the genotype, treatment, and GxT interaction effects, respectively. We use the *m*_*g*_, *m*_*t*_, and *m*_*g* × *t*_ indicator variables to specify the inclusion and exclusion of the genotype, treatment, and GxT interaction terms, respectively. The **m**_*j*_ vector refers to the *j*-th model (*j* = 1, …, 8). The eight models represent different types of GxT interactions.


To account for the inherent nonlinear relationship between the genotype and the tranformed molecular count phenotype, we use nonlinear regression. The eight models can be formulated in the same way using the **m** vectors. See our manuscript (**Publication**) for prior specification, posterior computation, and other details.

Although we primarily use the **m** vector notations to specify the types of GxT interaction, it is also useful to consider the following nomenclatures.

| **m** vector | Nomenclature |
| --- | --- |
| (0,0,0) | No effect |
| (1,0,0) | Genotype main effect only |
| (0,1,0) | Treatment main effect only |
| (1,1,0) | Genotype and treatment main effects only |
| (0,0,1) | Treatment-induced genotype effect |
| (1,0,1) | Treatment-altered genotype effect |
| (0,1,1) | Treatment main effect and treatment-induced genotype effect |
| (1,1,1) | Genotype and treatment main effects and interaction |

#### Installation

The *classifygxt* R package can be installed using *devtools*.

```
if (!requireNamespace("devtools", quietly = TRUE)) install.packages("devtools")
devtools::install_github("yharigaya/classifygxt")
```

#### License

- Full license
- GPL (>= 3)

#### Citation

- Citing classifygxt

#### Developers

- Yuriko Harigaya   
   Author, maintainer
- Michael Love   
   Author
- William Valdar   
   Author

Developed by Yuriko Harigaya, Michael Love, William Valdar.

Site built with pkgdown 2.0.9.
