## Supplemental File 1 for "Probabilistic classification of gene-by-treatment interactions on molecular count phenotypes": LICENSE.html

GNU General Public License • classifygxt       

Toggle navigation


classifygxt
0.0.1

- Get started
- Reference
- Changelog

### GNU General Public License

*Version 3, 29 June 2007*  
*Copyright © 2007 Free Software Foundation, Inc. <http://fsf.org/>*

Everyone is permitted to copy and distribute verbatim copies of this license document, but changing it is not allowed.

#### Preamble

The GNU General Public License is a free, copyleft license for software and other kinds of works.

The licenses for most software and other practical works are designed to take away your freedom to share and change the works. By contrast, the GNU General Public License is intended to guarantee your freedom to share and change all versions of a program–to make sure it remains free software for all its users. We, the Free Software Foundation, use the GNU General Public License for most of our software; it applies also to any other work released this way by its authors. You can apply it to your programs, too.

When we speak of free software, we are referring to freedom, not price. Our General Public Licenses are designed to make sure that you have the freedom to distribute copies of free software (and charge for them if you wish), that you receive source code or can get it if you want it, that you can change the software or use pieces of it in new free programs, and that you know you can do these things.

To protect your rights, we need to prevent others from denying you these rights or asking you to surrender the rights. Therefore, you have certain responsibilities if you distribute copies of the software, or if you modify it: responsibilities to respect the freedom of others.

For example, if you distribute copies of such a program, whether gratis or for a fee, you must pass on to the recipients the same freedoms that you received. You must make sure that they, too, receive or can get the source code. And you must show them these terms so they know their rights.

Developers that use the GNU GPL protect your rights with two steps: **(1)** assert copyright on the software, and **(2)** offer you this License giving you legal permission to copy, distribute and/or modify it.

For the developers’ and authors’ protection, the GPL clearly explains that there is no warranty for this free software. For both users’ and authors’ sake, the GPL requires that modified versions be marked as changed, so that their problems will not be attributed erroneously to authors of previous versions.

Some devices are designed to deny users access to install or run modified versions of the software inside them, although the manufacturer can do so. This is fundamentally incompatible with the aim of protecting users’ freedom to change the software. The systematic pattern of such abuse occurs in the area of products for individuals to use, which is precisely where it is most unacceptable. Therefore, we have designed this version of the GPL to prohibit the practice for those products. If such problems arise substantially in other domains, we stand ready to extend this provision to those domains in future versions of the GPL, as needed to protect the freedom of users.

Finally, every program is threatened constantly by software patents. States should not allow patents to restrict development and use of software on general-purpose computers, but in those that do, we wish to avoid the special danger that patents applied to a free program could make it effectively proprietary. To prevent this, the GPL assures that patents cannot be used to render the program non-free.

The precise terms and conditions for copying, distribution and modification follow.

#### TERMS AND CONDITIONS

##### 0. Definitions

“This License” refers to version 3 of the GNU General Public License.

“Copyright” also means copyright-like laws that apply to other kinds of works, such as semiconductor masks.

“The Program” refers to any copyrightable work licensed under this License. Each licensee is addressed as “you”. “Licensees” and “recipients” may be individuals or organizations.

To “modify” a work means to copy from or adapt all or part of the work in a fashion requiring copyright permission, other than the making of an exact copy. The resulting work is called a “modified version” of the earlier work or a work “based on” the earlier work.

A “covered work” means either the unmodified Program or a work based on the Program.

To “propagate” a work means to do anything with it that, without permission, would make you directly or secondarily liable for infringement under applicable copyright law, except executing it on a computer or modifying a private copy. Propagation includes copying, distribution (with or without modification), making available to the public, and in some countries other activities as well.

To “convey” a work means any kind of propagation that enables other parties to make or receive copies. Mere interaction with a user through a computer network, with no transfer of a copy, is not conveying.

An interactive user interface displays “Appropriate Legal Notices” to the extent that it includes a convenient and prominently visible feature that **(1)** displays an appropriate copyright notice, and **(2)** tells the user that there is no warranty for the work (except to the extent that warranties are provided), that licensees may convey the work under this License, and how to view a copy of this License. If the interface presents a list of user commands or options, such as a menu, a prominent item in the list meets this criterion.

##### 1. Source Code

The “source code” for a work means the preferred form of the work for making modifications to it. “Object code” means any non-source form of a work.

A “Standard Interface” means an interface that either is an official standard defined by a recognized standards body, or, in the case of interfaces specified for a particular programming language, one that is widely used among developers working in that language.

The “System Libraries” of an executable work include anything, other than the work as a whole, that **(a)** is included in the normal form of packaging a Major Component, but which is not part of that Major Component, and **(b)** serves only to enable use of the work with that Major Component, or to implement a Standard Interface for which an implementation is available to the public in source code form. A “Major Component”, in this context, means a major essential component (kernel, window system, and so on) of the specific operating system (if any) on which the executable work runs, or a compiler used to produce the work, or an object code interpreter used to run it.

The “Corresponding Source” for a work in object code form means all the source code needed to generate, install, and (for an executable work) run the object code and to modify the work, including scripts to control those activities. However, it does not include the work’s System Libraries, or general-purpose tools or generally available free programs which are used unmodified in performing those activities but which are not part of the work. For example, Corresponding Source includes interface definition files associated with source files for the work, and the source code for shared libraries and dynamically linked subprograms that the work is specifically designed to require, such as by intimate data communication or control flow between those subprograms and other parts of the work.

The Corresponding Source need not include anything that users can regenerate automatically from other parts of the Corresponding Source.

The Corresponding Source for a work in source code form is that same work.

##### 2. Basic Permissions

All rights granted under this License are granted for the term of copyright on the Program, and are irrevocable provided the stated conditions are met. This License explicitly affirms your unlimited permission to run the unmodified Program. The output from running a covered work is covered by this License only if the output, given its content, constitutes a covered work. This License acknowledges your rights of fair use or other equivalent, as provided by copyright law.

You may make, run and propagate covered works that you do not convey, without conditions so long as your license otherwise remains in force. You may convey covered works to others for the sole purpose of having them make modifications exclusively for you, or provide you with facilities for running those works, provided that you comply with the terms of this License in conveying all material for which you do not control copyright. Those thus making or running the covered works for you must do so exclusively on your behalf, under your direction and control, on terms that prohibit them from making any copies of your copyrighted material outside their relationship with you.

Conveying under any other circumstances is permitted solely under the conditions stated below. Sublicensing is not allowed; section 10 makes it unnecessary.

##### 3. Protecting Users’ Legal Rights From Anti-Circumvention Law

No covered work shall be deemed part of an effective technological measure under any applicable law fulfilling obligations under article 11 of the WIPO copyright treaty adopted on 20 December 1996, or similar laws prohibiting or restricting circumvention of such measures.

When you convey a covered work, you waive any legal power to forbid circumvention of technological measures to the extent such circumvention is effected by exercising rights under this License with respect to the covered work, and you disclaim any intention to limit operation or modification of the work as a means of enforcing, against the work’s users, your or third parties’ legal rights to forbid circumvention of technological measures.

##### 4. Conveying Verbatim Copies

You may convey verbatim copies of the Program’s source code as you receive it, in any medium, provided that you conspicuously and appropriately publish on each copy an appropriate copyright notice; keep intact all notices stating that this License and any non-permissive terms added in accord with section 7 apply to the code; keep intact all notices of the absence of any warranty; and give all recipients a copy of this License along with the Program.

You may charge any price or no price for each copy that you convey, and you may offer support or warranty protection for a fee.

##### 5. Conveying Modified Source Versions

You may convey a work based on the Program, or the modifications to produce it from the Program, in the form of source code under the terms of section 4, provided that you also meet all of these conditions:

- **a)** The work must carry prominent notices stating that you modified it, and giving a relevant date.
- **b)** The work must carry prominent notices stating that it is released under this License and any conditions added under section 7. This requirement modifies the requirement in section 4 to “keep intact all notices”.
- **c)** You must license the entire work, as a whole, under this License to anyone who comes into possession of a copy. This License will therefore apply, along with any applicable section 7 additional terms, to the whole of the work, and all its parts, regardless of how they are packaged. This License gives no permission to license the work in any other way, but it does not invalidate such permission if you have separately received it.
- **d)** If the work has interactive user interfaces, each must display Appropriate Legal Notices; however, if the Program has interactive interfaces that do not display Appropriate Legal Notices, your work need not make them do so.

A compilation of a covered work with other separate and independent works, which are not by their nature extensions of the covered work, and which are not combined with it such as to form a larger program, in or on a volume of a storage or distribution medium, is called an “aggregate” if the compilation and its resulting copyright are not used to limit the access or legal rights of the compilation’s users beyond what the individual works permit. Inclusion of a covered work in an aggregate does not cause this License to apply to the other parts of the aggregate.

##### 6. Conveying Non-Source Forms

You may convey a covered work in object code form under the terms of sections 4 and 5, provided that you also convey the machine-readable Corresponding Source under the terms of this License, in one of these ways:

- **a)** Convey the object code in, or embodied in, a physical product (including a physical distribution medium), accompanied by the Corresponding Source fixed on a durable physical medium customarily used for software interchange.
- **b)** Convey the object code in, or embodied in, a physical product (including a physical distribution medium), accompanied by a written offer, valid for at least three years and valid for as long as you offer spare parts or customer support for that product model, to give anyone who possesses the object code either **(1)** a copy of the Corresponding Source for all the software in the product that is covered by this License, on a durable physical medium customarily used for software interchange, for a price no more than your reasonable cost of physically performing this conveying of source, or **(2)** access to copy the Corresponding Source from a network server at no charge.
- **c)** Convey individual copies of the object code with a copy of the written offer to provide the Corresponding Source. This alternative is allowed only occasionally and noncommercially, and only if you received the object code with such an offer, in accord with subsection 6b.
- **d)** Convey the object code by offering access from a designated place (gratis or for a charge), and offer equivalent access to the Corresponding Source in the same way through the same place at no further charge. You need not require recipients to copy the Corresponding Source along with the object code. If the place to copy the object code is a network server, the Corresponding Source may be on a different server (operated by you or a third party) that supports equivalent copying facilities, provided you maintain clear directions next to the object code saying where to find the Corresponding Source. Regardless of what server hosts the Corresponding Source, you remain obligated to ensure that it is available for as long as needed to satisfy these requirements.
- **e)** Convey the object code using peer-to-peer transmission, provided you inform other peers where the object code and Corresponding Source of the work are being offered to the general public at no charge under subsection 6d.

A separable portion of the object code, whose source code is excluded from the Corresponding Source as a System Library, need not be included in conveying the object code work.

A “User Product” is either **(1)** a “consumer product”, which means any tangible personal property which is normally used for personal, family, or household purposes, or **(2)** anything designed or sold for incorporation into a dwelling. In determining whether a product is a consumer product, doubtful cases shall be resolved in favor of coverage. For a particular product received by a particular user, “normally used” refers to a typical or common use of that class of product, regardless of the status of the particular user or of the way in which the particular user actually uses, or expects or is expected to use, the product. A product is a consumer product regardless of whether the product has substantial commercial, industrial or non-consumer uses, unless such uses represent the only significant mode of use of the product.

“Installation Information” for a User Product means any methods, procedures, authorization keys, or other information required to install and execute modified versions of a covered work in that User Product from a modified version of its Corresponding Source. The information must suffice to ensure that the continued functioning of the modified object code is in no case prevented or interfered with solely because modification has been made.

If you convey an object code work under this section in, or with, or specifically for use in, a User Product, and the conveying occurs as part of a transaction in which the right of possession and use of the User Product is transferred to the recipient in perpetuity or for a fixed term (regardless of how the transaction is characterized), the Corresponding Source conveyed under this section must be accompanied by the Installation Information. But this requirement does not apply if neither you nor any third party retains the ability to install modified object code on the User Product (for example, the work has been installed in ROM).

The requirement to provide Installation Information does not include a requirement to continue to provide support service, warranty, or updates for a work that has been modified or installed by the recipient, or for the User Product in which it has been modified or installed. Access to a network may be denied when the modification itself materially and adversely affects the operation of the network or violates the rules and protocols for communication across the network.

Corresponding Source conveyed, and Installation Information provided, in accord with this section must be in a format that is publicly documented (and with an implementation available to the public in source code form), and must require no special password or key for unpacking, reading or copying.

##### 7. Additional Terms

“Additional permissions” are terms that supplement the terms of this License by making exceptions from one or more of its conditions. Additional permissions that are applicable to the entire Program shall be treated as though they were included in this License, to the extent that they are valid under applicable law. If additional permissions apply only to part of the Program, that part may be used separately under those permissions, but the entire Program remains governed by this License without regard to the additional permissions.

When you convey a copy of a covered work, you may at your option remove any additional permissions from that copy, or from any part of it. (Additional permissions may be written to require their own removal in certain cases when you modify the work.) You may place additional permissions on material, added by you to a covered work, for which you have or can give appropriate copyright permission.

Notwithstanding any other provision of this License, for material you add to a covered work, you may (if authorized by the copyright holders of that material) supplement the terms of this License with terms:

- **a)** Disclaiming warranty or limiting liability differently from the terms of sections 15 and 16 of this License; or
- **b)** Requiring preservation of specified reasonable legal notices or author attributions in that material or in the Appropriate Legal Notices displayed by works containing it; or
- **c)** Prohibiting misrepresentation of the origin of that material, or requiring that modified versions of such material be marked in reasonable ways as different from the original version; or
- **d)** Limiting the use for publicity purposes of names of licensors or authors of the material; or
- **e)** Declining to grant rights under trademark law for use of some trade names, trademarks, or service marks; or
- **f)** Requiring indemnification of licensors and authors of that material by anyone who conveys the material (or modified versions of it) with contractual assumptions of liability to the recipient, for any liability that these contractual assumptions directly impose on those licensors and authors.

All other non-permissive additional terms are considered “further restrictions” within the meaning of section 10. If the Program as you received it, or any part of it, contains a notice stating that it is governed by this License along with a term that is a further restriction, you may remove that term. If a license document contains a further restriction but permits relicensing or conveying under this License, you may add to a covered work material governed by the terms of that license document, provided that the further restriction does not survive such relicensing or conveying.

If you add terms to a covered work in accord with this section, you must place, in the relevant source files, a statement of the additional terms that apply to those files, or a notice indicating where to find the applicable terms.

Additional terms, permissive or non-permissive, may be stated in the form of a separately written license, or stated as exceptions; the above requirements apply either way.

##### 8. Termination

You may not propagate or modify a covered work except as expressly provided under this License. Any attempt otherwise to propagate or modify it is void, and will automatically terminate your rights under this License (including any patent licenses granted under the third paragraph of section 11).

However, if you cease all violation of this License, then your license from a particular copyright holder is reinstated **(a)** provisionally, unless and until the copyright holder explicitly and finally terminates your license, and **(b)** permanently, if the copyright holder fails to notify you of the violation by some reasonable means prior to 60 days after the cessation.

Moreover, your license from a particular copyright holder is reinstated permanently if the copyright holder notifies you of the violation by some reasonable means, this is the first time you have received notice of violation of this License (for any work) from that copyright holder, and you cure the violation prior to 30 days after your receipt of the notice.

Termination of your rights under this section does not terminate the licenses of parties who have received copies or rights from you under this License. If your rights have been terminated and not permanently reinstated, you do not qualify to receive new licenses for the same material under section 10.

##### 9. Acceptance Not Required for Having Copies

You are not required to accept this License in order to receive or run a copy of the Program. Ancillary propagation of a covered work occurring solely as a consequence of using peer-to-peer transmission to receive a copy likewise does not require acceptance. However, nothing other than this License grants you permission to propagate or modify any covered work. These actions infringe copyright if you do not accept this License. Therefore, by modifying or propagating a covered work, you indicate your acceptance of this License to do so.

##### 10. Automatic Licensing of Downstream Recipients

Each time you convey a covered work, the recipient automatically receives a license from the original licensors, to run, modify and propagate that work, subject to this License. You are not responsible for enforcing compliance by third parties with this License.

An “entity transaction” is a transaction transferring control of an organization, or substantially all assets of one, or subdividing an organization, or merging organizations. If propagation of a covered work results from an entity transaction, each party to that transaction who receives a copy of the work also receives whatever licenses to the work the party’s predecessor in interest had or could give under the previous paragraph, plus a right to possession of the Corresponding Source of the work from the predecessor in interest, if the predecessor has it or can get it with reasonable efforts.

You may not impose any further restrictions on the exercise of the rights granted or affirmed under this License. For example, you may not impose a license fee, royalty, or other charge for exercise of rights granted under this License, and you may not initiate litigation (including a cross-claim or counterclaim in a lawsuit) alleging that any patent claim is infringed by making, using, selling, offering for sale, or importing the Program or any portion of it.

##### 11. Patents

A “contributor” is a copyright holder who authorizes use under this License of the Program or a work on which the Program is based. The work thus licensed is called the contributor’s “contributor version”.

A contributor’s “essential patent claims” are all patent claims owned or controlled by the contributor, whether already acquired or hereafter acquired, that would be infringed by some manner, permitted by this License, of making, using, or selling its contributor version, but do not include claims that would be infringed only as a consequence of further modification of the contributor version. For purposes of this definition, “control” includes the right to grant patent sublicenses in a manner consistent with the requirements of this License.

Each contributor grants you a non-exclusive, worldwide, royalty-free patent license under the contributor’s essential patent claims, to make, use, sell, offer for sale, import and otherwise run, modify and propagate the contents of its contributor version.

In the following three paragraphs, a “patent license” is any express agreement or commitment, however denominated, not to enforce a patent (such as an express permission to practice a patent or covenant not to sue for patent infringement). To “grant” such a patent license to a party means to make such an agreement or commitment not to enforce a patent against the party.

If you convey a covered work, knowingly relying on a patent license, and the Corresponding Source of the work is not available for anyone to copy, free of charge and under the terms of this License, through a publicly available network server or other readily accessible means, then you must either **(1)** cause the Corresponding Source to be so available, or **(2)** arrange to deprive yourself of the benefit of the patent license for this particular work, or **(3)** arrange, in a manner consistent with the requirements of this License, to extend the patent license to downstream recipients. “Knowingly relying” means you have actual knowledge that, but for the patent license, your conveying the covered work in a country, or your recipient’s use of the covered work in a country, would infringe one or more identifiable patents in that country that you have reason to believe are valid.

If, pursuant to or in connection with a single transaction or arrangement, you convey, or propagate by procuring conveyance of, a covered work, and grant a patent license to some of the parties receiving the covered work authorizing them to use, propagate, modify or convey a specific copy of the covered work, then the patent license you grant is automatically extended to all recipients of the covered work and works based on it.

A patent license is “discriminatory” if it does not include within the scope of its coverage, prohibits the exercise of, or is conditioned on the non-exercise of one or more of the rights that are specifically granted under this License. You may not convey a covered work if you are a party to an arrangement with a third party that is in the business of distributing software, under which you make payment to the third party based on the extent of your activity of conveying the work, and under which the third party grants, to any of the parties who would receive the covered work from you, a discriminatory patent license **(a)** in connection with copies of the covered work conveyed by you (or copies made from those copies), or **(b)** primarily for and in connection with specific products or compilations that contain the covered work, unless you entered into that arrangement, or that patent license was granted, prior to 28 March 2007.

Nothing in this License shall be construed as excluding or limiting any implied license or other defenses to infringement that may otherwise be available to you under applicable patent law.

##### 12. No Surrender of Others’ Freedom

If conditions are imposed on you (whether by court order, agreement or otherwise) that contradict the conditions of this License, they do not excuse you from the conditions of this License. If you cannot convey a covered work so as to satisfy simultaneously your obligations under this License and any other pertinent obligations, then as a consequence you may not convey it at all. For example, if you agree to terms that obligate you to collect a royalty for further conveying from those to whom you convey the Program, the only way you could satisfy both those terms and this License would be to refrain entirely from conveying the Program.

##### 13. Use with the GNU Affero General Public License

Notwithstanding any other provision of this License, you have permission to link or combine any covered work with a work licensed under version 3 of the GNU Affero General Public License into a single combined work, and to convey the resulting work. The terms of this License will continue to apply to the part which is the covered work, but the special requirements of the GNU Affero General Public License, section 13, concerning interaction through a network will apply to the combination as such.

##### 14. Revised Versions of this License

The Free Software Foundation may publish revised and/or new versions of the GNU General Public License from time to time. Such new versions will be similar in spirit to the present version, but may differ in detail to address new problems or concerns.

Each version is given a distinguishing version number. If the Program specifies that a certain numbered version of the GNU General Public License “or any later version” applies to it, you have the option of following the terms and conditions either of that numbered version or of any later version published by the Free Software Foundation. If the Program does not specify a version number of the GNU General Public License, you may choose any version ever published by the Free Software Foundation.

If the Program specifies that a proxy can decide which future versions of the GNU General Public License can be used, that proxy’s public statement of acceptance of a version permanently authorizes you to choose that version for the Program.

Later license versions may give you additional or different permissions. However, no additional obligations are imposed on any author or copyright holder as a result of your choosing to follow a later version.

##### 15. Disclaimer of Warranty

THERE IS NO WARRANTY FOR THE PROGRAM, TO THE EXTENT PERMITTED BY APPLICABLE LAW. EXCEPT WHEN OTHERWISE STATED IN WRITING THE COPYRIGHT HOLDERS AND/OR OTHER PARTIES PROVIDE THE PROGRAM “AS IS” WITHOUT WARRANTY OF ANY KIND, EITHER EXPRESSED OR IMPLIED, INCLUDING, BUT NOT LIMITED TO, THE IMPLIED WARRANTIES OF MERCHANTABILITY AND FITNESS FOR A PARTICULAR PURPOSE. THE ENTIRE RISK AS TO THE QUALITY AND PERFORMANCE OF THE PROGRAM IS WITH YOU. SHOULD THE PROGRAM PROVE DEFECTIVE, YOU ASSUME THE COST OF ALL NECESSARY SERVICING, REPAIR OR CORRECTION.

##### 16. Limitation of Liability

IN NO EVENT UNLESS REQUIRED BY APPLICABLE LAW OR AGREED TO IN WRITING WILL ANY COPYRIGHT HOLDER, OR ANY OTHER PARTY WHO MODIFIES AND/OR CONVEYS THE PROGRAM AS PERMITTED ABOVE, BE LIABLE TO YOU FOR DAMAGES, INCLUDING ANY GENERAL, SPECIAL, INCIDENTAL OR CONSEQUENTIAL DAMAGES ARISING OUT OF THE USE OR INABILITY TO USE THE PROGRAM (INCLUDING BUT NOT LIMITED TO LOSS OF DATA OR DATA BEING RENDERED INACCURATE OR LOSSES SUSTAINED BY YOU OR THIRD PARTIES OR A FAILURE OF THE PROGRAM TO OPERATE WITH ANY OTHER PROGRAMS), EVEN IF SUCH HOLDER OR OTHER PARTY HAS BEEN ADVISED OF THE POSSIBILITY OF SUCH DAMAGES.

##### 17. Interpretation of Sections 15 and 16

If the disclaimer of warranty and limitation of liability provided above cannot be given local legal effect according to their terms, reviewing courts shall apply local law that most closely approximates an absolute waiver of all civil liability in connection with the Program, unless a warranty or assumption of liability accompanies a copy of the Program in return for a fee.

*END OF TERMS AND CONDITIONS*

#### How to Apply These Terms to Your New Programs

If you develop a new program, and you want it to be of the greatest possible use to the public, the best way to achieve this is to make it free software which everyone can redistribute and change under these terms.

To do so, attach the following notices to the program. It is safest to attach them to the start of each source file to most effectively state the exclusion of warranty; and each file should have at least the “copyright” line and a pointer to where the full notice is found.

```
<one line to give the program's name and a brief idea of what it does.>
Copyright (C) <year>  <name of author>

This program is free software: you can redistribute it and/or modify
it under the terms of the GNU General Public License as published by
the Free Software Foundation, either version 3 of the License, or
(at your option) any later version.

This program is distributed in the hope that it will be useful,
but WITHOUT ANY WARRANTY; without even the implied warranty of
MERCHANTABILITY or FITNESS FOR A PARTICULAR PURPOSE.  See the
GNU General Public License for more details.

You should have received a copy of the GNU General Public License
along with this program.  If not, see <http://www.gnu.org/licenses/>.
```

Also add information on how to contact you by electronic and paper mail.

If the program does terminal interaction, make it output a short notice like this when it starts in an interactive mode:

```
<program>  Copyright (C) <year>  <name of author>
This program comes with ABSOLUTELY NO WARRANTY; for details type 'show w'.
This is free software, and you are welcome to redistribute it
under certain conditions; type 'show c' for details.
```

The hypothetical commands `show w` and `show c` should show the appropriate parts of the General Public License. Of course, your program’s commands might be different; for a GUI interface, you would use an “about box”.

You should also get your employer (if you work as a programmer) or school, if any, to sign a “copyright disclaimer” for the program, if necessary. For more information on this, and how to apply and follow the GNU GPL, see <http://www.gnu.org/licenses/>.

The GNU General Public License does not permit incorporating your program into proprietary programs. If your program is a subroutine library, you may consider it more useful to permit linking proprietary applications with the library. If this is what you want to do, use the GNU Lesser General Public License instead of this License. But first, please read <http://www.gnu.org/philosophy/why-not-lgpl.html>.

#### Contents

Developed by Yuriko Harigaya, Michael Love, William Valdar.

Site built with pkgdown 2.0.9.
