## Supplemental File 1 for "Probabilistic classification of gene-by-treatment interactions on molecular count phenotypes": index.html

Changelog • classifygxt       

Toggle navigation


classifygxt
0.0.1

- Get started
- Reference
- Changelog

### Changelog

#### classifygxt 0.0.1

- Added a `NEWS.md` file to track changes to the package.

#### Contents

Developed by Yuriko Harigaya, Michael Love, William Valdar.

Site built with pkgdown 2.0.9.
