## Supplemental File 1 for "Probabilistic classification of gene-by-treatment interactions on molecular count phenotypes": pkgdown.js

```
/* http://gregfranko.com/blog/jquery-best-practices/ */
(function($) {
  $(function() {

    $('.navbar-fixed-top').headroom();

    $('body').css('padding-top', $('.navbar').height() + 10);
    $(window).resize(function(){
      $('body').css('padding-top', $('.navbar').height() + 10);
    });

    $('[data-toggle="tooltip"]').tooltip();

    var cur_path = paths(location.pathname);
    var links = $("#navbar ul li a");
    var max_length = -1;
    var pos = -1;
    for (var i = 0; i < links.length; i++) {
      if (links[i].getAttribute("href") === "#")
        continue;
      // Ignore external links
      if (links[i].host !== location.host)
        continue;

      var nav_path = paths(links[i].pathname);

      var length = prefix_length(nav_path, cur_path);
      if (length > max_length) {
        max_length = length;
        pos = i;
      }
    }

    // Add class to parent <li>, and enclosing <li> if in dropdown
    if (pos >= 0) {
      var menu_anchor = $(links[pos]);
      menu_anchor.parent().addClass("active");
      menu_anchor.closest("li.dropdown").addClass("active");
    }
  });

  function paths(pathname) {
    var pieces = pathname.split("/");
    pieces.shift(); // always starts with /

    var end = pieces[pieces.length - 1];
    if (end === "index.html" || end === "")
      pieces.pop();
    return(pieces);
  }

  // Returns -1 if not found
  function prefix_length(needle, haystack) {
    if (needle.length > haystack.length)
      return(-1);

    // Special case for length-0 haystack, since for loop won't run
    if (haystack.length === 0) {
      return(needle.length === 0 ? 0 : -1);
    }

    for (var i = 0; i < haystack.length; i++) {
      if (needle[i] != haystack[i])
        return(i);
    }

    return(haystack.length);
  }

  /* Clipboard --------------------------*/

  function changeTooltipMessage(element, msg) {
    var tooltipOriginalTitle=element.getAttribute('data-original-title');
    element.setAttribute('data-original-title', msg);
    $(element).tooltip('show');
    element.setAttribute('data-original-title', tooltipOriginalTitle);
  }

  if(ClipboardJS.isSupported()) {
    $(document).ready(function() {
      var copyButton = "<button type='button' class='btn btn-primary btn-copy-ex' type = 'submit' title='Copy to clipboard' aria-label='Copy to clipboard' data-toggle='tooltip' data-placement='left auto' data-trigger='hover' data-clipboard-copy><i class='fa fa-copy'></i></button>";

      $("div.sourceCode").addClass("hasCopyButton");

      // Insert copy buttons:
      $(copyButton).prependTo(".hasCopyButton");

      // Initialize tooltips:
      $('.btn-copy-ex').tooltip({container: 'body'});

      // Initialize clipboard:
      var clipboardBtnCopies = new ClipboardJS('[data-clipboard-copy]', {
        text: function(trigger) {
          return trigger.parentNode.textContent.replace(/\n#>[^\n]*/g, "");
        }
      });

      clipboardBtnCopies.on('success', function(e) {
        changeTooltipMessage(e.trigger, 'Copied!');
        e.clearSelection();
      });

      clipboardBtnCopies.on('error', function() {
        changeTooltipMessage(e.trigger,'Press Ctrl+C or Command+C to copy');
      });
    });
  }
})(window.jQuery || window.$)
```
