## Supplemental File 1 for "Probabilistic classification of gene-by-treatment interactions on molecular count phenotypes": classifygxt-package.html

The 'classifygxt' package. — classifygxt-package • classifygxt       

Toggle navigation


classifygxt
0.0.1

- Get started
- Reference
- Changelog

### The 'classifygxt' package.

`classifygxt-package.Rd`

A DESCRIPTION OF THE PACKAGE

#### Contents

Developed by Yuriko Harigaya, Michael Love, William Valdar.

Site built with pkgdown 2.0.9.
