## Supplemental File 1 for "Probabilistic classification of gene-by-treatment interactions on molecular count phenotypes": do_bms.html

Perform BMS for classifying GxT interactions — do\_bms • classifygxt       

Toggle navigation


classifygxt
0.0.1

- Get started
- Reference
- Changelog

### Perform BMS for classifying GxT interactions

`do_bms.Rd`

This function takes as input individual phenotype and genotype data
and performs BMS for a feature-SNP pair (or a SNP). The output
includes posterior probabilities of different types of GxT
interactions as well as the log marginal likelihood, which can be
used for optimizing the hyperparameter values by an empirical Bayes
approach. BMS is performed by either MCMC followed by bridge
sampling or MAP estimation followed by Laplace approximation. In
the latter case, a relative marginal likelihood is returned. That
is, the log marginal likelihood value is shifted by a
constant. Note that this shift does not affect the posterior
probabilities or hyperparameter optimization.
`tu.lambda` must be supplied when modeling a random effect.

```
do_bms(
  data,
  fn,
  rint = FALSE,
  method,
  p.m = NULL,
  phi,
  phi0 = sqrt(1000),
  kappa = 0.002,
  nu = 0.002,
  kappa.u = 0.002,
  nu.u = 0.002,
  tu.lambda = NULL,
  summary = TRUE,
  seed = 1,
  n.cores = 4
)
```

#### Arguments

data
:   A list containing vectors of subjects, genotypes,
    phenotypes, and treatment indicators, which must be named
    "subject", "g", "y", and "t", respectively.

fn
:   A character string specifying the function. This must be
    one of "nonlinear" and "linear", corresponding to nonlinear and
    linear models, respectively.

rint
:   A logical as to whether the phenotypes need to be
    RINT-transformed. This cannot be set to `TRUE` if `fn` is
    set to "nonlinear".

method
:   A character string spcifying the method for parameter
    estimation and computation of the marginal likelihood. This
    must be eigher "mcmc.bs", for MCMC followed by bridge sampling,
    or "map.lap", for MAP estimation
    followed by Laplace approximation.

p.m
:   A vector of hyperparameters of the model prior.

phi
:   A vector of hyperparameters of the effect prior.

phi0
:   A scalar specifying the hyperparameter on the intercept.

kappa
:   A scalar specifying the hyperparameter of the gamma
    prior on the residual error precision.

nu
:   A scalar specifying the hyperparameter of the gamma prior
    on the residual error precision.

kappa.u
:   A scalar specifying the hyperparameter of the gamma
    prior on the random intercept.

nu.u
:   A scalar specifying the hyperparameter of the gamma
    prior on the random intercept.

tu.lambda
:   A list obtained from
    `get_tu_lambda`. It must contain the transposed
    eigen vector matrix and the eigen values of the covariance
    matrix. The element names must be "tU" and "lambda".

summary
:   A Boolean variable indicating whether a summary or
    MCMC samples should be stored. This is only applicable when
    `method` is set to "mcmc.bs".

seed
:   An integer specifying a seed for RNG.

n.cores
:   An integer specifying the number of cores.

#### Value

A list object containing:

- `fn` - A character string specifying the function.
- `ranef` - A logical.
- `rint` - A logical as to whether the phenotypes have
  been RINT-transformed.
- `p.m` - A vector of hyperparameters of the model prior.
- `seed` - A integer specifying a seed fo RNG.
- `ln.p.y.given.m` - A named vector of the log marginal
  likelihood given each model. If `method` is set to
  "map.lap", the value is shifted by a constant.
- `ln.p.y` - A scalar value of the log marginal
  likelihood. If `method` is set to "map.lap", the value is
  shifted by a constant.
- `p.m.given.y` - A named vector of posterior probability of the
  models.
- `ml.errors` - A named vector of errors in the log marginal
  likelihood. This element is included only when `method` is
  set to "mcmc.bs".
- `stan.list` - A list of `stanfit` objects or data frames
  containing MCMC results for all eight models. This element is inclued only when
  `method` is set to "mcmc.bs". If `summary` is set to
  `TRUE`, a list of data frames containing posterior summary
  is returned. If the option is set to `FALSE`, a list of
  `stanfit` objects is returned.
- `optim.list` - A list of outputs from the `optim`
  function from the `stat` package containing MAP
  estimates and Hessian. This element is included only when
  `method` is set to "map.lap".

#### Contents

Developed by Yuriko Harigaya, Michael Love, William Valdar.

Site built with pkgdown 2.0.9.
