## Supplemental File 1 for "Probabilistic classification of gene-by-treatment interactions on molecular count phenotypes": format_gp.html

Prepare data for a genotype-phenotype plot — format\_gp • classifygxt       

Toggle navigation


classifygxt
0.0.1

- Get started
- Reference
- Changelog

### Prepare data for a genotype-phenotype plot

`format_gp.Rd`

This is a function to prepare data for visualization using
`make_gp_plot`.

```
format_gp(data, fit, seed = 1)
```

#### Arguments

data
:   A list containing phenotype, genotype, treatment, and
    subject, which must be named "y", "g", "t", and "subject ",
    respectively.

fit
:   A list obtained from the `do_bms`.

seed
:   A seed for RNG.

#### Value

A list of data frames.

#### Contents

Developed by Yuriko Harigaya, Michael Love, William Valdar.

Site built with pkgdown 2.0.9.
