## Supplemental File 1 for "Probabilistic classification of gene-by-treatment interactions on molecular count phenotypes": format_pp.html

Prepare data for a barplot of posterior probability — format\_pp • classifygxt       

Toggle navigation


classifygxt
0.0.1

- Get started
- Reference
- Changelog

### Prepare data for a barplot of posterior probability

`format_pp.Rd`

This is a function to prepare data for visualization using
`make_pp_plot`.

```
format_pp(fit, co = FALSE)
```

#### Arguments

fit
:   A list obtained from the `do_bms`.

co
:   A logical varialbe as to whether to visualize the
    probability of crossover interaction. If this is set to
    `TRUE`, `summary` must be set to `FALSE` when
    running `do_bms`.

#### Value

A data frame.

#### Contents

Developed by Yuriko Harigaya, Michael Love, William Valdar.

Site built with pkgdown 2.0.9.
