## Supplemental File 1 for "Probabilistic classification of gene-by-treatment interactions on molecular count phenotypes": get_co.html

Compute posterior probability of crossover interaction — get\_co • classifygxt       

Toggle navigation


classifygxt
0.0.1

- Get started
- Reference
- Changelog

### Compute posterior probability of crossover interaction

`get_co.Rd`

This is a function to compute posterior probability of the
crossover interaction. It takes the output of
`do_bms` as input. `summary` must be set to
`FALSE` when calling `do_bms`. By default, the
joint conditional probabilities are returend. If `prob`
is set to "conditional", the conditional probability given each
model is returned. If the option is set to "marginal", a scalar
value is returned.

```
get_co(fit, prob = "joint")
```

#### Arguments

fit
:   A list obtained from the `do_bms`. `summary`
    must be set to `FALSE` when calling `do_bms`.

prob
:   A character string specifying one of "conditional",
    "joint", and "marginal".

#### Value

A named vector or scalar of posterior probability.

#### Contents

Developed by Yuriko Harigaya, Michael Love, William Valdar.

Site built with pkgdown 2.0.9.
