## Supplemental File 1 for "Probabilistic classification of gene-by-treatment interactions on molecular count phenotypes": get_est.html

Extract parameter estimates — get\_est • classifygxt       

Toggle navigation


classifygxt
0.0.1

- Get started
- Reference
- Changelog

### Extract parameter estimates

`get_est.Rd`

This is a function to extract parameter estimates from
the output from `do_bms`. If `model` is not
specified, estimates for the MAP model is returned.

```
get_est(fit, model = NULL)
```

#### Arguments

fit
:   A list obtained from the `do_bms`.

model
:   A character string or integer specifying the model.

#### Value

A named vector of parameter estimates.

#### Contents

Developed by Yuriko Harigaya, Michael Love, William Valdar.

Site built with pkgdown 2.0.9.
