## Supplemental File 1 for "Probabilistic classification of gene-by-treatment interactions on molecular count phenotypes": get_map.html

Get MAP estimates and Hessian — get\_map • classifygxt       

Toggle navigation


classifygxt
0.0.1

- Get started
- Reference
- Changelog

### Get MAP estimates and Hessian

`get_map.Rd`

Get MAP estimates and Hessian

```
get_map(
  data,
  fn.gp,
  phi,
  phi0 = sqrt(1000),
  kappa = 0.002,
  nu = 0.002,
  kappa.u = 0.002,
  nu.u = 0.002,
  tu.lambda = NULL
)
```

#### Arguments

data
:   A list containing phenotype, genotype, treatment, and
    subject. The elements must be named "y", "g", "t", and
    "subject".

fn.gp
:   A character string specifying the function to model
    the relationship between the genotype and phenotype. This must
    be one of "nonlinear" and "linear", corresponding to nonlinear
    and linear models, respectively.

phi
:   A vector of hyperparameters of the effect prior.

phi0
:   A scalar specifying the hyperparameter on the intercept.

kappa
:   A scalar specifying the hyperparameter of the gamma prior on the residual error precision.

nu
:   A scalar specifying the hyperparameter of the gamma prior
    on the residual error precision.

kappa.u
:   A scalar specifying the hyperparameter of the gamma
    prior on the random intercept.

nu.u
:   A scalar specifying the hyperparameter of the gamma
    prior on the random intercept.

tu.lambda
:   A list containing the transposed eigenvector
    matrix and the eigenvalues of the covariance matrix. The
    element names must be "tU" and "lambda".

#### Value

A list object containing outputs from `optim` in the
`stats` package for the eight models.

#### Contents

Developed by Yuriko Harigaya, Michael Love, William Valdar.

Site built with pkgdown 2.0.9.
