## Supplemental File 1 for "Probabilistic classification of gene-by-treatment interactions on molecular count phenotypes": get_pp.html

Extract posterior probability — get\_pp • classifygxt       

Toggle navigation


classifygxt
0.0.1

- Get started
- Reference
- Changelog

### Extract posterior probability

`get_pp.Rd`

This is a function to extract posterior probability from
the output from `do_bms`. It optionally
aggregates the model categories.

```
get_pp(fit, aggregate = "none")
```

#### Arguments

fit
:   A list obtained from the `do_bms`.

aggregate
:   An optional character string specifying
    whether and how to aggregate the model categories.
    This must be one of "none", "genotype", and "treatment".

#### Value

a named vector of posterior probability

#### Contents

Developed by Yuriko Harigaya, Michael Love, William Valdar.

Site built with pkgdown 2.0.9.
