## Supplemental File 1 for "Probabilistic classification of gene-by-treatment interactions on molecular count phenotypes": get_sign.html

Get posterior probability accounting for the sign of effect sizes — get\_sign • classifygxt       

Toggle navigation


classifygxt
0.0.1

- Get started
- Reference
- Changelog

### Get posterior probability accounting for the sign of effect sizes

`get_sign.Rd`

This is a function to compute posterior probability
of the 27 models accounting for the sign of effect sizes.
It takes the output of `do_bms` as input.

```
get_sign(fit)
```

#### Arguments

fit
:   A list obtained from the `do_bms`.

#### Value

A named vector of posterior probability.

#### Contents

Developed by Yuriko Harigaya, Michael Love, William Valdar.

Site built with pkgdown 2.0.9.
