## Supplemental File 1 for "Probabilistic classification of gene-by-treatment interactions on molecular count phenotypes": get_sign_names.html

Get the names of the 27 models — get\_sign\_names • classifygxt       

Toggle navigation


classifygxt
0.0.1

- Get started
- Reference
- Changelog

### Get the names of the 27 models

`get_sign_names.Rd`

This function returns an ordered vector of character strings
corresponding to the names of the 27 models accounting for the
sign of effect sizes.

```
get_sign_names()
```

#### Value

A vector containing the eight model names.

#### Contents

Developed by Yuriko Harigaya, Michael Love, William Valdar.

Site built with pkgdown 2.0.9.
