## Supplemental File 1 for "Probabilistic classification of gene-by-treatment interactions on molecular count phenotypes": get_tu_lambda.html

Get eigenvectors and eigenvalues of the covariance matrix — get\_tu\_lambda • classifygxt       

Toggle navigation


classifygxt
0.0.1

- Get started
- Reference
- Changelog

### Get eigenvectors and eigenvalues of the covariance matrix

`get_tu_lambda.Rd`

This function computes eigenvectors and eigenvalues of the
covariance matrix, which are needed for running `do_bms` when
modeling a random effect. `kinship` must be set to the default
if a subject-specific random effect will be modeled. `kinship`
must be specified if a polygenic (kinship) random effect will be
modeled.

```
get_tu_lambda(data, kinship = NULL)
```

#### Arguments

data
:   A list containing subjects, genotypes and treatment
    indicators, which must be named "subject" and "t",
    respectively. The list can contain extra elements, such as
    genotypes and phenotypes.

kinship
:   A matrix containing pairwise genetic
    relatedness between subjects. The row and column names must match the
    set of unique elements of "subject" in "data" in the
    corresponding order (i.e., `unique(data$subject)`). This
    is set to `NULL` by default, in which case the identity
    matrix is used.

#### Value

A list object containing:

- `tU` - A matrix containing the transposed eigenvectors
  of the covariance matrix.
- `lambda` - A vector containing the eigenvalues of the
  covariance matrix.

#### Contents

Developed by Yuriko Harigaya, Michael Love, William Valdar.

Site built with pkgdown 2.0.9.
