## Supplemental File 1 for "Probabilistic classification of gene-by-treatment interactions on molecular count phenotypes": index.html

Function reference • classifygxt       

Toggle navigation


classifygxt
0.0.1

- Get started
- Reference
- Changelog

### Reference

| All functions | |
| --- | --- |
|  |  |
| --- | --- |
| `classifygxt-package` `classifygxt` | The 'classifygxt' package. |
| `do_bms()` | Perform BMS for classifying GxT interactions |
| `format_gp()` | Prepare data for a genotype-phenotype plot |
| `format_pp()` | Prepare data for a barplot of posterior probability |
| `get_co()` | Compute posterior probability of crossover interaction |
| `get_est()` | Extract parameter estimates |
| `get_map()` | Get MAP estimates and Hessian |
| `get_model_names()` | Get the names of the eight models |
| `get_pp()` | Extract posterior probability |
| `get_sign()` | Get posterior probability accounting for the sign of effect sizes |
| `get_sign_names()` | Get the names of the 27 models |
| `get_tu_lambda()` | Get eigenvectors and eigenvalues of the covariance matrix |
| `make_data()` | Generate data for simulation analysis |
| `make_gp_plot()` | Make a genotype-phenotype plot with model fit |
| `make_heatmap()` | Make a heatmap of posterior probability |
| `make_pp_plot()` | Make a barplot of posterior probability |

#### Contents

Developed by Yuriko Harigaya, Michael Love, William Valdar.

Site built with pkgdown 2.0.9.
