## Supplemental File 1 for "Probabilistic classification of gene-by-treatment interactions on molecular count phenotypes": make_data.html

Generate data for simulation analysis — make\_data • classifygxt       

Toggle navigation


classifygxt
0.0.1

- Get started
- Reference
- Changelog

### Generate data for simulation analysis

`make_data.Rd`

Generate data for simulation analysis

```
make_data(
  num,
  anno,
  fn,
  lb.maf = 0.05,
  ub.maf = 0.5,
  filter.geno = TRUE,
  sd,
  b0 = 0,
  sigma = 1,
  ranef = FALSE,
  sigma.u = NULL,
  kinship = NULL,
  seed = 1
)
```

#### Arguments

num
:   A named, ordered integer vector specifying the numbers of
    simulations in the eight model categories. The model names in the
    correct order can be obtained using
    `get_model_names`.

anno
:   A data frame containing the subjects (character
    strings or integers) and the treatment conditions (0 or 1) in
    the first and second columns, respectively. The columns must be
    named as "subject" and "condition".

fn
:   A character string specifying the function. This must be
    one of "nonlinear" and "linear", corresponding to nonlinear and
    linear models, respectively.

lb.maf
:   A scalar specifying the lower bound of MAF.

ub.maf
:   A scalar specifying the upper bound of MAF.

filter.geno
:   A Boolean variable as to whether to ensure that
    all genotype levels have at least one observation.

sd
:   A vetor of length three specifying the effect size
    standard deviations.

b0
:   A scalar specifying the intercept.

sigma
:   A scalar specifying the residual error standard
    deviation.

ranef
:   A Boolean variable as to whether to include random effect.

sigma.u
:   A scalar specifying the random intercept standard
    deviation. If `ranef` is `TRUE`, this is set to sqrt(0.2)
    by default.

kinship
:   A matrix containing pairwise genetic relatedness
    between individuals. If `ranef` is `TRUE`, this is set to
    an identity matrix by default.

seed
:   A seed for RNG.

#### Value

A list of lists containing:

- `y` - A vector of phenotypes.
- `g` - A vector of genotypes.
- `t` - A vector of treatment indicators.
- `subject` - A vector of subject.
- `index` - An integer specifying one of the eight
  model. The order of models corresponding to the indices dan be
  obtained using `get_model_names`.
- `maf` - A scalar specifying the minor allele frequency
  used for generating the genotype data.
- `beta` - A named numeric vector specifying the true
  coefficient values used for generating the phenotype data. The
  "b0" element represents the intercept. The "b1", "b2", and "b3"
  elements respectively represent the genotype, treatment, and
  interaction effect sizes.

#### Contents

Developed by Yuriko Harigaya, Michael Love, William Valdar.

Site built with pkgdown 2.0.9.
