## Supplemental File 1 for "Probabilistic classification of gene-by-treatment interactions on molecular count phenotypes": make_gp_plot.html

Make a genotype-phenotype plot with model fit — make\_gp\_plot • classifygxt       

Toggle navigation


classifygxt
0.0.1

- Get started
- Reference
- Changelog

### Make a genotype-phenotype plot with model fit

`make_gp_plot.Rd`

This function generates a genotype-phenotype plot with regression
lines in the control and treated conditions based on the MAP model.

```
make_gp_plot(gp, title = NULL, seed = 1)
```

#### Arguments

gp
:   A list object obtained from the `format_gp`
    function.

title
:   A character string specifying a title.

seed
:   A seed for RNG.

#### Value

A `ggplot2` object

#### Contents

Developed by Yuriko Harigaya, Michael Love, William Valdar.

Site built with pkgdown 2.0.9.
