## Supplemental File 1 for "Probabilistic classification of gene-by-treatment interactions on molecular count phenotypes": make_heatmap.html

Make a heatmap of posterior probability — make\_heatmap • classifygxt       

Toggle navigation


classifygxt
0.0.1

- Get started
- Reference
- Changelog

### Make a heatmap of posterior probability

`make_heatmap.Rd`

This is a function to make a heatmap of posterior probability for a
set of feature-SNP pairs.

```
make_heatmap(input)
```

#### Arguments

input
:   A matrix containing posterior probability. Rows and
    columns must represent feature-SNP pairs and model categories,
    respectively.

#### Value

A `ggplot2` object.

#### Contents

Developed by Yuriko Harigaya, Michael Love, William Valdar.

Site built with pkgdown 2.0.9.
