## Supplemental File 1 for "Probabilistic classification of gene-by-treatment interactions on molecular count phenotypes": make_pp_plot.html

Make a barplot of posterior probability — make\_pp\_plot • classifygxt       

Toggle navigation


classifygxt
0.0.1

- Get started
- Reference
- Changelog

### Make a barplot of posterior probability

`make_pp_plot.Rd`

This function takes output from `format_pp` as input
and generates a barplot of posterior probability.

```
make_pp_plot(pp)
```

#### Arguments

pp
:   A data frame obtained from `format_pp`. For
    visualizing the posterior of crossocer interaction, `co`
    must be set to `TRUE` when running running
    `format_pp`.

#### Value

A `ggplot2` object.

#### Contents

Developed by Yuriko Harigaya, Michael Love, William Valdar.

Site built with pkgdown 2.0.9.
