## Supplemental File 1 for "Probabilistic classification of gene-by-treatment interactions on molecular count phenotypes": lm.stan

```
data {
  int<lower=0> N; // number of observations
  vector[N] y; // vector of observations
  vector<lower=0, upper=2>[N] x1; // vector of genotypes coded as {0, 1, 2}
  vector<lower=0, upper=1>[N] x2; // vector of indicators denoting the treatment
  vector<lower=0, upper=2>[N] x3; // vector of interactions
  int<lower=0, upper=1> m1; // {0,1} selecting coefficient of genotype
  int<lower=0, upper=1> m2; // {0,1} selecting coefficient of treatment
  int<lower=0, upper=1> m3; // {0,1} selecting coefficient of interaction
  real<lower=0> phi1; // scaled prior standard deviation of effects
  real<lower=0> phi2; // scaled prior standard deviation of effects
  real<lower=0> phi3; // scaled prior standard deviation of effects
}
parameters {
  real b0; // intercept
  real b1; // coefficient of genotype
  real b2; // coefficient of treatment
  real b3; // coefficient of interaction
  real<lower=0> sigma_sq_inv; // precision of error
}
transformed parameters {
  real<lower=0> sigma;
  sigma = sigma_sq_inv^(-0.5); // standard deviation of error
}
model {
  target += gamma_lpdf(sigma_sq_inv | 0.001, 0.001); // prior for precision of error
  target += normal_lpdf(b0 | 0, sigma*sqrt(1000)); // prior for intercept
  target += normal_lpdf(b1 | 0, sigma*phi1); // prior for coefficient of treatment; untruncated
  target += normal_lpdf(b2 | 0, sigma*phi2); // prior for coefficient of genotype; untruncated
  target += normal_lpdf(b3 | 0, sigma*phi3); // prior for coefficient of interaction; untruncated
  target += normal_lpdf(y | b0 + m1*x1*b1 + m2*x2*b2 + m3*x3*b3, sigma); // likelihood model
}
```
