## Supplemental File 1 for "Probabilistic classification of gene-by-treatment interactions on molecular count phenotypes": lmm.stan

```
data {
  int<lower=0> N; // individuals
  matrix[N, N] tU;
  // U are eigenvectors of random effect covariance 
  vector[N] lambda; // eigenvalues of random effect covariance
  vector[N] y; // vector of observations
  vector<lower=0, upper=2>[N] x1; // vector of genotypes coded as {0, 1, 2}
  vector<lower=0, upper=1>[N] x2; // vector of indicators denoting the treatment
  vector<lower=0, upper=2>[N] x3; // vector of interactions
  int<lower=0, upper=1> m1; // {0, 1}
  int<lower=0, upper=1> m2; // {0, 1}
  int<lower=0, upper=1> m3; // {0, 1}
  real<lower=0> phi1; // scaled prior standard deviation of effects
  real<lower=0> phi2; // scaled prior standard deviation of effects
  real<lower=0> phi3; // scaled prior standard deviation of effects
}
parameters {
  real b0; // intercept
  real b1; // coefficient of genotype
  real b2; // coefficient of treatment
  real b3; // coefficient of interaction
  real<lower=0> sigma2;
  real<lower=0> sigma2_u;
}
model {
  vector[N] o; // diagonal elements
  vector[N] R;
  vector[N] r_mu;
  vector[N] mu;
  real sigma;
  sigma = sqrt(sigma2);
  mu = b0 + m1 * x1 * b1 + m2 * x2 * b2 + m3 * x3 * b3;
  o = sigma2_u * lambda + sigma2; 
  R = tU * (y - mu);
  target += gamma_lpdf(sigma2^(-1) | 0.001, 0.001);
  // prior for precision of error
  target += gamma_lpdf(sigma2_u^(-1) | 0.001, 0.001);
  // prior for precision of random effect
  target += normal_lpdf(b0 | 0, sigma * sqrt(1000));
  // prior for intercept
  target += normal_lpdf(b1 | 0, sigma * phi1);
  // prior for coefficient of genotype; untruncated
  target += normal_lpdf(b2 | 0, sigma * phi2);
  // prior for coefficient of treatment; untruncated
  target += normal_lpdf(b3 | 0, sigma * phi3);
  // prior for coefficient of interaction; untruncated
  target += - 0.5 * sum(log(o));
  target += - 0.5 * sum(R .* R ./ o );
  target += - 0.5 * N * (2 * pi());
}
```
