## Supplemental File 1 for "Probabilistic classification of gene-by-treatment interactions on molecular count phenotypes": classifygxt-package.R

```
#' The 'classifygxt' package.
#'
#' @description A DESCRIPTION OF THE PACKAGE
#'
#' @docType package
#' @name classifygxt-package
#' @aliases classifygxt
#' @useDynLib classifygxt, .registration = TRUE
#' @import methods
#' @import Rcpp
#' @importFrom rstan sampling
#' @importFrom rstantools rstan_config
#' @importFrom RcppParallel RcppParallelLibs
#'
#' @references
#' Stan Development Team (2022). RStan: the R interface to Stan. R package version 2.21.5. https://mc-stan.org
#'
NULL
```
