## Supplemental File 1 for "Probabilistic classification of gene-by-treatment interactions on molecular count phenotypes": core.R

```
#' Generate data for simulation analysis
#'
#' @export
#' @import mvtnorm
#' @importFrom magrittr "%>%"
#' @importFrom stats runif rbinom rnorm
#' @param num A named, ordered integer vector specifying the numbers of
#' simulations in the eight model categories. The model names in the
#'     correct order can be obtained using
#'     \code{\link{get_model_names}}.
#' @param fn A character string specifying the function. This must be
#'     one of "nonlinear" and "linear", corresponding to nonlinear and
#'     linear models, respectively.
#' @param lb.maf A scalar specifying the lower bound of MAF.
#' @param ub.maf A scalar specifying the upper bound of MAF.
#' @param filter.geno A Boolean variable as to whether to ensure that
#'     all genotype levels have at least one observation.
#' @param anno A data frame containing the subjects (character
#'     strings or integers) and the treatment conditions (0 or 1) in
#'     the first and second columns, respectively. The columns must be
#'     named as "subject" and "condition".
#' @param sd A vetor of length three specifying the effect size
#'     standard deviations.
#' @param b0 A scalar specifying the intercept.
#' @param sigma A scalar specifying the residual error standard
#'     deviation.
#' @param ranef A Boolean variable as to whether to include random effect.
#' @param sigma.u A scalar specifying the random intercept standard
#'     deviation. If \code{ranef} is \code{TRUE}, this is set to sqrt(0.2)
#'     by default.
#' @param kinship A matrix containing pairwise genetic relatedness
#'     between individuals. If \code{ranef} is \code{TRUE}, this is set to
#'     an identity matrix by default.
#' @param seed A seed for RNG.
#'
#' @return A list of lists containing:
#' \itemize{
#' \item{\code{y} - A vector of phenotypes.}
#' \item{\code{g} - A vector of genotypes.}
#' \item{\code{t} - A vector of treatment indicators.}
#' \item{\code{subject} - A vector of subject.}
#' \item{\code{index} - An integer specifying one of the eight
#'     model. The order of models corresponding to the indices dan be
#'     obtained using \code{\link{get_model_names}}.}
#' \item{\code{maf} - A scalar specifying the minor allele frequency
#'     used for generating the genotype data.}
#' \item{\code{beta} - A named numeric vector specifying the true
#'     coefficient values used for generating the phenotype data. The
#'     "b0" element represents the intercept. The "b1", "b2", and "b3"
#'     elements respectively represent the genotype, treatment, and
#'     interaction effect sizes. }
#' }
#'
make_data <- function(num, anno,
                      fn,
                      lb.maf=0.05, ub.maf=0.5,
                      filter.geno=TRUE,
                      sd, b0=0, sigma=1,
                      ranef=FALSE,
                      sigma.u=NULL,
                      kinship=NULL,
                      seed=1) {

    set.seed(seed)

    # if (is.null(num)) {
    #     num <- rep(1000, 8)
    # }

    if (!ranef) {
        if (!is.null(sigma.u)) {
            stop("`ranef` must be set to `TRUE` when `sigma.u` is supplied")
        }
        if (!is.null(kinship)) {
            stop("`ranef` must be set to `TRUE` when `kinship` is supplied")
        }
    }

    # extract info from the anno object
    subject <- anno$subject
    t <- anno$condition
    n <- nrow(anno) # the total number of samples
    n.sub <- length(unique(subject))

    sd.g <- sd[1]; sd.t <- sd[2]; sd.gxt <- sd[3]

    # get the covariance matrix
    if (ranef) {
        if (is.null(kinship)) {
            A <- diag(n.sub)
            rownames(A) <- colnames(A) <- seq_len(n.sub)
        } else {
            A <- kinship
        }

        if (is.null(sigma.u)) {
            sigma.u <- sqrt(0.2)
        }

        # generate the incidence matrix
        Z <- matrix(0, nrow=n, ncol=n.sub)
        for (i in seq_len(n)) {
            for (j in seq_len(n.sub)) {
                if (subject[i] == j) Z[i, j] <- 1
            }
        }

        Sigma.u <- Z %*% A %*% t(Z)
        cov <-
            sigma.u^2 * Sigma.u +
            sigma^2 * diag(n)
    }

    # get a matrix containing m vectors in rows
    model.name <- get_model_mat()

    # create a object to store data and info
    data.list.list <- vector("list", length(num))

    for (j in seq_along(num)) {

        n.data <- num[j]
        if (n.data == 0) {
            data.list.list[[j]] <- NULL
            next
        }

        index <- j
        data.list <- vector("list", length(n.data))

        for (i in seq_len(n.data)) {

            # get the m vector
            m.vec <- as.numeric(model.name[index, ])

            # draw maf
            maf <- runif(1, lb.maf, ub.maf)

            # simulate genotype
            geno.vec <- 0
            if (!filter.geno) {
                geno <- rbinom(n=n.sub, size=2, prob=maf)
            } else {
                while (any(geno.vec == 0)) {
                    geno <- rbinom(n=n.sub, size=2, prob=maf)
                    geno.vec <- factor(
                        geno, levels=c(0, 1, 2)) %>%
                        table %>%
                        as.numeric
                }
            }

            # get predictors
            g <- rep(geno, each=2) # genotype

            b1 <- ifelse(
                m.vec[1] == 1, rnorm(n=1, mean=0, sd=sd.g), 0)
            b2 <- ifelse(
                m.vec[2] == 1, rnorm(n=1, mean=0, sd=sd.t), 0)
            b3 <- ifelse(
                m.vec[3] == 1, rnorm(n=1, mean=0, sd=sd.gxt), 0)
            beta <- c(b0, b1, b2, b3) %>%
                `names<-`(c("b0", "b1", "b2", "b3"))

            y.mean <- get_mean(
                g=g, t=t, param=beta, m=m.vec, fn=fn)

            if (!ranef) {
                y <- y.mean +
                    rnorm(n=2 * n.sub, mean=0, sd=sigma)
            } else if (ranef){
                u.mean <- rep(0, n)
                y <- y.mean +
                    rmvnorm(n=1, mean=u.mean, sigma=cov) %>%
                    as.numeric
            }

            data.list[[i]] <- list(
                y=y, g=g, t=t, subject=subject,
                index=index, maf=maf, beta=beta)

        }

        data.list.list[[j]] <- data.list
    }

    data.list <- do.call("c", data.list.list)
    data.list

}

#' Get eigenvectors and eigenvalues of the covariance matrix
#'
#' This function computes eigenvectors and eigenvalues of the
#' covariance matrix, which are needed for running \code{\link{do_bms}} when
#' modeling a random effect. \code{kinship} must be set to the default
#' if a subject-specific random effect will be modeled. \code{kinship}
#' must be specified if a polygenic (kinship) random effect will be
#' modeled.
#'
#' @export
#' @import bridgesampling matrixStats
#' @importFrom rstan summary
#' @importFrom magrittr "%>%"
#' @param data A list containing subjects, genotypes and treatment
#'     indicators, which must be named "subject" and "t",
#'     respectively. The list can contain extra elements, such as
#'     genotypes and phenotypes.
#' @param kinship A matrix containing pairwise genetic
#'     relatedness between subjects. The row and column names must match the
#'     set of unique elements of "subject" in "data" in the
#'     corresponding order (i.e., \code{unique(data$subject)}). This
#'     is set to \code{NULL} by default, in which case the identity
#'     matrix is used.
#'
#' @return A list object containing:
#' \itemize{
#' \item{\code{tU} - A matrix containing the transposed eigenvectors
#'     of the covariance matrix.}
#' \item{\code{lambda} - A vector containing the eigenvalues of the
#'     covariance matrix.}
#' }
#'
get_tu_lambda <- function(data,
                          kinship=NULL) {

    # check input types
    # g <- data$g
    t <- data$t
    #  <- data$y
    n <- length(t)
    if (!is.numeric(t)) {
        stop("`data` has an incorrect format")
    }

    subject <- data$subject
    if (!(is.character(subject) | is.integer(subject))) {
        stop("`data` has an incorrect format")
    }

    sub.name <- unique(subject)
    n.sub <- length(sub.name)
    sub.vec <- subject %>%
        factor(levels=sub.name) %>%
        as.numeric

    if (is.null(kinship)) {
        A <- diag(n.sub)
    } else if (is.matrix(kinship)) {
        A <- kinship
        if (is.null(colnames(A)) | is.null(rownames(A))) {
            stop(paste(
                "the kinship matrix must have",
                "the row and column names"))
        }
        if (!all(colnames(A) == rownames(A))) {
            stop(paste(
                "the row and column names of",
                "the kiship matrix must be identical"))
        }
        if (!all(rownames(A) == sub.name)) {
            stop(paste(
                 "the row names of",
                 "the kiship matrix must match",
                 "the subject"))
        }
    }

    # generate the incidence matrix
    Z <- matrix(0, nrow=n, ncol=n.sub)
    for (i in seq_len(n)) {
        for (j in seq_len(n.sub)) {
            if (sub.vec[i] == j)
                Z[i, j] <- 1
        }
    }

    # get the covariance matrix
    Sigma.u <- Z %*% A %*% t(Z)

    # do eigen decomposition
    eigen.sample.kernel <- eigen(Sigma.u, symmetric=TRUE)
    tU <- t(eigen.sample.kernel$vectors)
    lambda <- eigen.sample.kernel$values

    output <- list(tU=tU, lambda=lambda)
    output
}

#' Perform BMS for classifying GxT interactions
#'
#' This function takes as input individual phenotype and genotype data
#' and performs BMS for a feature-SNP pair (or a SNP). The output
#' includes posterior probabilities of different types of GxT
#' interactions as well as the log marginal likelihood, which can be
#' used for optimizing the hyperparameter values by an empirical Bayes
#' approach. BMS is performed by either MCMC followed by bridge
#' sampling or MAP estimation followed by Laplace approximation. In
#' the latter case, a relative marginal likelihood is returned. That
#' is, the log marginal likelihood value is shifted by a
#' constant. Note that this shift does not affect the posterior
#' probabilities or hyperparameter optimization.
#' \code{tu.lambda} must be supplied when modeling a random effect.
#'
#' @export
#' @import bridgesampling matrixStats
#' @importFrom rstan summary
#' @importFrom stats qnorm
#' @param data A list containing vectors of subjects, genotypes,
#'     phenotypes, and treatment indicators, which must be named
#'     "subject", "g", "y", and "t", respectively.
#' @param fn A character string specifying the function. This must be
#'     one of "nonlinear" and "linear", corresponding to nonlinear and
#'     linear models, respectively.
#' @param rint A logical as to whether the phenotypes need to be
#'     RINT-transformed. This cannot be set to \code{TRUE} if \code{fn} is
#'     set to "nonlinear".
#' @param method A character string spcifying the method for parameter
#'     estimation and computation of the marginal likelihood. This
#'     must be eigher "mcmc.bs", for MCMC followed by bridge sampling,
#'     or "map.lap", for MAP estimation
#'     followed by Laplace approximation.
#' @param p.m A vector of hyperparameters of the model prior.
#' @param phi A vector of hyperparameters of the effect prior.
#' @param phi0 A scalar specifying the hyperparameter on the intercept.
#' @param kappa A scalar specifying the hyperparameter of the gamma
#'     prior on the residual error precision.
#' @param nu A scalar specifying the hyperparameter of the gamma prior
#'     on the residual error precision.
#' @param kappa.u A scalar specifying the hyperparameter of the gamma
#'     prior on the random intercept.
#' @param nu.u A scalar specifying the hyperparameter of the gamma
#'     prior on the random intercept.
#' @param tu.lambda A list obtained from
#'     \code{\link{get_tu_lambda}}. It must contain the transposed
#'     eigen vector matrix and the eigen values of the covariance
#'     matrix. The element names must be "tU" and "lambda".
#' @param summary A Boolean variable indicating whether a summary or
#'     MCMC samples should be stored. This is only applicable when
#'     \code{method} is set to "mcmc.bs".
#' @param seed An integer specifying a seed for RNG.
#' @param n.cores An integer specifying the number of cores.
#'
#' @return A list object containing:
#' \itemize{
#' \item{\code{fn} -  A character string specifying the function.}
#' \item{\code{ranef} - A logical.}
#' \item{\code{rint} - A logical as to whether the phenotypes have
#'     been RINT-transformed.}
#' \item{\code{p.m} - A vector of hyperparameters of the model prior.}
#' \item{\code{seed} - A integer specifying a seed fo RNG.}
#' \item{\code{ln.p.y.given.m} - A named vector of the log marginal
#'     likelihood given each model. If \code{method} is set to
#'     "map.lap", the value is shifted by a constant.}
#' \item{\code{ln.p.y} - A scalar value of the log marginal
#'     likelihood. If \code{method} is set to "map.lap", the value is
#'     shifted by a constant.}
#' \item{\code{p.m.given.y} - A named vector of posterior probability of the
#'     models.}
#' \item{\code{ml.errors} - A named vector of errors in the log marginal
#'     likelihood. This element is included only when \code{method} is
#'     set to "mcmc.bs".}
#' \item{\code{stan.list} - A list of \code{stanfit} objects or data frames
#'     containing MCMC results for all eight models. This element is inclued only when
#'     \code{method} is set to "mcmc.bs". If \code{summary} is set to
#'     \code{TRUE}, a list of data frames containing posterior summary
#'     is returned. If the option is set to \code{FALSE}, a list of
#'     \code{stanfit} objects is returned.}
#' \item{\code{optim.list} - A list of outputs from the \code{optim}
#'     function from the \code{stat} package containing MAP
#'     estimates and Hessian. This element is included only when
#'     \code{method} is set to "map.lap".}
#' }
#'
do_bms <- function(data,
                   fn,
                   rint=FALSE,
                   method,
                   p.m=NULL,
                   phi, phi0=sqrt(1e3),
                   kappa=0.002, nu=0.002,
                   kappa.u=0.002, nu.u=0.002,
                   tu.lambda=NULL,
                   summary=TRUE,
                   seed=1, n.cores=4) {

    # check input formats
    g <- data$g
    t <- data$t
    y <- data$y
    n <- length(y)
    if (!(is.numeric(g) & is.numeric(t) & is.numeric(y))) {
        stop("`data` has an incorrect format")
    }

    if (rint) {
        if (fn == "nonlinear") {
            stop("`rint` cannot be `TRUE` if `fn` is 'nonlinear'")
        }
        # perform RINT-transformation
        y <- qnorm((rank(y) - (1/2)) / length(y))
        data$y <- y
    }

    if (!(method %in% c("mcmc.bs", "map.lap"))) {
        stop("`method` must be eigher 'mcmc.bs' or 'map.lap'")
    }

    if (!(is.numeric(phi) & length(phi) == 3)) {
        stop("`phi` has an incorrect format")
    } else {
        phi1 <- phi[1]; phi2 <- phi[2]; phi3 <- phi[3]
    }

    if (is.null(tu.lambda)) {
        ranef <- FALSE
    } else if (class(tu.lambda) == "list") {
        ranef <- TRUE
    } else {
        stop("`tu.lambda` has an incorrect format")
    }

    # set a uniform prior over the models
    model.mat <- get_model_mat()
    model.name <- get_model_names()
    if (is.null(p.m)) {
        p.m <- rep(1/8, 8)
    }
    names(p.m) <- model.name

    if (method == "mcmc.bs") {

        # set the number of cores
        options(core=n.cores)

        # specify the data for stan
        data.stan <- list(
            N=n, y=y,
            phi1=phi1, phi2=phi2, phi3=phi3,
            phi0=phi0, kappa=kappa, nu=nu,
            kappa.u=kappa.u, nu.u=nu.u)

        if (fn == "nonlinear") {
            data.stan$x1 <- (1 - g/2) * (1 - t)
            data.stan$x2 <- (g/2) * (1 - t)
            data.stan$x3 <- (1 - g/2) * t
            data.stan$x4 <- (g/2) * t
        } else if (fn == "linear") {
            data.stan$x1 <- g
            data.stan$x2 <- t
            data.stan$x3 <- g * t
        }

        if (ranef) {
            data.stan$tU <- tu.lambda$tU
            data.stan$lambda <- tu.lambda$lambda
        }

        # specify objects to store the results
        ln.p.y.given.m <- rep(NA, length(model.name))
        names(ln.p.y.given.m) <- model.name
        ml.errors <-  rep(NA, length(model.name))
        names(ml.errors) <- model.name
        stan.list <- vector("list", length(model.name))
        names(stan.list) <- model.name

        # iterate through all of the models
        for (i in seq_along(p.m)) {
            # cat(i, "\t")
            data.stan$m1 <- model.mat[i, 1]
            data.stan$m2 <- model.mat[i, 2]
            data.stan$m3 <- model.mat[i, 3]
            # call the model using stan
            # with increased adapt_delta
            # to avoid warnings about divergence
            # set refresh=0 to suppress output
            if (fn == "nonlinear" & ranef) {
                fit <- sampling(
                    object=stanmodels$nlmm, data=data.stan, seed=seed,
                    control=list(adapt_delta=0.99), refresh=0)
            } else if (fn == "nonlinear" & !ranef) {
                fit <- sampling(
                    object=stanmodels$nl, data=data.stan, seed=seed,
                    control=list(adapt_delta=0.99), refresh=0)
            } else if (fn == "linear" & ranef) {
                fit <- sampling(
                    object=stanmodels$lmm, data=data.stan, seed=seed,
                    control=list(adapt_delta=0.99), refresh=0)
            } else if (fn == "linear" & !ranef) {
                fit <- sampling(
                    object=stanmodels$lm, data=data.stan, seed=seed,
                    control=list(adapt_delta=0.99), refresh=0)
            }
            # bridge sampling to estimate marginal likelihood;
            # store estimate and error
            fit.bridge <- bridge_sampler(fit, silent=TRUE)
            ln.p.y.given.m[i] <- fit.bridge$logml
            ml.errors[i] <- as.numeric(
                strsplit(error_measures(fit.bridge)$percentage, "%"))
            if (summary) {
                stan.list[[i]] <- summary(fit)$summary
            } else {
                stan.list[[i]] <- fit
            }
            rm(fit); rm(fit.bridge)
        }

        ln.p.y <- logSumExp(ln.p.y.given.m + log(p.m))
        p.m.given.y <- exp(ln.p.y.given.m + log(p.m) - ln.p.y)

        output <- list(
            fn=fn,
            ranef=ranef,
            rint=rint,
            p.m=p.m,
            seed=seed,
            ln.p.y.given.m=ln.p.y.given.m,
            ln.p.y=ln.p.y,
            p.m.given.y=p.m.given.y,
            ml.errors=ml.errors,
            stan.list=stan.list)

    } else if (method == "map.lap") {
        if (!ranef) {
            optim.list <- get_map(
                data=data, fn.gp=fn,
                phi=phi, phi0=phi0,
                kappa=kappa, nu=nu)
        } else if (ranef) {
            optim.list <- get_map(
                data=data, fn.gp=fn,
                phi=phi, phi0=phi0,
                kappa=kappa, nu=nu,
                kappa.u=kappa.u, nu.u=nu.u,
                tu.lambda=tu.lambda)
        }

        ln.p.y.given.m <- sapply(optim.list, get_marginal, n=n) %>%
            `names<-`(model.name)
        ln.p.y <- logSumExp(ln.p.y.given.m + log(p.m), na.rm=TRUE)
        p.m.given.y <- exp(ln.p.y.given.m + log(p.m) - ln.p.y) %>%
            `names<-`(model.name)

        output <- list(
            fn=fn,
            ranef=ranef,
            rint=rint,
            p.m=p.m,
            seed=seed,
            ln.p.y.given.m=ln.p.y.given.m,
            ln.p.y=ln.p.y,
            p.m.given.y=p.m.given.y,
            optim.list=optim.list)
    }

    output

}

#' Get MAP estimates and Hessian
#'
#' @export
#' @importFrom magrittr "%>%"
#' @importFrom stats lm coef optim dnorm dgamma
#' @param data A list containing phenotype, genotype, treatment, and
#'     subject. The elements must be named "y", "g", "t", and
#'     "subject".
#' @param fn.gp A character string specifying the function to model
#'     the relationship between the genotype and phenotype. This must
#'     be one of "nonlinear" and "linear", corresponding to nonlinear
#'     and linear models, respectively.
#' @param phi A vector of hyperparameters of the effect prior.
#' @param phi0 A scalar specifying the hyperparameter on the intercept.
#' @param kappa A scalar specifying the hyperparameter of the gamma prior on the residual error precision.
#' @param nu A scalar specifying the hyperparameter of the gamma prior
#'     on the residual error precision.
#' @param kappa.u A scalar specifying the hyperparameter of the gamma
#'     prior on the random intercept.
#' @param nu.u A scalar specifying the hyperparameter of the gamma
#'     prior on the random intercept.
#' @param tu.lambda A list containing the transposed eigenvector
#'     matrix and the eigenvalues of the covariance matrix. The
#'     element names must be "tU" and "lambda".
#'
#' @return A list object containing outputs from \code{optim} in the
#' \code{stats} package for the eight models.
get_map <- function(data,
                    fn.gp,
                    phi, phi0=sqrt(1e3),
                    kappa=0.002, nu=0.002,
                    kappa.u=0.002, nu.u=0.002,
                    tu.lambda=NULL) {

    if (is.null(tu.lambda)) {
        ranef <- FALSE
    } else if (class(tu.lambda) == "list") {
        ranef <- TRUE
    } else {
        stop("`tu.lambda` has an incorrect format")
    }

    model.name <- get_model_names()
    res.list <- vector("list", length(model.name))
    names(res.list) <- model.name

    if (!ranef) {
        init.list <- get_init(data=data, ranef=ranef)
    } else if (ranef) {
        # get 3 sets of initial values per model
        init.list.list <- get_init(data=data, ranef=ranef)
    }

    # l.fn <- get_models()
    # l.obj <- get_objectives()
    # l.bounds <- get_bounds()
    m.vec.list <- get_m_vectors()

    phi1 <- phi[1]; phi2 <- phi[2]; phi3 <- phi[3]
    hyper <- list(
        phi0=phi0,
        phi1=phi1,
        phi2=phi2,
        phi3=phi3,
        kappa=kappa, nu=nu,
        kappa.u=kappa.u, nu.u=nu.u)

    # consider using nest()
    for (i in seq_along(model.name)) {
        m <- m.vec.list[[i]]
        data$m1 <- m[1]
        data$m2 <- m[2]
        data$m3 <- m[3]

        warn0 <- getOption("warn")
        options(warn=-1)
        if (!ranef) {
            res.list[[i]] <- try(
                optim(
                    par=init.list[[i]],
                    fn=get_obj,
                    data=data, # further arguments for fn
                    hyper=hyper, # further arguments for fn
                    fn.gp=fn.gp, # further arguments for fn
                    ranef=ranef, # further arguments for fn
                    tu.lambda=tu.lambda, # further arguments for fn
                    method="BFGS",
                    hessian=TRUE))
        } else if (ranef) {
            init.list <- init.list.list[[i]]
            for (j in seq_along(init.list)) {
                tmp <- try(
                    optim(
                        par=init.list[[j]],
                        fn=get_obj,
                        data=data,
                        hyper=hyper,
                        fn.gp=fn.gp,
                        ranef=ranef,
                        tu.lambda=tu.lambda,
                        method="BFGS",
                        hessian=TRUE))
                if (class(tmp) != "try-error") break
            }
            res.list[[i]] <- tmp
        }
        on.exit(options(warn=warn0))
    }
    res.list
}

#' Extract parameter estimates
#'
#' This is a function to extract parameter estimates from
#' the output from \code{\link{do_bms}}. If \code{model} is not
#' specified, estimates for the MAP model is returned.
#'
#' @export
#' @importFrom magrittr "%>%"
#' @param fit A list obtained from the \code{\link{do_bms}}.
#' @param model A character string or integer specifying the model.
#'
#' @return A named vector of parameter estimates.
get_est <- function(fit,
                    model=NULL) {

    model.name <- get_model_names()
    model.mat <- get_model_mat()

    if (is.null(model)) {
        # message("returning estimates for the posterior mode")
        model.index <- which.max(fit$p.m.given.y)
    } else if (model %in% model.name) {
        model.index <- which(model.name == model)
    } else if (model %in% seq_len(8)) {
        model.index <- model
    } else {
        stop("`model` is in an incorrect format")
    }

    ranef <- fit$ranef

    if (is.list(fit$stan.list) & fit$ranef == FALSE) {

        fit <- fit$stan.list[[model.index]]
        if (class(fit)[1] == "stanfit") {
            fit <- rstan::summary(fit)$summary
        }

        beta.name <- c("b0", "b1", "b2", "b3")
        sd.name <- c("sigma")

        beta <- fit[beta.name, "mean"]
        m <- model.mat[model.index, ] %>% as.numeric
        # replace values with zero's when they are not non-zero
        beta <- c(1, m) * beta
        sd <- fit[sd.name, "mean"]

        output <- c(beta, sd)
        names(output) <- c(beta.name, sd.name)

    } else if (is.list(fit$stan.list) & fit$ranef == TRUE) {

        fit <- fit$stan.list[[model.index]]
        if (class(fit)[1] == "stanfit") {
            fit <- rstan::summary(fit)$summary
        }

        beta.name <- c("b0", "b1", "b2", "b3")
        var.name <- c("sigma2", "sigma2_u")
        sd.name <- c("sigma", "sigma_u")

        beta <- fit[beta.name, "mean"]
        m <- model.mat[model.index, ] %>% as.numeric
        # replace values with zero's when they are not non-zero
        beta <- c(1, m) * beta
        sd <- sqrt(fit[var.name, "mean"])

        output <- c(beta, sd)
        names(output) <- c(beta.name, sd.name)

    } else if (is.list(fit$optim.list) & fit$ranef == FALSE) {

        fit <- fit$optim.list[[model.index]]
        # beta.name <- c("b0", "bg", "bt", "bgt")
        beta.name <- c("b0", "b1", "b2", "b3")
        sd.name <- c("sigma")

        beta <- fit$par[names(fit$par) %in% beta.name]
        # print(beta)
        for (i in beta.name) {
            if (is.na(beta[i])) {
                beta[i] <- 0
            }
        }
        beta <- beta[beta.name]
        sd <- fit$par[names(fit$par) %in% sd.name]

        output <- c(beta, sd)
        names(output) <- c(beta.name, sd.name)

    } else if (is.list(fit$optim.list) & fit$ranef == TRUE) {

        fit <- fit$optim.list[[model.index]]
        # beta.name <- c("b0", "bg", "bt", "bgt")
        beta.name <- c("b0", "b1", "b2", "b3")
        sd.name <- c("sigma", "sigma_u")

        beta <- fit$par[names(fit$par) %in% beta.name]
        for (i in beta.name) {
            if (is.na(beta[i])) {
                beta[i] <- 0
            }
        }
        beta <- beta[beta.name]
        sd <- fit$par[names(fit$par) %in% sd.name]

        output <- c(beta, sd)
        names(output) <- c(beta.name, sd.name)

    }

    output

}

#' Extract posterior probability
#'
#' This is a function to extract posterior probability from
#' the output from \code{\link{do_bms}}. It optionally
#' aggregates the model categories.
#'
#' @export
#' @importFrom magrittr "%>%"
#' @param fit A list obtained from the \code{\link{do_bms}}.
#' @param aggregate An optional character string specifying
#' whether and how to aggregate the model categories.
#' This must be one of "none", "genotype", and "treatment".
#'
#' @return a named vector of posterior probability
get_pp <- function(fit,
                   aggregate="none") {

    if (!(aggregate %in% c("none", "genotype", "treatment"))) {
        stop("`aggregate` must be one of 'none', 'genotype', and 'treatment'")
    }

    pp <- fit$p.m.given.y

    if (aggregate == "none") {
        names(pp) <- get_model_names()
    } else if (aggregate != "none") {
        if (aggregate == "genotype") {
            cat.list <- list(
                nogxt=1:4,
                induced=c(5, 7),
                altered=c(6, 8))
        } else if (aggregate == "treatment") {
            cat.list <- list(
                nogxt=1:4,
                restricted=5:6,
                varying=7:8)
        }
        pp <- sapply(cat.list, function(x) sum(pp[x]))
    }
    pp
}

#' Compute posterior probability of crossover interaction
#'
#' This is a function to compute posterior probability of the
#' crossover interaction. It takes the output of
#' \code{\link{do_bms}} as input. \code{summary} must be set to
#' \code{FALSE} when calling \code{\link{do_bms}}. By default, the
#' joint conditional probabilities are returend. If \code{prob}
#' is set to "conditional", the conditional probability given each
#' model is returned. If the option is set to "marginal", a scalar
#' value is returned.
#'
#' @export
#' @importFrom magrittr "%>%"
#' @param fit A list obtained from the \code{\link{do_bms}}. `summary`
#'     must be set to \code{FALSE} when calling \code{\link{do_bms}}.
#' @param prob A character string specifying one of "conditional",
#'     "joint", and "marginal".
#'
#' @return A named vector or scalar of posterior probability.
get_co <- function(fit,
                   prob="joint") {

    stan.list <- fit$stan.list

    if (class(stan.list[[1]]) != "stanfit") {
        stop("`summary` must be set to `FALSE` when calling `do_bms()` to use this function")
    }

    if (!(prob %in% c("joint", "conditional", "marginal"))) {
        stop("`prob` must be one of 'joint', 'conditional', and 'marginal'")
    }

    model.name <- get_model_names()
    model.mat <- get_model_mat()

    fn <- fit$fn
    pp <- fit$p.m.given.y
    p.co.given.m <- rep(NA, 2)

    for (j in 1:2) {
        n.co <- 0 # number of samples with the coerged pattern
        m <- model.mat[(j + 6), ]
        post <- rstan::extract(stan.list[[(j + 6)]], permute=FALSE)
        n.iter <- dim(post)[1]
        n.chain <- dim(post)[2]
        # for the k-th iteration
        for (k in seq_len(n.iter)) {
            # for l-th chain
            for (l in seq_len(n.chain)) {
                beta <- post[k, l, 1:4]
                # names(b) <- c("b0", "bg", "bt", "bgt")
                names(beta) <- c("b0", "b1", "b2", "b3")
                e00 <- get_mean(g=0, t=0, param=beta, m=m, fn=fn)
                e01 <- get_mean(g=0, t=1, param=beta, m=m, fn=fn)
                e20 <- get_mean(g=2, t=0, param=beta, m=m, fn=fn)
                e21 <- get_mean(g=2, t=1, param=beta, m=m, fn=fn)
                # print((e01 - e00) * (e21 - e20))
                if ((e01 - e00) * (e21 - e20) < 0) {
                    n.co <- n.co + 1
                }
            }
        }
        p.co.given.m[j] <- n.co / (n.iter * n.chain)
    }

    p.co.given.m <- c(rep(0, 6), p.co.given.m)
    p.co <- p.co.given.m * pp

    if (prob == "joint") {
        output <- p.co
        names(output) <- model.name
    } else if (prob == "conditional") {
        output <- p.co.given.m
        names(output) <- model.name
    } else if (prob == "marginal") {
        output <- sum(p.co)
        unname(output)
    }

    output

}

#' Get posterior probability accounting for the sign of effect sizes
#'
#' This is a function to compute posterior probability
#' of the 27 models accounting for the sign of effect sizes.
#' It takes the output of \code{\link{do_bms}} as input.
#'
#' @export
#' @importFrom magrittr "%>%"
#' @param fit A list obtained from the \code{\link{do_bms}}.
#'
#' @return A named vector of posterior probability.
#'
get_sign <- function(fit) {

    # reorder the models
    sign.mat <- get_sign_mat()
    abs.mat <- sign.mat %>% abs
    min.mat <- get_model_mat()

    sign.name <- sign.mat %>% get_names_from_matrix
    abs.name <- abs.mat %>% get_names_from_matrix
    min.name <- min.mat %>% get_names_from_matrix

    abs.factor <- factor(abs.name, levels=min.name)
    model.order <- abs.factor %>% order
    # model.cumsum <- abs.factor %>% table %>% cumsum

    p.m.given.y <- fit$p.m.given.y

    output <- rep(0, length(sign.name))
    names(output) <- sign.name

    if (is.list(fit$stan.list)) {
        for (j in seq_along(fit$stan.list)) {
            tmp <- fit$stan.list[[j]]
            if (class(tmp)[1] == "stanfit") {
                tmp <- rstan::summary(tmp)$summary
            }
            par <- tmp[, "mean"]
            model <- p.m.given.y[j] %>%
                names %>%
                strsplit(",") %>%
                unlist
            if ("b1" %in% names(par) && par["b1"] < 0) {
                model[1] <- "-1"
            }
            if ("b2" %in% names(par) && par["b2"] < 0) {
                model[2] <- "-1"
            }
            if ("b3" %in% names(par) && par["b3"] < 0) {
                model[3] <- "-1"
            }
            model <- paste(model, collapse=",")
            index.new <- (sign.name == model) %>% which
            output[index.new] <- p.m.given.y[j]
        }
    } else if (is.list(fit$optim.list)) {
        for (j in seq_along(fit$optim.list)) {
            par <- fit$optim.list[[j]]$par
            model <- p.m.given.y[j] %>%
                names %>%
                strsplit(",") %>%
                unlist
            if ("b1" %in% names(par) && par["b1"] < 0) {
                model[1] <- "-1"
            }
            if ("b2" %in% names(par) && par["b2"] < 0) {
                model[2] <- "-1"
            }
            if ("b3" %in% names(par) && par["b3"] < 0) {
                model[3] <- "-1"
            }
            model <- paste(model, collapse=",")
            index.new <- (sign.name == model) %>% which
            output[index.new] <- p.m.given.y[j]
        }
    }

    output[model.order]

}
```
