## Supplemental File 1 for "Probabilistic classification of gene-by-treatment interactions on molecular count phenotypes": helper.R

```
get_obj <- function(param, data, hyper,
                    fn.gp, ranef, tu.lambda) {

    sigma <- param["sigma"]
    y <- data$y
    g <- data$g
    t <- data$t
    n <- length(y)
    # note that m1, m2, and m3 are needed later
    m1 <- data$m1; m2 <- data$m2; m3 <- data$m3
    m <- c(m1=m1, m2=m2, m3=m3)
    beta <- get_param(param, m)
    b0 <- beta[1]; b1 <- beta[2]
    b2 <- beta[3]; b3 <- beta[4]

    phi0 <- hyper$phi0 # intercept
    phi1 <- hyper$phi1 # genotype
    phi2 <- hyper$phi2 # treatment
    phi3 <- hyper$phi3 # interaction
    kappa <- hyper$kappa
    nu <- hyper$nu
    mu <- get_mean(g=g, t=t, param=param, m=m, fn=fn.gp)

    if (ranef) {
        sigma.u <- param["sigma_u"]
        tU <- tu.lambda$tU
        lambda <- tu.lambda$lambda
        kappa.u <- hyper$kappa.u
        nu.u <- hyper$nu.u
        o <- sigma.u^2 * lambda + sigma^2
        R <- tU %*% (y - mu)
    }

    if (!ranef) {
        nxh <- - sum(dnorm(x=y, mean=mu, sd=sigma, log=TRUE))
    } else if (ranef) {
        nxh <- (1/2) * sum(R * R / o)
        nxh <- nxh + (1/2) * sum(log(o))
        nxh <- nxh + (1/2) * n * log(2 * pi)
        nxh <- nxh - dnorm(x=b0, mean=0, sd=phi0 * sigma, log=TRUE)
    }

    if (m1 == 1) {
        nxh <- nxh - dnorm(x=b1, mean=0, sd=phi1 * sigma, log=TRUE)
    }
    if (m2 == 1) {
        nxh <- nxh - dnorm(x=b2, mean=0, sd=phi2 * sigma, log=TRUE)
    }
    if (m3 == 1) {
        nxh <- nxh -
            dnorm(x=b3, mean=0, sd=phi3 * sigma, log=TRUE)
    }
    nxh <- nxh -
        dgamma(
            x=sigma^(-2), shape=kappa/2, rate=nu/2, log=TRUE)

    if (ranef) {
        nxh <- nxh -
            dgamma(
                x=sigma.u^(-2), shape=kappa.u/2,
                rate=nu.u/2, log=TRUE)
    }

    (1/n) * unname(nxh) # h
}

get_init <- function(data, ranef) {
    d <- data.frame(y=data$y, g=data$g, t=data$t)
    terms <- c("g", "t", "g:t")
    coefs <- c("b1", "b2", "b3")
    model.mat <- get_model_mat()

    if (!ranef) {
        init.list <- list("vector", nrow(model.mat))
    } else if (ranef) {
        init.list.list <- list("vector", nrow(model.mat))
        scale <- c(0.2, 1, 5)
    }
    for (i in 1:nrow(model.mat)) {
        m <- model.mat[i, ]
        terms.i <- terms[m == 1]
        if (i == 1) {
            frml <- "y ~ 1"
        } else {
            frml <- paste(
                c("y ~ 1", paste("+", terms.i)), collapse=" ")
        }

        coefs.i <- coefs[m == 1]

        if (!ranef) {
            coefs.i <- c("b0", coefs.i, "sigma")
        } else if (ranef) {
            coefs.i <- c("b0", coefs.i, "sigma", "sigma_u")
        }

        lm.fit <- lm(frml, data=d)

        if (!ranef) {
            init.list[[i]] <- c(coef(lm.fit), sigma(lm.fit)) %>%
                `names<-`(coefs.i)
        } else if (ranef) {
            sigma <- sigma(lm.fit)
            init.list <- list("vector", length(scale))
            for (j in seq_along(scale)) {
                sigma.u <- sigma * scale[j]
                init.list[[j]] <- c(
                    coef(lm.fit), sigma, sigma.u) %>%
                    `names<-`(coefs.i)
            }
            init.list.list[[i]] <- init.list
        }
    }

    if (!ranef) {
        return(init.list)
    } else if (ranef) {
        return(init.list.list)
    }

}

get_marginal <- function(fit, n) {
    p <- length(fit$par)
    inv.hessian <- solve(fit$hessian)
    h <- fit$value
    log.i.hat <- - n * h +
        (p/2) * log((2 * pi) / n) +
        (1/2) * log(det(inv.hessian))
    log.i.hat
}

get_m_vectors <- function() {
    model.mat <- get_model_mat()
    m.list <- vector("list", nrow(model.mat))
    for (i in 1:nrow(model.mat)) {
        m.list[[i]] <- model.mat[i, ]
    }
    m.list
}

get_model_mat <- function() {
    model.mat <- cbind(
        m1=rep(rep(c(0, 1), each=2^0), times=2^2),
        m2=rep(rep(c(0, 1), each=2^1), times=2^1),
        m3=rep(rep(c(0, 1), each=2^2), times=2^0)
    )
    model.mat
}

get_sign_mat <- function() {
    model.mat <- cbind(
        m1=rep(rep(c(0, -1, 1), each=3^0), times=3^2),
        m2=rep(rep(c(0, -1, 1), each=3^1), times=3^1),
        m3=rep(rep(c(0, -1, 1), each=3^2), times=3^0)
    )
    model.mat
}

get_names_from_matrix <- function(m) {
    apply(m, 1,
          function(x) paste(as.character(x), collapse=","))
}

get_mean <- function(g, t, param, m, fn) {
    beta <- get_param(param, m)
    b0 <- beta[1]; b1 <- beta[2]
    b2 <- beta[3]; b3 <- beta[4]

    if (!(fn %in% c("linear", "nonlinear"))) {
        stop("fn must be either 'linear' or 'nonlinear'")
    }

    if (fn == "linear") {
        output <- b0 + b1 * g + b2 * t + b3 * g * t
    } else if (fn == "nonlinear") {
        tmp <-  exp(b0) * (1 - (g/2)) * (1 - t) +
            exp(b0 + 2 * b1) * (g/2) * (1 - t) +
            exp(b0 + b2) * (1 - (g/2)) * t +
            exp(b0 + 2 * b1 + b2 + 2 * b3) * (g/2) * t
        output <- log(tmp)
    }
    output
}

get_param <- function(param, m) {
    beta <- rep(0, 4)
    beta[1] <- param["b0"]
    if (m[1] == 1) beta[2] <- param["b1"]
    if (m[2] == 1) beta[3] <- param["b2"]
    if (m[3] == 1) beta[4] <- param["b3"]
    beta
}
```
