## Supplemental File 1 for "Probabilistic classification of gene-by-treatment interactions on molecular count phenotypes": plot.R

```
#' Make a genotype-phenotype plot with model fit
#'
#' This function generates a genotype-phenotype plot with regression
#' lines in the control and treated conditions based on the MAP model.
#'
#' @export
#' @import ggplot2
#' @importFrom grDevices rgb
#' @importFrom scales number_format
#' @param gp A list object obtained from the \code{\link{format_gp}}
#'     function.
#' @param title A character string specifying a title.
#' @param seed A seed for RNG.
#'
#' @return A \code{ggplot2} object
make_gp_plot <- function(gp,
                         title=NULL,
                         seed=1) {

    cols <- c(
        rgb(97/255, 104/255, 108/255),
        rgb(191/255, 144/255, 0/255))

    input <- gp$input
    fit <- gp$fit

    set.seed(seed)
    p <- ggplot() +
        geom_jitter(
            mapping=aes(x=.data$x, y=.data$y, colour=.data$t),
            data=input,
            width=0.1, alpha=0.2) +
        geom_line(
            mapping=aes(x=.data$x, y=.data$y, colour=.data$t),
            data=fit) +
        xlab("Genotype") +
        ylab("Phenotype") +
        labs(title=title) +
        labs(fill="Treatment") +
        scale_color_manual(
            labels=c("Control", "Treated"),
            values=cols) +
        scale_x_continuous(breaks=c(0, 1, 2)) +
        scale_y_continuous(
            labels=scales::number_format(accuracy=0.1))

    p <- p +
        theme(
            aspect.ratio=1,
            plot.title=element_text(size=rel(1), hjust=0.5),
            legend.position="bottom",
            legend.title=element_blank(),
            legend.key=element_rect(
                colour="transparent", fill="transparent"),
            panel.grid.major=element_blank(),
            panel.grid.minor=element_blank(),
            panel.background=element_blank(),
            panel.border=element_rect(
                colour="gray", fill=NA, linewidth=1))

    p

}

#' Prepare data for a genotype-phenotype plot
#'
#' This is a function to prepare data for visualization using
#' \code{\link{make_gp_plot}}.
#'
#' @export
#' @import ggplot2
#' @importFrom magrittr "%>%"
#' @importFrom stats qnorm
#' @param data A list containing phenotype, genotype, treatment, and
#'     subject, which must be named "y", "g", "t", and "subject ",
#'     respectively.
#' @param fit A list obtained from the \code{\link{do_bms}}.
#' @param seed A seed for RNG.
#'
#' @return A list of data frames.
#'
format_gp <- function(data,
                      fit,
                      seed=1) {

    fn <- fit$fn
    rint <- fit$rint
    model.name <- get_model_names()
    model.mat <- get_model_mat()

    xs <- seq(0, 2, 0.01)
    # cols <- c(
    #     rgb(97/255, 104/255, 108/255),
    #     rgb(191/255, 144/255, 0/255))

    # g <- data$g
    # t <- data$t
    # y <- data$y
    # n <- length(y)

    input.df <- data.frame(
        y=data$y,
        x=data$g,
        t=data$t %>% factor
    )

    if (fn == "nonlinear") {
        get_line <- compute_nl
    } else if (fn == "linear" & !(rint)) {
        get_line <- compute_ln
    } else if (fn == "linear" & rint) {
        get_line <- compute_ln
        y <- input.df$y
        # perform RINT-transformation
        y.rint <- qnorm((rank(y) - (1/2)) / length(y))
        input.df$y <- y.rint
    }

    pp.vec <- fit$p.m.given.y
    chosen <- pp.vec %>% which.max

    if (class(fit$stan.list) == "list") {
        res <- fit$stan.list[[chosen]]
        if (class(res)[1] == "stanfit") {
            res <- rstan::summary(res)$summary
        }

        coef.vec <- c("b0", "b1", "b2", "b3")
        beta <- res[coef.vec, "mean"]
        m.vec <- model.mat[chosen, ] %>% as.numeric
        y.c <- sapply(xs, get_line, t=0,
                      beta=beta, m=m.vec) # control
        y.t <- sapply(xs, get_line, t=1,
                      beta=beta, m=m.vec) # treated
        fit.df <- data.frame(
            y=c(y.c, y.t),
            x=rep(xs, times=2),
            t=rep(c(0, 1), each=length(xs)) %>% factor)
    } else if (class(fit$optim.list) == "list") {
        res <- fit$optim.list[[chosen]]
        beta <- res$par[!names(res$par) %in% c("sigma", "sigma_u")]
        # coef.vec <- paste0("b", c(0, 1:3))
        coef.vec <- c("b0", "b1", "b2", "b3")
        for (coef in coef.vec) {
            if (is.na(beta[coef])) beta[coef] <- 0
        }
        beta <- beta[coef.vec]
        m.vec <- model.mat[chosen, ] %>% as.numeric
        y.c <- sapply(xs, get_line, t=0,
            beta=beta, m=m.vec) # control
        y.t <- sapply(xs, get_line, t=1,
            beta=beta, m=m.vec) # treated
        fit.df <- data.frame(
            y=c(y.c, y.t),
            x=rep(xs, times=2),
            t=rep(c(0, 1), each=length(xs)) %>% factor)
    }

    output <- list(input=input.df, fit=fit.df, fn=fn)
    output

}

#' Make a barplot of posterior probability
#'
#' This function takes output from \code{\link{format_pp}} as input
#' and generates a barplot of posterior probability.
#'
#' @export
#' @import ggplot2
#' @param pp A data frame obtained from \code{\link{format_pp}}. For
#'     visualizing the posterior of crossocer interaction, \code{co}
#'     must be set to \code{TRUE} when running running
#'     \code{\link{format_pp}}.
#'
#' @return A \code{ggplot2} object.
#'
make_pp_plot <- function(pp) {

    xs <- seq(0, 2, 0.01)
    cols <- c(
        rgb(200/255, 200/255, 200/255),
        rgb(107/255, 174/255, 214/255),
        rgb(54/255, 100/255, 139/255))

    # model.name <- get_model_names()
    # model.fact <- factor(model.name, levels=model.name)

    if (length(levels(pp$cat)) == 2) {
        co <- FALSE
    } else if (length(levels(pp$cat)) == 3) {
        co <- TRUE
    } else {
        stop("`pp` is in an incorrect format")
    }

    if (!co) {

        # cat.vec <- c("No GxT", "GxT")
        # cat.fact <- factor(cat.vec, levels=cat.vec)
        # d <- data.frame(
        #     value=fit$p.m.given.y,
        #     name=model.fact,
        #     cat=rep(cat.fact, each=4))

        p <- ggplot() +
            geom_col(
                mapping=aes(
                    x=.data$name, y=.data$value,
                    fill=.data$category),
                data=pp) +
            ylim(0, 1) +
            xlab("") +
            ylab("Posterior probability")

        p <- p + theme(
            aspect.ratio=1,
            legend.position="bottom",
            legend.title=element_blank(),
            axis.text.x=element_text(
                angle=90, vjust=0.5, hjust=1),
            panel.grid.major=element_blank(),
            panel.grid.minor=element_blank(),
            panel.background=element_blank(),
            panel.border=element_rect(
                colour="gray", fill=NA))

        p <- p +
            # facet_wrap(vars(method), scales="free_y") +
            scale_fill_manual(values=cols[1:2])

    } else {

        p <- ggplot() +
            geom_col(
                mapping=aes(
                    x=.data$name, y=.data$value,
                    fill=.data$category),
                data=pp,
                position="stack") +
            ylim(0, 1) +
            xlab("") +
            ylab("Posterior probability") # +
            # scale_y_continuous(
            #     labels=scales::number_format(accuracy=0.01))
        p <- p + theme(
            aspect.ratio=1,
            legend.position="bottom",
            legend.title=element_blank(),
            axis.text.x=element_text(
                angle=90, vjust=0.5, hjust=1),
            panel.grid.major=element_blank(),
            panel.grid.minor=element_blank(),
            panel.background=element_blank(),
            panel.border=element_rect(
                colour="gray", fill=NA, linewidth=1))
        p <- p +
            # facet_wrap(vars(method), scales="free_y") +
            scale_fill_manual(values=cols)

    }

    p

}

#' Prepare data for a barplot of posterior probability
#'
#' This is a function to prepare data for visualization using
#' \code{\link{make_pp_plot}}.
#'
#' @export
#' @import ggplot2
#' @importFrom magrittr "%>%"
#' @importFrom stats qnorm
#' @param fit A list obtained from the \code{\link{do_bms}}.
#' @param co A logical varialbe as to whether to visualize the
#'     probability of crossover interaction. If this is set to
#'     \code{TRUE}, \code{summary} must be set to \code{FALSE} when
#'     running \code{\link{do_bms}}.
#'
#' @return A data frame.
format_pp <- function(fit,
                      co=FALSE) {

    model.name <- get_model_names()
    model.fact <- factor(model.name, levels=model.name)

    if (!co) {

        cat.vec <- c("No GxT", "GxT")
        cat.fact <- factor(cat.vec, levels=cat.vec)
        output <- data.frame(
            name=model.fact,
            value=fit$p.m.given.y,
            category=rep(cat.fact, each=4))
        rownames(output) <- NULL

    } else {

        format_co <- function(fit) {
            p.m.given.y <- get_pp(fit)
            # p.co.given.m <- get_co(fit)
            # p.non <- p.m.given.y * (1 - p.co.given.m)
            # p.co <- p.m.given.y * p.co.given.m
            p.co <- get_co(fit) # joint
            p.non <- p.m.given.y - p.co
            p <- c(p.non, p.co)
            p
        }

        cat.vec <- c("No GxT", "Non-crossover", "Crossover")
        cat.fact <- factor(cat.vec, levels=cat.vec)

        pp <- fit %>% format_co
        pp <- c(pp[1:4], rep(0, 8), pp[5:16])
        output <- data.frame(
            name=rep(model.fact, times=3),
            value=pp,
            category=rep(cat.fact, each=8))
        rownames(output) <- NULL

    }

    output

}

#' Make a heatmap of posterior probability
#'
#' This is a function to make a heatmap of posterior probability for a
#' set of feature-SNP pairs.
#'
#' @export
#' @import ggplot2 hrbrthemes viridis
#' @importFrom magrittr "%>%"
#' @param input A matrix containing posterior probability. Rows and
#'     columns must represent feature-SNP pairs and model categories,
#'     respectively.
#'
#' @return A \code{ggplot2} object.
make_heatmap <- function(input) {
    m <- input %>% as.matrix
    pp.vec <- c(m)
    pair <- m %>% nrow %>% seq_len
    model <- colnames(m)
    d <- expand.grid(X=pair, Y=model) %>%
        `colnames<-`(c("pair", "model"))
    d$pp <- pp.vec

    # p <- d %>%
    #    ggplot(aes(x=pair, y=model, fill=pp)) +
    #    geom_tile()

    p <- ggplot() +
        geom_tile(
            mapping=aes(
                x=.data$pair,
                y=.data$model,
                fill=.data$pp),
            data=d)

    # p <- p + scale_fill_viridis(discrete=FALSE)
    p <- p + scale_fill_viridis(
                 discrete=FALSE,
                 limits=c(0, 1),
                 breaks=c(0, 0.5, 1),
                 labels=c("0.0", "0.5", "1.0"))
    p <- p + scale_y_discrete(limits=rev)
    # p <- p + geom_tile(color="white", size=0.1)
    # p <- p + geom_tile(color="white", size=0.01)
    p <- p + labs(y="Model", x="Feature-SNP pairs")

    # p <- p + theme(plot.title=element_text(size=rel(1.2), hjust=0.5))
    # p <- p + theme(theme(plot.margin=unit(c(1, 1, 1, 1), "cm")))
    # p <- p + theme(theme(plot.margin=margin(r=0)))
    p <- p + theme(axis.ticks=element_blank())
    p <- p + theme(axis.line=element_blank())
    p <- p + theme(
                 axis.text.y=element_text(
                     hjust=1, margin=margin(r=-15), size=rel(1)))
    p <- p + theme(axis.text.x=element_blank())
    p <- p + theme(
        panel.grid.major=element_blank(),
        panel.grid.minor=element_blank(),
        panel.border=element_blank(),
        panel.background=element_blank())
    p <- p + theme(panel.spacing.x=unit(0, "cm"))
    p <- p + theme(panel.spacing.y=unit(0, "cm"))
    p <- p + theme(legend.position="none")

    p
}

compute_nl <- function(g, t, beta, m) {
    b0 <- beta[1]; b1 <- beta[2]; b2 <- beta[3]; b3 <- beta[4]
    m1 <- m[1]; m2 <- m[2]; m3 <- m[3]
    x1 <- (1 - g/2) * (1 - t)
    x2 <- (g/2) * (1 - t)
    x3 <- (1 - g/2) * (t)
    x4 <- (g/2) * (t)
    r.mu <-
        x1 +
        exp(2 * m1 * b1) * x2 +
        exp(m2 * b2) * x3 +
        exp(2 * m1 * b1 + m2 * b2 + 2 * m3 * b3) * x4
    y <- b0 + log(r.mu)
    y
}

compute_ln <- function(g, t, beta, m) {
    b0 <- beta[1]; b1 <- beta[2]; b2 <- beta[3]; b3 <- beta[4]
    m1 <- m[1]; m2 <- m[2]; m3 <- m[3]
    y <- b0 + m1 * b1 * g + m2 * b2 * t + m3 * b3 * g * t
    y
}
```
