## Supplemental File 1 for "Probabilistic classification of gene-by-treatment interactions on molecular count phenotypes": stanmodels.R

```
# Generated by rstantools.  Do not edit by hand.

# names of stan models
stanmodels <- c("lm", "lmm", "nl", "nlmm")

# load each stan module
Rcpp::loadModule("stan_fit4lm_mod", what = TRUE)
Rcpp::loadModule("stan_fit4lmm_mod", what = TRUE)
Rcpp::loadModule("stan_fit4nl_mod", what = TRUE)
Rcpp::loadModule("stan_fit4nlmm_mod", what = TRUE)

# instantiate each stanmodel object
stanmodels <- sapply(stanmodels, function(model_name) {
  # create C++ code for stan model
  stan_file <- if(dir.exists("stan")) "stan" else file.path("inst", "stan")
  stan_file <- file.path(stan_file, paste0(model_name, ".stan"))
  stanfit <- rstan::stanc_builder(stan_file,
                                  allow_undefined = TRUE,
                                  obfuscate_model_name = FALSE)
  stanfit$model_cpp <- list(model_cppname = stanfit$model_name,
                            model_cppcode = stanfit$cppcode)
  # create stanmodel object
  methods::new(Class = "stanmodel",
               model_name = stanfit$model_name,
               model_code = stanfit$model_code,
               model_cpp = stanfit$model_cpp,
               mk_cppmodule = function(x) get(paste0("rstantools_model_", model_name)))
})
```
