## Supplemental File 1 for "Probabilistic classification of gene-by-treatment interactions on molecular count phenotypes": utils.R

```
#' Get the names of the eight models
#'
#' This function returns an ordered vector of character strings
#' corresponding to the names of the eight models.
#'
#' @export
#' @return A vector containing the eight model names.
#'
get_model_names <- function() {
    model.mat <- get_model_mat()
    get_names_from_matrix(model.mat)
}

#' Get the names of the 27 models
#'
#' This function returns an ordered vector of character strings
#' corresponding to the names of the 27 models accounting for the
#' sign of effect sizes.
#'
#' @export
#' @return A vector containing the eight model names.
#'
get_sign_names <- function() {
    sign.mat <- get_sign_mat()
    get_names_from_matrix(sign.mat)
}
```
