## Supplemental File 1 for "Probabilistic classification of gene-by-treatment interactions on molecular count phenotypes": RcppExports.cpp

```
// Generated by using Rcpp::compileAttributes() -> do not edit by hand
// Generator token: 10BE3573-1514-4C36-9D1C-5A225CD40393

#include <RcppEigen.h>
#include <Rcpp.h>

using namespace Rcpp;

#ifdef RCPP_USE_GLOBAL_ROSTREAM
Rcpp::Rostream<true>&  Rcpp::Rcout = Rcpp::Rcpp_cout_get();
Rcpp::Rostream<false>& Rcpp::Rcerr = Rcpp::Rcpp_cerr_get();
#endif


RcppExport SEXP _rcpp_module_boot_stan_fit4lm_mod();
RcppExport SEXP _rcpp_module_boot_stan_fit4lmm_mod();
RcppExport SEXP _rcpp_module_boot_stan_fit4nl_mod();
RcppExport SEXP _rcpp_module_boot_stan_fit4nlmm_mod();

static const R_CallMethodDef CallEntries[] = {
    {"_rcpp_module_boot_stan_fit4lm_mod", (DL_FUNC) &_rcpp_module_boot_stan_fit4lm_mod, 0},
    {"_rcpp_module_boot_stan_fit4lmm_mod", (DL_FUNC) &_rcpp_module_boot_stan_fit4lmm_mod, 0},
    {"_rcpp_module_boot_stan_fit4nl_mod", (DL_FUNC) &_rcpp_module_boot_stan_fit4nl_mod, 0},
    {"_rcpp_module_boot_stan_fit4nlmm_mod", (DL_FUNC) &_rcpp_module_boot_stan_fit4nlmm_mod, 0},
    {NULL, NULL, 0}
};

RcppExport void R_init_classifygxt(DllInfo *dll) {
    R_registerRoutines(dll, NULL, CallEntries, NULL, NULL);
    R_useDynamicSymbols(dll, FALSE);
}
```
