## Supplemental File 1 for "Probabilistic classification of gene-by-treatment interactions on molecular count phenotypes": make_test_data.R

```
suppressPackageStartupMessages(library(devtools))

document()
load_all()

file.noranef <- "data.noranef.rds"
file.ranef <- "data.ranef.rds"

num <- rep(1, 8)

n.sub <- 80 # number of subjects
anno <- data.frame(
    subject=rep(seq_len(n.sub), each=2),
    condition=rep(c(0, 1), times=n.sub))

sd.g <- 1.5
sd.t <- 2.0
sd.gxt <- 1.0
sd <- c(sd.g, sd.t, sd.gxt)

# without random effect
data.noranef <- make_data(
    anno=anno, fn="nonlinear",
    num=num, sd=sd)

saveRDS(data.noranef, file=file.noranef)

# with random effect
ranef <- TRUE
sigma.u <- sqrt(0.2)
kinship <- NULL

data.ranef <- make_data(
    anno=anno, fn="nonlinear",
    num=num, sd=sd,
    ranef=ranef, sigma.u=sigma.u)

saveRDS(data.ranef, file=file.ranef)

# do bms
file.bs.nl <- "mcmc.bs.nl.rds"
file.bs.lm <- "mcmc.bs.lm.rds"
file.bs.nlmm <- "mcmc.bs.nlmm.rds"
file.bs.lmm <- "mcmc.bs.lmm.rds"
file.lap.nl <- "map.lap.nl.rds"
file.lap.lm <- "map.lap.lm.rds"
file.lap.nlmm <- "map.lap.nlmm.rds"
file.lap.lmm <- "map.lap.lmm.rds"

# set hyperparameter values
phi1 <- 1.5
phi2 <- 2.0
phi3 <- 1.0
phi <- c(phi1, phi2, phi3)

p.m <- rep(1/8, 8)

data <- data.noranef
seed <- 1
n.cores <- 4

k <- 8
data <- data[[k]]

res.bs.nl <- do_bms(
    data=data, p.m=p.m, phi=phi,
    fn="nonlinear", rint=FALSE, method="mcmc.bs")

res.bs.lm <- do_bms(
    data=data, p.m=p.m, phi=phi,
    fn="linear", rint=FALSE, method="mcmc.bs")

# get a list of tU and lambda
tu.lambda <- get_tu_lambda(data)

# with donor random effect (no polygenic)
res.bs.nlmm <- do_bms(
    data=data, p.m=p.m, phi=phi,
    fn="nonlinear", rint=FALSE, method="mcmc.bs",
    tu.lambda=tu.lambda)

res.bs.lmm <- do_bms(
    data=data, p.m=p.m, phi=phi,
    fn="linear", rint=FALSE, method="mcmc.bs",
    tu.lambda=tu.lambda)

# save results
saveRDS(res.bs.nl, file=file.bs.nl)
saveRDS(res.bs.lm, file=file.bs.lm)
saveRDS(res.bs.nlmm, file=file.bs.nlmm)
saveRDS(res.bs.lmm, file=file.bs.lmm)

# with map + lap
res.lap.nl <- do_bms(
    data=data, p.m=p.m, phi=phi,
    fn="nonlinear", rint=FALSE, method="map.lap")

res.lap.lm <- do_bms(
    data=data, p.m=p.m, phi=phi,
    fn="linear", rint=FALSE, method="map.lap")

# with donor random effect (no polygenic)
res.lap.nlmm <- do_bms(
    data=data, p.m=p.m, phi=phi,
    fn="nonlinear", rint=FALSE, method="map.lap",
    tu.lambda=tu.lambda)

res.lap.lmm <- do_bms(
    data=data, p.m=p.m, phi=phi,
    fn="linear", rint=FALSE, method="map.lap",
    tu.lambda=tu.lambda)

# save results
saveRDS(res.lap.nl, file=file.lap.nl)
saveRDS(res.lap.lm, file=file.lap.lm)
saveRDS(res.lap.nlmm, file=file.lap.nlmm)
saveRDS(res.lap.lmm, file=file.lap.lmm)

# get estimates
get_est(res.bs.nl) %>%
    saveRDS("est.bs.nl.rds")
get_est(res.bs.nlmm) %>%
    saveRDS("est.bs.nlmm.rds")
get_est(res.bs.nl) %>%
    saveRDS("est.bs.nl.sample.rds")
get_est(res.bs.nlmm) %>%
    saveRDS("est.bs.nlmm.sample.rds")
get_est(res.lap.nl) %>%
    saveRDS("est.lap.nl.rds")
get_est(res.lap.nlmm) %>%
    saveRDS("est.lap.nlmm.rds")

# get posterior probability
# of aggregated categories
get_pp(res.bs.nl, aggregate="genotype") %>%
    saveRDS("pp.gen.bs.nl.rds")
get_pp(res.bs.nl, aggregate="treatment") %>%
    saveRDS("pp.tre.bs.nl.rds")
get_pp(res.lap.nl, aggregate="genotype") %>%
    saveRDS("pp.gen.lap.nl.rds")
get_pp(res.lap.nl, aggregate="treatment") %>%
    saveRDS("pp.tre.lap.nl.rds")

# get posterior probability of crossover interaction
# without random effect
res.bs.nl.sample <- do_bms(
    data=data, p.m=p.m, phi=phi,
    fn="nonlinear", rint=FALSE, method="mcmc.bs",
    summary=FALSE)

# with donor random effect (no polygenic)
res.bs.nlmm.sample <- do_bms(
    data=data, p.m=p.m, phi=phi,
    fn="nonlinear", rint=FALSE, method="mcmc.bs",
    tu.lambda=tu.lambda,
    summary=FALSE)

get_co(res.bs.nl.sample) %>%
    saveRDS("pp.co.nl.rds")
get_co(res.bs.nlmm.sample) %>%
    saveRDS("pp.co.nlmm.rds")

# get posterior probability accounting for signs
get_sign(res.bs.nl) %>%
    saveRDS("pp.sign.bs.nl.rds")
get_sign(res.bs.nlmm) %>%
    saveRDS("pp.sign.bs.nlmm.rds")
get_sign(res.bs.nl.sample) %>%
    saveRDS("pp.sign.bs.nl.sample.rds")
get_sign(res.bs.nlmm.sample) %>%
    saveRDS("pp.sign.bs.nlmm.sample.rds")
get_sign(res.lap.nl) %>%
    saveRDS("pp.sign.lap.nl.rds")
get_sign(res.lap.nlmm) %>%
    saveRDS("pp.sign.lap.nlmm.rds")
```
