## Supplemental File 1 for "Probabilistic classification of gene-by-treatment interactions on molecular count phenotypes": test_do_bms.R

```
# set tolerance for expect_equal()
tolerance <- 1e-3

# set hyperparameter values
phi1 <- 1.5
phi2 <- 2.0
phi3 <- 1.0
phi <- c(phi1, phi2, phi3)

p.m <- rep(1/8, 8)

data <- readRDS("fixtures/data.noranef.rds")
seed <- 1
n.cores <- 4

k <- 8
data <- data[[k]]

# get a list of tU and lambda
tu.lambda <- get_tu_lambda(data)

test_that("MCMC and bridge sampling in do_bms works for nonlinear models without random effect", {
    res <- readRDS("fixtures/mcmc.bs.nl.rds")
    tmp <- do_bms(
        data=data, p.m=p.m, phi=phi,
        fn="nonlinear", rint=FALSE, method="mcmc.bs")
    # expect_equal(tmp, res, tolerance=tolerance)
    expect_equal(
        tmp$ln.p.y.given.m, res$ln.p.y.given.m, tolerance=tolerance)
})

test_that("MCMC and bridge sampling in do_bms works for linear models without random effect", {
    res <- readRDS("fixtures/mcmc.bs.lm.rds")
    tmp <- do_bms(
        data=data, p.m=p.m, phi=phi,
        fn="linear", rint=FALSE, method="mcmc.bs")
    # expect_equal(tmp, res, tolerance=tolerance)
    expect_equal(
        tmp$ln.p.y.given.m, res$ln.p.y.given.m, tolerance=tolerance)
})

test_that("MCMC and bridge sampling in do_bms works for nonlinear models with random effect", {
    res <- readRDS("fixtures/mcmc.bs.nlmm.rds")
    tmp <- do_bms(
        data=data, p.m=p.m, phi=phi,
        fn="nonlinear", rint=FALSE, method="mcmc.bs",
        tu.lambda=tu.lambda)
    # expect_equal(tmp, res, tolerance=tolerance)
    expect_equal(
        tmp$ln.p.y.given.m, res$ln.p.y.given.m, tolerance=tolerance)

})

test_that("MCMC and bridge sampling in do_bms works for linear models with random effect", {
    res <- readRDS("fixtures/mcmc.bs.lmm.rds")
    tmp <- do_bms(
        data=data, p.m=p.m, phi=phi,
        fn="linear", rint=FALSE, method="mcmc.bs",
        tu.lambda=tu.lambda)
    # expect_equal(tmp, res, tolerance=tolerance)
    expect_equal(
        tmp$ln.p.y.given.m, res$ln.p.y.given.m, tolerance=tolerance)
})

test_that("MAP estimation and Laplace approximation in do_bms works for nonlinear models without random effect", {
    res <- readRDS("fixtures/map.lap.nl.rds")
    tmp <- do_bms(
        data=data, p.m=p.m, phi=phi,
        fn="nonlinear", rint=FALSE, method="map.lap")
    expect_equal(
        tmp$ln.p.y.given.m, res$ln.p.y.given.m, tolerance=tolerance)
})

test_that("MAP estimation and Laplace approximation in do_bms works for linear models without random effect", {
    res <- readRDS("fixtures/map.lap.lm.rds")
    tmp <- do_bms(
        data=data, p.m=p.m, phi=phi,
        fn="linear", rint=FALSE, method="map.lap")
    expect_equal(
        tmp$ln.p.y.given.m, res$ln.p.y.given.m, tolerance=tolerance)
})

test_that("MAP estimation and Laplace approximation in do_bms works for nonlinear models with random effect", {
    res <- readRDS("fixtures/map.lap.nlmm.rds")
    tmp <- do_bms(
        data=data, p.m=p.m, phi=phi,
        fn="nonlinear", rint=FALSE, method="map.lap",
        tu.lambda=tu.lambda)
    expect_equal(
        tmp$ln.p.y.given.m, res$ln.p.y.given.m, tolerance=tolerance)
})

test_that("MAP estimation and Laplace approximation in do_bms works for linear models with random effect", {
    res <- readRDS("fixtures/map.lap.lmm.rds")
    tmp <- do_bms(
        data=data, p.m=p.m, phi=phi,
        fn="linear", rint=FALSE, method="map.lap",
        tu.lambda=tu.lambda)
    expect_equal(
        tmp$ln.p.y.given.m, res$ln.p.y.given.m, tolerance=tolerance)
})
```
