## Supplemental File 1 for "Probabilistic classification of gene-by-treatment interactions on molecular count phenotypes": test_get_co.R

```
# set tolerance for expect_equal()
tolerance <- 2e-2

phi1 <- 1.5
phi2 <- 2.0
phi3 <- 1.0
phi <- c(phi1, phi2, phi3)
p.m <- rep(1/8, 8)
k <- 8

data.list <- readRDS("fixtures/data.noranef.rds")
data <- data.list[[k]]
tu.lambda <- get_tu_lambda(data)

test_that("get_co works without random effect", {
    res <- readRDS("fixtures/pp.co.nl.rds")
    mcmc.bs.nl.sample <- do_bms(
        data=data, p.m=p.m, phi=phi,
        fn="nonlinear", rint=FALSE, method="mcmc.bs",
        summary=FALSE)
    tmp <- get_co(mcmc.bs.nl.sample)
    expect_equal(tmp, res, tolerance=tolerance)
})

test_that("get_co works with random effect", {
    res <- readRDS("fixtures/pp.co.nlmm.rds")
    mcmc.bs.nlmm.sample <- do_bms(
        data=data, p.m=p.m, phi=phi,
        fn="nonlinear", rint=FALSE, method="mcmc.bs",
        tu.lambda=tu.lambda,
        summary=FALSE)
    tmp <- get_co(mcmc.bs.nlmm.sample)
    expect_equal(tmp, res, tolerance=tolerance)
})
```
