## Supplemental File 1 for "Probabilistic classification of gene-by-treatment interactions on molecular count phenotypes": test_get_est.R

```
# set tolerance for expect_equal()
tolerance <- 1e-3

mcmc.bs.nl <- readRDS("fixtures/mcmc.bs.nl.rds")
mcmc.bs.nlmm <- readRDS("fixtures/mcmc.bs.nlmm.rds")
map.lap.nl <- readRDS("fixtures/map.lap.nl.rds")
map.lap.nlmm <- readRDS("fixtures/map.lap.nlmm.rds")

test_that("get_est works for a summary from MCMC without random effect", {
    res <- readRDS("fixtures/est.bs.nl.rds")
    tmp <- get_est(mcmc.bs.nl)
    expect_equal(tmp, res, tolerance=tolerance)
})

test_that("get_est works for a summary from MCMC with random effect", {
    res <- readRDS("fixtures/est.bs.nlmm.rds")
    tmp <- get_est(mcmc.bs.nlmm)
    expect_equal(tmp, res, tolerance=tolerance)
})

test_that("get_est works for MAP estimates without random effect", {
    res <- readRDS("fixtures/est.lap.nl.rds")
    tmp <- get_est(map.lap.nl)
    expect_equal(tmp, res, tolerance=tolerance)
})

test_that("get_est works for MAP estimates with random effect", {
    res <- readRDS("fixtures/est.lap.nlmm.rds")
    tmp <- get_est(map.lap.nlmm)
    expect_equal(tmp, res, tolerance=tolerance)
})
```
