## Supplemental File 1 for "Probabilistic classification of gene-by-treatment interactions on molecular count phenotypes": test_get_pp.R

```
# set tolerance for expect_equal()
tolerance <- 1e-3

mcmc.bs.nl <- readRDS("fixtures/mcmc.bs.nl.rds")
map.lap.nl <- readRDS("fixtures/map.lap.nl.rds")

test_that("get_pp works for results from MCMC and bridge sampling with the genotype option", {
    res <- readRDS("fixtures/pp.gen.bs.nl.rds")
    tmp <- get_pp(mcmc.bs.nl, aggregate="genotype")
    expect_equal(tmp, res, tolerance=tolerance)
})

test_that("get_pp works for results from MCMC and bridge sampling with the treatment option", {
    res <- readRDS("fixtures/pp.tre.bs.nl.rds")
    tmp <- get_pp(mcmc.bs.nl, aggregate="treatment")
    expect_equal(tmp, res, tolerance=tolerance)
})

test_that("get_pp works for results from MAP estimation and Laplace approximation with the genotype option", {
    res <- readRDS("fixtures/pp.gen.lap.nl.rds")
    tmp <- get_pp(map.lap.nl, aggregate="genotype")
    expect_equal(tmp, res, tolerance=tolerance)
})

test_that("get_pp works for results from MAP estimation and Laplace approximation with the treatment option", {
    res <- readRDS("fixtures/pp.tre.lap.nl.rds")
    tmp <- get_pp(map.lap.nl, aggregate="treatment")
    expect_equal(tmp, res, tolerance=tolerance)
})
```
