## Supplemental File 1 for "Probabilistic classification of gene-by-treatment interactions on molecular count phenotypes": test_make_data.R

```
num <- rep(1, 8)
n.sub <- 80 # number of subjects
anno <- data.frame(
    subject=rep(seq_len(n.sub), each=2),
    condition=rep(c(0, 1), times=n.sub))

sd.g <- 1.5
sd.t <- 2.0
sd.gxt <- 1.0
sd <- c(sd.g, sd.t, sd.gxt)

# without random effect
test_that("make_data works without random effect", {
    data.noranef <- readRDS("fixtures/data.noranef.rds")
    tmp <- make_data(
        anno=anno, fn="nonlinear",
        num=num, sd=sd)
    expect_equal(tmp, data.noranef)
})

# with random effect
test_that("make_data works with random effect", {
    ranef <- TRUE
    sigma.u <- sqrt(0.2)
    kinship <- NULL
    data.ranef <- readRDS("fixtures/data.ranef.rds")
    tmp <- make_data(
        anno=anno, fn="nonlinear",
        num=num, sd=sd,
        ranef=ranef, sigma.u=sigma.u)
    expect_equal(tmp, data.ranef)
})
```
