## Supplemental File 1 for "Probabilistic classification of gene-by-treatment interactions on molecular count phenotypes": classifygxt.Rmd

````
---
title: "ClassifyGxT - classifying gene-by-treatment interactions"
author: "Yuriko Harigaya, Michael Love, William Valdar"
date: "`r format(Sys.Date())`"
header_includes:
    - \newcommand{\Expect}{\mathrm{E}}
    - \newcommand{\Var}{\mathrm{Var}}
    - \newcommand{\Cov}{\mathrm{Cov}}
    - \newcommand{\Norm}{\mathrm{N}}
    - \newcommand{\iid}{\overset{\mathrm{iid}}{\sim}}
    - \newcommand{\veps}{\varepsilon}
    - \newcommand{\bveps}{\boldsymbol{\varepsilon}}
    - \newcommand{\bA}{\mathbf{A}}
    - \newcommand{\bK}{\mathbf{K}}
    - \newcommand{\bI}{\mathbf{I}}
    - \newcommand{\bB}{\mathbf{B}}
    - \newcommand{\bR}{\mathbf{R}}
    - \newcommand{\bV}{\mathbf{V}}
    - \newcommand{\bX}{\mathbf{X}}
    - \newcommand{\bY}{\mathbf{Y}}
    - \newcommand{\bZ}{\mathbf{Z}}
    - \newcommand{\bU}{\mathbf{U}}
    - \newcommand{\bO}{\mathbf{O}}
    - \newcommand{\bT}{\mathbf{T}}
    - \newcommand{\bG}{\mathbf{G}}
    - \newcommand{\bSigma}{\boldsymbol{\Sigma}}
    - \newcommand{\bOmega}{\boldsymbol{\Omega}}
    - \newcommand{\balpha}{\boldsymbol{\alpha}}
    - \newcommand{\bbeta}{\boldsymbol{\beta}}
    - \newcommand{\bmu}{\boldsymbol{\mu}}
    - \newcommand{\btheta}{\boldsymbol{\theta}}
    - \newcommand{\bgamma}{\boldsymbol{\gamma}}
    - \newcommand{\bpsi}{\boldsymbol{\psi}}
    - \newcommand{\bphi}{\boldsymbol{\phi}}
    - \newcommand{\bnu}{\boldsymbol{\nu}}
    - \newcommand{\bu}{\mathbf{u}}
    - \newcommand{\bv}{\mathbf{v}}
    - \newcommand{\bw}{\mathbf{w}}
    - \newcommand{\bx}{\mathbf{x}}
    - \newcommand{\by}{\mathbf{y}}
    - \newcommand{\bz}{\mathbf{z}}
    - \newcommand{\bk}{\mathbf{k}}
    - \newcommand{\bm}{\mathbf{m}}
    - \newcommand{\bnull}{\mathbf{0}} 
    - \newcommand{\bone}{\mathbf{1}}
output: rmarkdown::html_vignette
bibliography: ref.bib
vignette: >
  %\VignetteIndexEntry{ClassifyGxT - classifying gene-by-treatment interactions}
  %\VignetteEngine{knitr::rmarkdown}
  %\VignetteEncoding{UTF-8}
---

```{r, include = FALSE}
knitr::opts_chunk$set(
  collapse=TRUE,
  comment="#>"
)
```

# Introduction

The ClassifyGxT software is an implementation of a Bayesian model selection (BMS) framework for classifying GxT interactions.
The method was developed primarily for molecular count phenotypes, such as RNA-seq and ATAC-seq data, although it can be used for other types of phenotypes.

- The input is a list of feature-SNP pairs for which significant GxT interactions have been identified by the standard response molecular QTL mapping procedure (see @hipsci_consortium_shared_2018 and @matoba_wnt_2023 for example).
For each of the feature-SNP pairs, individual genotype and phenotype data are required.
- The main output is the posterior probability for different GxT types. See **Overview** on the main page for the model categories representing the types of GxT interactions.
- The package provides functions to process a single feature-SNP pair.
In practice, we recommend running parallel jobs on the computer cluster with appropriate grouping.

# Input data

For each feature-SNP pair, a list object containing the following elements needs to be generated:

- `y`: A vector of length $n$ containing preprocessed molecular count phenotypes, where $n$ is the number of samples. See **Data processing** below for preprocessing.
- `g`: A vector of length $n$ containing genotypes coded as ${0, 1, 2}$ to represent the number of minor (alternative) alleles or the imputation-based allelic dosage in $[0, 2]$. 
- `t`: A vector of indicator variables for the treatment.
- `subject`: An optional character or numeric vector corresponding to the subjects. This is necessary when the model includes donor or polygenic (kinship) random effects.
- `feat.id`: An optional character string representing the feature.
- `snp.id`: An optional character string representing the SNP.

Note that the samples must be in the same order in the `y`, `g`, `t`, and `subject` elements and that the list can contain additional elements.

# Data preprocessing

For molecular count phenotypes, the raw count data from sequencing experiments can be processed as follows:

- Step 1: make a matrix of feature counts with rows and columns representing features and samples, respectively
- Step 2: scale the count data by the library size according to @palowitch_estimation_2018
- Step 3: optionally filter features based on the scaled counts according to @matoba_wnt_2023
- Step 4: transform using the function $\log(x + 1)$
- Step 5: regress out the treatment indicator (and optionally other nuisance factors, such as sex and age of the subjects)
- Step 6: perform principal component analysis (PCA)
- Step 7: regress out an appropriate number of PCs from the original, transformed data (from Step 4)

# Loading packages

```{r packages, cache=FALSE, message=FALSE}
library(ggplot2)
library(classifygxt)
```

# Generating data

In this section, we simulate data using `make_data()`.
The simulated data will have the same format as discussed in the previous section and will be used for demonstrating how to run BMS in the next sections.
For simplicity, we omit random effects.
See the function documentation (`?make_data`) for how to include random effects.

We first specify the number of feature-SNP pairs for each of the eight models.

```{r}
(model.name <- get_model_names())
```

For this demonstration, we only make data for 10 pairs for each model.

```{r}
num <- rep(10, 8)
names(num) <- model.name
num
```

With this specification, the output will be a list of 80 lists.
Each of the lists corresponds to each feature-SNP pair.
The first ten lists are based on `"0,0,0"`,
the next ten are based on `"1,0,0"`, and so forth. 

The genotype, treatment, and interaction effects will be drawn from Normal distributions with a zero mean and user-specified standard deviations.
These values represent "typical" magnitudes of the effects.
We specify standard deviations of the effects as follows.

```{r}
sd.g <- 1.5 # genotype
sd.t <- 2.0 # treatment
sd.gxt <- 1.0 # interaction
sd <- c(sd.g, sd.t, sd.gxt)
```

Note that the residual error standard deviation $\sigma$, which represents a typical magnitude of noise, is set to 1 by default.

The following code generates a data frame specifying the mapping between samples, subjects, and treatment conditions.

```{r}
n.sub <- 80 # number of subjects
anno <- data.frame(
    subject=rep(seq_len(n.sub), each=2),
    condition=rep(c(0, 1), times=n.sub))
head(anno)
```

Now we can generate fake data using `make_data()`.

```{r}
data.list <- make_data(
    anno=anno, fn="nonlinear",
    num=num, sd=sd)
```

# Performing BMS

In this section, we perform BMS using `do_bms()`. 
For simplicity, we do not model random effects.
See **Including donor random effects** below for how to include random effects.

We first specify the "model prior."
Note that this is an optional argument and that, by default, a uniform prior is used.
This specification reflects a prior belief that all models are equally likely.
Here, we explicitly specify the default model prior for demonstration purposes.

```{r}
p.m <- rep(1/8, 8)
```

We also specify the "effect prior" by choosing the values of the $\phi_g$, $\phi_t$, and $\phi_{g \times t}$ hyperparameters, which correspond to the genotype, treatment, and GxT interaction effects, respectively.
The hyperparameters represent our prior beliefs about the effects relative to the residual error standard deviation (noise).
Specifically, we place priors,
\begin{equation}
  \beta_g \sim \Norm(0, \phi_g^2 \sigma^2), \quad 
  \beta_t \sim \Norm(0, \phi_t^2 \sigma^2), \quad
  \beta_{g \times t} \sim \Norm(0, \phi_{g \times t}^2 \sigma^2).
\end{equation}
In practice, we recommend optimizing the values via an empirical Bayes approach (see **Optimizing the effect prior hyperparameters** below).
Here, we set them to the same values as the effect standard deviations in the data-generating models.
This is a reasonable choice since we set the residual error standard deviation to 1 when generating the data.

```{r}
phi.g <- 1.5 # genotype
phi.t <- 2.0 # treatment
phi.gxt <- 1.0 # interaction
phi <- c(phi.g, phi.t, phi.gxt)
```

The following code performs BMS for the 71st data using `do_bms()` with nonlinear regression (`fn="nonlinear"`).
Recall that the 71-80th data were generated based on the eighth model, `"1,1,1"` (see **Generating data** above).
The `method` input argument must be set to either `"mcmc.bs"`, which represents Markov Chain Monte Carlo (MCMC) followed by bridge sampling, or `"map.lap"`, which represents MAP estimation followed by Laplace approximation.
Although the latter method can be orders of magnitude faster, we recommend using `"mcmc.bs"` for the final results, if possible, since it is not straightforward to obtain error bounds for Laplace approximation.

```{r}
k <- 71
data <- data.list[[k]]
res <- do_bms(
    data=data, p.m=p.m, phi=phi,
    fn="nonlinear", method="map.lap")
```

This returns a list object containing the following elements.
See the function documentation (`?do_bms`) for details.

```{r}
names(res)
```

For multiple feature-SNP pairs, we recommend processing the data in batches using a workflow such as [Snakemake](https://snakemake.readthedocs.io/en/stable/).
Within a batch, BMS can be performed in serial or parallel processes.

For a moderate number of feature-SNP pairs, the following code can be used.

```{r, warning=FALSE, message=FALSE}
res.list <- lapply(
    X=data.list, FUN=do_bms, p.m=p.m, phi=phi,
    fn="nonlinear", method="map.lap")
```

## Extracting the posterior probability of the models

We can extract the posterior probability of the model as follows.

```{r}
(pp <- get_pp(res))
```

Note that, in this particular example, the highest posterior probability is assigned to the fourth model `"1,1,0"`, even though the data-generating model is "`1,1,1`".
However, we will see that BMS tends to choose the correct models across multiple feature-SNP pairs (**Heatmap of the posterior probability of the models** below). 

The following code can be used to extract the posterior probability from a list object storing results for multiple feature-SNP pairs.

```{r}
pp.mat <- sapply(X=res.list, FUN=get_pp)
```

In this case, the output is a matrix that contains rows and columns corresponding to the models and the feature-SNP pairs, respectively.

# Including random effects

In analyses of experimental data, it is ofen desirable to include random effects in the model. 
To include donor random effects, we first need to run `get_tu_lambda()`.

```{r}
tu.lambda <- get_tu_lambda(data)
```

We then use the output as input when running `do_bms()`.

```{r}
res.ranef <- do_bms(
    data=data, p.m=p.m, phi=phi,
    fn="nonlinear", method="map.lap",
    tu.lambda=tu.lambda)
```

We can also include polygenic random effects rather than donor random effects to accout for the genetic relatedness and popuplation structure. 
See the function documentation (`?get_tu_lambda`) for details.

# Visualizing the results

## Generating a genotype-phenotype (GP) plot

We recommend visually inspecting the model fit by generating scatter plots, which we call a "GP" plot, for each feature-SNP pair (or SNP).
To make a GP plot, we first create a list of data frames using
`format_gp()`.

```{r}
gp.plot <- format_gp(data=data, fit=res)
```

We then create a `ggplot2` object using `make_gp_plot()`.
The appearance of the plot can be modified as usual.

```{r, fig.width=4, fig.height=4, fig.align="center"}
p1 <- make_gp_plot(gp=gp.plot)
p1
```

## Generating a posterior probability (PP) plot

It is also useful to generate a barplot of posterior probability, which we call a "PP" plot.
To make a PP plot, we first create a data frame using `format_pp()`.

```{r}
pp.plot <- format_pp(fit=res)
```

We then create a `ggplot2` object using `make_pp_plot()`.
The appearance of the plot can be modified as usual.

```{r, fig.width=4, fig.height=4, fig.align="center"}
p2 <- make_pp_plot(pp=pp.plot) 
p2
```

## Heatmap of the posterior probability of the models

We can generate a heatmap to visualize posterior probability across multiple feature-SNP pairs using `make_heatmap()`.
Note that we transpose the matrix using `t()` in the following code.

```{r, fig.width=5, fig.height=5, fig.align="center"}
p3 <- make_heatmap(t(pp.mat)) 
p3 + theme(legend.position="bottom",
           legend.title=element_blank())
```

As expected, we see that the highest probability tends to be assigned to the correct (i.e., data-generating) model.
That is, the MAP model tends to be `"0,0,0"` for the first ten feature-SNP pairs in the left-most columns, `"1,0,0"` for the 11-20th pairs, and so forth.

# Optimizing the effect prior hyperparameters

We recommend optimizing the effect prior hyperparmeters by an empirical Bayes approach.
In this approach, we obtain the $\phi_g$, $\phi_t$, and $\phi_{g \times t}$ hyperparameter values that maximize the sum of $\log$-transformed marginal likelihood across all feature-SNP pairs that are being analyzed. 
The most conceptually straightforward method is to perform a grid search (see our manuscript for details).
We recommend using a workflow such as [Snakemake](https://snakemake.readthedocs.io/en/stable/).
For faster computation, we use MAP estimation and Laplace approximation, setting `method="map.lap"` (see **Performing BMS** above).
From the BMS result for a feature-SNP pair, the $\log$ of the marginal likelihood can be extracted as follows.

```{r}
(ln.p.y <- res$ln.p.y)
```

Since error bounds for Laplace approximation are not easily obtained, we recommend rerunning MCMC and bridge sampling with optimal hyperparameter values for the final result.

# Session information

```{r}
sessionInfo()
```

# References
````
