## Supplemental File 1 for "Probabilistic classification of gene-by-treatment interactions on molecular count phenotypes": ref.bib

@article{crouse_bayesian_2022,
	title = {A {Bayesian} model selection approach to mediation analysis},
	volume = {18},
	issn = {1553-7404},
	url = {https://dx.plos.org/10.1371/journal.pgen.1010184},
	doi = {10.1371/journal.pgen.1010184},
	abstract = {Genetic studies often seek to establish a causal chain of events originating from genetic variation through to molecular and clinical phenotypes. When multiple phenotypes share a common genetic association, one phenotype may act as an intermediate for the genetic effects on the other. Alternatively, the phenotypes may be causally unrelated but share genetic loci. Mediation analysis represents a class of causal inference approaches used to determine which of these scenarios is most plausible. We have developed a general approach to mediation analysis based on Bayesian model selection and have implemented it in an R package, bmediatR. Bayesian model selection provides a flexible framework that can be tailored to different analyses. Our approach can incorporate prior information about the likelihood of models and the strength of causal effects. It can also accommodate multiple genetic variants or multi-state haplotypes. Our approach reports posterior probabilities that can be useful in interpreting uncertainty among competing models. We compared bmediatR with other popular methods, including the Sobel test, Mendelian randomization, and Bayesian network analysis using simulated data. We found that bmediatR performed as well or better than these alternatives in most scenarios. We applied bmediatR to proteome data from Diversity Outbred (DO) mice, a multi-parent population, and demonstrate the power of mediation with multi-state haplotypes. We also applied bmediatR to data from human cell lines to identify transcripts that are mediated through or are expressed independently from local chromatin accessibility. We demonstrate that Bayesian model selection provides a powerful and versatile approach to identify causal relationships in genetic studies using model organism or human data.},
	language = {en},
	number = {5},
	urldate = {2023-06-14},
	journal = {PLOS Genetics},
	author = {Crouse, Wesley L. and Keele, Gregory R. and Gastonguay, Madeleine S. and Churchill, Gary A. and Valdar, William},
	editor = {Kutalik, Zoltán},
	month = may,
	year = {2022},
	pages = {e1010184},
	file = {journal.pgen.1010184.pdf:/Users/yurikoharigaya/Library/CloudStorage/Dropbox/love_valdar_lab/papers/mediation/journal.pgen.1010184.pdf:application/pdf},
}

@article{aygun_inferring_2023,
	title = {Inferring cell-type-specific causal gene regulatory networks during human neurogenesis},
	abstract = {Background:  Genetic variation influences both chromatin accessibility, assessed in chromatin accessibility quantitative trait loci (caQTL) studies, and gene expression, assessed in expression QTL (eQTL) studies. Genetic variants can impact either nearby genes (cis-eQTLs) or distal genes (trans-eQTLs). Colocalization between caQTL and eQTL, or cis- and trans-eQTLs suggests that they share causal variants. However, pairwise colocalization between these molecular QTLs does not guarantee a causal relationship. Mediation analysis can be applied to assess the evidence supporting causality versus independence between molecular QTLs. Given that the function of QTLs can be cell-type-specific, we performed mediation analyses to find epigenetic and distal regulatory causal pathways for genes within two major cell types of the developing human cortex, progenitors and neurons.
Results:  We find that the expression of 168 and 38 genes is mediated by chromatin accessibility in progenitors and neurons, respectively. We also find that the expression of 11 and 12 downstream genes is mediated by upstream genes in progenitors and neurons. Moreover, we discover that a genetic locus associated with inter-individual differences in brain structure shows evidence for mediation of SLC26A7 through chromatin accessibility, identifying molecular mechanisms of a common variant association to a brain trait.
Conclusions:  In this study, we identify cell-type-specific causal gene regulatory networks whereby the impacts of variants on gene expression were mediated by chromatin accessibility or distal gene expression. Identification of these causal paths will enable identifying and prioritizing actionable regulatory targets perturbing these key processes during neurodevelopment.},
	language = {en},
	author = {Aygün, Nil},
	year = {2023},
	file = {s13059-023-02959-0.pdf:/Users/yurikoharigaya/Library/CloudStorage/Dropbox/love_valdar_lab/papers/mediation/s13059-023-02959-0.pdf:application/pdf},
}

@techreport{matoba_wnt_2023,
	type = {preprint},
	title = {Wnt activity reveals context-specific genetic effects on gene regulation in neural progenitors},
	url = {http://biorxiv.org/lookup/doi/10.1101/2023.02.07.527357},
	abstract = {Abstract
          Gene regulatory effects in bulk-post mortem brain tissues are undetected at many non-coding brain trait-associated loci. We hypothesized that context-specific genetic variant function during stimulation of a developmental signaling pathway would explain additional regulatory mechanisms. We measured chromatin accessibility and gene expression following activation of the canonical Wnt pathway in primary human neural progenitors from 82 donors. TCF/LEF motifs, brain structure-, and neuropsychiatric disorder-associated variants were enriched within Wnt-responsive regulatory elements (REs). Genetically influenced REs were enriched in genomic regions under positive selection along the human lineage. Stimulation of the Wnt pathway increased the detection of genetically influenced REs/genes by 66.2\%/52.7\%, and led to the identification of 397 REs primed for effects on gene expression. Context-specific molecular quantitative trait loci increased brain-trait colocalizations by up to 70\%, suggesting that genetic variant effects during early neurodevelopmental patterning lead to differences in adult brain and behavioral traits.},
	language = {en},
	urldate = {2023-06-14},
	institution = {Genetics},
	author = {Matoba, Nana and Le, Brandon D and Valone, Jordan M and Wolter, Justin M and Mory, Jessica and Liang, Dan and Aygün, Nil and Broadaway, K Alaine and Bond, Marielle L and Mohlke, Karen L and Zylka, Mark J and Love, Michael I and Stein, Jason L},
	month = feb,
	year = {2023},
	doi = {10.1101/2023.02.07.527357},
	file = {2023.02.07.527357v2.full.pdf:/Users/yurikoharigaya/Library/CloudStorage/Dropbox/love_valdar_lab/papers/eqtl/2023.02.07.527357v2.full.pdf:application/pdf},
}

@article{palowitch_estimation_2018,
	title = {Estimation of cis-{eQTL} effect sizes using a log of linear model: {Effect} {Sizes} of cis-{eQTLs} {Using} a {Log} of {Linear} {Model}},
	volume = {74},
	issn = {0006341X},
	shorttitle = {Estimation of cis-{eQTL} effect sizes using a log of linear model},
	url = {https://onlinelibrary.wiley.com/doi/10.1111/biom.12810},
	doi = {10.1111/biom.12810},
	abstract = {The study of expression Quantitative Trait Loci (eQTL) is an important problem in genomics and biomedicine. While detection (testing) of eQTL associations has been widely studied, less work has been devoted to the estimation of eQTL eﬀect size. To reduce false positives, detection methods frequently rely on linear modeling of rank-based normalized or log-transformed gene expression data. Unfortunately, these approaches do not correspond to the simplest model of eQTL action, and thus yield estimates of eQTL association that can be uninterpretable and inaccurate. In this article, we propose a new, log-of-linear model for eQTL action, termed ACME, that captures allelic contributions to cis-acting eQTLs in an additive fashion, yielding eﬀect size estimates that correspond to a biologically coherent model of cis-eQTLs. We describe a non-linear least-squares algorithm to ﬁt the model by maximum likelihood, and obtain corresponding p-values. We perform careful investigation of the model using a combination of simulated data and data from the Genotype Tissue Expression (GTEx) project. Our results reveal little evidence for dominance eﬀects, a parsimonious result that accords with a simple biological model for allele-speciﬁc expression and supports use of the ACME model. We show that Type-I error is wellcontrolled under our approach in a realistic setting, so that rank-based normalizations are unnecessary. Furthermore, we show that such normalizations can be detrimental to power and estimation accuracy under the proposed model. We then show, through eﬀect size analyses of whole-genome cis-eQTLs in the GTEx data, that using standard normalizations instead of ACME noticeably aﬀects the ranking and sign of estimates.},
	language = {en},
	number = {2},
	urldate = {2023-06-14},
	journal = {Biometrics},
	author = {Palowitch, John and Shabalin, Andrey and Zhou, Yi-Hui and Nobel, Andrew B. and Wright, Fred A.},
	month = jun,
	year = {2018},
	pages = {616--625},
	file = {Biometrics - 2017 - Palowitch - Estimation of cis‐eQTL effect sizes using a log of linear model.pdf:/Users/yurikoharigaya/Library/CloudStorage/Dropbox/love_valdar_lab/papers/eqtl/Biometrics - 2017 - Palowitch - Estimation of cis‐eQTL effect sizes using a log of linear model.pdf:application/pdf},
}

@article{sun_statistical_2012,
	title = {A {Statistical} {Framework} for {eQTL} {Mapping} {Using} {RNA}-seq {Data}},
	volume = {68},
	issn = {0006341X},
	url = {https://onlinelibrary.wiley.com/doi/10.1111/j.1541-0420.2011.01654.x},
	doi = {10.1111/j.1541-0420.2011.01654.x},
	abstract = {RNA-seq may replace gene expression microarrays in the near future. Using RNA-seq, the expression of a gene can be estimated using the total number of sequence reads mapped to that gene, known as the total read count (TReC). Traditional expression quantitative trait locus (eQTL) mapping methods, such as linear regression, can be applied to TReC measurements after they are properly normalized. In this article, we show that eQTL mapping, by directly modeling TReC using discrete distributions, has higher statistical power than the two-step approach: data normalization followed by linear regression. In addition, RNA-seq provides information on allele-speciﬁc expression (ASE) that is not available from microarrays. By combining the information from TReC and ASE, we can computationally distinguish cis- and trans-eQTL and further improve the power of cis-eQTL mapping. Both simulation and real data studies conﬁrm the improved power of our new methods. We also discuss the design issues of RNA-seq experiments. Speciﬁcally, we show that by combining TReC and ASE measurements, it is possible to minimize cost and retain the statistical power of cis-eQTL mapping by reducing sample size while increasing the number of sequence reads per sample. In addition to RNA-seq data, our method can also be employed to study the genetic basis of other types of sequencing data, such as chromatin immunoprecipitation followed by DNA sequencing data. In this article, we focus on eQTL mapping of a single gene using the association-based method. However, our method establishes a statistical framework for future developments of eQTL mapping methods using RNA-seq data (e.g., linkage-based eQTL mapping), and the joint study of multiple genetic markers and/or multiple genes.},
	language = {en},
	number = {1},
	urldate = {2023-06-14},
	journal = {Biometrics},
	author = {Sun, Wei},
	month = mar,
	year = {2012},
	pages = {1--11},
	file = {j.1541-0420.2011.01654.x.pdf:/Users/yurikoharigaya/Library/CloudStorage/Dropbox/love_valdar_lab/papers/eqtl/j.1541-0420.2011.01654.x.pdf:application/pdf},
}

@article{zhou_pca_2022,
	title = {{PCA} outperforms popular hidden variable inference methods for molecular {QTL} mapping},
	volume = {23},
	issn = {1474-760X},
	url = {https://genomebiology.biomedcentral.com/articles/10.1186/s13059-022-02761-4},
	doi = {10.1186/s13059-022-02761-4},
	abstract = {Background:  Estimating and accounting for hidden variables is widely practiced as an important step in molecular quantitative trait locus (molecular QTL, henceforth “QTL”) analysis for improving the power of QTL identification. However, few benchmark studies have been performed to evaluate the efficacy of the various methods developed for this purpose.
Results:  Here we benchmark popular hidden variable inference methods including surrogate variable analysis (SVA), probabilistic estimation of expression residuals (PEER), and hidden covariates with prior (HCP) against principal component analysis (PCA)—a well-established dimension reduction and factor discovery method—via 362 synthetic and 110 real data sets. We show that PCA not only underlies the statistical methodology behind the popular methods but is also orders of magnitude faster, better-performing, and much easier to interpret and use.
Conclusions:  To help researchers use PCA in their QTL analysis, we provide an R package PCAForQTL along with a detailed guide, both of which are freely available at https://github.com/heatherjzhou/PCAForQTL. We believe that using PCA rather than SVA, PEER, or HCP will substantially improve and simplify hidden variable inference in QTL mapping as well as increase the transparency and reproducibility of QTL research.},
	language = {en},
	number = {1},
	urldate = {2023-06-14},
	journal = {Genome Biology},
	author = {Zhou, Heather J. and Li, Lei and Li, Yumei and Li, Wei and Li, Jingyi Jessica},
	month = oct,
	year = {2022},
	pages = {210},
	file = {s13059-022-02761-4.pdf:/Users/yurikoharigaya/Library/CloudStorage/Dropbox/love_valdar_lab/papers/eqtl/s13059-022-02761-4.pdf:application/pdf},
}

@article{mohammadi_quantifying_2017,
	title = {Quantifying the regulatory effect size of \textit{cis} -acting genetic variation using allelic fold change},
	volume = {27},
	issn = {1088-9051, 1549-5469},
	url = {http://genome.cshlp.org/lookup/doi/10.1101/gr.216747.116},
	doi = {10.1101/gr.216747.116},
	abstract = {Mapping
              cis
              -acting expression quantitative trait loci (
              cis
              -eQTL) has become a popular approach for characterizing proximal genetic regulatory variants. In this paper, we describe and characterize log allelic fold change (aFC), the magnitude of expression change associated with a given genetic variant, as a biologically interpretable unit for quantifying the effect size of
              cis
              -eQTLs and a mathematically convenient approach for systematic modeling of
              cis
              -regulation. This measure is mathematically independent from expression level and allele frequency, additive, applicable to multiallelic variants, and generalizable to multiple independent variants. We provide efficient tools and guidelines for estimating aFC from both eQTL and allelic expression data sets and apply it to Genotype Tissue Expression (GTEx) data. We show that aFC estimates independently derived from eQTL and allelic expression data are highly consistent, and identify technical and biological correlates of eQTL effect size. We generalize aFC to analyze genes with two eQTLs in GTEx and show that in nearly all cases the two eQTLs act independently in regulating gene expression. In summary, aFC is a solid measure of
              cis
              -regulatory effect size that allows quantitative interpretation of cellular regulatory events from population data, and it is a valuable approach for investigating novel aspects of eQTL data sets.},
	language = {en},
	number = {11},
	urldate = {2023-06-14},
	journal = {Genome Research},
	author = {Mohammadi, Pejman and Castel, Stephane E. and Brown, Andrew A. and Lappalainen, Tuuli},
	month = nov,
	year = {2017},
	pages = {1872--1884},
	file = {Genome Res.-2017-Mohammadi-1872-84.pdf:/Users/yurikoharigaya/Library/CloudStorage/Dropbox/love_valdar_lab/papers/transformation/Genome Res.-2017-Mohammadi-1872-84.pdf:application/pdf},
}

@article{kim-hellmuth_cell_2020,
	title = {Cell type–specific genetic regulation of gene expression across human tissues},
	volume = {369},
	issn = {0036-8075, 1095-9203},
	url = {https://www.science.org/doi/10.1126/science.aaz8528},
	doi = {10.1126/science.aaz8528},
	abstract = {Cell type composition, estimated from bulk tissue, maps the cellular specificity of genetic variants.
          , 
            Cell type–specific quantitative trait loci
            
              Understanding how human genetic variation affects phenotype requires tissue- or even cell type–specific measurements. Kim-Hellmuth
              et al.
              used computational methods to identify cell-type proportions within bulk tissues in the Genotype-Tissue Expression (GTEx) project dataset to identify cell-type interaction quantitative trait loci and map these to genetic variants correlated with expression or splicing differences between individuals. By characterizing the cellular context, this study illustrates how genetic variants that operate in a cell type–specific manner affect gene regulation and can be linked to complex traits. This deconvolution and analysis of cell types from bulk tissues allows greater precision in understanding how phenotypes are linked to genetic variation.
            
            
              Science
              , this issue p.
              eaaz8528
            
          , 
            
              INTRODUCTION
              Efforts to map quantitative trait loci (QTLs) across human tissues by the GTEx Consortium and others have identified expression and splicing QTLs (eQTLs and sQTLs, respectively) for a majority of genes. However, these studies were largely performed with gene expression measurements from bulk tissue samples, thus obscuring the cellular specificity of genetic regulatory effects and in turn limiting their functional interpretation. Identifying the cell type (or types) in which a QTL is active will be key to uncovering the molecular mechanisms that underlie complex trait variation. Recent studies demonstrated the feasibility of identifying cell type–specific QTLs from bulk tissue RNA-sequencing data by using computational estimates of cell type proportions. To date, such approaches have only been applied to a limited number of cell types and tissues. By applying this methodology to GTEx tissues for a diverse set of cell types, we aim to characterize the cellular specificity of genetic effects across human tissues and to describe the contribution of these effects to complex traits.
            
            
              RATIONALE
              A growing number of in silico cell type deconvolution methods and associated reference panels with cell type–specific marker genes enable the robust estimation of the enrichment of specific cell types from bulk tissue gene expression data. We benchmarked and used enrichment estimates for seven cell types (adipocytes, epithelial cells, hepatocytes, keratinocytes, myocytes, neurons, and neutrophils) across 35 tissues from the GTEx project to map QTLs that are specific to at least one cell type. We mapped such cell type–interaction QTLs for expression and splicing (ieQTLs and isQTLs, respectively) by testing for interactions between genotype and cell type enrichment.
            
            
              RESULTS
              Using 43 pairs of tissues and cell types, we found 3347 protein-coding and long intergenic noncoding RNA (lincRNA) genes with an ieQTL and 987 genes with an isQTL (at 5\% false discovery rate in each pair). To validate these findings, we tested the QTLs for replication in available external datasets and applied an independent validation using allele-specific expression from eQTL heterozygotes. We analyzed the cell type–interaction QTLs for patterns of tissue sharing and found that ieQTLs are enriched for genes with tissue-specific eQTLs and are generally not shared across unrelated tissues, suggesting that tissue-specific eQTLs originate in tissue-specific cell types. Last, we tested the ieQTLs and isQTLs for colocalization with genetic associations for 87 complex traits. We show that cell type–interaction QTLs are enriched for complex trait associations and identify colocalizations for hundreds of loci that were undetected in bulk tissue, corresponding to an increase of {\textgreater}50\% over colocalizations with standard QTLs. Our results also reveal the cellular specificity and potential origin for a similar number of colocalized standard QTLs.
            
            
              CONCLUSION
              The ieQTLs and isQTLs identified for seven cell types across GTEx tissues suggest that the large majority of cell type–specific QTLs remains to be discovered. Our colocalization results indicate that comprehensive mapping of cell type–specific QTLs will be highly valuable for gaining a mechanistic understanding of complex trait associations. We anticipate that the approaches presented here will complement studies mapping QTLs in single cells.
              
                
                  Detection of cell type–specific effects on gene expression.
                  The enrichment of seven cell types is calculated across GTEx tissues, enabling mapping of cell type–interaction QTLs for expression and splicing by testing for significant interactions between genotypes and cell type enrichments. Linking these QTLs to complex trait associations enables discovery of {\textgreater}50\% more colocalizations compared with standard QTLs and reveals the cellular specificity of traits.
                
                
              
            
          , 
            The Genotype-Tissue Expression (GTEx) project has identified expression and splicing quantitative trait loci in cis (QTLs) for the majority of genes across a wide range of human tissues. However, the functional characterization of these QTLs has been limited by the heterogeneous cellular composition of GTEx tissue samples. We mapped interactions between computational estimates of cell type abundance and genotype to identify cell type–interaction QTLs for seven cell types and show that cell type–interaction expression QTLs (eQTLs) provide finer resolution to tissue specificity than bulk tissue cis-eQTLs. Analyses of genetic associations with 87 complex traits show a contribution from cell type–interaction QTLs and enables the discovery of hundreds of previously unidentified colocalized loci that are masked in bulk tissue.},
	language = {en},
	number = {6509},
	urldate = {2023-06-17},
	journal = {Science},
	author = {Kim-Hellmuth, Sarah and Aguet, François and Oliva, Meritxell and Muñoz-Aguirre, Manuel and Kasela, Silva and Wucher, Valentin and Castel, Stephane E. and Hamel, Andrew R. and Viñuela, Ana and Roberts, Amy L. and Mangul, Serghei and Wen, Xiaoquan and Wang, Gao and Barbeira, Alvaro N. and Garrido-Martín, Diego and Nadel, Brian B. and Zou, Yuxin and Bonazzola, Rodrigo and Quan, Jie and Brown, Andrew and Martinez-Perez, Angel and Soria, José Manuel and {GTEx Consortium} and Getz, Gad and Dermitzakis, Emmanouil T. and Small, Kerrin S. and Stephens, Matthew and Xi, Hualin S. and Im, Hae Kyung and Guigó, Roderic and Segrè, Ayellet V. and Stranger, Barbara E. and Ardlie, Kristin G. and Lappalainen, Tuuli and Aguet, François and Anand, Shankara and Ardlie, Kristin G. and Gabriel, Stacey and Getz, Gad A. and Graubert, Aaron and Hadley, Kane and Handsaker, Robert E. and Huang, Katherine H. and Kashin, Seva and Li, Xiao and MacArthur, Daniel G. and Meier, Samuel R. and Nedzel, Jared L. and Nguyen, Duyen T. and Segrè, Ayellet V. and Todres, Ellen and Balliu, Brunilda and Barbeira, Alvaro N. and Battle, Alexis and Bonazzola, Rodrigo and Brown, Andrew and Brown, Christopher D. and Castel, Stephane E. and Conrad, Donald F. and Cotter, Daniel J. and Cox, Nancy and Das, Sayantan and De Goede, Olivia M. and Dermitzakis, Emmanouil T. and Einson, Jonah and Engelhardt, Barbara E. and Eskin, Eleazar and Eulalio, Tiffany Y. and Ferraro, Nicole M. and Flynn, Elise D. and Fresard, Laure and Gamazon, Eric R. and Garrido-Martín, Diego and Gay, Nicole R. and Gloudemans, Michael J. and Guigó, Roderic and Hame, Andrew R. and He, Yuan and Hoffman, Paul J. and Hormozdiari, Farhad and Hou, Lei and Im, Hae Kyung and Jo, Brian and Kasela, Silva and Kellis, Manolis and Kim-Hellmuth, Sarah and Kwong, Alan and Lappalainen, Tuuli and Li, Xin and Liang, Yanyu and Mangul, Serghei and Mohammadi, Pejman and Montgomery, Stephen B. and Muñoz-Aguirre, Manuel and Nachun, Daniel C. and Nobel, Andrew B. and Oliva, Meritxell and Park, YoSon and Park, Yongjin and Parsana, Princy and Rao, Abhiram S. and Reverter, Ferran and Rouhana, John M. and Sabatti, Chiara and Saha, Ashis and Stephens, Matthew and Stranger, Barbara E. and Strober, Benjamin J. and Teran, Nicole A. and Viñuela, Ana and Wang, Gao and Wen, Xiaoquan and Wright, Fred and Wucher, Valentin and Zou, Yuxin and Ferreira, Pedro G. and Li, Gen and Melé, Marta and Yeger-Lotem, Esti and Barcus, Mary E. and Bradbury, Debra and Krubit, Tanya and McLean, Jeffrey A. and Qi, Liqun and Robinson, Karna and Roche, Nancy V. and Smith, Anna M. and Sobin, Leslie and Tabor, David E. and Undale, Anita and Bridge, Jason and Brigham, Lori E. and Foster, Barbara A. and Gillard, Bryan M. and Hasz, Richard and Hunter, Marcus and Johns, Christopher and Johnson, Mark and Karasik, Ellen and Kopen, Gene and Leinweber, William F. and McDonald, Alisa and Moser, Michael T. and Myer, Kevin and Ramsey, Kimberley D. and Roe, Brian and Shad, Saboor and Thomas, Jeffrey A. and Walters, Gary and Washington, Michael and Wheeler, Joseph and Jewell, Scott D. and Rohrer, Daniel C. and Valley, Dana R. and Davis, David A. and Mash, Deborah C. and Branton, Philip A. and Barker, Laura K. and Gardiner, Heather M. and Mosavel, Maghboeba and Siminoff, Laura A. and Flicek, Paul and Haeussler, Maximilian and Juettemann, Thomas and Kent, W. James and Lee, Christopher M. and Powell, Conner C. and Rosenbloom, Kate R. and Ruffier, Magali and Sheppard, Dan and Taylor, Kieron and Trevanion, Stephen J. and Zerbino, Daniel R. and Abell, Nathan S. and Akey, Joshua and Chen, Lin and Demanelis, Kathryn and Doherty, Jennifer A. and Feinberg, Andrew P. and Hansen, Kasper D. and Hickey, Peter F. and Jasmine, Farzana and Jiang, Lihua and Kaul, Rajinder and Kibriya, Muhammad G. and Li, Jin Billy and Li, Qin and Lin, Shin and Linder, Sandra E. and Pierce, Brandon L. and Rizzardi, Lindsay F. and Skol, Andrew D. and Smith, Kevin S. and Snyder, Michael and Stamatoyannopoulos, John and Tang, Hua and Wang, Meng and Carithers, Latarsha J. and Guan, Ping and Koester, Susan E. and Little, A. Roger and Moore, Helen M. and Nierras, Concepcion R. and Rao, Abhi K. and Vaught, Jimmie B. and Volpi, Simona},
	month = sep,
	year = {2020},
	pages = {eaaz8528},
	file = {Full Text:/Users/yurikoharigaya/Zotero/storage/HYR5PKNM/Kim-Hellmuth et al. - 2020 - Cell type–specific genetic regulation of gene expr.pdf:application/pdf},
}

@article{kawaguchi_improved_2023,
	title = {Improved two‐step testing of genome‐wide gene–environment interactions},
	volume = {47},
	issn = {0741-0395, 1098-2272},
	url = {https://onlinelibrary.wiley.com/doi/10.1002/gepi.22509},
	doi = {10.1002/gepi.22509},
	abstract = {Two‐step tests for gene–environment (G × E) interactions exploit marginal single‐nucleotide polymorphism (SNP) effects to improve the power of a genome‐wide interaction scan. They combine a screening step based on marginal effects used to “bin” SNPs for weighted hypothesis testing in the second step to deliver greater power over single‐step tests while preserving the genome‐wide Type I error. However, the presence of many SNPs with detectable marginal effects on the trait of interest can reduce power by “displacing” true interactions with weaker marginal effects and by adding to the number of tests that need to be corrected for multiple testing. We introduce a new significance‐based allocation into bins for Step‐2 G × E testing that overcomes the displacement issue and propose a computationally efficient approach to account for multiple testing within bins. Simulation results demonstrate that these simple improvements can provide substantially greater power than current methods under several scenarios. An application to a multistudy collaboration for understanding colorectal cancer reveals a G × Sex interaction located near the SMAD7 gene.},
	language = {en},
	number = {2},
	urldate = {2023-07-05},
	journal = {Genetic Epidemiology},
	author = {Kawaguchi, Eric S. and Kim, Andre E. and Lewinger, Juan Pablo and Gauderman, W. James},
	month = mar,
	year = {2023},
	pages = {152--166},
	file = {Genetic Epidemiology - 2022 - Kawaguchi - Improved two‐step testing of genome‐wide gene environment interactions.pdf:/Users/yurikoharigaya/Library/CloudStorage/Dropbox/love_valdar_lab/papers/interaction/Genetic Epidemiology - 2022 - Kawaguchi - Improved two‐step testing of genome‐wide gene environment interactions.pdf:application/pdf},
}

@article{huang_power_2018,
	title = {Power, false discovery rate and {Winner}’s {Curse} in {eQTL} studies},
	volume = {46},
	issn = {0305-1048, 1362-4962},
	url = {https://academic.oup.com/nar/article/46/22/e133/5090771},
	doi = {10.1093/nar/gky780},
	abstract = {Investigation of the genetic architecture of gene expression traits has aided interpretation of disease and trait-associated genetic variants; however, key aspects of expression quantitative trait loci (eQTL) study design and analysis remain understudied. We used extensive, empirically driven simulations to explore eQTL study design and the performance of various analysis strategies. Across multiple testing correction methods, false discoveries of genes with eQTLs (eGenes) were substantially inﬂated when false discovery rate (FDR) control was applied to all tests and only appropriately controlled using hierarchical procedures. All multiple testing correction procedures had low power and inﬂated FDR for eGenes whose causal SNPs had small allele frequencies using small sample sizes (e.g. frequency {\textless}10\% in 100 samples), indicating that even moderately low frequency eQTL SNPs (eSNPs) in these studies are enriched for false discoveries. In scenarios with ≥80\% power, the top eSNP was the true simulated eSNP 90\% of the time, but substantially less frequently for very common eSNPs (minor allele frequencies {\textgreater}25\%). Overestimation of eQTL effect sizes, so-called ‘Winner’s Curse’, was common in low and moderate power settings. To address this, we developed a bootstrap method (BootstrapQTL) that led to more accurate effect size estimation. These insights provide a foundation for future eQTL studies, especially those with sampling constraints and subtly different conditions.},
	language = {en},
	number = {22},
	urldate = {2023-07-05},
	journal = {Nucleic Acids Research},
	author = {Huang, Qin Qin and Ritchie, Scott C and Brozynska, Marta and Inouye, Michael},
	month = dec,
	year = {2018},
	pages = {e133--e133},
	file = {gky780.pdf:/Users/yurikoharigaya/Library/CloudStorage/Dropbox/love_valdar_lab/papers/eqtl/gky780.pdf:application/pdf},
}

@article{the_gtex_consortium_gtex_2020,
	title = {The {GTEx} {Consortium} atlas of genetic regulatory effects across human tissues},
	volume = {369},
	issn = {0036-8075, 1095-9203},
	url = {https://www.science.org/doi/10.1126/science.aaz1776},
	doi = {10.1126/science.aaz1776},
	abstract = {The Genotype-Tissue Expression (GTEx) project dissects how genetic variation affects gene expression and splicing.
          , 
            A survey of transcription across tissues
            
              Some human genetic variants affect the amount of RNA produced and the splicing of gene transcripts, crucial steps in development and maintaining a healthy individual. However, some of these changes only occur in a small number of tissues within the body. The Genotype-Tissue Expression (GTEx) project has been expanded over time, and, looking at the final data in version 8, Aguet
              et al.
              present a deep characterization of genetic associations and gene expression and splicing in 838 individuals over 49 tissues (see the Perspective by Wilson). This large study was able to characterize the details underlying many aspects of gene expression and provides a resource with which to better understand the fundamental molecular mechanisms of how genetic variants affect gene regulation and complex traits in humans.
            
            
              Science
              , this issue p.
              1318
              ; see also p.
              1298
            
          , 
            The Genotype-Tissue Expression (GTEx) project was established to characterize genetic effects on the transcriptome across human tissues and to link these regulatory mechanisms to trait and disease associations. Here, we present analyses of the version 8 data, examining 15,201 RNA-sequencing samples from 49 tissues of 838 postmortem donors. We comprehensively characterize genetic associations for gene expression and splicing in cis and trans, showing that regulatory associations are found for almost all genes, and describe the underlying molecular mechanisms and their contribution to allelic heterogeneity and pleiotropy of complex traits. Leveraging the large diversity of tissues, we provide insights into the tissue specificity of genetic effects and show that cell type composition is a key factor in understanding gene regulatory mechanisms in human tissues.},
	language = {en},
	number = {6509},
	urldate = {2023-08-08},
	journal = {Science},
	author = {{The GTEx Consortium} and Aguet, François and Anand, Shankara and Ardlie, Kristin G. and Gabriel, Stacey and Getz, Gad A. and Graubert, Aaron and Hadley, Kane and Handsaker, Robert E. and Huang, Katherine H. and Kashin, Seva and Li, Xiao and MacArthur, Daniel G. and Meier, Samuel R. and Nedzel, Jared L. and Nguyen, Duyen T. and Segrè, Ayellet V. and Todres, Ellen and Balliu, Brunilda and Barbeira, Alvaro N. and Battle, Alexis and Bonazzola, Rodrigo and Brown, Andrew and Brown, Christopher D. and Castel, Stephane E. and Conrad, Donald F. and Cotter, Daniel J. and Cox, Nancy and Das, Sayantan and De Goede, Olivia M. and Dermitzakis, Emmanouil T. and Einson, Jonah and Engelhardt, Barbara E. and Eskin, Eleazar and Eulalio, Tiffany Y. and Ferraro, Nicole M. and Flynn, Elise D. and Fresard, Laure and Gamazon, Eric R. and Garrido-Martín, Diego and Gay, Nicole R. and Gloudemans, Michael J. and Guigó, Roderic and Hame, Andrew R. and He, Yuan and Hoffman, Paul J. and Hormozdiari, Farhad and Hou, Lei and Im, Hae Kyung and Jo, Brian and Kasela, Silva and Kellis, Manolis and Kim-Hellmuth, Sarah and Kwong, Alan and Lappalainen, Tuuli and Li, Xin and Liang, Yanyu and Mangul, Serghei and Mohammadi, Pejman and Montgomery, Stephen B. and Muñoz-Aguirre, Manuel and Nachun, Daniel C. and Nobel, Andrew B. and Oliva, Meritxell and Park, YoSon and Park, Yongjin and Parsana, Princy and Rao, Abhiram S. and Reverter, Ferran and Rouhana, John M. and Sabatti, Chiara and Saha, Ashis and Stephens, Matthew and Stranger, Barbara E. and Strober, Benjamin J. and Teran, Nicole A. and Viñuela, Ana and Wang, Gao and Wen, Xiaoquan and Wright, Fred and Wucher, Valentin and Zou, Yuxin and Ferreira, Pedro G. and Li, Gen and Melé, Marta and Yeger-Lotem, Esti and Barcus, Mary E. and Bradbury, Debra and Krubit, Tanya and McLean, Jeffrey A. and Qi, Liqun and Robinson, Karna and Roche, Nancy V. and Smith, Anna M. and Sobin, Leslie and Tabor, David E. and Undale, Anita and Bridge, Jason and Brigham, Lori E. and Foster, Barbara A. and Gillard, Bryan M. and Hasz, Richard and Hunter, Marcus and Johns, Christopher and Johnson, Mark and Karasik, Ellen and Kopen, Gene and Leinweber, William F. and McDonald, Alisa and Moser, Michael T. and Myer, Kevin and Ramsey, Kimberley D. and Roe, Brian and Shad, Saboor and Thomas, Jeffrey A. and Walters, Gary and Washington, Michael and Wheeler, Joseph and Jewell, Scott D. and Rohrer, Daniel C. and Valley, Dana R. and Davis, David A. and Mash, Deborah C. and Branton, Philip A. and Barker, Laura K. and Gardiner, Heather M. and Mosavel, Maghboeba and Siminoff, Laura A. and Flicek, Paul and Haeussler, Maximilian and Juettemann, Thomas and Kent, W. James and Lee, Christopher M. and Powell, Conner C. and Rosenbloom, Kate R. and Ruffier, Magali and Sheppard, Dan and Taylor, Kieron and Trevanion, Stephen J. and Zerbino, Daniel R. and Abell, Nathan S. and Akey, Joshua and Chen, Lin and Demanelis, Kathryn and Doherty, Jennifer A. and Feinberg, Andrew P. and Hansen, Kasper D. and Hickey, Peter F. and Jasmine, Farzana and Jiang, Lihua and Kaul, Rajinder and Kibriya, Muhammad G. and Li, Jin Billy and Li, Qin and Lin, Shin and Linder, Sandra E. and Pierce, Brandon L. and Rizzardi, Lindsay F. and Skol, Andrew D. and Smith, Kevin S. and Snyder, Michael and Stamatoyannopoulos, John and Tang, Hua and Wang, Meng and Carithers, Latarsha J. and Guan, Ping and Koester, Susan E. and Little, A. Roger and Moore, Helen M. and Nierras, Concepcion R. and Rao, Abhi K. and Vaught, Jimmie B. and Volpi, Simona},
	month = sep,
	year = {2020},
	pages = {1318--1330},
	file = {Full Text:/Users/yurikoharigaya/Zotero/storage/AFLQWFHF/The GTEx Consortium et al. - 2020 - The GTEx Consortium atlas of genetic regulatory ef.pdf:application/pdf},
}

@article{buenrostro_atac-seq_2015,
	title = {{ATAC}-seq: {A} {Method} for {Assaying} {Chromatin} {Accessibility} {Genome}-{Wide}},
	volume = {109},
	issn = {1934-3647},
	shorttitle = {{ATAC}-seq},
	doi = {10.1002/0471142727.mb2129s109},
	abstract = {This unit describes Assay for Transposase-Accessible Chromatin with high-throughput sequencing (ATAC-seq), a method for mapping chromatin accessibility genome-wide. This method probes DNA accessibility with hyperactive Tn5 transposase, which inserts sequencing adapters into accessible regions of chromatin. Sequencing reads can then be used to infer regions of increased accessibility, as well as to map regions of transcription-factor binding and nucleosome position. The method is a fast and sensitive alternative to DNase-seq for assaying chromatin accessibility genome-wide, or to MNase-seq for assaying nucleosome positions in accessible regions of the genome.},
	language = {eng},
	journal = {Current Protocols in Molecular Biology},
	author = {Buenrostro, Jason D. and Wu, Beijing and Chang, Howard Y. and Greenleaf, William J.},
	month = jan,
	year = {2015},
	pmid = {25559105},
	pmcid = {PMC4374986},
	keywords = {ATAC-seq, Chromatin, chromatin accessibility, Cytological Techniques, High-Throughput Nucleotide Sequencing, Molecular Biology, Protein Binding, Recombination, Genetic, Staining and Labeling, transposase, Transposases},
	pages = {21.29.1--21.29.9},
	file = {Accepted Version:/Users/yurikoharigaya/Zotero/storage/N6YH7R2N/Buenrostro et al. - 2015 - ATAC-seq A Method for Assaying Chromatin Accessib.pdf:application/pdf},
}

@article{mortazavi_mapping_2008,
	title = {Mapping and quantifying mammalian transcriptomes by {RNA}-{Seq}},
	volume = {5},
	issn = {1548-7091, 1548-7105},
	url = {https://www.nature.com/articles/nmeth.1226},
	doi = {10.1038/nmeth.1226},
	language = {en},
	number = {7},
	urldate = {2023-08-08},
	journal = {Nature Methods},
	author = {Mortazavi, Ali and Williams, Brian A and McCue, Kenneth and Schaeffer, Lorian and Wold, Barbara},
	month = jul,
	year = {2008},
	pages = {621--628},
}

@article{hipsci_consortium_shared_2018,
	title = {Shared genetic effects on chromatin and gene expression indicate a role for enhancer priming in immune response},
	volume = {50},
	issn = {1061-4036, 1546-1718},
	url = {https://www.nature.com/articles/s41588-018-0046-7},
	doi = {10.1038/s41588-018-0046-7},
	language = {en},
	number = {3},
	urldate = {2023-08-09},
	journal = {Nature Genetics},
	author = {{HIPSCI Consortium} and Alasoo, Kaur and Rodrigues, Julia and Mukhopadhyay, Subhankar and Knights, Andrew J. and Mann, Alice L. and Kundu, Kousik and Hale, Christine and Dougan, Gordon and Gaffney, Daniel J.},
	month = mar,
	year = {2018},
	pages = {424--431},
	file = {Accepted Version:/Users/yurikoharigaya/Zotero/storage/XW3VS7PN/HIPSCI Consortium et al. - 2018 - Shared genetic effects on chromatin and gene expre.pdf:application/pdf},
}

@article{knowles_determining_2018,
	title = {Determining the genetic basis of anthracycline-cardiotoxicity by molecular response {QTL} mapping in induced cardiomyocytes},
	volume = {7},
	issn = {2050-084X},
	url = {https://elifesciences.org/articles/33480},
	doi = {10.7554/eLife.33480},
	abstract = {Anthracycline-induced cardiotoxicity (ACT) is a key limiting factor in setting optimal chemotherapy regimes, with almost half of patients expected to develop congestive heart failure given high doses. However, the genetic basis of sensitivity to anthracyclines remains unclear. We created a panel of iPSC-derived cardiomyocytes from 45 individuals and performed RNA-seq after 24 hr exposure to varying doxorubicin dosages. The transcriptomic response is substantial: the majority of genes are differentially expressed and over 6000 genes show evidence of differential splicing, the later driven by reduced splicing fidelity in the presence of doxorubicin. We show that inter-individual variation in transcriptional response is predictive of in vitro cell damage, which in turn is associated with in vivo ACT risk. We detect 447 response-expression quantitative trait loci (QTLs) and 42 response-splicing QTLs, which are enriched in lower ACT GWAS
              
                p
              
              -values, supporting the in vivo relevance of our map of genetic regulation of cellular response to anthracyclines.
            
          , 
            Many cancers, including leukaemia, lymphoma and breast cancer, are treated with potent chemotherapy drugs such as anthracyclines. However, anthracyclines have strong side effects known as anthracycline cardiotoxicity, which affect the health of the heart. Almost half of the patients given high doses of anthracyclines develop chronic heart failure.
            While anthracycline cardiotoxicity is very common, people’s genes may contribute to how sensitive they are to these drugs but it is not understood which genes can cause this effect. Previous studies using only a small number of participants have not been able to pin down the genetic factors that make some patients respond well to anthracyclines, and others prone to developing heart failure when taking these drugs.
            To find out which genes affect anthracycline cardiotoxicity, Knowles, Burrows et al. transformed blood cells from 45 individuals into stem cells, which were then developed into heart muscle cells. Then, the activity of genes was analyzed by measuring the amount of RNA (the template molecules used to make proteins) produced by those genes.
            After the cells had been exposed for 24 hours to the anthracycline drug doxorubicin, hundreds of gene activity differences could be found in the heart muscle cells between individuals. Some of these differences were linked to poorer health of the cells after treatment with the drug. As a result, a number of genetic variants that could predispose patients to the side effects of doxorubicin were discovered. The experiments also revealed how doxorubicin disrupts an important process that separates ‘junk’ parts of the RNA from the parts that are used as a template for proteins.
            Being able to predict who is likely to be sensitive to drugs such as doxorubicin could help doctors to tailor chemotherapy treatments more effectively, minimising the risk of heart failure. In future, larger studies could lead to accurate predictions of a patient’s response to a particular chemotherapy drug to personalize their cancer treatment.},
	language = {en},
	urldate = {2023-08-09},
	journal = {eLife},
	author = {Knowles, David A and Burrows, Courtney K and Blischak, John D and Patterson, Kristen M and Serie, Daniel J and Norton, Nadine and Ober, Carole and Pritchard, Jonathan K and Gilad, Yoav},
	month = may,
	year = {2018},
	pages = {e33480},
	file = {Full Text:/Users/yurikoharigaya/Zotero/storage/UZZCGZZR/Knowles et al. - 2018 - Determining the genetic basis of anthracycline-car.pdf:application/pdf},
}

@article{beasley_rank-based_2009,
	title = {Rank-based inverse normal transformations are increasingly used, but are they merited?},
	volume = {39},
	issn = {1573-3297},
	doi = {10.1007/s10519-009-9281-0},
	abstract = {Many complex traits studied in genetics have markedly non-normal distributions. This often implies that the assumption of normally distributed residuals has been violated. Recently, inverse normal transformations (INTs) have gained popularity among genetics researchers and are implemented as an option in several software packages. Despite this increasing use, we are unaware of extensive simulations or mathematical proofs showing that INTs have desirable statistical properties in the context of genetic studies. We show that INTs do not necessarily maintain proper Type 1 error control and can also reduce statistical power in some circumstances. Many alternatives to INTs exist. Therefore, we contend that there is a lack of justification for performing parametric statistical procedures on INTs with the exceptions of simple designs with moderate to large sample sizes, which makes permutation testing computationally infeasible and where maximum likelihood testing is used. Rigorous research evaluating the utility of INTs seems warranted.},
	language = {eng},
	number = {5},
	journal = {Behavior Genetics},
	author = {Beasley, T. Mark and Erickson, Stephen and Allison, David B.},
	month = sep,
	year = {2009},
	pmid = {19526352},
	pmcid = {PMC2921808},
	keywords = {Alleles, Humans, Models, Theoretical, Chromosome Mapping, Computer Simulation, Genetic Diseases, Inborn, Genetic Techniques, Genetics, Homozygote, Likelihood Functions, Models, Genetic, Models, Statistical, Phenotype, Reproducibility of Results, Research Design},
	pages = {580--595},
	file = {Full Text:/Users/yurikoharigaya/Zotero/storage/NZ2LI2EB/Beasley et al. - 2009 - Rank-based inverse normal transformations are incr.pdf:application/pdf},
}

@article{benjamini_controlling_1995,
	title = {Controlling the {False} {Discovery} {Rate}: {A} {Practical} and {Powerful} {Approach} to {Multiple} {Testing}},
	volume = {57},
	issn = {00359246},
	shorttitle = {Controlling the {False} {Discovery} {Rate}},
	url = {https://onlinelibrary.wiley.com/doi/10.1111/j.2517-6161.1995.tb02031.x},
	doi = {10.1111/j.2517-6161.1995.tb02031.x},
	language = {en},
	number = {1},
	urldate = {2023-08-09},
	journal = {Journal of the Royal Statistical Society: Series B (Methodological)},
	author = {Benjamini, Yoav and Hochberg, Yosef},
	month = jan,
	year = {1995},
	pages = {289--300},
}

@article{davis_efficient_2016,
	title = {An {Efficient} {Multiple}-{Testing} {Adjustment} for {eQTL} {Studies} that {Accounts} for {Linkage} {Disequilibrium} between {Variants}},
	volume = {98},
	issn = {1537-6605},
	doi = {10.1016/j.ajhg.2015.11.021},
	abstract = {Methods for multiple-testing correction in local expression quantitative trait locus (cis-eQTL) studies are a trade-off between statistical power and computational efficiency. Bonferroni correction, though computationally trivial, is overly conservative and fails to account for linkage disequilibrium between variants. Permutation-based methods are more powerful, though computationally far more intensive. We present an alternative correction method called eigenMT, which runs over 500 times faster than permutations and has adjusted p values that closely approximate empirical ones. To achieve this speed while also maintaining the accuracy of permutation-based methods, we estimate the effective number of independent variants tested for association with a particular gene, termed Meff, by using the eigenvalue decomposition of the genotype correlation matrix. We employ a regularized estimator of the correlation matrix to ensure Meff is robust and yields adjusted p values that closely approximate p values from permutations. Finally, using a common genotype matrix, we show that eigenMT can be applied with even greater efficiency to studies across tissues or conditions. Our method provides a simpler, more efficient approach to multiple-testing correction than existing methods and fits within existing pipelines for eQTL discovery.},
	language = {eng},
	number = {1},
	journal = {American Journal of Human Genetics},
	author = {Davis, Joe R. and Fresard, Laure and Knowles, David A. and Pala, Mauro and Bustamante, Carlos D. and Battle, Alexis and Montgomery, Stephen B.},
	month = jan,
	year = {2016},
	pmid = {26749306},
	pmcid = {PMC4716687},
	keywords = {Humans, Quantitative Trait Loci, Linkage Disequilibrium},
	pages = {216--224},
	file = {Full Text:/Users/yurikoharigaya/Zotero/storage/56B4P76M/Davis et al. - 2016 - An Efficient Multiple-Testing Adjustment for eQTL .pdf:application/pdf},
}

@article{carpenter_stan_2017,
	title = {\textit{{Stan}} : {A} {Probabilistic} {Programming} {Language}},
	volume = {76},
	issn = {1548-7660},
	shorttitle = {\textit{{Stan}}},
	url = {http://www.jstatsoft.org/v76/i01/},
	doi = {10.18637/jss.v076.i01},
	language = {en},
	number = {1},
	urldate = {2023-08-09},
	journal = {Journal of Statistical Software},
	author = {Carpenter, Bob and Gelman, Andrew and Hoffman, Matthew D. and Lee, Daniel and Goodrich, Ben and Betancourt, Michael and Brubaker, Marcus and Guo, Jiqiang and Li, Peter and Riddell, Allen},
	year = {2017},
	file = {Full Text:/Users/yurikoharigaya/Zotero/storage/3AW9EBIU/Carpenter et al. - 2017 - Stan  A Probabilistic Programming Language.pdf:application/pdf},
}

@article{gronau_bridgesampling_2020,
	title = {\textbf{bridgesampling} : {An} \textit{{R}} {Package} for {Estimating} {Normalizing} {Constants}},
	volume = {92},
	issn = {1548-7660},
	shorttitle = {\textbf{bridgesampling}},
	url = {http://www.jstatsoft.org/v92/i10/},
	doi = {10.18637/jss.v092.i10},
	language = {en},
	number = {10},
	urldate = {2023-08-09},
	journal = {Journal of Statistical Software},
	author = {Gronau, Quentin F. and Singmann, Henrik and Wagenmakers, Eric-Jan},
	year = {2020},
	file = {Full Text:/Users/yurikoharigaya/Zotero/storage/Q7I7G66E/Gronau et al. - 2020 - bridgesampling  An R Package for Es.pdf:application/pdf},
}

@article{moore_linear_2019,
	title = {A linear mixed-model approach to study multivariate gene-environment interactions},
	volume = {51},
	issn = {1546-1718},
	doi = {10.1038/s41588-018-0271-0},
	abstract = {Different exposures, including diet, physical activity, or external conditions can contribute to genotype-environment interactions (G×E). Although high-dimensional environmental data are increasingly available and multiple exposures have been implicated with G×E at the same loci, multi-environment tests for G×E are not established. Here, we propose the structured linear mixed model (StructLMM), a computationally efficient method to identify and characterize loci that interact with one or more environments. After validating our model using simulations, we applied StructLMM to body mass index in the UK Biobank, where our model yields previously known and novel G×E signals. Finally, in an application to a large blood eQTL dataset, we demonstrate that StructLMM can be used to study interactions with hundreds of environmental variables.},
	language = {eng},
	number = {1},
	journal = {Nature Genetics},
	author = {Moore, Rachel and Casale, Francesco Paolo and Jan Bonder, Marc and Horta, Danilo and {BIOS Consortium} and Franke, Lude and Barroso, Inês and Stegle, Oliver},
	month = jan,
	year = {2019},
	pmid = {30478441},
	pmcid = {PMC6354905},
	keywords = {Humans, Quantitative Trait Loci, Computer Simulation, Models, Genetic, Algorithms, Environment, Gene-Environment Interaction, Genotype, Linear Models},
	pages = {180--186},
	file = {Accepted Version:/Users/yurikoharigaya/Zotero/storage/MXU63EMD/Moore et al. - 2019 - A linear mixed-model approach to study multivariat.pdf:application/pdf},
}

@book{ritz_nonlinear_2009,
	address = {New York, NY},
	title = {Nonlinear {Regression} with {R}},
	isbn = {978-0-387-09615-5 978-0-387-09616-2},
	url = {http://link.springer.com/10.1007/978-0-387-09616-2},
	language = {en},
	urldate = {2023-11-02},
	publisher = {Springer New York},
	editor = {Ritz, Christian and Streibig, Jens Carl},
	year = {2009},
	doi = {10.1007/978-0-387-09616-2},
	file = {Submitted Version:/Users/yurikoharigaya/Zotero/storage/34GP325G/Ritz and Streibig - 2009 - Nonlinear Regression with R.pdf:application/pdf},
}

@book{gill_bayesian_2015,
	address = {Boca Raton},
	edition = {Third edition},
	series = {Chapman \& {Hall}/{CRC} statistics in the social and behavioral sciences series},
	title = {Bayesian methods: a social and behavioral sciences approach},
	isbn = {978-1-4398-6248-3},
	shorttitle = {Bayesian methods},
	publisher = {CRC Press, Taylor \& Francis Group},
	author = {Gill, Jeff},
	year = {2015},
	keywords = {Bayesian statistical decision theory, Social sciences, Statistical methods},
}

@article{servin_imputation-based_2007,
	title = {Imputation-based analysis of association studies: candidate regions and quantitative traits},
	volume = {3},
	issn = {1553-7404},
	shorttitle = {Imputation-based analysis of association studies},
	doi = {10.1371/journal.pgen.0030114},
	abstract = {We introduce a new framework for the analysis of association studies, designed to allow untyped variants to be more effectively and directly tested for association with a phenotype. The idea is to combine knowledge on patterns of correlation among SNPs (e.g., from the International HapMap project or resequencing data in a candidate region of interest) with genotype data at tag SNPs collected on a phenotyped study sample, to estimate ("impute") unmeasured genotypes, and then assess association between the phenotype and these estimated genotypes. Compared with standard single-SNP tests, this approach results in increased power to detect association, even in cases in which the causal variant is typed, with the greatest gain occurring when multiple causal variants are present. It also provides more interpretable explanations for observed associations, including assessing, for each SNP, the strength of the evidence that it (rather than another correlated SNP) is causal. Although we focus on association studies with quantitative phenotype and a relatively restricted region (e.g., a candidate gene), the framework is applicable and computationally practical for whole genome association studies. Methods described here are implemented in a software package, Bim-Bam, available from the Stephens Lab website http://stephenslab.uchicago.edu/software.html.},
	language = {eng},
	number = {7},
	journal = {PLoS genetics},
	author = {Servin, Bertrand and Stephens, Matthew},
	month = jul,
	year = {2007},
	pmid = {17676998},
	pmcid = {PMC1934390},
	keywords = {Humans, Chromosome Mapping, Models, Statistical, Phenotype, Genotype, Bayes Theorem, Computational Biology, Data Interpretation, Statistical, Genomics, Models, Biological, Odds Ratio, Polymorphism, Genetic, Polymorphism, Single Nucleotide, Software},
	pages = {e114},
	file = {Full Text:/Users/yurikoharigaya/Zotero/storage/7AMBFX4D/Servin and Stephens - 2007 - Imputation-based analysis of association studies .pdf:application/pdf},
}

@article{flutre_statistical_2013,
	title = {A statistical framework for joint {eQTL} analysis in multiple tissues},
	volume = {9},
	issn = {1553-7404},
	doi = {10.1371/journal.pgen.1003486},
	abstract = {Mapping expression Quantitative Trait Loci (eQTLs) represents a powerful and widely adopted approach to identifying putative regulatory variants and linking them to specific genes. Up to now eQTL studies have been conducted in a relatively narrow range of tissues or cell types. However, understanding the biology of organismal phenotypes will involve understanding regulation in multiple tissues, and ongoing studies are collecting eQTL data in dozens of cell types. Here we present a statistical framework for powerfully detecting eQTLs in multiple tissues or cell types (or, more generally, multiple subgroups). The framework explicitly models the potential for each eQTL to be active in some tissues and inactive in others. By modeling the sharing of active eQTLs among tissues, this framework increases power to detect eQTLs that are present in more than one tissue compared with "tissue-by-tissue" analyses that examine each tissue separately. Conversely, by modeling the inactivity of eQTLs in some tissues, the framework allows the proportion of eQTLs shared across different tissues to be formally estimated as parameters of a model, addressing the difficulties of accounting for incomplete power when comparing overlaps of eQTLs identified by tissue-by-tissue analyses. Applying our framework to re-analyze data from transformed B cells, T cells, and fibroblasts, we find that it substantially increases power compared with tissue-by-tissue analysis, identifying 63\% more genes with eQTLs (at FDR = 0.05). Further, the results suggest that, in contrast to previous analyses of the same data, the majority of eQTLs detectable in these data are shared among all three tissues.},
	language = {eng},
	number = {5},
	journal = {PLoS genetics},
	author = {Flutre, Timothée and Wen, Xiaoquan and Pritchard, Jonathan and Stephens, Matthew},
	month = may,
	year = {2013},
	pmid = {23671422},
	pmcid = {PMC3649995},
	keywords = {Models, Theoretical, Quantitative Trait Loci, Chromosome Mapping, Gene Expression, Genome-Wide Association Study, Organ Specificity},
	pages = {e1003486},
	file = {Full Text:/Users/yurikoharigaya/Zotero/storage/NN7QSC4M/Flutre et al. - 2013 - A statistical framework for joint eQTL analysis in.pdf:application/pdf},
}

@article{fairfax_innate_2014,
	title = {Innate {Immune} {Activity} {Conditions} the {Effect} of {Regulatory} {Variants} upon {Monocyte} {Gene} {Expression}},
	volume = {343},
	issn = {0036-8075, 1095-9203},
	url = {https://www.science.org/doi/10.1126/science.1246949},
	doi = {10.1126/science.1246949},
	abstract = {Immune Variation
            
              It is difficult to determine the mechanistic consequences of context-dependent genetic variants, some of which may be related to disease (see the Perspective by
              
                Gregersen
              
              ). Two studies now report on the effects of stimulating immunological monocytes and dendritic cells with proteins that can elicit a response to bacterial or viral infection and assess the functional links between genetic variants and profiles of gene expression.
              
                M. N. Lee
                et al.
              
              (
              10.1126/science.1246980
              ) analyzed the expression of more than 400 genes, in dendritic cells from 30 healthy subjects, which revealed how expression quantitative trait loci (eQTLs) affect gene expression within the interferon-β and the Toll-like receptor 3 and 4 pathways.
              
                Fairfax
                et al.
              
              (
              10.1126/science.1246949
              ) performed a genome-wide analysis to show that many eQTLs affected monocyte gene expression in a stimulus- or time-specific manner.
            
          , 
            
              Analysis of the transcriptional responses during induced innate immune activity in primary human monocytes is explained.
              
                [Also see Perspective by
                Gregersen
                ]
              
            
          , 
            
              To systematically investigate the impact of immune stimulation upon regulatory variant activity, we exposed primary monocytes from 432 healthy Europeans to interferon-γ (IFN-γ) or differing durations of lipopolysaccharide and mapped expression quantitative trait loci (eQTLs). More than half of cis-eQTLs identified, involving hundreds of genes and associated pathways, are detected specifically in stimulated monocytes. Induced innate immune activity reveals multiple master regulatory trans-eQTLs including the major histocompatibility complex (MHC), coding variants altering enzyme and receptor function, an IFN-β cytokine network showing temporal specificity, and an interferon regulatory factor 2 (IRF2) transcription factor–modulated network. Induced eQTL are significantly enriched for genome-wide association study loci, identifying context-specific associations to putative causal genes including
              CARD9
              ,
              ATM
              , and
              IRF8
              . Thus, applying pathophysiologically relevant immune stimuli assists resolution of functional genetic variants.},
	language = {en},
	number = {6175},
	urldate = {2023-11-02},
	journal = {Science},
	author = {Fairfax, Benjamin P. and Humburg, Peter and Makino, Seiko and Naranbhai, Vivek and Wong, Daniel and Lau, Evelyn and Jostins, Luke and Plant, Katharine and Andrews, Robert and McGee, Chris and Knight, Julian C.},
	month = mar,
	year = {2014},
	pages = {1246949},
	file = {Accepted Version:/Users/yurikoharigaya/Zotero/storage/FMGNZ4XQ/Fairfax et al. - 2014 - Innate Immune Activity Conditions the Effect of Re.pdf:application/pdf},
}

@article{kim_characterizing_2014,
	title = {Characterizing the genetic basis of innate immune response in {TLR4}-activated human monocytes},
	volume = {5},
	issn = {2041-1723},
	url = {https://www.nature.com/articles/ncomms6236},
	doi = {10.1038/ncomms6236},
	abstract = {Abstract
            
              Toll-like receptors (TLRs) play a key role in innate immunity. Apart from their function in host defense, dysregulation in TLR signalling can confer risk to autoimmune diseases, septic shock or cancer. Here we report genetic variants and transcripts that are active only during TLR signalling and contribute to interindividual differences in immune response. Comparing unstimulated versus TLR4-stimulated monocytes reveals 1,471 expression quantitative trait loci (eQTLs) that are unique to TLR4 stimulation. Among these we find functional SNPs for the expression of
              NEU4
              ,
              CCL14
              ,
              CBX3
              and
              IRF5
              on TLR4 activation. Furthermore, we show that SNPs conferring risk to primary biliary cirrhosis (PBC), inflammatory bowel disease (IBD) and celiac disease are immune response eQTLs for
              PDGFB
              and
              IL18R1
              . Thus,
              PDGFB
              and
              IL18R1
              represent plausible candidates for studying the pathophysiology of these disorders in the context of TLR4 activation. In summary, this study presents novel insights into the genetic basis of the innate immune response and exemplifies the value of eQTL studies in the context of exogenous cell stimulation.},
	language = {en},
	number = {1},
	urldate = {2023-11-02},
	journal = {Nature Communications},
	author = {Kim, Sarah and Becker, Jessica and Bechheim, Matthias and Kaiser, Vera and Noursadeghi, Mahdad and Fricker, Nadine and Beier, Esther and Klaschik, Sven and Boor, Peter and Hess, Timo and Hofmann, Andrea and Holdenrieder, Stefan and Wendland, Jens R. and Fröhlich, Holger and Hartmann, Gunther and Nöthen, Markus M. and Müller-Myhsok, Bertram and Pütz, Benno and Hornung, Veit and Schumacher, Johannes},
	month = oct,
	year = {2014},
	pages = {5236},
	file = {Full Text:/Users/yurikoharigaya/Zotero/storage/B86MFGXQ/Kim et al. - 2014 - Characterizing the genetic basis of innate immune .pdf:application/pdf},
}

@article{lee_common_2014,
	title = {Common {Genetic} {Variants} {Modulate} {Pathogen}-{Sensing} {Responses} in {Human} {Dendritic} {Cells}},
	volume = {343},
	issn = {0036-8075, 1095-9203},
	url = {https://www.science.org/doi/10.1126/science.1246980},
	doi = {10.1126/science.1246980},
	abstract = {Immune Variation
            
              It is difficult to determine the mechanistic consequences of context-dependent genetic variants, some of which may be related to disease (see the Perspective by
              
                Gregersen
              
              ). Two studies now report on the effects of stimulating immunological monocytes and dendritic cells with proteins that can elicit a response to bacterial or viral infection and assess the functional links between genetic variants and profiles of gene expression.
              
                M. N. Lee
                et al.
              
              (
              10.1126/science.1246980
              ) analyzed the expression of more than 400 genes, in dendritic cells from 534 healthy subjects, which revealed how expression quantitative trait loci (eQTLs) affect gene expression within the interferon-β and the Toll-like receptor 3 and 4 pathways.
              
                Fairfax
                et al.
              
              (
              10.1126/science.1246949
              ) performed a genome-wide analysis to show that many eQTLs affected monocyte gene expression in a stimulus- or time-specific manner.
            
          , 
            
              Mapping of human host-pathogen gene-by-environment interactions identifies pathogen-specific loci.
              
                [Also see Perspective by
                Gregersen
                ]
              
            
          , 
            
              Little is known about how human genetic variation affects the responses to environmental stimuli in the context of complex diseases. Experimental and computational approaches were applied to determine the effects of genetic variation on the induction of pathogen-responsive genes in human dendritic cells. We identified 121 common genetic variants associated in cis with variation in expression responses to
              Escherichia coli
              lipopolysaccharide, influenza, or interferon-β (IFN-β). We localized and validated causal variants to binding sites of pathogen-activated STAT (signal transducer and activator of transcription) and IRF (IFN-regulatory factor) transcription factors. We also identified a common variant in
              IRF7
              that is associated in trans with type I IFN induction in response to influenza infection. Our results reveal common alleles that explain interindividual variation in pathogen sensing and provide functional annotation for genetic variants that alter susceptibility to inflammatory diseases.},
	language = {en},
	number = {6175},
	urldate = {2023-11-02},
	journal = {Science},
	author = {Lee, Mark N. and Ye, Chun and Villani, Alexandra-Chloé and Raj, Towfique and Li, Weibo and Eisenhaure, Thomas M. and Imboywa, Selina H. and Chipendo, Portia I. and Ran, F. Ann and Slowikowski, Kamil and Ward, Lucas D. and Raddassi, Khadir and McCabe, Cristin and Lee, Michelle H. and Frohlich, Irene Y. and Hafler, David A. and Kellis, Manolis and Raychaudhuri, Soumya and Zhang, Feng and Stranger, Barbara E. and Benoist, Christophe O. and De Jager, Philip L. and Regev, Aviv and Hacohen, Nir},
	month = mar,
	year = {2014},
	pages = {1246980},
	file = {Accepted Version:/Users/yurikoharigaya/Zotero/storage/UVRCCIU8/Lee et al. - 2014 - Common Genetic Variants Modulate Pathogen-Sensing .pdf:application/pdf},
}

@article{caliskan_host_2015,
	title = {Host {Genetic} {Variation} {Influences} {Gene} {Expression} {Response} to {Rhinovirus} {Infection}},
	volume = {11},
	issn = {1553-7404},
	url = {https://dx.plos.org/10.1371/journal.pgen.1005111},
	doi = {10.1371/journal.pgen.1005111},
	language = {en},
	number = {4},
	urldate = {2023-11-02},
	journal = {PLOS Genetics},
	author = {Çalışkan, Minal and Baker, Samuel W. and Gilad, Yoav and Ober, Carole},
	editor = {Gibson, Greg},
	month = apr,
	year = {2015},
	pages = {e1005111},
	file = {Full Text:/Users/yurikoharigaya/Zotero/storage/R9RARTDC/Çalışkan et al. - 2015 - Host Genetic Variation Influences Gene Expression .pdf:application/pdf},
}

@article{nedelec_genetic_2016,
	title = {Genetic {Ancestry} and {Natural} {Selection} {Drive} {Population} {Differences} in {Immune} {Responses} to {Pathogens}},
	volume = {167},
	issn = {00928674},
	url = {https://linkinghub.elsevier.com/retrieve/pii/S0092867416313071},
	doi = {10.1016/j.cell.2016.09.025},
	language = {en},
	number = {3},
	urldate = {2023-11-02},
	journal = {Cell},
	author = {Nédélec, Yohann and Sanz, Joaquín and Baharian, Golshid and Szpiech, Zachary A. and Pacis, Alain and Dumaine, Anne and Grenier, Jean-Christophe and Freiman, Andrew and Sams, Aaron J. and Hebert, Steven and Pagé Sabourin, Ariane and Luca, Francesca and Blekhman, Ran and Hernandez, Ryan D. and Pique-Regi, Roger and Tung, Jenny and Yotova, Vania and Barreiro, Luis B.},
	month = oct,
	year = {2016},
	pages = {657--669.e21},
	file = {Full Text:/Users/yurikoharigaya/Zotero/storage/YHPNBHCR/Nédélec et al. - 2016 - Genetic Ancestry and Natural Selection Drive Popul.pdf:application/pdf},
}

@techreport{wolter_cellular_2022,
	type = {preprint},
	title = {Cellular genome wide association study identifies common genetic variation influencing lithium induced neural progenitor proliferation},
	url = {http://biorxiv.org/lookup/doi/10.1101/2022.01.31.478307},
	abstract = {Abstract
          
            Lithium is used in the treatment of bipolar disorder (BD) and is known to increase neural progenitor cell (NPC) proliferation. Though the mechanism of lithium’s therapeutic effect is not understood, evidence suggests that genetic variation influences response to treatment. Here, we used a library of genetically diverse human NPCs to identify common genetic variants that modulate lithium induced proliferation. We identified a locus on chr3p21.1 associated with lithium induced proliferation that colocalizes with BD risk. One lithium responsive gene,
            GNL3
            , was detected within the locus. The allele associated with increased baseline and lithium-induced
            GNL3
            expression was also associated with increased lithium-induced NPC proliferation. Experimental manipulation of
            GNL3
            expression using CRISPRa/i in NPCs showed that
            GNL3
            was necessary for lithium’s full proliferative effects, and sufficient to induce proliferation without lithium treatment. In all, our data suggest that
            GNL3
            expression sensitizes NPCs for a stronger proliferative response to lithium.},
	language = {en},
	urldate = {2023-11-02},
	institution = {Genetics},
	author = {Wolter, Justin M. and Le, Brandon D. and Matoba, Nana and Lafferty, Michael J. and Aygün, Nil and Liang, Dan and Courtney, Kenan and Piven, Joseph and Zylka, Mark J. and Stein, Jason L.},
	month = feb,
	year = {2022},
	doi = {10.1101/2022.01.31.478307},
	file = {Submitted Version:/Users/yurikoharigaya/Zotero/storage/2FL8N5BS/Wolter et al. - 2022 - Cellular genome wide association study identifies .pdf:application/pdf},
}

@article{mosedale_identification_2019,
	title = {Identification of {Candidate} {Risk} {Factor} {Genes} for {Human} {Idelalisib} {Toxicity} {Using} a {Collaborative} {Cross} {Approach}},
	volume = {172},
	issn = {1096-0929},
	doi = {10.1093/toxsci/kfz199},
	abstract = {Idelalisib is a phosphatidylinositol 3-kinase inhibitor highly selective for the delta isoform that has shown good efficacy in treating chronic lymphocytic leukemia and follicular lymphoma. In clinical trials, however, idelalisib was associated with rare, but potentially serious liver and lung toxicities. In this study, we used the Collaborative Cross (CC) mouse population to identify genetic factors associated with the drug response that may inform risk management strategies for idelalisib in humans. Eight male mice (4 matched pairs) from 50 CC lines were treated once daily for 14 days by oral gavage with either vehicle or idelalisib at a dose selected to achieve clinically relevant peak plasma concentrations (150 mg/kg/day). The drug was well tolerated across all CC lines, and there were no observations of overt liver injury. Differences across CC lines were seen in drug concentration in plasma samples collected at the approximate Tmax on study Days 1, 7, and 14. There were also small but statistically significant treatment-induced alterations in plasma total bile acids and microRNA-122, and these may indicate early hepatocellular stress required for immune-mediated hepatotoxicity in humans. Idelalisib treatment further induced significant elevations in the total cell count of terminal bronchoalveolar lavage fluid, which may be analogous to pneumonitis observed in the clinic. Genetic mapping identified loci associated with interim plasma idelalisib concentration and the other 3 treatment-related endpoints. Thirteen priority candidate quantitative trait genes identified in CC mice may now guide interrogation of risk factors for adverse drug responses associated with idelalisib in humans.},
	language = {eng},
	number = {2},
	journal = {Toxicological Sciences: An Official Journal of the Society of Toxicology},
	author = {Mosedale, Merrie and Cai, Yanwei and Eaddy, John Scott and Corty, Robert W. and Nautiyal, Manisha and Watkins, Paul B. and Valdar, William},
	month = dec,
	year = {2019},
	pmid = {31501888},
	pmcid = {PMC6876541},
	keywords = {Quantitative Trait Loci, Chromosome Mapping, Polymorphism, Single Nucleotide, Animals, Antineoplastic Agents, Biomarkers, Bronchoalveolar Lavage Fluid, Chemical and Drug Induced Liver Injury, Collaborative Cross, Dose-Response Relationship, Drug, drug-induced liver injury, idelalisib, Liver Function Tests, Lung Injury, Mice, Inbred Strains, MicroRNAs, Oxidative Stress, Phosphatidylinositol 3-Kinase, precision medicine, Protein Kinase Inhibitors, Purines, Quinazolinones, Risk Factors, Species Specificity, toxicogenetics, Toxicogenetics},
	pages = {265--278},
	file = {Full Text:/Users/yurikoharigaya/Zotero/storage/4C49YADJ/Mosedale et al. - 2019 - Identification of Candidate Risk Factor Genes for .pdf:application/pdf},
}

@article{barreiro_deciphering_2012,
	title = {Deciphering the genetic architecture of variation in the immune response to \textit{{Mycobacterium} tuberculosis} infection},
	volume = {109},
	issn = {0027-8424, 1091-6490},
	url = {https://pnas.org/doi/full/10.1073/pnas.1115761109},
	doi = {10.1073/pnas.1115761109},
	abstract = {Tuberculosis (TB) is a major public health problem. One-third of the world's population is estimated to be infected with
              Mycobacterium tuberculosis
              (MTB), the etiological agent causing TB, and active disease kills nearly 2 million individuals worldwide every year. Several lines of evidence indicate that interindividual variation in susceptibility to TB has a heritable component, yet we still know little about the underlying genetic architecture. To address this, we performed a genome-wide mapping study of loci that are associated with functional variation in immune response to MTB. Specifically, we characterized transcript and protein expression levels and mapped expression quantitative trait loci (eQTL) in primary dendritic cells (DCs) from 65 individuals, before and after infection with MTB. We found 198 response eQTL, namely loci that were associated with variation in gene expression levels in either untreated or MTB-infected DCs, but not both. These response eQTL are associated with natural regulatory variation that likely affects (directly or indirectly) host interaction with MTB. Indeed, when we integrated our data with results from a genome-wide association study (GWAS) for pulmonary TB, we found that the response eQTL were more likely to be genetically associated with the disease. We thus identified a number of candidate loci, including the MAPK phosphatase
              DUSP14
              in particular, that are promising susceptibility genes to pulmonary TB.},
	language = {en},
	number = {4},
	urldate = {2023-11-02},
	journal = {Proceedings of the National Academy of Sciences},
	author = {Barreiro, Luis B. and Tailleux, Ludovic and Pai, Athma A. and Gicquel, Brigitte and Marioni, John C. and Gilad, Yoav},
	month = jan,
	year = {2012},
	pages = {1204--1209},
	file = {Full Text:/Users/yurikoharigaya/Zotero/storage/8PWCS3RS/Barreiro et al. - 2012 - Deciphering the genetic architecture of variation .pdf:application/pdf},
}

@article{li_inter-individual_2016,
	title = {Inter-individual variability and genetic influences on cytokine responses to bacteria and fungi},
	volume = {22},
	issn = {1078-8956, 1546-170X},
	url = {https://www.nature.com/articles/nm.4139},
	doi = {10.1038/nm.4139},
	language = {en},
	number = {8},
	urldate = {2023-11-02},
	journal = {Nature Medicine},
	author = {Li, Yang and Oosting, Marije and Deelen, Patrick and Ricaño-Ponce, Isis and Smeekens, Sanne and Jaeger, Martin and Matzaraki, Vasiliki and Swertz, Morris A and Xavier, Ramnik J and Franke, Lude and Wijmenga, Cisca and Joosten, Leo A B and Kumar, Vinod and Netea, Mihai G},
	month = aug,
	year = {2016},
	pages = {952--960},
	file = {Accepted Version:/Users/yurikoharigaya/Zotero/storage/CJHE7BLG/Li et al. - 2016 - Inter-individual variability and genetic influence.pdf:application/pdf},
}

@article{the_lifelines_cohort_study_genetic_2015,
	title = {Genetic studies of body mass index yield new insights for obesity biology},
	volume = {518},
	issn = {0028-0836, 1476-4687},
	url = {https://www.nature.com/articles/nature14177},
	doi = {10.1038/nature14177},
	language = {en},
	number = {7538},
	urldate = {2023-11-03},
	journal = {Nature},
	author = {{The LifeLines Cohort Study} and {The ADIPOGen Consortium} and {The AGEN-BMI Working Group} and {The CARDIOGRAMplusC4D Consortium} and {The CKDGen Consortium} and {The GLGC} and {The ICBP} and {The MAGIC Investigators} and {The MuTHER Consortium} and {The MIGen Consortium} and {The PAGE Consortium} and {The ReproGen Consortium} and {The GENIE Consortium} and {The International Endogene Consortium} and Locke, Adam E. and Kahali, Bratati and Berndt, Sonja I. and Justice, Anne E. and Pers, Tune H. and Day, Felix R. and Powell, Corey and Vedantam, Sailaja and Buchkovich, Martin L. and Yang, Jian and Croteau-Chonka, Damien C. and Esko, Tonu and Fall, Tove and Ferreira, Teresa and Gustafsson, Stefan and Kutalik, Zoltán and Luan, Jian’an and Mägi, Reedik and Randall, Joshua C. and Winkler, Thomas W. and Wood, Andrew R. and Workalemahu, Tsegaselassie and Faul, Jessica D. and Smith, Jennifer A. and Hua Zhao, Jing and Zhao, Wei and Chen, Jin and Fehrmann, Rudolf and Hedman, Åsa K. and Karjalainen, Juha and Schmidt, Ellen M. and Absher, Devin and Amin, Najaf and Anderson, Denise and Beekman, Marian and Bolton, Jennifer L. and Bragg-Gresham, Jennifer L. and Buyske, Steven and Demirkan, Ayse and Deng, Guohong and Ehret, Georg B. and Feenstra, Bjarke and Feitosa, Mary F. and Fischer, Krista and Goel, Anuj and Gong, Jian and Jackson, Anne U. and Kanoni, Stavroula and Kleber, Marcus E. and Kristiansson, Kati and Lim, Unhee and Lotay, Vaneet and Mangino, Massimo and Mateo Leach, Irene and Medina-Gomez, Carolina and Medland, Sarah E. and Nalls, Michael A. and Palmer, Cameron D. and Pasko, Dorota and Pechlivanis, Sonali and Peters, Marjolein J. and Prokopenko, Inga and Shungin, Dmitry and Stančáková, Alena and Strawbridge, Rona J. and Ju Sung, Yun and Tanaka, Toshiko and Teumer, Alexander and Trompet, Stella and Van Der Laan, Sander W. and Van Setten, Jessica and Van Vliet-Ostaptchouk, Jana V. and Wang, Zhaoming and Yengo, Loïc and Zhang, Weihua and Isaacs, Aaron and Albrecht, Eva and Ärnlöv, Johan and Arscott, Gillian M. and Attwood, Antony P. and Bandinelli, Stefania and Barrett, Amy and Bas, Isabelita N. and Bellis, Claire and Bennett, Amanda J. and Berne, Christian and Blagieva, Roza and Blüher, Matthias and Böhringer, Stefan and Bonnycastle, Lori L. and Böttcher, Yvonne and Boyd, Heather A. and Bruinenberg, Marcel and Caspersen, Ida H. and Ida Chen, Yii-Der and Clarke, Robert and Warwick Daw, E. and De Craen, Anton J. M. and Delgado, Graciela and Dimitriou, Maria and Doney, Alex S. F. and Eklund, Niina and Estrada, Karol and Eury, Elodie and Folkersen, Lasse and Fraser, Ross M. and Garcia, Melissa E. and Geller, Frank and Giedraitis, Vilmantas and Gigante, Bruna and Go, Alan S. and Golay, Alain and Goodall, Alison H. and Gordon, Scott D. and Gorski, Mathias and Grabe, Hans-Jörgen and Grallert, Harald and Grammer, Tanja B. and Gräßler, Jürgen and Grönberg, Henrik and Groves, Christopher J. and Gusto, Gaëlle and Haessler, Jeffrey and Hall, Per and Haller, Toomas and Hallmans, Goran and Hartman, Catharina A. and Hassinen, Maija and Hayward, Caroline and Heard-Costa, Nancy L. and Helmer, Quinta and Hengstenberg, Christian and Holmen, Oddgeir and Hottenga, Jouke-Jan and James, Alan L. and Jeff, Janina M. and Johansson, Åsa and Jolley, Jennifer and Juliusdottir, Thorhildur and Kinnunen, Leena and Koenig, Wolfgang and Koskenvuo, Markku and Kratzer, Wolfgang and Laitinen, Jaana and Lamina, Claudia and Leander, Karin and Lee, Nanette R. and Lichtner, Peter and Lind, Lars and Lindström, Jaana and Sin Lo, Ken and Lobbens, Stéphane and Lorbeer, Roberto and Lu, Yingchang and Mach, François and Magnusson, Patrik K. E. and Mahajan, Anubha and McArdle, Wendy L. and McLachlan, Stela and Menni, Cristina and Merger, Sigrun and Mihailov, Evelin and Milani, Lili and Moayyeri, Alireza and Monda, Keri L. and Morken, Mario A. and Mulas, Antonella and Müller, Gabriele and Müller-Nurasyid, Martina and Musk, Arthur W. and Nagaraja, Ramaiah and Nöthen, Markus M. and Nolte, Ilja M. and Pilz, Stefan and Rayner, Nigel W. and Renstrom, Frida and Rettig, Rainer and Ried, Janina S. and Ripke, Stephan and Robertson, Neil R. and Rose, Lynda M. and Sanna, Serena and Scharnagl, Hubert and Scholtens, Salome and Schumacher, Fredrick R. and Scott, William R. and Seufferlein, Thomas and Shi, Jianxin and Vernon Smith, Albert and Smolonska, Joanna and Stanton, Alice V. and Steinthorsdottir, Valgerdur and Stirrups, Kathleen and Stringham, Heather M. and Sundström, Johan and Swertz, Morris A. and Swift, Amy J. and Syvänen, Ann-Christine and Tan, Sian-Tsung and Tayo, Bamidele O. and Thorand, Barbara and Thorleifsson, Gudmar and Tyrer, Jonathan P. and Uh, Hae-Won and Vandenput, Liesbeth and Verhulst, Frank C. and Vermeulen, Sita H. and Verweij, Niek and Vonk, Judith M. and Waite, Lindsay L. and Warren, Helen R. and Waterworth, Dawn and Weedon, Michael N. and Wilkens, Lynne R. and Willenborg, Christina and Wilsgaard, Tom and Wojczynski, Mary K. and Wong, Andrew and Wright, Alan F. and Zhang, Qunyuan and Brennan, Eoin P. and Choi, Murim and Dastani, Zari and Drong, Alexander W. and Eriksson, Per and Franco-Cereceda, Anders and Gådin, Jesper R. and Gharavi, Ali G. and Goddard, Michael E. and Handsaker, Robert E. and Huang, Jinyan and Karpe, Fredrik and Kathiresan, Sekar and Keildson, Sarah and Kiryluk, Krzysztof and Kubo, Michiaki and Lee, Jong-Young and Liang, Liming and Lifton, Richard P. and Ma, Baoshan and McCarroll, Steven A. and McKnight, Amy J. and Min, Josine L. and Moffatt, Miriam F. and Montgomery, Grant W. and Murabito, Joanne M. and Nicholson, George and Nyholt, Dale R. and Okada, Yukinori and Perry, John R. B. and Dorajoo, Rajkumar and Reinmaa, Eva and Salem, Rany M. and Sandholm, Niina and Scott, Robert A. and Stolk, Lisette and Takahashi, Atsushi and Tanaka, Toshihiro and Van’T Hooft, Ferdinand M. and Vinkhuyzen, Anna A. E. and Westra, Harm-Jan and Zheng, Wei and Zondervan, Krina T. and Heath, Andrew C. and Arveiler, Dominique and Bakker, Stephan J. L. and Beilby, John and Bergman, Richard N. and Blangero, John and Bovet, Pascal and Campbell, Harry and Caulfield, Mark J. and Cesana, Giancarlo and Chakravarti, Aravinda and Chasman, Daniel I. and Chines, Peter S. and Collins, Francis S. and Crawford, Dana C. and Adrienne Cupples, L. and Cusi, Daniele and Danesh, John and De Faire, Ulf and Den Ruijter, Hester M. and Dominiczak, Anna F. and Erbel, Raimund and Erdmann, Jeanette and Eriksson, Johan G. and Farrall, Martin and Felix, Stephan B. and Ferrannini, Ele and Ferrières, Jean and Ford, Ian and Forouhi, Nita G. and Forrester, Terrence and Franco, Oscar H. and Gansevoort, Ron T. and Gejman, Pablo V. and Gieger, Christian and Gottesman, Omri and Gudnason, Vilmundur and Gyllensten, Ulf and Hall, Alistair S. and Harris, Tamara B. and Hattersley, Andrew T. and Hicks, Andrew A. and Hindorff, Lucia A. and Hingorani, Aroon D. and Hofman, Albert and Homuth, Georg and Kees Hovingh, G. and Humphries, Steve E. and Hunt, Steven C. and Hyppönen, Elina and Illig, Thomas and Jacobs, Kevin B. and Jarvelin, Marjo-Riitta and Jöckel, Karl-Heinz and Johansen, Berit and Jousilahti, Pekka and Wouter Jukema, J. and Jula, Antti M. and Kaprio, Jaakko and Kastelein, John J. P. and Keinanen-Kiukaanniemi, Sirkka M. and Kiemeney, Lambertus A. and Knekt, Paul and Kooner, Jaspal S. and Kooperberg, Charles and Kovacs, Peter and Kraja, Aldi T. and Kumari, Meena and Kuusisto, Johanna and Lakka, Timo A. and Langenberg, Claudia and Le Marchand, Loic and Lehtimäki, Terho and Lyssenko, Valeriya and Männistö, Satu and Marette, André and Matise, Tara C. and McKenzie, Colin A. and McKnight, Barbara and Moll, Frans L. and Morris, Andrew D. and Morris, Andrew P. and Murray, Jeffrey C. and Nelis, Mari and Ohlsson, Claes and Oldehinkel, Albertine J. and Ong, Ken K. and Madden, Pamela A. F. and Pasterkamp, Gerard and Peden, John F. and Peters, Annette and Postma, Dirkje S. and Pramstaller, Peter P. and Price, Jackie F. and Qi, Lu and Raitakari, Olli T. and Rankinen, Tuomo and Rao, D. C. and Rice, Treva K. and Ridker, Paul M. and Rioux, John D. and Ritchie, Marylyn D. and Rudan, Igor and Salomaa, Veikko and Samani, Nilesh J. and Saramies, Jouko and Sarzynski, Mark A. and Schunkert, Heribert and Schwarz, Peter E. H. and Sever, Peter and Shuldiner, Alan R. and Sinisalo, Juha and Stolk, Ronald P. and Strauch, Konstantin and Tönjes, Anke and Trégouët, David-Alexandre and Tremblay, Angelo and Tremoli, Elena and Virtamo, Jarmo and Vohl, Marie-Claude and Völker, Uwe and Waeber, Gérard and Willemsen, Gonneke and Witteman, Jacqueline C. and Carola Zillikens, M. and Adair, Linda S. and Amouyel, Philippe and Asselbergs, Folkert W. and Assimes, Themistocles L. and Bochud, Murielle and Boehm, Bernhard O. and Boerwinkle, Eric and Bornstein, Stefan R. and Bottinger, Erwin P. and Bouchard, Claude and Cauchi, Stéphane and Chambers, John C. and Chanock, Stephen J. and Cooper, Richard S. and De Bakker, Paul I. W. and Dedoussis, George and Ferrucci, Luigi and Franks, Paul W. and Froguel, Philippe and Groop, Leif C. and Haiman, Christopher A. and Hamsten, Anders and Hui, Jennie and Hunter, David J. and Hveem, Kristian and Kaplan, Robert C. and Kivimaki, Mika and Kuh, Diana and Laakso, Markku and Liu, Yongmei and Martin, Nicholas G. and März, Winfried and Melbye, Mads and Metspalu, Andres and Moebus, Susanne and Munroe, Patricia B. and Njølstad, Inger and Oostra, Ben A. and Palmer, Colin N. A. and Pedersen, Nancy L. and Perola, Markus and Pérusse, Louis and Peters, Ulrike and Power, Chris and Quertermous, Thomas and Rauramaa, Rainer and Rivadeneira, Fernando and Saaristo, Timo E. and Saleheen, Danish and Sattar, Naveed and Schadt, Eric E. and Schlessinger, David and Eline Slagboom, P. and Snieder, Harold and Spector, Tim D. and Thorsteinsdottir, Unnur and Stumvoll, Michael and Tuomilehto, Jaakko and Uitterlinden, André G. and Uusitupa, Matti and Van Der Harst, Pim and Walker, Mark and Wallaschofski, Henri and Wareham, Nicholas J. and Watkins, Hugh and Weir, David R. and Wichmann, H-Erich and Wilson, James F. and Zanen, Pieter and Borecki, Ingrid B. and Deloukas, Panos and Fox, Caroline S. and Heid, Iris M. and O’Connell, Jeffrey R. and Strachan, David P. and Stefansson, Kari and Van Duijn, Cornelia M. and Abecasis, Gonçalo R. and Franke, Lude and Frayling, Timothy M. and McCarthy, Mark I. and Visscher, Peter M. and Scherag, André and Willer, Cristen J. and Boehnke, Michael and Mohlke, Karen L. and Lindgren, Cecilia M. and Beckmann, Jacques S. and Barroso, Inês and North, Kari E. and Ingelsson, Erik and Hirschhorn, Joel N. and Loos, Ruth J. F. and Speliotes, Elizabeth K.},
	month = feb,
	year = {2015},
	pages = {197--206},
	file = {Accepted Version:/Users/yurikoharigaya/Zotero/storage/DZU97BP3/The LifeLines Cohort Study et al. - 2015 - Genetic studies of body mass index yield new insig.pdf:application/pdf},
}

@article{bycroft_uk_2018,
	title = {The {UK} {Biobank} resource with deep phenotyping and genomic data},
	volume = {562},
	issn = {0028-0836, 1476-4687},
	url = {https://www.nature.com/articles/s41586-018-0579-z},
	doi = {10.1038/s41586-018-0579-z},
	language = {en},
	number = {7726},
	urldate = {2023-11-03},
	journal = {Nature},
	author = {Bycroft, Clare and Freeman, Colin and Petkova, Desislava and Band, Gavin and Elliott, Lloyd T. and Sharp, Kevin and Motyer, Allan and Vukcevic, Damjan and Delaneau, Olivier and O’Connell, Jared and Cortes, Adrian and Welsh, Samantha and Young, Alan and Effingham, Mark and McVean, Gil and Leslie, Stephen and Allen, Naomi and Donnelly, Peter and Marchini, Jonathan},
	month = oct,
	year = {2018},
	pages = {203--209},
	file = {Full Text:/Users/yurikoharigaya/Zotero/storage/2WNNZYIS/Bycroft et al. - 2018 - The UK Biobank resource with deep phenotyping and .pdf:application/pdf},
}

@article{young_multiple_2016,
	title = {Multiple novel gene-by-environment interactions modify the effect of {FTO} variants on body mass index},
	volume = {7},
	issn = {2041-1723},
	url = {https://www.nature.com/articles/ncomms12724},
	doi = {10.1038/ncomms12724},
	abstract = {Abstract
            
              Genetic studies have shown that obesity risk is heritable and that, of the many common variants now associated with body mass index, those in an intron of the fat mass and obesity-associated (
              FTO
              ) gene have the largest effect. The size of the UK Biobank, and its joint measurement of genetic, anthropometric and lifestyle variables, offers an unprecedented opportunity to assess gene-by-environment interactions in a way that accounts for the dependence between different factors. We jointly examine the evidence for interactions between
              FTO
              (rs1421085) and various lifestyle and environmental factors. We report interactions between the
              FTO
              variant and each of: frequency of alcohol consumption (
              P
              =3.0 × 10
              −4
              ); deviations from mean sleep duration (
              P
              =8.0 × 10
              −4
              ); overall diet (
              P
              =5.0 × 10
              −6
              ), including added salt (
              P
              =1.2 × 10
              −3
              ); and physical activity (
              P
              =3.1 × 10
              −4
              ).},
	language = {en},
	number = {1},
	urldate = {2023-11-03},
	journal = {Nature Communications},
	author = {Young, Alexander I. and Wauthier, Fabian and Donnelly, Peter},
	month = sep,
	year = {2016},
	pages = {12724},
	file = {Full Text:/Users/yurikoharigaya/Zotero/storage/D2P3MIZK/Young et al. - 2016 - Multiple novel gene-by-environment interactions mo.pdf:application/pdf},
}

@book{hoff_first_2009,
	address = {New York, NY},
	series = {Springer {Texts} in {Statistics}},
	title = {A {First} {Course} in {Bayesian} {Statistical} {Methods}},
	isbn = {978-0-387-92299-7 978-0-387-92407-6},
	url = {http://link.springer.com/10.1007/978-0-387-92407-6},
	urldate = {2023-11-03},
	publisher = {Springer New York},
	author = {Hoff, Peter D.},
	year = {2009},
	doi = {10.1007/978-0-387-92407-6},
	file = {Full Text:/Users/yurikoharigaya/Zotero/storage/5HISC9YC/Hoff - 2009 - A First Course in Bayesian Statistical Methods.pdf:application/pdf},
}

@book{wakefield_bayesian_2013,
	address = {New York, NY},
	series = {Springer {Series} in {Statistics}},
	title = {Bayesian and {Frequentist} {Regression} {Methods}},
	isbn = {978-1-4419-0924-4 978-1-4419-0925-1},
	url = {https://link.springer.com/10.1007/978-1-4419-0925-1},
	language = {en},
	urldate = {2023-11-03},
	publisher = {Springer New York},
	author = {Wakefield, Jon},
	year = {2013},
	doi = {10.1007/978-1-4419-0925-1},
}

@book{gelman_bayesian_2014,
	address = {Boca Raton},
	edition = {Third edition},
	series = {Chapman \& {Hall}/{CRC} texts in statistical science},
	title = {Bayesian data analysis},
	isbn = {978-1-4398-4095-5},
	abstract = {"Preface This book is intended to have three roles and to serve three associated audiences: an introductory text on Bayesian inference starting from first principles, a graduate text on effective current approaches to Bayesian modeling and computation in statistics and related fields, and a handbook of Bayesian methods in applied statistics for general users of and researchers in applied statistics. Although introductory in its early sections, the book is definitely not elementary in the sense of a first text in statistics. The mathematics used in our book is basic probability and statistics, elementary calculus, and linear algebra. A review of probability notation is given in Chapter 1 along with a more detailed list of topics assumed to have been studied. The practical orientation of the book means that the reader's previous experience in probability, statistics, and linear algebra should ideally have included strong computational components. To write an introductory text alone would leave many readers with only a taste of the conceptual elements but no guidance for venturing into genuine practical applications, beyond those where Bayesian methods agree essentially with standard non-Bayesian analyses. On the other hand, we feel it would be a mistake to present the advanced methods without first introducing the basic concepts from our data-analytic perspective. Furthermore, due to the nature of applied statistics, a text on current Bayesian methodology would be incomplete without a variety of worked examples drawn from real applications. To avoid cluttering the main narrative, there are bibliographic notes at the end of each chapter and references at the end of the book"--},
	publisher = {CRC Press},
	author = {Gelman, Andrew},
	year = {2014},
	keywords = {Bayesian statistical decision theory, MATHEMATICS / Probability \& Statistics / General},
}

@incollection{fiske_data_2010,
	edition = {1},
	title = {Data {Analysis} in {Social} {Psychology}: {Recent} and {Recurring} {Issues}},
	isbn = {978-0-470-13747-5 978-0-470-56111-9},
	shorttitle = {Data {Analysis} in {Social} {Psychology}},
	url = {https://onlinelibrary.wiley.com/doi/10.1002/9780470561119.socpsy001004},
	language = {en},
	urldate = {2023-11-17},
	booktitle = {Handbook of {Social} {Psychology}},
	publisher = {Wiley},
	author = {Judd, Charles M. and Kenny, David A.},
	editor = {Fiske, Susan T. and Gilbert, Daniel T. and Lindzey, Gardner},
	month = jun,
	year = {2010},
	doi = {10.1002/9780470561119.socpsy001004},
}

@article{mackinnon_mediation_2007,
	title = {Mediation analysis},
	volume = {58},
	issn = {0066-4308},
	doi = {10.1146/annurev.psych.58.110405.085542},
	abstract = {Mediating variables are prominent in psychological theory and research. A mediating variable transmits the effect of an independent variable on a dependent variable. Differences between mediating variables and confounders, moderators, and covariates are outlined. Statistical methods to assess mediation and modern comprehensive approaches are described. Future directions for mediation analysis are discussed.},
	language = {eng},
	journal = {Annual Review of Psychology},
	author = {MacKinnon, David P. and Fairchild, Amanda J. and Fritz, Matthew S.},
	year = {2007},
	pmid = {16968208},
	pmcid = {PMC2819368},
	keywords = {Humans, Models, Statistical, Analysis of Variance, Biomedical Research, Causality, Effect Modifier, Epidemiologic, Longitudinal Studies, Mathematical Computing, Models, Psychological, Psychological Theory, Treatment Outcome},
	pages = {593--614},
	file = {Accepted Version:/Users/yurikoharigaya/Zotero/storage/GTZYEGX2/MacKinnon et al. - 2007 - Mediation analysis.pdf:application/pdf},
}

@article{judd_process_1981,
	title = {Process {Analysis}: {Estimating} {Mediation} in {Treatment} {Evaluations}},
	volume = {5},
	issn = {0193-841X, 1552-3926},
	shorttitle = {Process {Analysis}},
	url = {http://journals.sagepub.com/doi/10.1177/0193841X8100500502},
	doi = {10.1177/0193841X8100500502},
	abstract = {This article presents the rationale and procedures for conducting a process analysis in evaluation research. Such an analysis attempts to identify the process that mediates the effects of some treatment, by estimating the parameters of a causal chain between the treatment and some outcome variable. Two different procedures for estimating mediation are discussed. In addition we present procedures for examining whether a treatment exerts its effects, in part, by altering the mediating process that produces the outcome. Finally, the benefits of process analysis in evaluation research are underlined.},
	language = {en},
	number = {5},
	urldate = {2023-11-17},
	journal = {Evaluation Review},
	author = {Judd, Charles M. and Kenny, David A.},
	month = oct,
	year = {1981},
	pages = {602--619},
}

@article{pearl_interpretation_2014,
	title = {Interpretation and identification of causal mediation},
	volume = {19},
	issn = {1939-1463},
	doi = {10.1037/a0036434},
	abstract = {This article reviews the foundations of causal mediation analysis and offers a general and transparent account of the conditions necessary for the identification of natural direct and indirect effects, thus facilitating a more informed judgment of the plausibility of these conditions in specific applications. I show that the conditions usually cited in the literature are overly restrictive and can be relaxed substantially without compromising identification. In particular, I show that natural effects can be identified by methods that go beyond standard adjustment for confounders, applicable to observational studies in which treatment assignment remains confounded with the mediator or with the outcome. These identification conditions can be validated algorithmically from the diagrammatic description of one's model and are guaranteed to produce unbiased results whenever the description is correct. The identification conditions can be further relaxed in parametric models, possibly including interactions, and permit one to compare the relative importance of several pathways, mediated by interdependent variables.},
	language = {eng},
	number = {4},
	journal = {Psychological Methods},
	author = {Pearl, Judea},
	month = dec,
	year = {2014},
	pmid = {24885338},
	keywords = {Humans, Models, Statistical, Causality, Statistics as Topic},
	pages = {459--481},
}

@article{yuan_bayesian_2009,
	title = {Bayesian mediation analysis},
	volume = {14},
	issn = {1939-1463},
	doi = {10.1037/a0016972},
	abstract = {In this article, we propose Bayesian analysis of mediation effects. Compared with conventional frequentist mediation analysis, the Bayesian approach has several advantages. First, it allows researchers to incorporate prior information into the mediation analysis, thus potentially improving the efficiency of estimates. Second, under the Bayesian mediation analysis, inference is straightforward and exact, which makes it appealing for studies with small samples. Third, the Bayesian approach is conceptually simpler for multilevel mediation analysis. Simulation studies and analysis of 2 data sets are used to illustrate the proposed methods. (PsycINFO Database Record (c) 2009 APA, all rights reserved).},
	language = {eng},
	number = {4},
	journal = {Psychological Methods},
	author = {Yuan, Ying and MacKinnon, David P.},
	month = dec,
	year = {2009},
	pmid = {19968395},
	pmcid = {PMC2885293},
	keywords = {Humans, Bayes Theorem, Models, Psychological, Confidence Intervals, Psychology},
	pages = {301--322},
	file = {Accepted Version:/Users/yurikoharigaya/Zotero/storage/ZGBV982M/Yuan and MacKinnon - 2009 - Bayesian mediation analysis.pdf:application/pdf},
}

@article{oreper_reciprocal_2018,
	title = {Reciprocal {F1} {Hybrids} of {Two} {Inbred} {Mouse} {Strains} {Reveal} {Parent}-of-{Origin} and {Perinatal} {Diet} {Effects} on {Behavior} and {Expression}},
	volume = {8},
	issn = {2160-1836},
	doi = {10.1534/g3.118.200135},
	abstract = {Parent-of-origin effects (POE) in mammals typically arise from maternal effects or imprinting. In some instances, such POE have been associated with psychiatric disorders, as well as with changes in a handful of animal behaviors. However, POE on complex traits such as behavior remain largely uncharacterized. Moreover, although both behavior and epigenetic effects are known to be modified by perinatal environmental exposures such as nutrient deficiency, the architecture of such environment-by-POE is mostly unexplored. To study POE and environment-by-POE, we employ a relatively neglected but especially powerful experimental system for POE-detection: reciprocal F1 hybrids (RF1s). We exposed female NOD/ShiLtJ×C57Bl/6J and C57Bl/6J×NOD/ShiLtJ mice, perinatally, to one of four different diets, then after weaning recorded a set of behaviors that model psychiatric disease. Whole-brain microarray expression data revealed an imprinting-enriched set of 15 genes subject to POE. The most-significant expression POE, on the non-imprinted gene Carmil1 (a.k.a. Lrrc16a), was validated using qPCR in the same and in a new set of mice. Several behaviors, especially locomotor behaviors, also showed POE. Bayesian mediation analysis suggested Carmil1 expression suppresses behavioral POE, and that the imprinted gene Airn suppresses POE on Carmil1 expression. A suggestive diet-by-POE was observed on percent center time in the open field test, and a significant diet-by-POE was observed on one imprinted gene, Mir341, and on 16 non-imprinted genes. The relatively small, tractable set of POE and diet-by-POE detected on behavior and expression here motivates further studies examining such effects across RF1s on multiple genetic backgrounds.},
	language = {eng},
	number = {11},
	journal = {G3 (Bethesda, Md.)},
	author = {Oreper, Daniel and Schoenrock, Sarah A. and McMullan, Rachel and Ervin, Robin and Farrington, Joseph and Miller, Darla R. and de Villena, Fernando Pardo-Manuel and Valdar, William and Tarantino, Lisa M.},
	month = nov,
	year = {2018},
	pmid = {30171036},
	pmcid = {PMC6222572},
	keywords = {Animals, Bayesian mediation analysis, Behavior, Animal, Brain, C57BL/6J, Diet, Female, Genomic Imprinting, imprinting, Male, Mice, Inbred C57BL, Mice, Inbred NOD, NOD/ShiLtJ, nutritional deficiencies, Stress, Psychological, Tissue Array Analysis},
	pages = {3447--3468},
	file = {Full Text:/Users/yurikoharigaya/Zotero/storage/RV7GM33M/Oreper et al. - 2018 - Reciprocal F1 Hybrids of Two Inbred Mouse Strains .pdf:application/pdf},
}

@article{song_bayesian_2020,
	title = {Bayesian shrinkage estimation of high dimensional causal mediation effects in omics studies},
	volume = {76},
	issn = {1541-0420},
	doi = {10.1111/biom.13189},
	abstract = {Causal mediation analysis aims to examine the role of a mediator or a group of mediators that lie in the pathway between an exposure and an outcome. Recent biomedical studies often involve a large number of potential mediators based on high-throughput technologies. Most of the current analytic methods focus on settings with one or a moderate number of potential mediators. With the expanding growth of -omics data, joint analysis of molecular-level genomics data with epidemiological data through mediation analysis is becoming more common. However, such joint analysis requires methods that can simultaneously accommodate high-dimensional mediators and that are currently lacking. To address this problem, we develop a Bayesian inference method using continuous shrinkage priors to extend previous causal mediation analysis techniques to a high-dimensional setting. Simulations demonstrate that our method improves the power of global mediation analysis compared to simpler alternatives and has decent performance to identify true nonnull contributions to the mediation effects of the pathway. The Bayesian method also helps us to understand the structure of the composite null cases for inactive mediators in the pathway. We applied our method to Multi-Ethnic Study of Atherosclerosis and identified DNA methylation regions that may actively mediate the effect of socioeconomic status on cardiometabolic outcomes.},
	language = {eng},
	number = {3},
	journal = {Biometrics},
	author = {Song, Yanyi and Zhou, Xiang and Zhang, Min and Zhao, Wei and Liu, Yongmei and Kardia, Sharon L. R. and Roux, Ana V. Diez and Needham, Belinda L. and Smith, Jennifer A. and Mukherjee, Bhramar},
	month = sep,
	year = {2020},
	pmid = {31733066},
	pmcid = {PMC7228845},
	keywords = {Models, Statistical, Bayes Theorem, Causality, Bayesian sparse models, continuous shrinkage, DNA Methylation, epigenetics, high-dimensional mediators, Mediation Analysis},
	pages = {700--710},
	file = {Full Text:/Users/yurikoharigaya/Zotero/storage/DHCCK6VY/Song et al. - 2020 - Bayesian shrinkage estimation of high dimensional .pdf:application/pdf},
}

@article{nuijten_default_2015,
	title = {A default {Bayesian} hypothesis test for mediation},
	volume = {47},
	issn = {1554-3528},
	doi = {10.3758/s13428-014-0470-2},
	abstract = {In order to quantify the relationship between multiple variables, researchers often carry out a mediation analysis. In such an analysis, a mediator (e.g., knowledge of a healthy diet) transmits the effect from an independent variable (e.g., classroom instruction on a healthy diet) to a dependent variable (e.g., consumption of fruits and vegetables). Almost all mediation analyses in psychology use frequentist estimation and hypothesis-testing techniques. A recent exception is Yuan and MacKinnon (Psychological Methods, 14, 301-322, 2009), who outlined a Bayesian parameter estimation procedure for mediation analysis. Here we complete the Bayesian alternative to frequentist mediation analysis by specifying a default Bayesian hypothesis test based on the Jeffreys-Zellner-Siow approach. We further extend this default Bayesian test by allowing a comparison to directional or one-sided alternatives, using Markov chain Monte Carlo techniques implemented in JAGS. All Bayesian tests are implemented in the R package BayesMed (Nuijten, Wetzels, Matzke, Dolan, \& Wagenmakers, 2014).},
	language = {eng},
	number = {1},
	journal = {Behavior Research Methods},
	author = {Nuijten, Michèle B. and Wetzels, Ruud and Matzke, Dora and Dolan, Conor V. and Wagenmakers, Eric-Jan},
	month = mar,
	year = {2015},
	pmid = {24903686},
	keywords = {Humans, Research Design, Bayes Theorem, Markov Chains, Monte Carlo Method, Multivariate Analysis, Negotiating},
	pages = {85--97},
	file = {Full Text:/Users/yurikoharigaya/Zotero/storage/C7NX4JLF/Nuijten et al. - 2015 - A default Bayesian hypothesis test for mediation.pdf:application/pdf},
}

@article{pickar-oliver_next_2019,
	title = {The next generation of {CRISPR}–{Cas} technologies and applications},
	volume = {20},
	issn = {1471-0072, 1471-0080},
	url = {https://www.nature.com/articles/s41580-019-0131-5},
	doi = {10.1038/s41580-019-0131-5},
	language = {en},
	number = {8},
	urldate = {2023-11-17},
	journal = {Nature Reviews Molecular Cell Biology},
	author = {Pickar-Oliver, Adrian and Gersbach, Charles A.},
	month = aug,
	year = {2019},
	pages = {490--507},
	file = {Accepted Version:/Users/yurikoharigaya/Zotero/storage/H4X5TZPN/Pickar-Oliver and Gersbach - 2019 - The next generation of CRISPR–Cas technologies and.pdf:application/pdf},
}

@article{wallace_eliciting_2020,
	title = {Eliciting priors and relaxing the single causal variant assumption in colocalisation analyses},
	volume = {16},
	issn = {1553-7404},
	url = {https://dx.plos.org/10.1371/journal.pgen.1008720},
	doi = {10.1371/journal.pgen.1008720},
	language = {en},
	number = {4},
	urldate = {2023-11-17},
	journal = {PLOS Genetics},
	author = {Wallace, Chris},
	editor = {Epstein, Michael P.},
	month = apr,
	year = {2020},
	pages = {e1008720},
	file = {Full Text:/Users/yurikoharigaya/Zotero/storage/LDAMTN4M/Wallace - 2020 - Eliciting priors and relaxing the single causal va.pdf:application/pdf},
}

@article{jin_pu1_2011,
	title = {{PU}.1 and {C}/{EBP}(alpha) synergistically program distinct response to {NF}-{kappaB} activation through establishing monocyte specific enhancers},
	volume = {108},
	issn = {1091-6490},
	doi = {10.1073/pnas.1017214108},
	abstract = {Unraveling the complexity of transcriptional programs coded by different cell types has been one of the central goals of cell biology. By using genome-wide location analysis, we examined how two different cell types generate different responses to the NF-κB signaling pathway. We showed that, after TNF-α treatment, the NF-κB p65 subunit binds to distinct genome locations and subsequently induces different subsets of genes in human monocytic THP-1 cells versus HeLa cells. Interestingly, the differential p65 binding in two cell types correlates with preexisting cell type-specific enhancers before TNF-α stimulation, marked by histone modifications. We also found that two transcription factors, PU.1 and C/EBPα, appear to synergistically mediate enhancer creation and affect NF-κB target selection in THP-1 cells. In HeLa cells, coexpression of PU.1 and C/EBPα conferred TNF-α responsiveness to a subset of THP-1-specific NF-κB target genes. These results suggest that the diversity of transcriptional programs in mammalian cells arises, at least in part, from preexisting enhancers that are established by cell-specific transcription factors.},
	language = {eng},
	number = {13},
	journal = {Proceedings of the National Academy of Sciences of the United States of America},
	author = {Jin, Fulai and Li, Yan and Ren, Bing and Natarajan, Rama},
	month = mar,
	year = {2011},
	pmid = {21402921},
	pmcid = {PMC3069155},
	keywords = {Humans, Animals, CCAAT-Enhancer-Binding Proteins, Enhancer Elements, Genetic, Gene Expression Regulation, Genome, HeLa Cells, Histones, Monocytes, NF-kappa B, p300-CBP Transcription Factors, Proto-Oncogene Proteins, Signal Transduction, Trans-Activators, Transcription Factors, Tumor Necrosis Factor-alpha},
	pages = {5290--5295},
	file = {Full Text:/Users/yurikoharigaya/Zotero/storage/T9UNV2UV/Jin et al. - 2011 - PU.1 and CEBP(alpha) synergistically program dist.pdf:application/pdf},
}

@article{heinz_simple_2010,
	title = {Simple combinations of lineage-determining transcription factors prime cis-regulatory elements required for macrophage and {B} cell identities},
	volume = {38},
	issn = {1097-4164},
	doi = {10.1016/j.molcel.2010.05.004},
	abstract = {Genome-scale studies have revealed extensive, cell type-specific colocalization of transcription factors, but the mechanisms underlying this phenomenon remain poorly understood. Here, we demonstrate in macrophages and B cells that collaborative interactions of the common factor PU.1 with small sets of macrophage- or B cell lineage-determining transcription factors establish cell-specific binding sites that are associated with the majority of promoter-distal H3K4me1-marked genomic regions. PU.1 binding initiates nucleosome remodeling, followed by H3K4 monomethylation at large numbers of genomic regions associated with both broadly and specifically expressed genes. These locations serve as beacons for additional factors, exemplified by liver X receptors, which drive both cell-specific gene expression and signal-dependent responses. Together with analyses of transcription factor binding and H3K4me1 patterns in other cell types, these studies suggest that simple combinations of lineage-determining transcription factors can specify the genomic sites ultimately responsible for both cell identity and cell type-specific responses to diverse signaling inputs.},
	language = {eng},
	number = {4},
	journal = {Molecular Cell},
	author = {Heinz, Sven and Benner, Christopher and Spann, Nathanael and Bertolino, Eric and Lin, Yin C. and Laslo, Peter and Cheng, Jason X. and Murre, Cornelis and Singh, Harinder and Glass, Christopher K.},
	month = may,
	year = {2010},
	pmid = {20513432},
	pmcid = {PMC2898526},
	keywords = {Animals, Male, Mice, Inbred C57BL, Proto-Oncogene Proteins, Trans-Activators, Transcription Factors, B-Lymphocytes, Binding Sites, Cell Lineage, Macrophages, Mice, Oligonucleotide Array Sequence Analysis, Regulatory Elements, Transcriptional},
	pages = {576--589},
	file = {Full Text:/Users/yurikoharigaya/Zotero/storage/UP252DR4/Heinz et al. - 2010 - Simple combinations of lineage-determining transcr.pdf:application/pdf},
}

@article{heinz_effect_2013,
	title = {Effect of natural genetic variation on enhancer selection and function},
	volume = {503},
	issn = {0028-0836, 1476-4687},
	url = {https://www.nature.com/articles/nature12615},
	doi = {10.1038/nature12615},
	language = {en},
	number = {7477},
	urldate = {2023-11-17},
	journal = {Nature},
	author = {Heinz, S. and Romanoski, C. E. and Benner, C. and Allison, K. A. and Kaikkonen, M. U. and Orozco, L. D. and Glass, C. K.},
	month = nov,
	year = {2013},
	pages = {487--492},
	file = {Accepted Version:/Users/yurikoharigaya/Zotero/storage/BVGKXDF5/Heinz et al. - 2013 - Effect of natural genetic variation on enhancer se.pdf:application/pdf},
}

@article{wang_epigenetic_2015,
	title = {Epigenetic priming of enhancers predicts developmental competence of {hESC}-derived endodermal lineage intermediates},
	volume = {16},
	issn = {1875-9777},
	doi = {10.1016/j.stem.2015.02.013},
	abstract = {Embryonic development relies on the capacity of progenitor cells to appropriately respond to inductive cues, a cellular property known as developmental competence. Here, we report that epigenetic priming of enhancers signifies developmental competence during endodermal lineage diversification. Chromatin mapping during pancreatic and hepatic differentiation of human embryonic stem cells revealed the en masse acquisition of a poised chromatin state at enhancers specific to endoderm-derived cell lineages in gut tube intermediates. Experimentally, the acquisition of this poised enhancer state predicts the ability of endodermal intermediates to respond to inductive signals. Furthermore, these enhancers are first recognized by the pioneer transcription factors FOXA1 and FOXA2 when competence is acquired, while subsequent recruitment of lineage-inductive transcription factors, such as PDX1, leads to enhancer and target gene activation. Together, our results identify the acquisition of a poised chromatin state at enhancers as a mechanism by which progenitor cells acquire developmental competence.},
	language = {eng},
	number = {4},
	journal = {Cell Stem Cell},
	author = {Wang, Allen and Yue, Feng and Li, Yan and Xie, Ruiyu and Harper, Thomas and Patel, Nisha A. and Muth, Kayla and Palmer, Jeffrey and Qiu, Yunjiang and Wang, Jinzhao and Lam, Dieter K. and Raum, Jeffrey C. and Stoffers, Doris A. and Ren, Bing and Sander, Maike},
	month = apr,
	year = {2015},
	pmid = {25842977},
	pmcid = {PMC4478079},
	keywords = {Humans, High-Throughput Nucleotide Sequencing, Organ Specificity, Biomarkers, Enhancer Elements, Genetic, Histones, Trans-Activators, Cell Lineage, Cell Differentiation, Cell Line, Chromatin Assembly and Disassembly, Embryonic Stem Cells, Endoderm, Epigenesis, Genetic, Gastrula, Gene Expression Regulation, Developmental, Hepatocyte Nuclear Factor 3-alpha, Hepatocyte Nuclear Factor 3-beta, Homeodomain Proteins, Pancreas, RNA, Small Interfering},
	pages = {386--399},
	file = {Full Text:/Users/yurikoharigaya/Zotero/storage/ES9GCE4Z/Wang et al. - 2015 - Epigenetic priming of enhancers predicts developme.pdf:application/pdf},
}

@article{liu_large-scale_2022,
	title = {Large-{Scale} {Hypothesis} {Testing} for {Causal} {Mediation} {Effects} with {Applications} in {Genome}-wide {Epigenetic} {Studies}},
	volume = {117},
	issn = {0162-1459},
	doi = {10.1080/01621459.2021.1914634},
	abstract = {In genome-wide epigenetic studies, it is of great scientific interest to assess whether the effect of an exposure on a clinical outcome is mediated through DNA methylations. However, statistical inference for causal mediation effects is challenged by the fact that one needs to test a large number of composite null hypotheses across the whole epigenome. Two popular tests, the Wald-type Sobel's test and the joint significant test using the traditional null distribution are underpowered and thus can miss important scientific discoveries. In this paper, we show that the null distribution of Sobel's test is not the standard normal distribution and the null distribution of the joint significant test is not uniform under the composite null of no mediation effect, especially in finite samples and under the singular point null case that the exposure has no effect on the mediator and the mediator has no effect on the outcome. Our results explain why these two tests are underpowered, and more importantly motivate us to develop a more powerful Divide-Aggregate Composite-null Test (DACT) for the composite null hypothesis of no mediation effect by leveraging epigenome-wide data. We adopted Efron's empirical null framework for assessing statistical significance of the DACT test. We showed analytically that the proposed DACT method had improved power, and could well control type I error rate. Our extensive simulation studies showed that, in finite samples, the DACT method properly controlled the type I error rate and outperformed Sobel's test and the joint significance test for detecting mediation effects. We applied the DACT method to the US Department of Veterans Affairs Normative Aging Study, an ongoing prospective cohort study which included men who were aged 21 to 80 years at entry. We identified multiple DNA methylation CpG sites that might mediate the effect of smoking on lung function with effect sizes ranging from -0.18 to -0.79 and false discovery rate controlled at level 0.05, including the CpG sites in the genes AHRR and F2RL3. Our sensitivity analysis found small residual correlations (less than 0.01) of the error terms between the outcome and mediator regressions, suggesting that our results are robust to unmeasured confounding factors.},
	language = {eng},
	number = {537},
	journal = {Journal of the American Statistical Association},
	author = {Liu, Zhonghua and Shen, Jincheng and Barfield, Richard and Schwartz, Joel and Baccarelli, Andrea A. and Lin, Xihong},
	year = {2022},
	pmid = {35989709},
	pmcid = {PMC9385159},
	keywords = {Causal mediation analysis, Composite null, Divide-aggregate composite-null test, Genome-wide epigenetic studies, Hypothesis testing, Indirect effects, Mediation effects, Proportions of true nulls},
	pages = {67--81},
	file = {Accepted Version:/Users/yurikoharigaya/Zotero/storage/3XBE7XVJ/Liu et al. - 2022 - Large-Scale Hypothesis Testing for Causal Mediatio.pdf:application/pdf},
}

@book{muller_regression_2003,
	address = {Hoboken, N.J},
	title = {Regression and {ANOVA}: an integrated approach using {SAS} software},
	isbn = {978-0-471-46943-8},
	shorttitle = {Regression and {ANOVA}},
	publisher = {J. Wiley},
	author = {Muller, Keith E. and Fetterman, Bethel A.},
	year = {2003},
	keywords = {Analysis of variance, Data processing, Regression analysis, SAS (Computer file)},
}

@article{akaike_new_1974,
	title = {A new look at the statistical model identification},
	volume = {19},
	copyright = {https://ieeexplore.ieee.org/Xplorehelp/downloads/license-information/IEEE.html},
	issn = {0018-9286},
	url = {http://ieeexplore.ieee.org/document/1100705/},
	doi = {10.1109/TAC.1974.1100705},
	language = {en},
	number = {6},
	urldate = {2024-04-23},
	journal = {IEEE Transactions on Automatic Control},
	author = {Akaike, H.},
	month = dec,
	year = {1974},
	pages = {716--723},
}

@article{schwarz_estimating_1978,
	title = {Estimating the {Dimension} of a {Model}},
	volume = {6},
	issn = {0090-5364},
	url = {https://projecteuclid.org/journals/annals-of-statistics/volume-6/issue-2/Estimating-the-Dimension-of-a-Model/10.1214/aos/1176344136.full},
	doi = {10.1214/aos/1176344136},
	number = {2},
	urldate = {2024-04-23},
	journal = {The Annals of Statistics},
	author = {Schwarz, Gideon},
	month = mar,
	year = {1978},
	file = {Full Text:/Users/yurikoharigaya/Zotero/storage/9YDF2QMV/Schwarz - 1978 - Estimating the Dimension of a Model.pdf:application/pdf},
}

@article{veyrieras_high-resolution_2008,
	title = {High-resolution mapping of expression-{QTLs} yields insight into human gene regulation},
	volume = {4},
	issn = {1553-7404},
	doi = {10.1371/journal.pgen.1000214},
	abstract = {Recent studies of the HapMap lymphoblastoid cell lines have identified large numbers of quantitative trait loci for gene expression (eQTLs). Reanalyzing these data using a novel Bayesian hierarchical model, we were able to create a surprisingly high-resolution map of the typical locations of sites that affect mRNA levels in cis. Strikingly, we found a strong enrichment of eQTLs in the 250 bp just upstream of the transcription end site (TES), in addition to an enrichment around the transcription start site (TSS). Most eQTLs lie either within genes or close to genes; for example, we estimate that only 5\% of eQTLs lie more than 20 kb upstream of the TSS. After controlling for position effects, SNPs in exons are approximately 2-fold more likely than SNPs in introns to be eQTLs. Our results suggest an important role for mRNA stability in determining steady-state mRNA levels, and highlight the potential of eQTL mapping as a high-resolution tool for studying the determinants of gene regulation.},
	language = {eng},
	number = {10},
	journal = {PLoS genetics},
	author = {Veyrieras, Jean-Baptiste and Kudaravalli, Sridhar and Kim, Su Yeon and Dermitzakis, Emmanouil T. and Gilad, Yoav and Stephens, Matthew and Pritchard, Jonathan K.},
	month = oct,
	year = {2008},
	pmid = {18846210},
	pmcid = {PMC2556086},
	keywords = {Humans, Quantitative Trait Loci, Chromosome Mapping, Models, Genetic, Bayes Theorem, Polymorphism, Single Nucleotide, Gene Expression Regulation, Databases, Genetic, Genetic Complementation Test, Genome, Human, Introns, RNA Stability, RNA, Messenger, Terminator Regions, Genetic, Transcription Initiation Site},
	pages = {e1000214},
	file = {Full Text:/Users/yurikoharigaya/Zotero/storage/SYFYUVRY/Veyrieras et al. - 2008 - High-resolution mapping of expression-QTLs yields .pdf:application/pdf},
}

@article{chenn_regulation_2002,
	title = {Regulation of cerebral cortical size by control of cell cycle exit in neural precursors},
	volume = {297},
	issn = {1095-9203},
	doi = {10.1126/science.1074192},
	abstract = {Transgenic mice expressing a stabilized beta-catenin in neural precursors develop enlarged brains with increased cerebral cortical surface area and folds resembling sulci and gyri of higher mammals. Brains from transgenic animals have enlarged lateral ventricles lined with neuroepithelial precursor cells, reflecting an expansion of the precursor population. Compared with wild-type precursors, a greater proportion of transgenic precursors reenter the cell cycle after mitosis. These results show that beta-catenin can function in the decision of precursors to proliferate or differentiate during mammalian neuronal development and suggest that beta-catenin can regulate cerebral cortical size by controlling the generation of neural precursor cells.},
	language = {eng},
	number = {5580},
	journal = {Science (New York, N.Y.)},
	author = {Chenn, Anjen and Walsh, Christopher A.},
	month = jul,
	year = {2002},
	pmid = {12130776},
	keywords = {Animals, Brain, Signal Transduction, Trans-Activators, Mice, Cell Differentiation, Antibodies, Monoclonal, Apoptosis, Basic Helix-Loop-Helix Transcription Factors, beta Catenin, Cell Adhesion Molecules, Neuronal, Cell Count, Cell Cycle, Cell Division, Cerebral Cortex, Cytoskeletal Proteins, DNA-Binding Proteins, Epithelium, Extracellular Matrix Proteins, Fungal Proteins, In Situ Hybridization, Ki-67 Antigen, Mice, Transgenic, Mitosis, Nerve Tissue Proteins, Neurons, Recombinant Fusion Proteins, Reelin Protein, Repressor Proteins, Saccharomyces cerevisiae Proteins, Serine Endopeptidases, Stem Cells, T-Box Domain Proteins},
	pages = {365--369},
}

@article{harrison-uy_wnt_2012,
	title = {Wnt {Signaling} and {Forebrain} {Development}},
	volume = {4},
	issn = {1943-0264},
	url = {http://cshperspectives.cshlp.org/lookup/doi/10.1101/cshperspect.a008094},
	doi = {10.1101/cshperspect.a008094},
	language = {en},
	number = {7},
	urldate = {2024-05-28},
	journal = {Cold Spring Harbor Perspectives in Biology},
	author = {Harrison-Uy, S. J. and Pleasure, S. J.},
	month = jul,
	year = {2012},
	pages = {a008094--a008094},
	file = {Full Text:/Users/yurikoharigaya/Zotero/storage/ALXKWQ7A/Harrison-Uy and Pleasure - 2012 - Wnt Signaling and Forebrain Development.pdf:application/pdf},
}

@article{evgrafov_gene_2020,
	title = {Gene {Expression} in {Patient}-{Derived} {Neural} {Progenitors} {Implicates} {WNT5A} {Signaling} in the {Etiology} of {Schizophrenia}},
	volume = {88},
	copyright = {https://www.elsevier.com/tdm/userlicense/1.0/},
	issn = {00063223},
	url = {https://linkinghub.elsevier.com/retrieve/pii/S000632232030010X},
	doi = {10.1016/j.biopsych.2020.01.005},
	language = {en},
	number = {3},
	urldate = {2024-05-28},
	journal = {Biological Psychiatry},
	author = {Evgrafov, Oleg V. and Armoskus, Chris and Wrobel, Bozena B. and Spitsyna, Valeria N. and Souaiaia, Tade and Herstein, Jennifer S. and Walker, Christopher P. and Nguyen, Joseph D. and Camarena, Adrian and Weitz, Jonathan R. and Kim, Jae Mun “Hugo” and Lopez Duarte, Edder and Wang, Kai and Simpson, George M. and Sobell, Janet L. and Medeiros, Helena and Pato, Michele T. and Pato, Carlos N. and Knowles, James A.},
	month = aug,
	year = {2020},
	pages = {236--247},
	file = {Submitted Version:/Users/yurikoharigaya/Zotero/storage/SYKA6L53/Evgrafov et al. - 2020 - Gene Expression in Patient-Derived Neural Progenit.pdf:application/pdf},
}

@article{topol_altered_2015,
	title = {Altered {WNT} {Signaling} in {Human} {Induced} {Pluripotent} {Stem} {Cell} {Neural} {Progenitor} {Cells} {Derived} from {Four} {Schizophrenia} {Patients}},
	volume = {78},
	copyright = {https://www.elsevier.com/tdm/userlicense/1.0/},
	issn = {00063223},
	url = {https://linkinghub.elsevier.com/retrieve/pii/S0006322315000670},
	doi = {10.1016/j.biopsych.2014.12.028},
	language = {en},
	number = {6},
	urldate = {2024-05-28},
	journal = {Biological Psychiatry},
	author = {Topol, Aaron and Zhu, Shijia and Tran, Ngoc and Simone, Anthony and Fang, Gang and Brennand, Kristen J.},
	month = sep,
	year = {2015},
	pages = {e29--e34},
	file = {Accepted Version:/Users/yurikoharigaya/Zotero/storage/84Y4524H/Topol et al. - 2015 - Altered WNT Signaling in Human Induced Pluripotent.pdf:application/pdf},
}

@article{marchetto_altered_2017,
	title = {Altered proliferation and networks in neural cells derived from idiopathic autistic individuals},
	volume = {22},
	issn = {1359-4184, 1476-5578},
	url = {https://www.nature.com/articles/mp201695},
	doi = {10.1038/mp.2016.95},
	language = {en},
	number = {6},
	urldate = {2024-05-28},
	journal = {Molecular Psychiatry},
	author = {Marchetto, M C and Belinson, H and Tian, Y and Freitas, B C and Fu, C and Vadodaria, K C and Beltrao-Braga, P C and Trujillo, C A and Mendes, A P D and Padmanabhan, K and Nunez, Y and Ou, J and Ghosh, H and Wright, R and Brennand, K J and Pierce, K and Eichenfield, L and Pramparo, T and Eyler, L T and Barnes, C C and Courchesne, E and Geschwind, D H and Gage, F H and Wynshaw-Boris, A and Muotri, A R},
	month = jun,
	year = {2017},
	pages = {820--835},
	file = {Accepted Version:/Users/yurikoharigaya/Zotero/storage/4XMKVQTY/Marchetto et al. - 2017 - Altered proliferation and networks in neural cells.pdf:application/pdf},
}

@article{caracci_wnt-catenin-dependent_2021,
	title = {Wnt/β-{Catenin}-{Dependent} {Transcription} in {Autism} {Spectrum} {Disorders}},
	volume = {14},
	issn = {1662-5099},
	url = {https://www.frontiersin.org/articles/10.3389/fnmol.2021.764756/full},
	doi = {10.3389/fnmol.2021.764756},
	abstract = {Autism spectrum disorders (ASD) is a heterogeneous group of neurodevelopmental disorders characterized by synaptic dysfunction and defects in dendritic spine morphology. In the past decade, an extensive list of genes associated with ASD has been identified by genome-wide sequencing initiatives. Several of these genes functionally converge in the regulation of the Wnt/β-catenin signaling pathway, a conserved cascade essential for stem cell pluripotency and cell fate decisions during development. Here, we review current information regarding the transcriptional program of Wnt/β-catenin signaling in ASD. First, we discuss that Wnt/β-catenin gain and loss of function studies recapitulate brain developmental abnormalities associated with ASD. Second, transcriptomic approaches using patient-derived induced pluripotent stem cells (iPSC) cells, featuring mutations in high confidence ASD genes, reveal a significant dysregulation in the expression of Wnt signaling components. Finally, we focus on the activity of chromatin-remodeling proteins and transcription factors considered high confidence ASD genes, including CHD8, ARID1B, ADNP, and TBR1, that regulate Wnt/β-catenin-dependent transcriptional activity in multiple cell types, including pyramidal neurons, interneurons and oligodendrocytes, cells which are becoming increasingly relevant in the study of ASD. We conclude that the level of Wnt/β-catenin signaling activation could explain the high phenotypical heterogeneity of ASD and be instrumental in the development of new diagnostics tools and therapies.},
	urldate = {2024-05-28},
	journal = {Frontiers in Molecular Neuroscience},
	author = {Caracci, Mario O. and Avila, Miguel E. and Espinoza-Cavieres, Francisca A. and López, Héctor R. and Ugarte, Giorgia D. and De Ferrari, Giancarlo V.},
	month = nov,
	year = {2021},
	pages = {764756},
	file = {Full Text:/Users/yurikoharigaya/Zotero/storage/EFW92W45/Caracci et al. - 2021 - Wntβ-Catenin-Dependent Transcription in Autism Sp.pdf:application/pdf},
}

@article{gogarten_genetic_2019,
	title = {Genetic association testing using the {GENESIS} {R}/{Bioconductor} package},
	volume = {35},
	copyright = {https://academic.oup.com/journals/pages/open\_access/funder\_policies/chorus/standard\_publication\_model},
	issn = {1367-4803, 1367-4811},
	url = {https://academic.oup.com/bioinformatics/article/35/24/5346/5536872},
	doi = {10.1093/bioinformatics/btz567},
	abstract = {Abstract
            
              Summary
              The Genomic Data Storage (GDS) format provides efficient storage and retrieval of genotypes measured by microarrays and sequencing. We developed GENESIS to perform various single- and aggregate-variant association tests using genotype data stored in GDS format. GENESIS implements highly flexible mixed models, allowing for different link functions, multiple variance components and phenotypic heteroskedasticity. GENESIS integrates cohesively with other R/Bioconductor packages to build a complete genomic analysis workflow entirely within the R environment.
            
            
              Availability and implementation
              https://bioconductor.org/packages/GENESIS; vignettes included.
            
            
              Supplementary information
              Supplementary data are available at Bioinformatics online.},
	language = {en},
	number = {24},
	urldate = {2024-05-28},
	journal = {Bioinformatics},
	author = {Gogarten, Stephanie M and Sofer, Tamar and Chen, Han and Yu, Chaoyu and Brody, Jennifer A and Thornton, Timothy A and Rice, Kenneth M and Conomos, Matthew P},
	editor = {Valencia, Alfonso},
	month = dec,
	year = {2019},
	pages = {5346--5348},
	file = {Full Text:/Users/yurikoharigaya/Zotero/storage/DH36CVCG/Gogarten et al. - 2019 - Genetic association testing using the GENESIS RBi.pdf:application/pdf},
}

@article{nelder_reformulation_1977,
	title = {A {Reformulation} of {Linear} {Models}},
	volume = {140},
	issn = {00359238},
	url = {https://www.jstor.org/stable/10.2307/2344517?origin=crossref},
	doi = {10.2307/2344517},
	number = {1},
	urldate = {2024-05-28},
	journal = {Journal of the Royal Statistical Society. Series A (General)},
	author = {Nelder, J. A.},
	year = {1977},
	pages = {48},
}

@article{heathcote_limits_2023,
	title = {The {Limits} of {Marginality}},
	volume = {6},
	issn = {2522-0861, 2522-087X},
	url = {https://link.springer.com/10.1007/s42113-021-00120-3},
	doi = {10.1007/s42113-021-00120-3},
	abstract = {Abstract
            The “marginality principle” for linear regression models states that when a higher order term is included, its constituent terms must also be included. The target article relies on this principle for the fixed-effects part of linear mixed models of ANOVA designs and considers the implication that if extended to combined fixed-and-random-effects models, model selection tests specific to some fixed-effects ANOVA terms are not possible. We review the basis for this principle for fixed-effects models and delineate its limits. We then consider its extension to combined fixed-and-random-effects models. We conclude that we have been unable to find in the literature, including the target article, and have ourselves been unable to construct any satisfactory argument against the use of incomplete ANOVA models. The only basis we could find requires one to assume that it is not possible to test point-null hypotheses, something we disagree with, and which we believe is incompatible with the Bayesian model-selection methods that are the basis of the target article.},
	language = {en},
	number = {1},
	urldate = {2024-05-28},
	journal = {Computational Brain \& Behavior},
	author = {Heathcote, Andrew and Matzke, Dora},
	month = mar,
	year = {2023},
	pages = {28--34},
	file = {Full Text:/Users/yurikoharigaya/Zotero/storage/ZWJEZL8R/Heathcote and Matzke - 2023 - The Limits of Marginality.pdf:application/pdf},
}

@article{morris_marginality_2023,
	title = {The marginality principle revisited: {Should} "higher-order" terms always be accompanied by "lower-order" terms in regression analyses?},
	volume = {65},
	issn = {1521-4036},
	shorttitle = {The marginality principle revisited},
	doi = {10.1002/bimj.202300069},
	abstract = {The marginality principle guides analysts to avoid omitting lower-order terms from models in which higher-order terms are included as covariates. Lower-order terms are viewed as "marginal" to higher-order terms. We consider how this principle applies to three cases: regression models that may include the ratio of two measured variables; polynomial transformations of a measured variable; and factorial arrangements of defined interventions. For each case, we show that which terms or transformations are considered to be lower-order, and therefore marginal, depends on the scale of measurement, which is frequently arbitrary. Understanding the implications of this point leads to an intuitive understanding of the curse of dimensionality. We conclude that the marginality principle may be useful to analysts in some specific cases but caution against invoking it as a context-free recipe.},
	language = {eng},
	number = {8},
	journal = {Biometrical Journal. Biometrische Zeitschrift},
	author = {Morris, Tim P. and van Smeden, Maarten and Pham, Tra My},
	month = dec,
	year = {2023},
	pmid = {37775940},
	pmcid = {PMC7615369},
	keywords = {Algorithms, curse of dimensionality, interactions, marginality principle, polynomials, principle of marginality, ratios, Regression Analysis},
	pages = {e2300069},
	file = {Full Text:/Users/yurikoharigaya/Zotero/storage/88SBPHRF/Morris et al. - 2023 - The marginality principle revisited Should highe.pdf:application/pdf},
}

@article{ferrari_bayesian_2021,
	title = {Bayesian {Factor} {Analysis} for {Inference} on {Interactions}},
	volume = {116},
	issn = {0162-1459},
	doi = {10.1080/01621459.2020.1745813},
	abstract = {This article is motivated by the problem of inference on interactions among chemical exposures impacting human health outcomes. Chemicals often co-occur in the environment or in synthetic mixtures and as a result exposure levels can be highly correlated. We propose a latent factor joint model, which includes shared factors in both the predictor and response components while assuming conditional independence. By including a quadratic regression in the latent variables in the response component, we induce flexible dimension reduction in characterizing main effects and interactions. We propose a Bayesian approach to inference under this Factor analysis for INteractions (FIN) framework. Through appropriate modifications of the factor modeling structure, FIN can accommodate higher order interactions. We evaluate the performance using a simulation study and data from the National Health and Nutrition Examination Survey (NHANES). Code is available on GitHub.},
	language = {eng},
	number = {535},
	journal = {Journal of the American Statistical Association},
	author = {Ferrari, Federico and Dunson, David B.},
	year = {2021},
	pmid = {34898761},
	pmcid = {PMC8654343},
	keywords = {Bayesian Modeling, Chemical Mixtures, Correlated Exposures, Quadratic regression, Statistical Interactions},
	pages = {1521--1532},
	file = {Accepted Version:/Users/yurikoharigaya/Zotero/storage/AGFSKRS2/Ferrari and Dunson - 2021 - Bayesian Factor Analysis for Inference on Interact.pdf:application/pdf},
}
