## Supplementary figures and images for "Probabilistic classification of gene-by-treatment interactions on molecular count phenotypes"

### bms.png

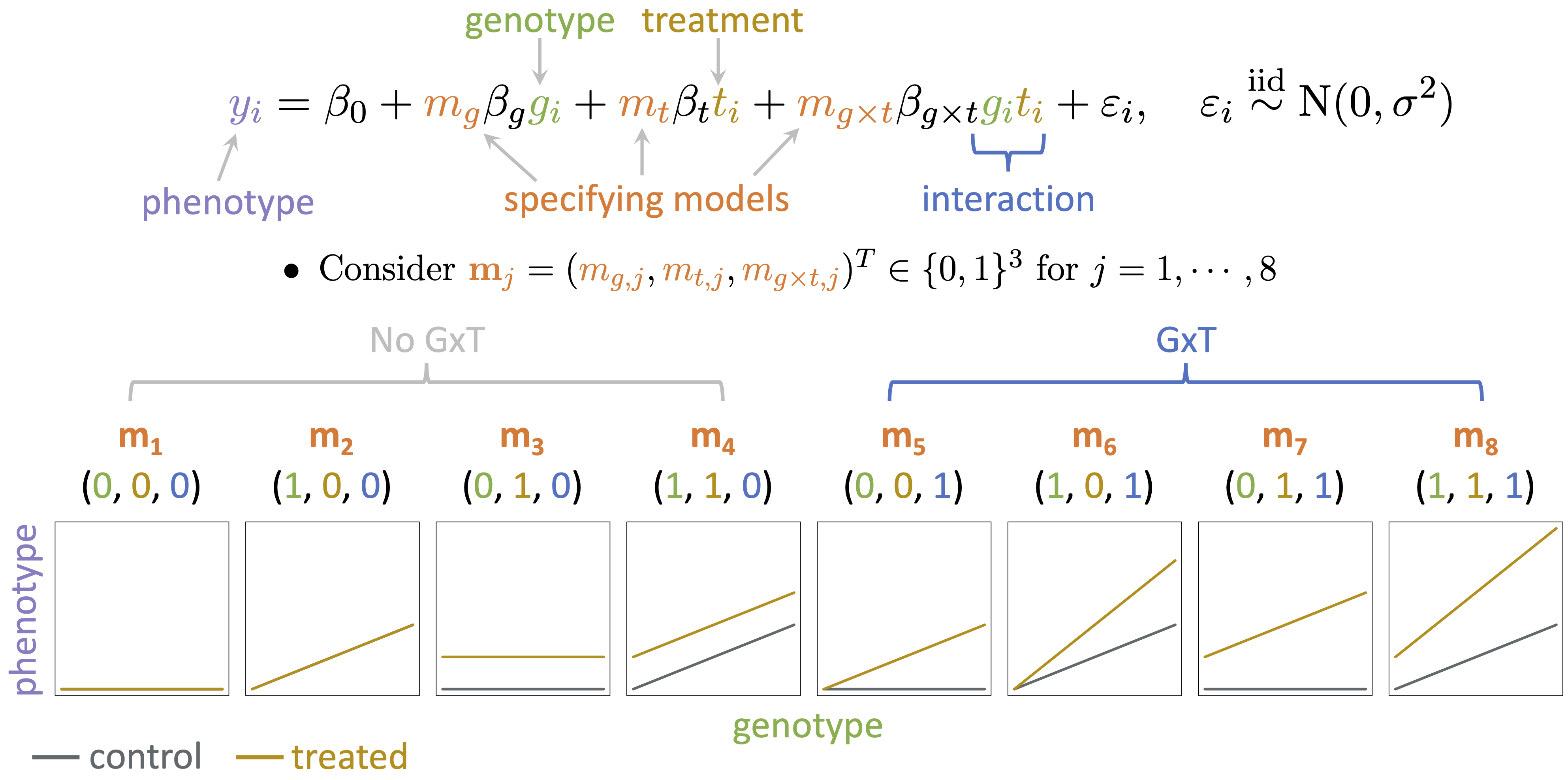

### unnamed-chunk-17-1.png

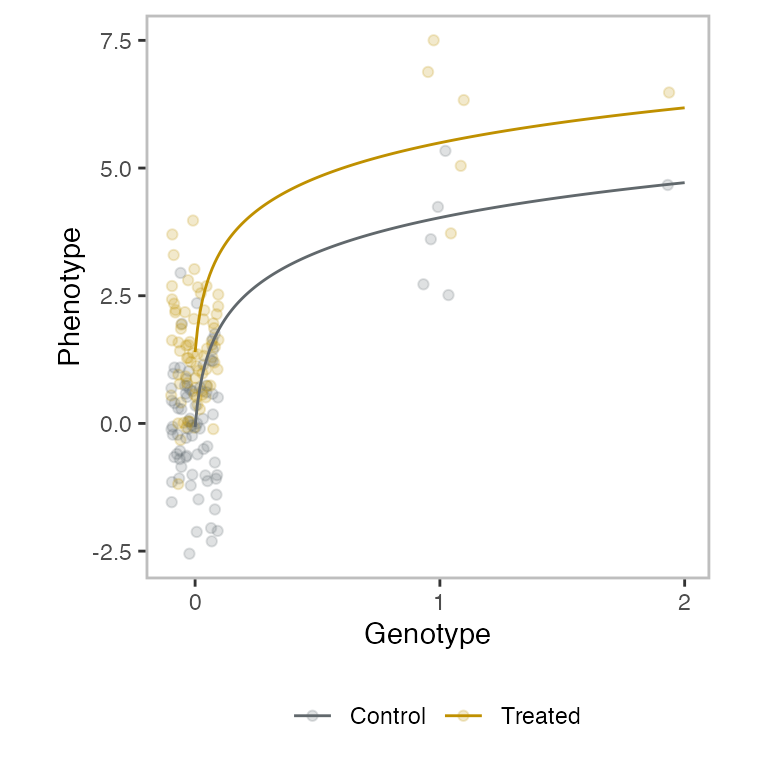

### unnamed-chunk-19-1.png

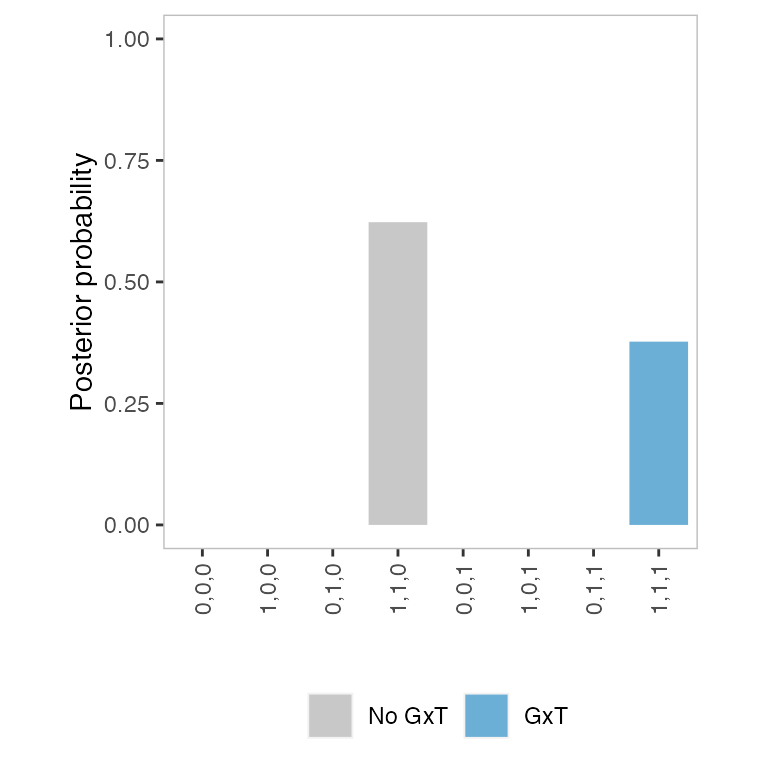

### unnamed-chunk-20-1.png

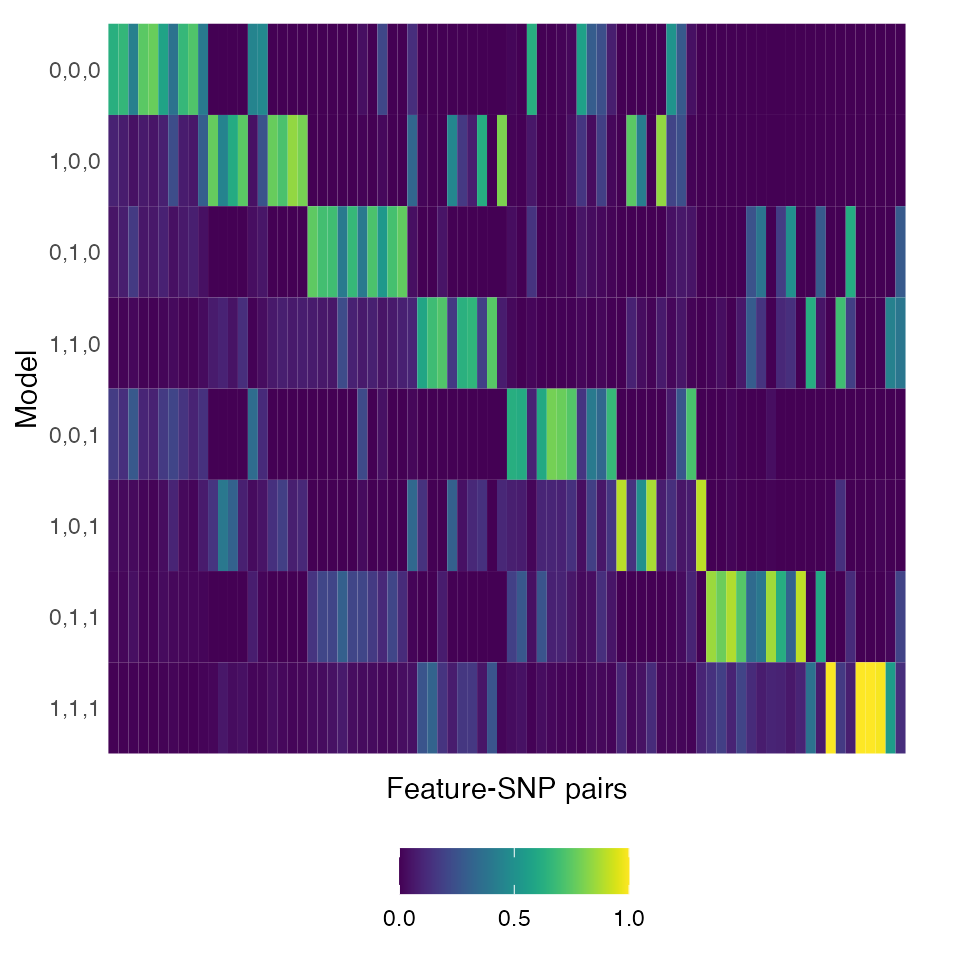
