## Supplemental File 2 for "Probabilistic classification of gene-by-treatment interactions on molecular count phenotypes": README.md

# classifygxt-paper

## About 

This repository contains code for analyses described in the following article on the ClassifyGxT method.

Yuriko Harigaya,
Nana Matoba,
Brandon D. Le,
Jordan M. Valone,
Jason L. Stein,
Michael I. Love\*,
William Valdar\*.
"Probabilistic classification of gene-by-treatment interactions on molecular count phenotypes."
(\* These authors contributed equally to this work.)

The ClassifyGxT software is available from <https://github.com/yharigaya/classifygxt>.

## Directories

* [sim](https://github.com/yharigaya/classifygxt-paper/tree/main/sim) - code for simulation analyses 
* [wntrna](https://github.com/yharigaya/classifygxt-paper/tree/main/wntrna) - code for analyzing response eQTL data in human neural progenitor cells (hNPCs) treated with CHIR, an activator of the Wnt pathway
* [wntatac](https://github.com/yharigaya/classifygxt-paper/tree/main/wntatac) - code for analyzing response caQTL data in hNPCs treated with CHIR
* [vis](https://github.com/yharigaya/classifygxt-paper/tree/main/vis) - code for visualization

See also `README.md` within each directory.
