## Supplemental File 2 for "Probabilistic classification of gene-by-treatment interactions on molecular count phenotypes": config.yaml

```
dir: "../derived_data/sim/"
fig_dir: "../figures/sim/"
lb: 25 # lower bound
ub: 250 # upper bound
step: 25 # step
lb_comp: 50
ub_comp: 250
step_comp: 50
size: 20 # batch size for mcmc
num_hp: 800 # for sensitivity analysis
num_post: 8000 # for post-processing
num_comp: 80 # for comparing two computatonal methods
seed: 1
num_core: 4 # number of cores
```
