## Supplemental File 2 for "Probabilistic classification of gene-by-treatment interactions on molecular count phenotypes": sbatch.yaml

```
---
__default__:
  time: "0-01:00"
  threads: 1
  mem: 2G
  queue: "general"
  error: "../logs/sim/%x_%j_%N.err"
  output: "../logs/sim/%x_%j_%N.log"
make_data:
  time: "0-00:30"
  threads: 1
  mem: 4G
  queue: "general"
optimize_hp:
  time: "0-04:00"
  threads: 1
  mem: 2G
  queue: "general"
map_lap_hp:
  time: "0-04:00"
  threads: 1
  mem: 2G
  queue: "general"
mcmc_bs_hp:
  time: "0-06:00"
  threads: 4
  mem: 2G
  queue: "general"
post:
  time: "0-06:00"
  threads: 4
  mem: 2G
  queue: "general"
get_co:
  time: "0-04:00"
  threads: 1
  mem: 4G
  queue: "general"
mcmc_bs_for_comp:
  time: "0-06:00"
  threads: 4
  mem: 2G
  queue: "general"
```
