## Supplemental File 2 for "Probabilistic classification of gene-by-treatment interactions on molecular count phenotypes": do_mcmc_bs.R

```
suppressPackageStartupMessages(library(tidyverse))
suppressPackageStartupMessages(library(classifygxt))

args <- commandArgs(TRUE)
file.data <- as.character(args[1])
file.tu.lambda <- as.character(args[2])
file.fit <- as.character(args[3])
seed <- as.integer(args[4])
n.cores <- as.integer(args[5])
method <- as.character(args[6])
ranef <- as.character(args[7])
hp <- as.character(args[8])
rstan <- as.character(args[9]) # summary or sample
start <- as.integer(args[10])
size <- as.integer(args[11])
dir.out <- as.character(args[12])

# read in the data
data.list <- file.data %>% readRDS
ln.p.y <- file.fit %>% readRDS

# specify model prior
p.m <- rep(1/8, 8)

# get phi values
opt <- ln.p.y[which.max(ln.p.y$ln.p.y), 1:3] %>%
    as.numeric

if (hp == "opt")
    phi <- opt
if (hp == "rest") {
    phi <- opt * 0.5
} else if (hp == "perm") {
    phi <- opt * 2
}

computation <- "mcmc.bs"

if (ranef == "ranef") {
    ranef <- TRUE
    tu.lambda <- file.tu.lambda %>% readRDS
} else if (ranef == "noranef") {
    ranef <- FALSE
    tu.lambda <- NULL
}    

if (method == "nl") {
    fn <- "nonlinear"
    rint <- FALSE
} else if (method == "lm") {
    fn <- "linear"
    rint <- FALSE
} else if (method == "rint") {    
    fn <- "linear"
    rint <- TRUE
}

if (rstan == "summary") {
    summary <- TRUE
} else if (rstan == "sample") {
    summary <- FALSE
}    

end <- start + size - 1
end <- min(end, length(data.list))

for (i in start:end) {

    file.out <- file.path(dir.out,
        paste0("res", str_pad(i, 4, pad="0"), ".rds"))
    if (file.exists(file.out)) {
        warning(paste("The file", file.out, "already exists."))
        next
    }

    data <- data.list[[i]]

    res <- do_bms(
        data=data, p.m=p.m, phi=phi,
        fn=fn, rint=rint, method=computation,
        tu.lambda=tu.lambda,
        summary=summary,
        n.cores=n.cores)
    
    saveRDS(res, file=file.out)
}
    
paste0(
    "complete",
    str_pad(start, 4, pad="0"), "_", size, ".txt") %>%
    file.path(dir.out, .) %>%
    file.create
```
