## Supplemental File 2 for "Probabilistic classification of gene-by-treatment interactions on molecular count phenotypes": do_mcmc_bs_for_comp.R

```
suppressPackageStartupMessages(library(tidyverse))
suppressPackageStartupMessages(library(reqtlpkg))

args <- commandArgs(TRUE)
data.file <- as.character(args[1])
tu.lambda.file <- as.character(args[2])
seed <- as.integer(args[3])
n.cores <- as.integer(args[4])
phi1 <- as.numeric(args[5])
phi2 <- as.numeric(args[6])
phi3 <- as.numeric(args[7])
method <- as.character(args[8])
ranef <- as.character(args[9])
rstan <- as.character(args[10])
start <- as.integer(args[11])
size <- as.integer(args[12])
out.dir <- as.character(args[13])
# out.file <- as.character(args[8])

# read in the data
data.list <- data.file %>% readRDS

# specify model prior
p.m <- rep(1/8, 8)

# get phi values
phi <- c(phi1, phi2, phi3)

computation <- "mcmc.bs"

if (ranef == "ranef") {
    ranef <- TRUE
    tu.lambda <- tu.lambda.file %>% readRDS
} else if (ranef == "noranef") {
    ranef <- FALSE
    tu.lambda <- NULL
}

if (method == "nl") {
    fn <- "nonlinear"
    rint <- FALSE
} else if (method == "lm") {
    fn <- "linear"
    rint <- FALSE
} else if (method == "rint") {
    fn <- "linear"
    rint <- TRUE
}

if (rstan == "summary") {
    summary <- TRUE
} else if (rstan == "sample") {
    summary <- FALSE
}

end <- start + size - 1
end <- min(end, length(data.list))


for (i in start:end) {

    out.file <- file.path(out.dir,
        paste0("res", str_pad(i, 4, pad="0"), ".rds"))
    if (file.exists(out.file)) {
        warning(paste("The file", out.file, "already exists."))
        next
    }

    data <- data.list[[i]]

    res <- do_bms(
        data=data, p.m=p.m, phi=phi,
        fn=fn, rint=rint, method=computation,
        tu.lambda=tu.lambda,
        summary=summary,
        n.cores=n.cores)

    saveRDS(res, file=out.file)

}

paste0(
    "complete",
    str_pad(start, 4, pad="0"), "_", size, ".txt") %>%
    file.path(out.dir, .) %>%
    file.create
```
