## Supplemental File 2 for "Probabilistic classification of gene-by-treatment interactions on molecular count phenotypes": get_calib.R

```
suppressPackageStartupMessages(library(tidyverse))
suppressPackageStartupMessages(library(classifygxt))

# define functions
get_vec <- function(d.pp, type) {

    m.pp <- d.pp %>%
    dplyr::select(-correct) %>%
    as.matrix
    
    if (type %in% c("nogxt", "interaction", "induced", "altered")) {
    
        if (type == "nogxt") {
            v.pattern <- c(1, 1, 1, 1, 0, 0, 0, 0)
        } else if (type == "interaction") {
            v.pattern <- c(0, 0, 0, 0, 1, 1, 1, 1)
        } else if (type == "induced") {
            v.pattern <- c(0, 0, 0, 0, 1, 0, 1, 0)
        } else if (type == "altered") {
            v.pattern <- c(0, 0, 0, 0, 0, 1, 0, 1)
        }
    } 

    v.truth <- (d.pp$correct %in% which(v.pattern == 1)) %>%
        as.numeric    
    v.pp <- rowSums(m.pp[, v.pattern == 1])

    output <- list(v.truth=v.truth, v.pp=v.pp)
    output    
}

# compute bs according to wiki
# compute_bs1 <- function(v.pp, v.truth) {
#     sum((v.pp - v.truth)^2) / length(v.truth)
# }    

# compute bs according to parmigiani
compute_bs2 <- function(v.pp, v.truth, v.bin=NULL) {
    if (is.null(v.bin)) {
        v.bin <- seq(0, 1, 0.1) # breaks
    }    
    # v.pi <- seq(0.05, 1, 0.1)
    n.bin <- length(v.bin) - 1 # number of bins

    v.bs.bin <- rep(0, n.bin)
    v.x <- rep(0, n.bin)
    v.n <- rep(0, n.bin)

    for (i in seq_len(n.bin)) {
        lb <- v.bin[i]
        ub <- v.bin[(i + 1)]
        pi.i <- mean(c(lb, ub)) # pi.i <- v.pi[i]
        sel <- v.pp >= lb & v.pp < ub
        if (all(!sel)) next
        v.pp.i <- v.pp[sel]
        v.truth.i <- v.truth[sel]
        v.n[i] <- length(v.pp.i)
        nu.i <- length(v.pp.i) / length(v.pp)
        x.i <- mean(v.truth.i == 1)
        v.x[i] <- x.i
        v.bs.bin[i] <-
            nu.i * (x.i * (pi.i - 1)^2 + (1 - x.i) * pi.i^2)
    }

    output <- list(
        v.pp=v.pp,
        v.truth=v.truth,
        v.bin=v.bin,
        v.x=v.x,
        v.n=v.n,
        v.bs.bin=v.bs.bin)

    output
}

args <- commandArgs(TRUE)
in.file <- as.character(args[1])
type <- as.character(args[2])
out.file <- as.character(args[3])

pp <- in.file %>% readRDS
pp <- na.omit(pp)

model.name <- get_model_names()
if (class(pp$correct) == "character") {
    index <- pp$correct %>%
        sapply(function(x) which(x == model.name))
    pp <- pp %>%
        mutate(correct=index)
}

vec.list <- pp %>% get_vec(type=type)
pp.vec <- vec.list$v.pp
truth.vec <- vec.list$v.truth

calib.list <- pp.vec %>%
    compute_bs2(v.truth=truth.vec)

saveRDS(calib.list, file=out.file)
```
