## Supplemental File 2 for "Probabilistic classification of gene-by-treatment interactions on molecular count phenotypes": get_calib_co.R

```
suppressPackageStartupMessages(library(tidyverse))
suppressPackageStartupMessages(library(classifygxt))

# compute bs according to wiki
# compute_bs1 <- function(v.pp, v.truth) {
#     sum((v.pp - v.truth)^2) / length(v.truth)
# }    

# compute bs according to parmigiani
compute_bs2 <- function(v.pp, v.truth, v.bin=NULL) {
    if (is.null(v.bin)) {
        v.bin <- seq(0, 1, 0.1) # breaks
    }    
    # v.pi <- seq(0.05, 1, 0.1)
    n.bin <- length(v.bin) - 1 # number of bins

    v.bs.bin <- rep(0, n.bin)
    v.x <- rep(0, n.bin)
    v.n <- rep(0, n.bin)

    for (i in seq_len(n.bin)) {
        lb <- v.bin[i]
        ub <- v.bin[(i + 1)]
        pi.i <- mean(c(lb, ub)) # pi.i <- v.pi[i]
        sel <- v.pp >= lb & v.pp < ub
        if (all(!sel)) next
        v.pp.i <- v.pp[sel]
        v.truth.i <- v.truth[sel]
        v.n[i] <- length(v.pp.i)
        nu.i <- length(v.pp.i) / length(v.pp)
        x.i <- mean(v.truth.i == 1)
        v.x[i] <- x.i
        v.bs.bin[i] <-
            nu.i * (x.i * (pi.i - 1)^2 + (1 - x.i) * pi.i^2)
    }

    output <- list(
        v.pp=v.pp,
        v.truth=v.truth,
        v.bin=v.bin,
        v.x=v.x,
        v.n=v.n,
        v.bs.bin=v.bs.bin)

    output
}

args <- commandArgs(TRUE)
in.file <- args[1]
true.file <- args[2]
out.file <- args[3]

p.co.vec <- in.file %>%
    readRDS %>%
    rowSums

true.vec <- true.file %>%
    readRDS %>%
    pluck("co")

calib.list <- p.co.vec %>%
    compute_bs2(v.truth=true.vec)

saveRDS(calib.list, file=out.file)
```
