## Supplemental File 2 for "Probabilistic classification of gene-by-treatment interactions on molecular count phenotypes": get_co.R

```
suppressPackageStartupMessages(library(tidyverse))
suppressPackageStartupMessages(library(classifygxt))

args <- commandArgs(TRUE)
in.dir <- as.character(args[1])
out.file <- as.character(args[2])

file.vec <- list.files(
    in.dir, pattern="^res", full.names=TRUE) %>%
    grep("rds$", ., value=TRUE)

# get the joint probability (default for get_co)
p.co <- file.vec %>%
    map(\(x) x %>% readRDS %>% get_co) %>%
    bind_rows %>%
    as.matrix

saveRDS(p.co, file=out.file)
```
