## Supplemental File 2 for "Probabilistic classification of gene-by-treatment interactions on molecular count phenotypes": get_lnpy_for_map_lap.R

```
suppressPackageStartupMessages(library(tidyverse))
suppressPackageStartupMessages(library(classifygxt))

args <- commandArgs(TRUE)
in.file <- as.character(args[1])
out.file <- as.character(args[2])

fit.list <- in.file %>% readRDS

ln.p.y.vec <- fit.list %>%
    sapply(function(x) x$ln.p.y)

saveRDS(ln.p.y.vec, file=out.file)
```
