## Supplemental File 2 for "Probabilistic classification of gene-by-treatment interactions on molecular count phenotypes": get_lnpy_for_map_lap_for_comp.R

```
suppressPackageStartupMessages(library(tidyverse))
suppressPackageStartupMessages(library(classifygxt))

args <- commandArgs(TRUE)
dir.prefix <- as.character(args[1])
lb <- as.integer(args[2])
ub <- as.integer(args[3])
step <- as.integer(args[4])
out.file <- as.character(args[5])

size <- 10

phi <- seq(lb / 100, ub / 100, step / 100) %>%
    round(2) %>%
    format(nsmall=2)
phi.matrix <- cbind(
    phi1=rep(phi, times=10^2),
    phi2=rep(rep(phi, each=10), times=10),
    phi3=rep(phi, each=10^2))
phi.name <- apply(phi.matrix, 1, paste, collapse="_") %>%
    sapply(function(x) paste0("phi", x)) %>%
    unname

# covert the characger matrix to numeric
phi.matrix <- apply(phi.matrix, 2, as.numeric)

# take the first `size` simulations for each model
subset <- rep(seq(0, 7, 1) * 1e3, each=size) +
    seq_len(size)

# create a vector to store the results
ln.p.y <- rep(NA, length(phi.name))
names(ln.p.y) <- phi.name 

for (i in seq_along(phi.name)) {

    ln.p.y[i] <- phi.name[i] %>%
        file.path(dir.prefix, ., "fit.rds") %>%
        readRDS %>%
        `[`(subset) %>%
        sapply(function(x) x$ln.p.y) %>%
        sum

}

saveRDS(ln.p.y, file=out.file)
```
