## Supplemental File 2 for "Probabilistic classification of gene-by-treatment interactions on molecular count phenotypes": get_lnpy_for_mcmc_bs.R

```
suppressPackageStartupMessages(library(tidyverse))
suppressPackageStartupMessages(library(classifygxt))

args <- commandArgs(TRUE)
in.dir <- as.character(args[1])
out.file <- as.character(args[2])

file.list <- list.files(
    in.dir, pattern="^res", full.names=TRUE)

ln.p.y.vec <- file.list %>%
    sapply(function(x) x %>%
                       readRDS %>%
                       pluck("ln.p.y")) %>%
    unname

saveRDS(ln.p.y.vec, file=out.file)
```
