## Supplemental File 2 for "Probabilistic classification of gene-by-treatment interactions on molecular count phenotypes": get_marginal_llk.R

```
suppressPackageStartupMessages(library(tidyverse))
suppressPackageStartupMessages(library(classifygxt))

args <- commandArgs(TRUE)
dir.prefix <- as.character(args[1])
lb <- as.integer(args[2])
ub <- as.integer(args[3])
step <- as.integer(args[4])
out.file <- as.character(args[5])

phi <- seq(lb / 100, ub / 100, step / 100) %>%
    round(2) %>%
    format(nsmall=2)
phi.matrix <- cbind(
    phi1=rep(phi, times=10^2),
    phi2=rep(rep(phi, each=10), times=10),
    phi3=rep(phi, each=10^2))
phi.name <- apply(phi.matrix, 1, paste, collapse="_") %>%
    sapply(function(x) paste0("phi", x)) %>%
    unname

# covert the characger matrix to numeric
phi.matrix <- apply(phi.matrix, 2, as.numeric)

ln.p.y <- phi.name %>% sapply(function(x)
    dir.prefix %>%
    file.path(x, "fit.rds") %>%
    readRDS %>%
    sapply(function(x) x$ln.p.y) %>%
    sum
    ) %>%
    `names<-`(phi.name) %>%
    cbind(phi.matrix, ln.p.y=.) %>%
    as.data.frame

saveRDS(ln.p.y, file=out.file)
```
