## Supplemental File 2 for "Probabilistic classification of gene-by-treatment interactions on molecular count phenotypes": get_pp_for_mcmc_bs.R

```
suppressPackageStartupMessages(library(tidyverse))
suppressPackageStartupMessages(library(classifygxt))

args <- commandArgs(TRUE)
in.dir <- as.character(args[1])
data.file <- as.character(args[2])
out.file <- as.character(args[3])

# read in and process the data
file.vec <- list.files(
    in.dir, pattern="^res", full.names=TRUE) %>%
    grep("rds$", ., value=TRUE)

pp <- file.vec %>%
    map(
        function(x) readRDS(x) %>%
                    get_pp) %>%
    do.call("rbind", .) %>%
    as.data.frame

data.list <- data.file %>% readRDS

index <- data.list %>%
    map_int(function(x) x$index)
model.name <- get_model_names()

pp <- pp %>%
    mutate(correct=model.name[index])

# save the result
saveRDS(pp, file=out.file)
```
