## Supplemental File 2 for "Probabilistic classification of gene-by-treatment interactions on molecular count phenotypes": get_rocc.R

```
suppressPackageStartupMessages(library(tidyverse))
suppressPackageStartupMessages(library(classifygxt))

args <- commandArgs(TRUE)
in.file <- as.character(args[1])
type <- as.character(args[2])
out.file <- as.character(args[3])

model.name <- get_model_names()

pp <- in.file %>% readRDS

if (class(pp$correct) == "character") {
    index <- pp$correct %>%
        sapply(function(x) which(x == model.name))
    pp <- pp %>%
        mutate(correct=index)
}

pp.mat <- pp %>%
    dplyr::select(-correct) %>%
    as.matrix
    
if (type == "nogxt") {
    pattern.vec <- c(1, 1, 1, 1, 0, 0, 0, 0)
} else if (type == "interaction") {
    pattern.vec <- c(0, 0, 0, 0, 1, 1, 1, 1)
} else if (type == "induced") {
    pattern.vec <- c(0, 0, 0, 0, 1, 0, 1, 0)
} else if (type == "altered") {
    pattern.vec <- c(0, 0, 0, 0, 0, 1, 0, 1)
}

truth.vec <- (pp$correct %in% which(pattern.vec == 1)) %>%
    as.numeric    

pp.vec <- rowSums(pp.mat[, pattern.vec == 1])
n.p <- sum(truth.vec == 1)
n.n <- sum(truth.vec == 0)

thresh.vec <- seq(0, 1, 0.001)
recall.vec <- prec.vec <- fpr.vec <-
    rep(NA, length(thresh.vec))
for (i in seq_along(thresh.vec)) {
    thresh <- thresh.vec[i]
    tp <- sum(pp.vec > thresh & truth.vec == 1)
    fp <- sum(pp.vec > thresh & truth.vec == 0)
    fn <- n.p - tp
    tn <- n.n - fp
    recall.vec[i] <- tp / (tp + fn)
    prec.vec[i] <- tp / (tp + fp)
    fpr.vec[i] <- fp / (fp + tn)        
}
d <- data.frame(
    recall=recall.vec,
    prec=prec.vec,
    fpr=fpr.vec)

aoc <- 0
n.thresh <- thresh.vec %>% length
for (i in 1:(n.thresh - 1)) {
    aoc <- aoc +
        recall.vec[i] * abs(fpr.vec[i + 1] - fpr.vec[i])
}

res.list <- list(roc=d, aoc=aoc)
saveRDS(res.list, file=out.file)
```
