## Supplemental File 2 for "Probabilistic classification of gene-by-treatment interactions on molecular count phenotypes": get_true_co.R

```
suppressPackageStartupMessages(library(tidyverse))
suppressPackageStartupMessages(library(classifygxt))

args <- commandArgs(TRUE)
in.file <- as.character(args[1])
out.file <- as.character(args[2])

data.list <- in.file %>% readRDS
n.data <- length(data.list)

true.co <- matrix(NA, nrow=n.data, ncol=2) %>%
    as.data.frame %>%
    `colnames<-`(c("index", "co"))
fn <- "nonlinear"
model.mat <- classifygxt:::get_model_mat()

for (i in seq_along(data.list)) {

    l <- data.list[[i]]
    true.co[i, 1] <- index <- l$index

    if (!is.null(l$b)) {
        beta <- l$b
    } else if (!is.null(l$beta)) {
        beta <- l$beta
    } else {
        stop("coefficients are not found")
    }

    m <- model.mat[index, ] %>% unname
    e00 <- classifygxt:::compute_nl(g=0, t=0, beta=beta, m=m)
    e01 <- classifygxt:::compute_nl(g=0, t=1, beta=beta, m=m)
    e20 <- classifygxt:::compute_nl(g=2, t=0, beta=beta, m=m)
    e21 <- classifygxt:::compute_nl(g=2, t=1, beta=beta, m=m)

    if ((e01 - e00) * (e21 - e20) < 0) {
        true.co[i, 2] <- 1
    } else {
        true.co[i, 2] <- 0
    }
}

saveRDS(true.co, file=out.file)
```
