## Supplemental File 2 for "Probabilistic classification of gene-by-treatment interactions on molecular count phenotypes": get_tu_lambda.R

```
suppressPackageStartupMessages(library(tidyverse))
suppressPackageStartupMessages(library(classifygxt))

args <- commandArgs(TRUE)
file.data <- as.character(args[1])
file.kin <- as.character(args[2]) # file name or "NA"
file.out <- as.character(args[3])

if (file.kin == "NA") {
    kinship <- NULL
} else {
    kinship <- file.kin %>% readRDS
}
    
# read in the data
data.list <- file.data %>% readRDS

# extract the first (or any) data
data <- data.list[[1]]

# get a list of tU and lambda
tu.lambda <- get_tu_lambda(data, kinship=kinship)

saveRDS(tu.lambda, file=file.out)
```
