## Supplemental File 2 for "Probabilistic classification of gene-by-treatment interactions on molecular count phenotypes": make_data.R

```
suppressPackageStartupMessages(library(tidyverse))
suppressPackageStartupMessages(library(classifygxt))

args <- commandArgs(TRUE)
ranef <- as.character(args[1])
file.out <- as.character(args[2])

num <- rep(1000, 8)

n.sub <- 80 # number of subjects
anno <- data.frame(
    subject=rep(seq_len(n.sub), each=2),
    condition=rep(c(0, 1), times=n.sub))

sd.g <- 1.5
sd.t <- 2.0
sd.gxt <- 1.0
sd <- c(sd.g, sd.t, sd.gxt)

if (ranef == "noranef") { # without random effect
    data.list <- make_data(
        anno=anno, fn="nonlinear",
        num=num, sd=sd)
} else if (ranef == "ranef") { # with random effect
    ranef <- TRUE
    sigma.u <- sqrt(0.2)
    kinship <- NULL
    data.list <- make_data(
        anno=anno, fn="nonlinear",
        num=num, sd=sd,
        ranef=ranef, sigma.u=sigma.u)
}

saveRDS(data.list, file=file.out)
```
