## Supplemental File 2 for "Probabilistic classification of gene-by-treatment interactions on molecular count phenotypes": optimize_hp.R

```
suppressPackageStartupMessages(library(tidyverse))
suppressPackageStartupMessages(library(classifygxt))

args <- commandArgs(TRUE)
file.data <- as.character(args[1])
seed <- as.integer(args[2])
phi1 <- as.numeric(args[3])
phi2 <- as.numeric(args[4])
phi3 <- as.numeric(args[5])
method <- as.character(args[6])
ranef <- as.character(args[7])
file.tu.lambda <- as.character(args[8])
file.out <- as.character(args[9])

# read in the data
data.list <- file.data %>% readRDS

# specify model prior
p.m <- rep(1/8, 8)

phi <- c(phi1, phi2, phi3)
computation <- "map.lap"

if (ranef == "ranef") {
    ranef <- TRUE
    tu.lambda <- file.tu.lambda %>% readRDS
} else if (ranef == "noranef") {
    ranef <- FALSE
    tu.lambda <- NULL
}    

if (method == "nl") {
    fn <- "nonlinear"
    rint <- FALSE
} else if (method == "lm") {
    fn <- "linear"
    rint <- FALSE
} else if (method == "rint") {    
    fn <- "linear"
    rint <- TRUE
}

res.list <- lapply(
    X=data.list,
    FUN=do_bms,
    p.m=p.m, phi=phi,
    fn=fn, rint=rint, method=computation,
    tu.lambda=tu.lambda)

saveRDS(res.list, file=file.out)
```
