## Supplemental File 2 for "Probabilistic classification of gene-by-treatment interactions on molecular count phenotypes": plot_calib.R

```
suppressPackageStartupMessages(library(tidyverse))

format_data <- function(l.bs, alpha=0.05) {
    v.bin <- l.bs$v.bin
    v.x <- l.bs$v.x
    v.n <- l.bs$v.n
    v.bs.bin <- l.bs$v.bs.bin

    n.bin <- length(v.bin) - 1
    size.bin <- 0.1
    v.pi <- (v.bin + (size.bin / 2)) %>%
        `[`(seq_len(n.bin))

    d <- data.frame(x=v.pi, y=v.x)
    
    se <- sqrt(v.pi * (1 - v.pi) / v.n)
    lb <- v.x - qnorm(1 - alpha/2) * se
    ub <- v.x + qnorm(1 - alpha/2) * se
    d.se <- data.frame(x=v.pi, y1=lb, y2=ub)

    output <- list(d=d, d.se=d.se)
    output
}

args <- commandArgs(TRUE)
in.dir <- as.character(args[1])
fig.dir <- as.character(args[2])
out.file <- as.character(args[3])
fig.file <- as.character(args[4])

if (!dir.exists(fig.dir)) dir.create(fig.dir, recursive=TRUE)

method.vec <- c("nl", "lm", "rint")
method.name <- c("log-NL", "log-LM", "RINT-LM")
n.method <- length(method.vec)

cat.vec <- c("nogxt", "induced", "altered")
cat.name <- c("No GxT", "Induced", "Altered")
n.cat <- length(cat.vec)

dir.vec <- file.path(in.dir, method.vec)
file.vec <- paste0(
    rep(dir.vec, each=n.method), "/calib.",
    rep(cat.vec, times=n.cat), ".rds")

calib.list <- file.vec %>%
    map(readRDS) %>%
    map(format_data) 

d <- calib.list %>%
    map(function(x) x$d) %>%
    do.call("rbind", .)

n.pt <- nrow(d) / length(file.vec)

method.fact <-
    rep(method.name, each=n.pt * n.cat) %>%
    factor(levels=method.name)
cat.fact <-
    rep(rep(cat.name, each=n.pt), times=n.method) %>%
    factor(levels=cat.name)

d <- d %>%
    mutate(method=method.fact) %>%
    mutate(cat=cat.fact)

d.se <- calib.list %>%
    map(\(x) x$d.se) %>%
    do.call("rbind", .) %>%
    mutate(method=method.fact) %>%
    mutate(cat=cat.fact)

output <- list(data=d, se=d.se)
saveRDS(output, file=out.file)

# plot
p <- ggplot() +
    geom_point(
        mapping=aes(x=x, y=y),
        data=d)
p <- p +
    geom_segment(
        mapping=aes(x=x, y=y1, xend=x, yend=y2),
        data=d.se)
p <- p +
    geom_abline(intercept=0, slope=1, colour="dodgerblue")
# p <- p + xlim(0, 1) + ylim(0, 1)
p <- p +
    xlab("Posterior probability") +
    ylab("Fraction of positives")
p <- p + scale_x_continuous(
             limits=c(0, 1),
             breaks=c(0, 0.5, 1),
             labels=c("0.0", "0.5", "1.0"))
p <- p + scale_y_continuous(
             limits=c(0, 1),
             breaks=c(0, 0.5, 1),
             labels=c("0.0", "0.5", "1.0"))
# p <- p + theme(aspect.ratio=1)
p <- p + theme(legend.position="none")
# p <- p + theme(panel.grid.major=element_blank())
# p <- p + theme(panel.grid.minor=element_blank())
p <- p + theme(panel.grid.major=element_line(
                   colour="gray", linewidth=rel(0.6)))
p <- p + theme(panel.grid.minor=element_line(
                   colour="gray", linewidth=rel(0.6)))
p <- p + theme(panel.background=element_blank())
p <- p + theme(panel.border=element_rect(
                   colour="gray", fill=NA, linewidth=1))
p <- p + facet_grid(cat ~ method)

p <- p + coord_equal(xlim=c(0, 1), ylim=c(0, 1))

pdf(fig.file, width=7, height=7)
p
dev.off()

# ggsave(filename=fig.file, width=7, height=7)
```
