## Supplemental File 2 for "Probabilistic classification of gene-by-treatment interactions on molecular count phenotypes": plot_co.R

```
suppressPackageStartupMessages(library(tidyverse))
suppressPackageStartupMessages(library(classifygxt))

cols <- c(rgb(230/255, 159/255, 0/255),
          rgb(86/255, 180/255, 233/255),
          rgb(0/255, 158/255, 115/255))

args <- commandArgs(TRUE)
true.file <- as.character(args[1])
in.dir <- as.character(args[2])
fig.dir <- as.character(args[3])
out.file <- as.character(args[4])
fig.file <- as.character(args[5])

method.vec <- c("nl", "lm", "rint")
method.name <- c("log-NL", "log-LM", "RINT-LM")
file.vec <- file.path(in.dir, method.vec, "p.co.rds")

true.co <- true.file %>% readRDS

p.co <- file.vec %>%
    map(readRDS) %>%
    map(rowSums) %>%
    bind_cols

d <- bind_cols(true.co, p.co) %>%
    `colnames<-`(
        c("index", "truth", method.name)) %>%
    pivot_longer(all_of(method.name)) %>%
    mutate(name=factor(name, levels=method.name)) %>%
    mutate(truth=factor(truth, levels=c("1", "0"))) %>%
    mutate(cat=factor("Crossover"))

saveRDS(d, file=out.file)

lab.truth <- c("Correct", "Incorrect") %>%
    `names<-`(c("1", "0"))
lab <- labeller(truth=lab.truth)

p <- d %>% ggplot(aes(x=name, y=value, color=name)) +
    geom_violin(scale="width") +
    # geom_boxplot(outlier.shape=NA, width=0.1) +
    stat_summary(
        fun=median, geom="point", size=1) +
    ylim(0, 1) +
    ylab("Posterior probability") +
    # facet_wrap(vars(correct), ncol=8, scales="free") +
    # theme_classic() +
    theme(axis.ticks.x=element_blank(),
          axis.text.x=element_blank(),
          axis.title.x=element_blank(),
          legend.title=element_blank(),
          legend.position="bottom") +
    facet_grid(
        vars(truth), vars(cat),
        labeller=lab, scales="free")
p <- p + scale_color_manual(values=cols)

pdf(file=fig.file, width=3, height=6)
p
dev.off()
```
