## Supplemental File 2 for "Probabilistic classification of gene-by-treatment interactions on molecular count phenotypes": plot_compare_pp.R

```
suppressPackageStartupMessages(library(tidyverse))
suppressPackageStartupMessages(library(classifygxt))

args <- commandArgs(TRUE)
mcmc.bs.dir <- as.character(args[1])
map.lap.dir <- as.character(args[2])
out.file <- as.character(args[3])
fig.file <- as.character(args[4])

method.level <- c("nl", "lm", "rint")
hp.level <- c("rest", "opt", "perm")
method.vec <-
    rep(c("nl", "lm", "rint"), each=3)
hp.vec <-
    rep(c("rest", "opt", "perm"), times=3)

mcmc.bs.list <-
    file.path(
        mcmc.bs.dir, method.vec, hp.vec, "pp.rds")
map.lap.list <-
    file.path(
        map.lap.dir, method.vec, hp.vec, "pp.rds")

df.list <- vector("list", length(mcmc.bs.list)) 

for (i in seq_along(mcmc.bs.list)) {

    mcmc.bs.file <- mcmc.bs.list[[i]]
    map.lap.file <- map.lap.list[[i]]
    mcmc.bs.mat <- mcmc.bs.file %>%
        readRDS %>%
        dplyr::select(-correct) %>%
        as.matrix
    map.lap.mat <- map.lap.file %>%
        readRDS %>%
        dplyr::select(-correct) %>%
        as.matrix
    
    if (nrow(mcmc.bs.mat) != nrow(map.lap.mat)) {
        # take the first 100 simulations for each model
        subset <- rep(
            seq(0, 7, 1) * 1e3, each=100) + 1:100
        map.lap.mat <- map.lap.mat[subset, ]
    }

    mcmc.bs.vec <- mcmc.bs.mat %>% c
    map.lap.vec <- map.lap.mat %>% c

    d <- data.frame(
        mcmc.bs=mcmc.bs.vec,
        map.lap=map.lap.vec,
        method=method.vec[i],
        hp=hp.vec[i])
    
    df.list[[i]] <- d
        
}

df <- df.list %>%
    bind_rows %>%
    mutate(method=factor(method, levels=method.level)) %>%
    mutate(hp=factor(hp, levels=hp.level))

# save the data frame
saveRDS(df, out.file)

# make plot
map.method <- c("log-NL", "log-LM", "RINT-LM") %>%
    `names<-`(method.level)
map.hp <- c("Restrictive", "Optimal", "Permissive") %>%
    `names<-`(hp.level)

p <- ggplot() +
    geom_point(
        mapping=aes(x=mcmc.bs, y=map.lap),
        data=df,
        size=0.01)
p <- p +
    xlab("MCMC & bridge sampling") +
    ylab("MAP & Laplace") 
p <- p +
    facet_grid(
        vars(method), vars(hp),
        labeller=labeller(
            method=map.method,
            hp=map.hp))
p <- p + coord_equal()
# p <- p + coord_equal(xlim=c(0, 1), ylim=c(0, 1))

pdf(fig.file, width=7, height=7)
p
dev.off()

# ggsave(filename=fig.file, plot=p, width=7, height=7)
```
