## Supplemental File 2 for "Probabilistic classification of gene-by-treatment interactions on molecular count phenotypes": plot_compare_sum_lnpy.R

```
suppressPackageStartupMessages(library(tidyverse))
suppressPackageStartupMessages(library(classifygxt))

args <- commandArgs(TRUE)
mcmc.bs.dir <- as.character(args[1])
map.lap.dir <- as.character(args[2])
fig.dir <- as.character(args[3])
out.file <- as.character(args[4])
fig.file <- as.character(args[5])

if (!dir.exists(fig.dir)) dir.create(fig.dir, recursive=TRUE)

method.level <- c("nl", "lm", "rint")

mcmc.bs.file.vec <-
    file.path(
        mcmc.bs.dir, method.level, "sum.lnpy.vec.rds")
map.lap.file.vec <-
    file.path(
        map.lap.dir, method.level, "sum.lnpy.vec.rds")

df.list <- vector("list", length(method.level)) 

for (i in seq_along(method.level)) {

    mcmc.bs.file <- mcmc.bs.file.vec[[i]]
    map.lap.file <- map.lap.file.vec[[i]]
    mcmc.bs.vec <- mcmc.bs.file %>%
        readRDS
    map.lap.vec <- map.lap.file %>%
        readRDS
    
    d <- data.frame(
        mcmc.bs=mcmc.bs.vec,
        map.lap=map.lap.vec,
        method=method.level[i])
    
    df.list[[i]] <- d
        
}

df <- df.list %>%
    bind_rows %>%
    mutate(method=factor(method, levels=method.level))

# save the data frame
saveRDS(df, out.file)

# ggplot2
map.method <- c("log-NL", "log-LM", "RINT-LM") %>%
    `names<-`(method.level)

p <- ggplot() +
    geom_point(
        mapping=aes(x=mcmc.bs, y=map.lap),
        data=df,
        size=0.01)
p <- p +
    xlab("MCMC & bridge sampling") +
    ylab("MAP & Laplace") 
p <- p + theme(aspect.ratio=1)
p <- p +
    facet_wrap(
        vars(method),
        labeller=labeller(
            method=map.method),
        scales="free")
# p <- p + coord_equal()
# p <- p + coord_equal(xlim=c(0, 1), ylim=c(0, 1))

ggsave(filename=fig.file, plot=p, width=9, height=3)
```
