## Supplemental File 2 for "Probabilistic classification of gene-by-treatment interactions on molecular count phenotypes": plot_hist.R

```
suppressPackageStartupMessages(library(dplyr))
suppressPackageStartupMessages(library(tibble))
suppressPackageStartupMessages(library(ggplot2))
suppressPackageStartupMessages(library(tidyr))
suppressPackageStartupMessages(library(magrittr))

args <- commandArgs(TRUE)
in.file <- as.character(args[1])
comp <- as.character(args[2])
out.dir <- as.character(args[3])
out.file <- as.character(args[4])

if (!dir.exists(out.dir)) dir.create(out.dir, recursive=TRUE)

if (comp == "mcmc-bs") {
    limits <- c(0, 20)
    breaks <- c(0, 10, 20)
    labels <- c("0", "10", "20")
} else if (comp == "map-lap") {    
    limits <- c(0, 200)
    breaks <- c(0, 100, 200)
    labels <- c("0", "100", "200")
}

dat <- readRDS(in.file)
dat <- dat %>%
    as_tibble() %>%
    mutate(sim=seq_len(n()),
           correct=as.character(correct))

p <- dat %>%
    pivot_longer(
        cols=1:8, names_to="model", values_to="pp") %>%
    group_by(sim) %>%
    mutate(top = model[which.max(pp)]) %>%
    filter(model == top) %>%
    ggplot(aes(pp)) +
    geom_histogram() +
    facet_grid(correct ~ top) # +
    # xlim(0, 1.1)
p <- p +
    xlab("Posterior probability") +
    ylab("Count")
# p <- p + labs(y="Model", x="Feature-SNP pairs")
# p <- p + theme(plot.title=element_text(size=rel(1.2), hjust=0.5))
# p <- p + theme(plot.margin=unit(c(1, 1, 1, 1), "cm"))
# p <- p + theme(plot.margin=margin(r=0))
p <- p + scale_x_continuous(
             limits=c(0, 1.05),
             breaks=c(0, 0.5, 1),
             labels=c("0.0", "0.5", "1.0"),
             oob=scales::squish)
             # labels=scales::number_format(accuracy=0.1))
p <- p + scale_y_continuous(
             limits=limits,
             breaks=breaks,
             labels=labels)
# p <- p + scale_y_continuous(
#              limits=c(0, 20),
#              breaks=c(0, 10, 20),
#              labels=c("0", "10", "20"))
# p <- p + theme(axis.ticks=element_blank())
# p <- p + theme(axis.line=element_blank())
p <- p + theme(
             axis.text.y=element_text(
                 hjust=1, size=rel(1)))
p <- p + theme(
             axis.text.x=element_text(
                 angle=90, vjust=0.5, hjust=1))
# p <- p + theme(
             # panel.grid.major=element_blank(),
             # panel.grid.minor=element_blank(),
             # panel.border=element_blank(),
             # panel.background=element_blank())
# p <- p + theme(panel.margin.x=unit(0, "cm"))
# p <- p + theme(panel.margin.y=unit(0, "cm"))
p <- p + theme(legend.position="none")

ggsave(filename=out.file, plot=p, width=7, height=7)
```
