## Supplemental File 2 for "Probabilistic classification of gene-by-treatment interactions on molecular count phenotypes": plot_rocc.R

```
suppressPackageStartupMessages(library(tidyverse))
suppressPackageStartupMessages(library(here))

args <- commandArgs(TRUE)
in.dir <- as.character(args[1])
data.file <- as.character(args[2])
fig.file <- as.character(args[3])

method.vec <- c("nl", "lm", "rint")
method.name <- c("log-NL", "log-LM", "RINT-LM")
n.method <- length(method.vec)

cat.vec <- c("nogxt", "induced", "altered")
cat.name <- c("No GxT", "Induced", "Altered")
n.cat <- length(cat.vec)

dir.vec <- file.path(in.dir, method.vec)
file.vec <- paste0(
    rep(dir.vec, each=n.method), "/rocc.",
    rep(cat.vec, times=n.cat), ".rds")

cols <- c(rgb(230/255, 159/255, 0/255),
          rgb(86/255, 180/255, 233/255),
          rgb(0/255, 158/255, 115/255))

# create a data frame
rocc.list <- lapply(file.vec, readRDS)
rocc <- rocc.list %>%
    map(function(x) x$roc) %>%
    do.call("rbind", .)
n.pt <- nrow(rocc) / length(file.vec)

method.fact <-
    rep(method.name, each=n.pt * n.cat) %>%
    factor(levels=method.name)
cat.fact <-
    rep(rep(cat.name, each=n.pt), times=n.method) %>%
    factor(levels=cat.name)
rocc <- rocc %>%
    mutate(method=method.fact) %>%
    mutate(cat=cat.fact)

saveRDS(rocc, file=data.file)

# plot
p <- ggplot() +
    geom_line(
        mapping=aes(x=fpr, y=recall, colour=method),
        data=rocc)
p <- p + geom_abline(intercept=0, slope=1, linetype=2)
p <- p + xlab("FPR") + ylab("TPR")

# modify scales
p <- p + scale_color_manual(values=cols)
# p <- p + scale_x_continuous(
#              labels=scales::number_format(accuracy=0.1))
# p <- p + scale_y_continuous(
#              labels=scales::number_format(accuracy=0.1))
p <- p + scale_x_continuous(
             breaks=c(0, 0.5, 1),
             labels=c("0.0", "0.5", "1.0"))
p <- p + scale_y_continuous(
             breaks=c(0, 0.5, 1),
             labels=c("0.0", "0.5", "1.0"))

# modify themes
p <- p + theme(aspect.ratio=1)
p <- p + theme(legend.position="bottom")
# p <- p + theme(panel.grid.major=element_blank())
# p <- p + theme(panel.grid.minor=element_blank())
p <- p + theme(panel.grid.major=element_line(
                   colour="gray", linewidth=rel(0.6)))
p <- p + theme(panel.grid.minor=element_line(
                   colour="gray", linewidth=rel(0.6)))
p <- p + theme(panel.background=element_blank())
p <- p + theme(panel.border=element_rect(
                   colour="gray", fill=NA, linewidth=1))

p <- p + facet_wrap(vars(cat))
p
ggsave(filename=fig.file, width=7, height=3)

# get aoc
# aoc.vec <- rocc.list %>%
#     map_dbl(function(x) x$aoc)
```
