## Supplemental File 2 for "Probabilistic classification of gene-by-treatment interactions on molecular count phenotypes": plot_vln.R

```
suppressPackageStartupMessages(library(tidyverse))
suppressPackageStartupMessages(library(classifygxt))

cols <- c(rgb(230/255, 159/255, 0/255),
          rgb(86/255, 180/255, 233/255),
          rgb(0/255, 158/255, 115/255))

args <- commandArgs(TRUE)
in.dir <- as.character(args[1])
fig.dir <- as.character(args[2])
out.file <- as.character(args[3])
fig.file <- as.character(args[4])

if (!dir.exists(fig.dir)) dir.create(fig.dir, recursive=TRUE)

method.vec <- c("nl", "lm", "rint")
method.name <- c("log-NL", "log-LM", "RINT-LM")
n.method <- length(method.vec)

cat.vec <- c("nogxt", "induced", "altered")
cat.name <- c("No GxT", "Induced", "Altered")
n.cat <- length(cat.vec)

model.name <- get_model_names()

# read in the data
dir.vec <- file.path(in.dir, method.vec, "opt")
file.vec <- paste0(dir.vec, "/pp.rds")
pp.list <- file.vec %>% map(readRDS)

n.dat <- nrow(pp.list[[1]])
pp <- pp.list %>% do.call("rbind", .)
pp$method <- rep(method.vec, each=n.dat) %>%
    factor(levels=method.vec)

if (class(pp$correct) != "character") {
    pp$correct <- model.name[pp$correct]
}

pp$correct <- pp$correct %>%
    factor(levels=model.name)

# get the posterior probability of the correct and incorrect models
dl <- pp %>%
    pivot_longer(-c(correct, method)) %>%
    mutate(name=factor(name, level=model.name))    
sel <- dl$correct == dl$name

dl.correct <- dl %>%
    filter(sel) %>%
    mutate(type="Correct")
dl.incorrect <- dl %>%
    filter(!sel) %>%
    mutate(type="Incorrect")

dl <- bind_rows(dl.correct, dl.incorrect) %>%
    mutate(type=factor(type, levels=c("Correct", "Incorrect")))

saveRDS(dl, file=out.file)

p <- dl %>%
    ggplot(aes(x=method, y=value, color=method)) +
    geom_violin(scale="width") +
    # geom_boxplot(outlier.shape=NA, width=0.1) +
    stat_summary(
        fun=median, geom="point", size=1) +
    ylim(0, 1) +
    ylab("Posterior probability") +
    facet_grid(
        vars(type), vars(name),
        # labeller=lab,
        scales="free") +
    # theme_classic() +
    theme(axis.ticks.x=element_blank(),
          axis.text.x=element_blank(),
          axis.title.x=element_blank(),
          legend.title=element_blank(),
          legend.position="bottom")
p <- p + scale_color_manual(values=cols)

p <- p + theme(panel.grid.major=element_line(
                   colour="gray", linewidth=rel(0.6)))
p <- p + theme(panel.grid.minor=element_line(
                   colour="gray", linewidth=rel(0.6)))
p <- p + theme(panel.background=element_blank())
p <- p + theme(panel.border=element_rect(
                   colour="gray", fill=NA, linewidth=1))
p <- p + theme(axis.ticks.x=element_blank())
p <- p + theme(axis.text.x=element_blank())
p <- p + theme(axis.title.x=element_blank())

pdf(file=fig.file, width=10, height=6)
p
dev.off()
```
