## Supplemental File 2 for "Probabilistic classification of gene-by-treatment interactions on molecular count phenotypes": plot_vln_aggr.R

```
suppressPackageStartupMessages(library(tidyverse))
suppressPackageStartupMessages(library(classifygxt))

cols <- c(rgb(230/255, 159/255, 0/255),
          rgb(86/255, 180/255, 233/255),
          rgb(0/255, 158/255, 115/255))

in.dir <- "../derived_data/sim/mcmc-bs-hp/ranef-ranef"
out.dir <- "../figures/sim/mcmc-bs-hp" 
fig.file <- "../figures/sim/mcmc-bs-hp/vln_aggr_ranef_ranef.pdf"

args <- commandArgs(TRUE)
in.dir <- as.character(args[1])
fig.dir <- as.character(args[2])
out.file <- as.character(args[3])
fig.file <- as.character(args[4])

if (!dir.exists(out.dir)) dir.create(out.dir, recursive=TRUE)

method.vec <- c("nl", "lm", "rint")
method.name <- c("log-NL", "log-LM", "RINT-LM")
n.method <- length(method.vec)

cat.vec <- c("nogxt", "ind", "alt")
cat.name <- c("No GxT", "Induced", "Altered")
n.cat <- length(cat.vec)

model.name <- get_model_names()

# read in the data
dir.vec <- file.path(in.dir, method.vec, "opt")
file.vec <- paste0(dir.vec, "/pp.rds")
pp.list <- file.vec %>% map(readRDS)

n.dat <- nrow(pp.list[[1]])
pp <- pp.list %>% do.call("rbind", .)
pp$method <- rep(method.vec, each=n.dat) %>%
    factor(levels=method.vec)

nogxt.vec <- 1:4
ind.vec <- c(5, 7)
alt.vec <- c(6, 8)
cat.list <- list(
    nogxt=nogxt.vec, ind=ind.vec, alt=alt.vec)

# aggregate columns
pp.aggr <- cat.list %>%
    map(function(x) pp[, x]) %>%
    map(rowSums) %>%
    bind_cols %>%
    mutate(method=pp$method)

pp.dl <- pp.aggr %>%
    pivot_longer(-method)

correct.vec <- pp %>%
    pull(correct) %>%
    map_int(function(x) which(x == model.name))

correct.aggr <- cat.list %>%
    map(function(x) as.numeric(correct.vec %in% x)) %>%
    bind_cols

correct.dl <- correct.aggr %>%
    pivot_longer(everything())

dl <- bind_cols(pp.dl, correct.dl) %>%
    dplyr::select(-4) %>%
    `colnames<-`(c("method", "name", "value", "type")) %>%
    mutate(name=factor(name, levels=cat.vec)) %>%
    mutate(type=factor(type, levels=c(1, 0)))

saveRDS(dl, file=out.file)

map.type <- c("Correct", "Incorrect") %>%
    `names<-`(c("1", "0"))
map.name <- c("No GxT", "Induced", "Altered") %>%
    `names<-`(levels(dl$name))

p <- dl %>%
    ggplot(aes(x=method, y=value, color=method)) +
    geom_violin(scale="width") +
    # geom_boxplot(outlier.shape=NA, width=0.1) +
    stat_summary(
        fun=median, geom="point", size=1) +
    ylim(0, 1) +
    ylab("Posterior probability") +
    facet_grid(
        vars(type), vars(name),
        labeller=labeller(
            type=map.type,
            name=map.name),
        scales="free") +
    # theme_classic() +
    theme(axis.ticks.x=element_blank(),
          axis.text.x=element_blank(),
          axis.title.x=element_blank(),
          legend.title=element_blank(),
          legend.position="bottom")
p <- p + scale_color_manual(
             values=cols,
             labels=method.name)
p <- p + theme(panel.grid.major=element_line(
                   colour="gray", linewidth=rel(0.6)))
p <- p + theme(panel.grid.minor=element_line(
                   colour="gray", linewidth=rel(0.6)))
p <- p + theme(panel.background=element_blank())
p <- p + theme(panel.border=element_rect(
                   colour="gray", fill=NA, linewidth=1))
p <- p + theme(axis.ticks.x=element_blank())
p <- p + theme(axis.text.x=element_blank())
p <- p + theme(axis.title.x=element_blank())

pdf(file=fig.file, width=6, height=6)
p
dev.off()
```
