## Supplemental File 2 for "Probabilistic classification of gene-by-treatment interactions on molecular count phenotypes": plot_vln_effect.R

```
suppressPackageStartupMessages(library(tidyverse))
suppressPackageStartupMessages(library(classifygxt))
suppressPackageStartupMessages(library(patchwork))

args <- commandArgs(TRUE)
in.dir <- as.character(args[1])
out.dir <- as.character(args[2])
fig.file <- as.character(args[3])

if (!dir.exists(out.dir)) dir.create(out.dir, recursive=TRUE)

method.vec <- c("nl", "lm", "rint")
method.name <- c("log-NL", "log-LM", "RINT-LM")
n.method <- length(method.vec)

hp.vec <- c("rest", "opt", "perm")
hp.name <- c("Restrictive", "Optimal", "Permissive")
n.hp <- length(hp.vec)

model.name <- get_model_names()

# read in the data
dir.vec <- file.path(in.dir, method.vec)
file.vec <- paste0(
    rep(dir.vec, each=n.method), "/",
    rep(hp.vec, times=n.hp), "/pp.rds")
pp.list <- file.vec %>% map(readRDS)

n.dat <- nrow(pp.list[[1]])
pp <- pp.list %>% do.call("rbind", .)

# correct
if (class(pp$correct) != "character") {
    pp$correct <- model.name[pp$correct]
}
pp$correct <- pp$correct %>%
    factor(levels=model.name)

# method, hp
method.fact <-
    rep(method.name, each=n.dat * n.hp) %>%
    factor(levels=method.name)
hp.fact <-
    rep(rep(hp.name, each=n.dat), times=n.method) %>%
    factor(levels=hp.name)

pp <- pp %>%
    mutate(method=method.fact) %>%
    mutate(hp=hp.fact)

dl <- pp %>%
    pivot_longer(-c(correct, method, hp)) %>%
    mutate(name=factor(name, level=model.name))

sel <- dl$correct == dl$name

dl.correct <- dl %>%
    filter(sel) %>%
    mutate(type="Correct")
dl.incorrect <- dl %>%
    filter(!sel) %>%
    mutate(type="Incorrect")

# plot
cols <- hcl.colors(n=10, palette="Berlin")[1:3]

p.correct <- dl.correct %>%
    ggplot(aes(x=hp, y=value, col=hp)) +
    geom_violin(scale="width") +
    stat_summary(
        fun=median, geom="point", size=1) +
    ylim(0, 1) +
    ylab("Posterior probability") +
    scale_color_manual(values=cols) +
    # facet_wrap(vars(correct), ncol=8, scales="free") +
    facet_grid(vars(method), vars(name)) +
    # theme_classic() +
    theme(axis.ticks.x=element_blank(),
          axis.text.x=element_blank(),
          axis.title.x=element_blank(),
          legend.title=element_blank(),
          legend.position="none")

p.incorrect <- dl.incorrect %>%
    ggplot(aes(x=hp, y=value, col=hp)) +
    geom_violin(scale="width") +
    stat_summary(
        fun=median, geom="point", size=1) +
    ylim(0, 1) +
    ylab("Posterior probability") +
    scale_color_manual(values=cols) +
    # facet_wrap(vars(correct), ncol=8, scales="free") +
    facet_grid(vars(method), vars(name)) +
    # theme_classic() +
    theme(axis.ticks.x=element_blank(),
          axis.text.x=element_blank(),
          axis.title.x=element_blank(),
          legend.title=element_blank(),
          legend.position="none")

(p.correct / p.incorrect) +
    plot_layout(guides="collect") &
    theme(legend.position="bottom") &
    plot_annotation(tag_levels="A") # &
    # theme(plot.tag=element_text(size=rel(1.1)))

ggsave(file=fig.file, width=8, height=12)
# ggsave(file=fig.file, width=7.2, height=10.8)
# ggsave(file=fig.file, width=10, height=16)
```
