## Supplemental File 2 for "Probabilistic classification of gene-by-treatment interactions on molecular count phenotypes": plot_vln_effect_aggr.R

```
suppressPackageStartupMessages(library(tidyverse))
suppressPackageStartupMessages(library(patchwork))
suppressPackageStartupMessages(library(classifygxt))

args <- commandArgs(TRUE)
in.dir <- as.character(args[1])
out.dir <- as.character(args[2])
fig.file <- as.character(args[3])

if (!dir.exists(out.dir)) dir.create(out.dir, recursive=TRUE)

method.vec <- c("nl", "lm", "rint")
method.name <- c("log-NL", "log-LM", "RINT-LM")
n.method <- length(method.vec)

hp.vec <- c("rest", "opt", "perm")
hp.name <- c("Restrictive", "Optimal", "Permissive")
n.hp <- length(hp.vec)

cat.vec <- c("nogxt", "ind", "alt")
cat.name <- c("No GxT", "Induced", "Altered")
n.cat <- length(cat.vec)

model.name <- get_model_names()

# read in the data
dir.vec <- file.path(in.dir, method.vec)
file.vec <- paste0(
    rep(dir.vec, each=n.method), "/",
    rep(hp.vec, times=n.hp), "/pp.rds")
pp.list <- file.vec %>% map(readRDS)

n.dat <- nrow(pp.list[[1]])
pp <- pp.list %>% do.call("rbind", .)

# correct
if (class(pp$correct) != "character") {
    pp$correct <- model.name[pp$correct]
}
pp$correct <- pp$correct %>%
    factor(levels=model.name)

method.fact <-
    rep(method.name, each=n.dat * n.hp) %>%
    factor(levels=method.name)
hp.fact <-
    rep(rep(hp.name, each=n.dat), times=n.method) %>%
    factor(levels=hp.name)

pp <- pp %>%
    mutate(method=method.fact) %>%
    mutate(hp=hp.fact)

cat.list <- list(
    nogxt=1:4, ind=c(5, 7), alt=c(6, 8))

# aggregate columns
pp.aggr <- cat.list %>%
    map(function(x) pp[, x]) %>%
    map(rowSums) %>%
    bind_cols %>%
    mutate(method=pp$method) %>%
    mutate(hp=pp$hp)

pp.dl <- pp.aggr %>%
    pivot_longer(-c(method, hp))

head(pp.dl)

correct.vec <- pp %>%
    pull(correct) %>%
    map_int(function(x) which(x == model.name))

correct.aggr <- cat.list %>%
    map(function(x) as.numeric(correct.vec %in% x)) %>%
    bind_cols

correct.dl <- correct.aggr %>%
    pivot_longer(everything())

dl <- correct.dl %>%
    dplyr::select(-name) %>%
    bind_cols(pp.dl, .) %>%
    `colnames<-`(
        c("method", "hp", "name", "value", "type")) %>%
    mutate(name=factor(name, levels=cat.vec)) %>%
    mutate(type=factor(type, levels=c(1, 0)))

map.type <- c("Correct", "Incorrect") %>%
    `names<-`(c("1", "0"))
map.name <- c("No GxT", "Induced", "Altered") %>%
    `names<-`(levels(dl$name))

# plot
cols <- hcl.colors(n=10, palette="Berlin")[1:3]

p.correct <- dl %>%
    filter(type == 1) %>%
    ggplot(aes(x=hp, y=value, col=hp)) +
    geom_violin(scale="width") +
    stat_summary(
        fun=median, geom="point", size=1) +
    ylim(0, 1) +
    ylab("Posterior probability") +
    scale_color_manual(values=cols) +
    # facet_wrap(vars(correct), ncol=8, scales="free") +
    facet_grid(
        vars(method), vars(name),
        labeller=labeller(
            type=map.type,
            name=map.name),
        scales="free") +
    theme(axis.ticks.x=element_blank(),
          axis.text.x=element_blank(),
          axis.title.x=element_blank(),
          legend.title=element_blank(),
          legend.position="none")

p.incorrect <- dl %>%
    filter(type == 0) %>%
    ggplot(aes(x=hp, y=value, col=hp)) +
    geom_violin(scale="width") +
    stat_summary(
        fun=median, geom="point", size=1) +
    ylim(0, 1) +
    ylab("Posterior probability") +
    scale_color_manual(values=cols) +
    # facet_wrap(vars(correct), ncol=8, scales="free") +
    facet_grid(
        vars(method), vars(name),
        labeller=labeller(
            type=map.type,
            name=map.name),
        scales="free") +
    theme(axis.ticks.x=element_blank(),
          axis.text.x=element_blank(),
          axis.title.x=element_blank(),
          legend.title=element_blank(),
          legend.position="none")

(p.correct | p.incorrect) +
    plot_layout(guides="collect") &
    theme(legend.position="bottom") &
    plot_annotation(tag_levels="A")

ggsave(file=fig.file, width=10, height=9)
```
