## Supplemental File 2 for "Probabilistic classification of gene-by-treatment interactions on molecular count phenotypes": snake.sh

```
#!/bin/bash

#SBATCH --job-name=snake
#SBATCH --time=08-00:00:00
#SBATCH --mem=2G

module load python/3.8.8

snakemake --snakefile Snakefile --latency-wait 60 --configfile config/config.yaml --cluster 'sbatch -N 1 -n {cluster.threads} --mem {cluster.mem} -t {cluster.time} -p {cluster.queue} -o {cluster.output}' --jobs 100 --cluster-config config/sbatch.yaml
```
