## Supplemental File 2 for "Probabilistic classification of gene-by-treatment interactions on molecular count phenotypes": Snakefile

```
lb = config["lb"] # lower bound
ub = config["ub"] # upper bound
step = config["step"] # step
lb_comp = config["lb_comp"]
ub_comp = config["ub_comp"]
step_comp = config["step_comp"]
size = config["size"] # batch size for mcmc
num_hp = config["num_hp"] # for sensitivity analysis
num_post = config["num_post"] # for post-processing
num_comp = config["num_comp"] # for comparing two computatonal methods
seed = config["seed"]
num_core = config["num_core"]

method = ["nl", "lm", "rint"]
gen = ["noranef", "ranef"]
fit = ["noranef", "ranef"]
comp = ["map-lap", "mcmc-bs"]
category = ["nogxt", "induced", "altered"]
hp =["rest", "opt", "perm"]

numbers = [x / 100.00 for x in range(lb, ub + 1, step)]
phi1 = ["{:1.2f}".format(x) for x in numbers]
phi2 = ["{:1.2f}".format(x) for x in numbers]
phi3 = ["{:1.2f}".format(x) for x in numbers]
numbers_comp = [x / 100.00 for x in range(lb_comp, ub_comp + 1, step_comp)]
phi1_comp = ["{:1.2f}".format(x) for x in numbers_comp]
phi2_comp = ["{:1.2f}".format(x) for x in numbers_comp]
phi3_comp = ["{:1.2f}".format(x) for x in numbers_comp]

start_hp = [format(number, "04") for number in range(1, num_hp + 1, size)] 
start_post = [format(number, "04") for number in range(1, num_post + 1, size)] 
start_comp = [format(number, "04") for number in range(1, num_comp + 1, size)]
num_each = num_hp / 8
num_comp_each = num_comp / 8

rule all:
    input:
        expand(config["fig_dir"]
               + "comparison/{gen}-{fit}/"
               "compare_sum_lnpy.pdf",
               gen = gen, fit = fit),
        expand(config["dir"]
               + "comparison/{gen}-{fit}/"
               "compare.sum.lnpy.rds",
               gen = gen, fit = fit),
        expand(config["dir"]
               + "comparison/map-lap/{gen}-{fit}/"
               "{method}/sum.lnpy.vec.rds",
               gen = gen, fit = fit,
               method = method),
        expand(config["dir"]
               + "comparison/mcmc-bs/{gen}-{fit}/"
               "{method}/sum.lnpy.vec.rds",
               gen = gen, fit = fit,
               method = method),
        expand(config["dir"]
               + "comparison/{gen}-{fit}/pp.rds",
               gen = gen, fit = fit),
        expand(config["fig_dir"]
               + "comparison/{gen}-{fit}/pp.pdf",
               gen = gen, fit = fit),
        expand(config["dir"]
               + "comparison/mcmc-bs/"
               "{gen}-{fit}/{method}/"
               "phi{phi1}_{phi2}_{phi3}/"
               "complete{start}_{size}.txt",
               gen = gen, fit = fit, method = method,
               phi1 = phi1_comp, phi2 = phi2_comp,
               phi3 = phi3_comp, start = start_comp,
               size = size),
        expand(config["dir"]
               + "comparison/data/data.{gen}.rds",
               gen = gen, size = size),
        expand(config["dir"]
               + "post/{gen}-{fit}/plot.rocc.co.rds",
               gen = gen, fit = fit),
        expand(config["fig_dir"]
               + "post/{gen}-{fit}/rocc_co.pdf",
               gen = gen, fit = fit),
        expand(config["dir"]
               + "post/{gen}-{fit}/{method}/rocc.co.rds",
               gen = gen, fit = fit, method = method),
        expand(config["dir"]
               + "post/{gen}-{fit}/plot.calib.co.rds",
               gen = gen, fit = fit),
        expand(config["fig_dir"]
               + "post/{gen}-{fit}/calib_co.pdf",
               gen = gen, fit = fit),
        expand(config["dir"]
               + "post/{gen}-{fit}/{method}/calib.co.rds",
               gen = gen, fit = fit, method = method),
        expand(config["dir"]
               + "post/{gen}-{fit}/vln.co.rds",
               gen = gen, fit = fit),
        expand(config["fig_dir"]
               + "post/{gen}-{fit}/vln_co.pdf",
               gen = gen, fit = fit),
        expand(config["dir"]
               + "post/true.co.{gen}.rds",
               gen = gen),
        expand(config["dir"]
               + "post/{gen}-{fit}/{method}/p.co.rds",
               gen = gen, fit = fit,
               method = method),
        expand(config["dir"]
               + "post/{gen}-{fit}/{method}/"
               + "fit/complete{start}_{size}.txt",
               gen = gen, fit = fit, method = method,
               start = start_post, size = size),
        expand(config["fig_dir"]
               + "{comp}/{gen}-{fit}/"
               "rocc.pdf",
               comp = comp, gen = gen, fit = fit),    
        expand(config["dir"]
               + "{comp}/{gen}-{fit}/"
               "{method}/rocc.{category}.rds",
               comp = comp, gen = gen, fit = fit,
               method = method, category = category),
        expand(config["fig_dir"]
               + "{comp}/{gen}-{fit}/"
               "calib.pdf",
               comp = comp, gen = gen, fit = fit),
        expand(config["dir"]
               + "{comp}/{gen}-{fit}/"
               "{method}/calib.{category}.rds",
               comp = comp, gen = gen, fit = fit,
               method = method,
               category = category),
        expand(config["fig_dir"]
               + "{comp}/{gen}-{fit}/"
               "vln_effect.pdf",
               comp = comp, gen = gen, fit = fit),        
        expand(config["dir"]
               + "{comp}/{gen}-{fit}/"
               "vln.aggr.rds",
               comp = comp, gen = gen, fit = fit),
        expand(config["dir"]
               + "{comp}/{gen}-{fit}/vln.rds",
               comp = comp, gen = gen, fit = fit),
        expand(config["fig_dir"]
               + "{comp}/{gen}-{fit}/"
               "vln_effect_aggr.pdf",
               comp = comp, gen = gen, fit = fit),
        expand(config["fig_dir"]
               + "{comp}/{gen}-{fit}/"
               "vln_aggr.pdf",
               comp = comp, gen = gen, fit = fit),
        expand(config["fig_dir"]
               + "{comp}/{gen}-{fit}/"
               "vln.pdf",
               comp = comp, gen = gen, fit = fit),        
        expand(config["fig_dir"]
               + "{comp}/{gen}-{fit}/{method}/"
               "hist.pdf",
               comp = comp, gen = gen, fit = fit,
               method = method),
        expand(config["dir"]
               + "mcmc-bs-hp/{gen}-{fit}/{method}/"
               + "{hp}/pp.rds",
               gen = gen, fit = fit,
               method = method, hp = hp),
        expand(config["dir"]
               + "map-lap-hp/{gen}-{fit}/{method}/"
               + "{hp}/pp.rds",
               gen = gen, fit = fit,
               method = method, hp = hp),    
        expand(config["dir"]
               + "map-lap-hp/{gen}-{fit}/{method}/"
               + "{hp}/lnpy.vec.rds",
               gen = gen, fit = fit,
               method = method, hp = hp),       
        expand(config["dir"]
               + "mcmc-bs-hp/{gen}-{fit}/{method}/"
               + "{hp}/lnpy.vec.rds",
               gen = gen, fit = fit,
               method = method, hp = hp),        
        expand(config["dir"]
               + "mcmc-bs-hp/{gen}-{fit}/{method}/"
               + "{hp}/fit/complete{start}_{size}.txt",
               gen = gen, fit = fit,
               method = method, hp = hp,
               start = start_hp, size = size),
        expand(config["dir"]
               + "map-lap-hp/{gen}-{fit}/{method}/"
               + "{hp}/fit.rds",
               gen = gen, fit = fit,
               method = method, hp = hp),
        expand(config["dir"]
               + "optim/{gen}-{fit}/{method}/"
               "ln.p.y.rds",
               gen = gen, fit = fit,
               method = method),
        expand(config["dir"]
               + "optim/{gen}-{fit}/{method}/"
               "phi{phi1}_{phi2}_{phi3}/fit.rds",
               gen = gen, fit = fit,
               method = method,
               phi1 = phi1, phi2 = phi2, phi3 = phi3),

rule plot_compare_sum_lnpy:
    input:
        expand(config["dir"]
               + "comparison/{comp}/{{gen}}-{{fit}}/"
               "{method}/sum.lnpy.vec.rds",
               comp = comp, method = method),
    output:
        data = config["dir"]
               + "comparison/{gen}-{fit}/"
               "compare.sum.lnpy.rds",
        fig = config["fig_dir"]
               + "comparison/{gen}-{fit}/"
               "compare_sum_lnpy.pdf",
    shell:
        "module add r/4.1.3; "
        "Rscript plot_compare_sum_lnpy.R"
        " " + config["dir"]
        + "comparison/mcmc-bs/"
        "{wildcards.gen}-{wildcards.fit}"
        " " + config["dir"]
        + "comparison/map-lap/"
        "{wildcards.gen}-{wildcards.fit}"
        " " + config["fig_dir"]
        + "comparison/{wildcards.gen}-{wildcards.fit}"
        " {output.data}"
        " {output.fig}"

rule get_lnpy_for_mcmc_bs_for_comp:
    input:
        expand(config["dir"]
               + "comparison/mcmc-bs/{{gen}}-{{fit}}/"
               "{{method}}/"
               "phi{phi1}_{phi2}_{phi3}/"
               "complete{start}_{size}.txt",
               phi1 = phi1_comp,
               phi2 = phi2_comp, phi3 = phi3_comp,
               start = start_comp, size = size),
    output:
        config["dir"]
        + "comparison/mcmc-bs/{gen}-{fit}/"
        "{method}/sum.lnpy.vec.rds",
    shell:
        "module add r/4.1.3; "
        "Rscript get_lnpy_for_mcmc_bs_for_comp.R"
        " " + config["dir"]
        + "comparison/mcmc-bs/"
        "{wildcards.gen}-{wildcards.fit}/"
        "{wildcards.method}"
        " {lb_comp}"
        " {ub_comp}"
        " {step_comp}"
        " {output}"

rule get_lnpy_for_map_lap_for_comp:
    input:
        expand(config["dir"]
               + "optim/{{gen}}-{{fit}}/{{method}}/"
               "phi{phi1}_{phi2}_{phi3}/fit.rds",
               phi1 = phi1_comp, phi2 = phi2_comp,
               phi3 = phi3_comp),
    output:
        config["dir"]
        + "comparison/map-lap/{gen}-{fit}/{method}/"
        + "sum.lnpy.vec.rds",
    shell:
        "module add gcc/11.2.0; "
        "module add r/4.1.3; "
        "Rscript get_lnpy_for_map_lap_for_comp.R"
        " " + config["dir"]
        + "optim/{wildcards.gen}-{wildcards.fit}/"
        "{wildcards.method}"
        " {lb_comp}"
        " {ub_comp}"
        " {step_comp}"
        " {output}"

rule mcmc_bs_for_comp:
    input:
        data = config["dir"]
        + "comparison/data/data.{gen}.rds",
        tu_lambda = config["dir"]
        + "data/tu.lambda.rds",
    output:
        config["dir"]
        + "comparison/mcmc-bs/{gen}-{fit}/{method}/"
        "phi{phi1}_{phi2}_{phi3}/"
        "complete{start}_{size}.txt"
    shell:
        "module add gcc/11.2.0; "
        "module add r/4.1.3; "
        "Rscript do_mcmc_bs_for_comp.R"
        " {input.data}"
        " {input.tu_lambda}"
        " {seed}"
        " {num_core}"
        " {wildcards.phi1}"
        " {wildcards.phi2}"
        " {wildcards.phi3}"
        " {wildcards.method}"
        " {wildcards.fit}"
        " summary"
        " {wildcards.start}"
        " {wildcards.size}"
        " " + config["dir"]
        + "comparison/mcmc-bs/"
        "{wildcards.gen}-{wildcards.fit}/"
        "{wildcards.method}/"
        "phi{wildcards.phi1}_"
        "{wildcards.phi2}_"
        "{wildcards.phi3}"
        
rule subset_data_for_comp:
    input:
        config["dir"] + "data/data.{gen}.rds",
    output:
        config["dir"] + "comparison/data/data.{gen}.rds",
    shell:
        "module add gcc/11.2.0; "
        "module add r/4.1.3; "
        "Rscript subset_data.R"
        " {input}"
        " {num_comp_each}" # num of pairs for each model
        " {output}"
        
rule plot_rocc_co:
    input:
        expand(config["dir"]
               + "post/{{gen}}-{{fit}}/"
               "{method}/rocc.co.rds",
               method = method),
    output:
        fig = config["fig_dir"]
        + "post/{gen}-{fit}/rocc_co.pdf",
        data = config["dir"]
        + "post/{gen}-{fit}/plot.rocc.co.rds",
    shell:
        "module add r/4.1.3; "
        "Rscript plot_rocc_co.R"
        " " + config["dir"]
        + "post/{wildcards.gen}-{wildcards.fit}"
        " " + config["fig_dir"]
        + "post/{wildcards.gen}-{wildcards.fit}"
        " {output.fig}"
        " {output.data}"

rule get_rocc_co:
    input:
        co = config["dir"]
        + "post/{gen}-{fit}/"
        "{method}/p.co.rds",
        true = config["dir"]
        + "post/true.co.{gen}.rds",
    output:
        config["dir"]
        + "post/{gen}-{fit}/{method}/rocc.co.rds",
    shell:
        "module add r/4.1.3; "
        "Rscript get_rocc_co.R"
        " {input.co}"
        " {input.true}"
        " {output}"

rule plot_calib_co:
    input:
        expand(config["dir"]
               + "post/{{gen}}-{{fit}}/"
               "{method}/calib.co.rds",
               method = method),
    output:
        fig = config["fig_dir"]
        + "post/{gen}-{fit}/calib_co.pdf",
        data = config["dir"]
        + "post/{gen}-{fit}/plot.calib.co.rds",
    shell:
        "module add r/4.1.3; "
        "Rscript plot_calib_co.R"
        " " + config["dir"]
        + "post/{wildcards.gen}-{wildcards.fit}"
        " " + config["fig_dir"]
        + "post/{wildcards.gen}-{wildcards.fit}"
        " {output.fig}"
        " {output.data}"

rule get_calib_co:
    input:
        co = config["dir"]
        + "post/{gen}-{fit}/"
        "{method}/p.co.rds",
        true = config["dir"]
        + "post/true.co.{gen}.rds",
    output:
        config["dir"]
        + "post/{gen}-{fit}/{method}/calib.co.rds",
    shell:
        "module add r/4.1.3; "
        "Rscript get_calib_co.R"
        " {input.co}"
        " {input.true}"
        " {output}"

rule plot_co:
    input:
        true = config["dir"]
        + "post/true.co.{gen}.rds",
        co = expand(config["dir"]
                    + "post/{{gen}}-{{fit}}/"
                    "{method}/p.co.rds",
                    method = method),
    output:
        data = config["dir"]
        + "post/{gen}-{fit}/vln.co.rds",
        fig = config["fig_dir"]
        + "post/{gen}-{fit}/vln_co.pdf"
    shell:
        "module add r/4.1.3; "
        "Rscript plot_co.R"
        " {input.true}"
        " " + config["dir"]
        + "post/"
        "{wildcards.gen}-{wildcards.fit}"
        " " + config["fig_dir"]
        + "post/"
        "{wildcards.gen}-{wildcards.fit}"
        " {output.data}"
        " {output.fig}"

rule get_true_co:
    input:
        config["dir"]
        + "data/data.{gen}.rds",
    output:
        config["dir"]
        + "post/true.co.{gen}.rds",
    shell:
        "module add r/4.1.3; "
        "Rscript get_true_co.R"
        " {input}"
        " {output}"

rule get_co:
    input:
        expand(config["dir"]
               + "post/{{gen}}-{{fit}}/{{method}}/"
               + "fit/complete{start}_{size}.txt",
               start = start_post, size = size),
    output:
        config["dir"]
        + "post/{gen}-{fit}/{method}/p.co.rds",
    shell:
        "module add r/4.1.3; "
        "Rscript get_co.R"
        " " + config["dir"]
        + "post/"
        "{wildcards.gen}-{wildcards.fit}/"
        "{wildcards.method}/fit"
        " {output}"
        
rule plot_hist:
    input:
        config["dir"]
        + "{comp}-hp/{gen}-{fit}/{method}/opt/"
        "pp.rds",
    output:
        config["fig_dir"]
        + "{comp}/{gen}-{fit}/{method}/"
        "hist.pdf",
    shell:
        "module add r/4.1.3; "
        "Rscript plot_hist.R"
        " {input}"
        " {wildcards.comp}"
        " " + config["fig_dir"]
        + "{wildcards.comp}/"
        "{wildcards.gen}-{wildcards.fit}/"
        "{wildcards.method}"
        " {output}"
        
rule plot_vln:
    input:
        expand(config["dir"]
               + "{{comp}}-hp/{{gen}}-{{fit}}/"
               "{method}/opt/pp.rds",
               method = method)
    output:
        data = config["dir"]
        + "{comp}/{gen}-{fit}/vln.rds",
        fig = config["fig_dir"]
        + "{comp}/{gen}-{fit}/vln.pdf"
    shell:
        "module add r/4.1.3; "
        "Rscript plot_vln.R"
        " " + config["dir"]
        + "{wildcards.comp}-hp/"
        "{wildcards.gen}-{wildcards.fit}"
        " " + config["fig_dir"]
        + "{wildcards.comp}/"
        "{wildcards.gen}-{wildcards.fit}"
        " {output.data}"
        " {output.fig}"

rule plot_vln_aggr:        
    input:
        expand(config["dir"]
               + "{{comp}}-hp/{{gen}}-{{fit}}/"
               "{method}/opt/pp.rds",
               method = method)
    output:
        data = config["dir"]
        + "{comp}/{gen}-{fit}/vln.aggr.rds",
        fig = config["fig_dir"]
        + "{comp}/{gen}-{fit}/vln_aggr.pdf",
    shell:
        "module add r/4.1.3; "
        "Rscript plot_vln_aggr.R"
        " " + config["dir"]
        + "{wildcards.comp}-hp/"
        "{wildcards.gen}-{wildcards.fit}"
        " " + config["fig_dir"]
        + "{wildcards.comp}/"
        "{wildcards.gen}-{wildcards.fit}"
        " {output.data}"
        " {output.fig}"

rule plot_vln_effect:
    input:
        expand(config["dir"]
               + "{{comp}}-hp/{{gen}}-{{fit}}/"
               "{method}/{hp}/pp.rds",
               method = method, hp = hp)
    output:
        config["fig_dir"]
        + "{comp}/{gen}-{fit}/vln_effect.pdf"
    shell:
        "module add r/4.1.3; "
        "Rscript plot_vln_effect.R"
        " " + config["dir"]
        + "{wildcards.comp}-hp/"
        "{wildcards.gen}-{wildcards.fit}"
        " " + config["fig_dir"]
        + "{wildcards.comp}/"
        "{wildcards.gen}-{wildcards.fit}"
        " {output}"
        
rule plot_vln_effect_aggr:
    input:
        expand(config["dir"]
               + "{{comp}}-hp/{{gen}}-{{fit}}/"
               "{method}/{hp}/pp.rds",
               method = method, hp = hp)
    output:
        config["fig_dir"]
        + "{comp}/{gen}-{fit}/vln_effect_aggr.pdf"
    shell:
        "module add r/4.1.3; "
        "Rscript plot_vln_effect_aggr.R"
        " " + config["dir"]
        + "{wildcards.comp}-hp/"
        "{wildcards.gen}-{wildcards.fit}"
        " " + config["fig_dir"]
        + "{wildcards.comp}/"
        "{wildcards.gen}-{wildcards.fit}"
        " {output}"

rule plot_rocc:
    input:
        expand(config["dir"]
               + "{{comp}}/{{gen}}-{{fit}}/"
               "{method}/rocc.{category}.rds",
               method = method,
               category = category),
    output:
        data = config["dir"]
        + "{comp}/{gen}-{fit}/"
        "rocc.rds",
        fig = config["fig_dir"]
        + "{comp}/{gen}-{fit}/rocc.pdf",
    shell:
        "module add r/4.1.3; "
        "Rscript plot_rocc.R"
        " " + config["dir"]
        + "{wildcards.comp}/"
        "{wildcards.gen}-{wildcards.fit}"
        " {output.data}"
        " {output.fig}"
        
rule get_rocc_for_mcmc_bs:
    input:
        config["dir"]
        + "mcmc-bs-hp/{gen}-{fit}/"
        "{method}/opt/pp.rds",
    output:    
        config["dir"]
        + "mcmc-bs/{gen}-{fit}/"
        "{method}/rocc.{category}.rds",
    shell:
        "module add r/4.1.3; "
        "Rscript get_rocc.R"
        " {input}"
        " {wildcards.category}"
        " {output}"

rule get_rocc_for_map_lap:
    input:
        config["dir"]
        + "map-lap-hp/{gen}-{fit}/"
        "{method}/opt/pp.rds",
    output:    
        config["dir"]
        + "map-lap/{gen}-{fit}/"
        "{method}/rocc.{category}.rds",
    shell:
        "module add r/4.1.3; "
        "Rscript get_rocc.R"
        " {input}"
        " {wildcards.category}"
        " {output}"

rule plot_calib:
    input:
        expand(config["dir"]
               + "{{comp}}/{{gen}}-{{fit}}/"
               "{method}/calib.{category}.rds",
               method = method,
               category = category),
    output:
        data = config["dir"]
        + "{comp}/{gen}-{fit}/calib.rds",
        fig = config["fig_dir"]
        + "{comp}/{gen}-{fit}/calib.pdf",
    shell:
        "module add r/4.1.3; "
        "Rscript plot_calib.R"
        " " + config["dir"]
        + "{wildcards.comp}/"
        "{wildcards.gen}-{wildcards.fit}"
        " " + config["fig_dir"]
        + "{wildcards.comp}/"
        "{wildcards.gen}-{wildcards.fit}"
        " {output.data}"
        " {output.fig}"
        
rule get_calib_for_mcmc_bs:
    input:
        config["dir"]
        + "mcmc-bs-hp/{gen}-{fit}/"
        "{method}/opt/pp.rds",
    output:
        config["dir"]
        + "mcmc-bs/{gen}-{fit}/"
        "{method}/calib.{category}.rds",
    shell:
        "module add r/4.1.3; "
        "Rscript get_calib.R"
        " {input}"
        " {wildcards.category}"
        " {output}"

rule get_calib_for_map_lap:
    input:
        config["dir"]
        + "map-lap-hp/{gen}-{fit}/"
        "{method}/opt/pp.rds",
    output:
        config["dir"]
        + "map-lap/{gen}-{fit}/"
        "{method}/calib.{category}.rds",
    shell:
        "module add r/4.1.3; "
        "Rscript get_calib.R"
        " {input}"
        " {wildcards.category}"
        " {output}"

rule get_lnpy_for_mcmc_bs:
    input:
        expand(config["dir"]
               + "mcmc-bs-hp/{{gen}}-{{fit}}/"
               "{{method}}/{{hp}}/fit/"
               "complete{start}_{size}.txt",
               start = start_hp, size = size),
    output:
        config["dir"]
        + "mcmc-bs-hp/{gen}-{fit}/"
        "{method}/{hp}/lnpy.vec.rds",
    shell:
        "module add r/4.1.3; "
        "Rscript get_lnpy_for_mcmc_bs.R"
        " " + config["dir"]
        + "mcmc-bs-hp/"
        "{wildcards.gen}-{wildcards.fit}/"
        "{wildcards.method}/{wildcards.hp}/fit"
        " {output}"
        
rule get_lnpy_for_map_lap:
    input:
        config["dir"]
        + "map-lap-hp/{gen}-{fit}/"
        "{method}/{hp}/fit.rds",
    output:
        config["dir"]
        + "map-lap-hp/{gen}-{fit}/"
        "{method}/{hp}/lnpy.vec.rds",
    shell:
        "module add r/4.1.3; "
        "Rscript get_lnpy_for_map_lap.R"
        " {input}"
        " {output}"

rule plot_compare_pp:
    input:
       expand(config["dir"] +
              "{comp}-hp/{{gen}}-{{fit}}/"
              "{method}/{hp}/pp.rds",
              comp = comp, method = method, hp = hp),
    output:
       data = config["dir"]
       + "comparison/{gen}-{fit}/pp.rds",
       fig = config["fig_dir"]
       + "comparison/{gen}-{fit}/pp.pdf"
    shell:
        "module add r/4.1.3; "
        "Rscript plot_compare_pp.R"
        " " + config["dir"]
        + "mcmc-bs-hp/" # mcmc.bs dir
        "{wildcards.gen}-{wildcards.fit}"        
        " " + config["dir"]
        + "map-lap-hp/" # map.lap dir
        "{wildcards.gen}-{wildcards.fit}"        
        " {output.data}"
        " {output.fig}"
        
rule get_pp_for_mcmc_bs:
    input:
        fit = expand(config["dir"]
               + "mcmc-bs-hp/{{gen}}-{{fit}}/"
               "{{method}}/{{hp}}/fit/"
               "complete{start}_{size}.txt",
               start = start_hp, size = size),
        data = config["dir"]
        + "data/subset.{gen}.rds"
    output:
        config["dir"]
        + "mcmc-bs-hp/{gen}-{fit}/{method}/"
        + "{hp}/pp.rds",
    shell:
        "module add gcc/11.2.0; "
        "module add r/4.1.3; "
        "Rscript get_pp_for_mcmc_bs.R"
        " " + config["dir"]
        + "mcmc-bs-hp/"
        "{wildcards.gen}-{wildcards.fit}/"
        "{wildcards.method}/"
        "{wildcards.hp}/fit"
        " {input.data}"
        " {output}"
        
rule get_pp_for_map_lap:
    input:
        fit = config["dir"]
        + "map-lap-hp/{gen}-{fit}/{method}/"
        + "{hp}/fit.rds",
        data = config["dir"]
        + "data/data.{gen}.rds"
    output:
        config["dir"]
        + "map-lap-hp/{gen}-{fit}/{method}/"
        + "{hp}/pp.rds",
    shell:
        "module add gcc/11.2.0; "
        "module add r/4.1.3; "
        "Rscript get_pp_for_map_lap.R"
        " {input.fit}"
        " {input.data}"
        " {output}"
        
rule post:
    input:
        data = config["dir"] + "data/data.{gen}.rds",
        tu_lambda = config["dir"] + "data/tu.lambda.rds",
        fit = config["dir"]
        + "optim/{gen}-{fit}/{method}/"
        + "ln.p.y.rds",
    output:
        config["dir"]
        + "post/{gen}-{fit}/{method}/"
        + "fit/complete{start}_{size}.txt",
    shell:
        "module add gcc/11.2.0; "
        "module add r/4.1.3; "
        "Rscript do_mcmc_bs.R"
        " {input.data}"
        " {input.tu_lambda}"
        " {input.fit}"
        " {seed}"
        " {num_core}"
        " {wildcards.method}"
        " {wildcards.fit}"
        " opt"
        " sample"
        " {wildcards.start}"
        " {wildcards.size}"
        " " + config["dir"]
        + "post/{wildcards.gen}-{wildcards.fit}/"
        "{wildcards.method}/fit"

rule mcmc_bs_hp:
    input:
        data = config["dir"] + "data/subset.{gen}.rds",
        tu_lambda = config["dir"] + "data/tu.lambda.rds",
        fit = config["dir"]
        + "optim/{gen}-{fit}/{method}/"
        + "ln.p.y.rds",
    output:
        config["dir"]
        + "mcmc-bs-hp/{gen}-{fit}/{method}/"
        + "{hp}/fit/complete{start}_{size}.txt",
    shell:
        "module add gcc/11.2.0; "
        "module add r/4.1.3; "
        "Rscript do_mcmc_bs.R"
        " {input.data}"
        " {input.tu_lambda}"
        " {input.fit}"
        " {seed}"
        " {num_core}"
        " {wildcards.method}"
        " {wildcards.fit}"
        " {wildcards.hp}"
        " summary"
        " {wildcards.start}"
        " {wildcards.size}"
        " " + config["dir"]
        + "mcmc-bs-hp/{wildcards.gen}-{wildcards.fit}/"
        "{wildcards.method}/{wildcards.hp}/fit"
        
rule map_lap_hp:
    input:
        data = config["dir"] + "data/data.{gen}.rds",
        tu_lambda = config["dir"] + "data/tu.lambda.rds",
        fit = config["dir"]
        + "optim/{gen}-{fit}/{method}/"
        + "ln.p.y.rds",
    output:
        config["dir"]
        + "map-lap-hp/{gen}-{fit}/{method}/"
        + "{hp}/fit.rds",
    shell:
        "module add gcc/11.2.0; "
        "module add r/4.1.3; "
        "Rscript do_map_lap.R"
        " {input.data}"
        " {input.tu_lambda}"
        " {input.fit}"
        " {seed}"
        " {wildcards.method}"
        " {wildcards.fit}"
        " {wildcards.hp}"
        " {output}"
        
rule subset_data:
    input:
        config["dir"] + "data/data.{gen}.rds",
    output:
        config["dir"] + "data/subset.{gen}.rds",
    shell:
        "module add gcc/11.2.0; "
        "module add r/4.1.3; "
        "Rscript subset_data.R"
        " {input}"
        " {num_each}" # num of pairs per model
        " {output}"
        
rule get_marginal_llk:
    input:
        expand(config["dir"]
               + "optim/{{gen}}-{{fit}}/{{method}}/"
               "phi{phi1}_{phi2}_{phi3}/fit.rds",
               phi1 = phi1, phi2 = phi2, phi3 = phi3),
    output:
        config["dir"]
        + "optim/{gen}-{fit}/{method}/"
        + "ln.p.y.rds",
    shell:
        "module add gcc/11.2.0; "
        "module add r/4.1.3; "
        "Rscript get_marginal_llk.R"
        " " + config["dir"]
        + "optim/"
        + "{wildcards.gen}-{wildcards.fit}/"
        + "{wildcards.method}/"
        " {lb}"
        " {ub}"
        " {step}"
        " {output}"

rule optimize_hp:
    input:
        data = config["dir"] + "data/data.{gen}.rds",
        tu_lambda = config["dir"] + "data/tu.lambda.rds"
    output:
        config["dir"]
        + "optim/{gen}-{fit}/{method}/"
        + "phi{phi1}_{phi2}_{phi3}/fit.rds",
    shell:
        "module add gcc/11.2.0; "
        "module add r/4.1.3; "
        "Rscript optimize_hp.R"
        " {input.data}"
        " {seed}"
        " {wildcards.phi1}"
        " {wildcards.phi2}"
        " {wildcards.phi3}"
        " {wildcards.method}"
        " {wildcards.fit}"
        " {input.tu_lambda}"
        " {output}"
        
rule get_tu_lambda:
    input:
        config["dir"] + "data/data.ranef.rds"
    output:
        config["dir"] + "data/tu.lambda.rds"
    shell:
        "module add gcc/11.2.0; "
        "module add r/4.1.3; "
        "Rscript get_tu_lambda.R"
        " {input}"
        " NA" # kinship matrix file name
        " {output}"
        
rule make_data:        
    output:
        config["dir"] + "data/data.{gen}.rds",
    shell:
        "module add gcc/11.2.0; "
        "module add r/4.1.3; "
        "Rscript make_data.R"
        " {wildcards.gen}"
        " {output}"
```
