## Supplemental File 2 for "Probabilistic classification of gene-by-treatment interactions on molecular count phenotypes": subset_data.R

```
suppressPackageStartupMessages(library(tidyverse))

args <- commandArgs(TRUE)
in.file <- as.character(args[1])
n.sim <- as.integer(args[2])
out.file <- as.character(args[3])

data.list <- in.file %>% readRDS

# take the first n.sim simulations for each model
subset <- rep(seq(0, 7, 1) * 1e3, each=n.sim) + seq_len(n.sim)
subset.list <- data.list[subset]

saveRDS(subset.list, file=out.file)
```
