## Supplemental File 2 for "Probabilistic classification of gene-by-treatment interactions on molecular count phenotypes": assemble.sh

```
#!/bin/bash

#SBATCH --job-name=snake
#SBATCH --time=1:00:00
#SBATCH --mem=2G

cd ../figures/vis

mkdir -p main-figs

cp sim_rocc_vln.pdf main-figs/
cp wntrna_gp.pdf main-figs/
cp wntrna_gp_co.pdf main-figs/

mkdir -p supp-figs

cp sim/map-lap/calib.pdf supp-figs/sim_calib_map_lap.pdf
cp sim/mcmc-bs/calib.pdf supp-figs/sim_calib_mcmc_bs.pdf
cp sim/map-lap/rocc.pdf supp-figs/sim_rocc_map_lap.pdf
cp sim/mcmc-bs/rocc.pdf supp-figs/sim_rocc_mcmc_bs.pdf

cp sim/map-lap/lm/hist.pdf supp-figs/sim_hist_map_lap_lm.pdf
cp sim/mcmc-bs/lm/hist.pdf supp-figs/sim_hist_mcmc_bs_lm.pdf
cp sim/map-lap/nl/hist.pdf supp-figs/sim_hist_map_lap_nl.pdf
cp sim/mcmc-bs/nl/hist.pdf supp-figs/sim_hist_mcmc_bs_nl.pdf
cp sim/map-lap/rint/hist.pdf supp-figs/sim_hist_map_lap_rint.pdf
cp sim/mcmc-bs/rint/hist.pdf supp-figs/sim_hist_mcmc_bs_rint.pdf

cp sim/map-lap/vln1.pdf supp-figs/sim_vln_map_lap1.pdf
cp sim/map-lap/vln2.pdf supp-figs/sim_vln_map_lap2.pdf
cp sim/mcmc-bs/vln1.pdf supp-figs/sim_vln_mcmc_bs1.pdf
cp sim/mcmc-bs/vln2.pdf supp-figs/sim_vln_mcmc_bs2.pdf
cp sim/map-lap/vln_aggr.pdf supp-figs/sim_vln_aggr_map_lap.pdf
cp sim/mcmc-bs/vln_aggr.pdf supp-figs/sim_vln_aggr_mcmc_bs.pdf

cp ../sim/map-lap/noranef-noranef/vln_effect.pdf supp-figs/sim_vln_effect_map_lap_noranef_noranef.pdf
cp ../sim/map-lap/noranef-ranef/vln_effect.pdf supp-figs/sim_vln_effect_map_lap_noranef_ranef.pdf
cp ../sim/map-lap/ranef-noranef/vln_effect.pdf supp-figs/sim_vln_effect_map_lap_ranef_noranef.pdf
cp ../sim/map-lap/ranef-ranef/vln_effect.pdf supp-figs/sim_vln_effect_map_lap_ranef_ranef.pdf
cp ../sim/mcmc-bs/noranef-noranef/vln_effect.pdf supp-figs/sim_vln_effect_mcmc_bs_noranef_noranef.pdf
cp ../sim/mcmc-bs/noranef-ranef/vln_effect.pdf supp-figs/sim_vln_effect_mcmc_bs_noranef_ranef.pdf
cp ../sim/mcmc-bs/ranef-noranef/vln_effect.pdf supp-figs/sim_vln_effect_mcmc_bs_ranef_noranef.pdf
cp ../sim/mcmc-bs/ranef-ranef/vln_effect.pdf supp-figs/sim_vln_effect_mcmc_bs_ranef_ranef.pdf

cp sim_compare_sum_lnpy.pdf supp-figs/
cp sim_compare_pp.pdf supp-figs/

cp wnt_llk.pdf supp-figs/
cp wntrna_compare_pp.pdf supp-figs/ 
cp wntrna_hm.pdf supp-figs/ 
cp wntatac_hm.pdf supp-figs/ 
cp wntatac_gp.pdf supp-figs/
cp wntatac_gp_co.pdf supp-figs/
```
