## Supplemental File 2 for "Probabilistic classification of gene-by-treatment interactions on molecular count phenotypes": config.yaml

```
sim_dir: "../derived_data/sim/"
wntrna_dir: "../derived_data/wntrna/"
wntatac_dir: "../derived_data/wntatac/"
dir: "../derived_data/vis/"
fig_dir: "../figures/vis/"
pc: [10]
num_wntrna_pair: 98
num_wntatac_pair: 1775
size: 10
```
