## Supplemental File 2 for "Probabilistic classification of gene-by-treatment interactions on molecular count phenotypes": plot_compare_pp.R

```
suppressPackageStartupMessages(library(tidyverse))
suppressPackageStartupMessages(library(classifygxt))

in.dir1 <- "../derived_data/wntrna/post/kin/pc10"
in.dir2 <- "../derived_data/wntrna/post/id/pc10"
fig.file <- "../figures/vis/wntrna_compare_pp.pdf"

# read in data
method.level <- c("nl", "lm", "rint")
file.vec1 <- file.path(in.dir1, method.level, "pp.rds")
file.vec2 <- file.path(in.dir2, method.level, "pp.rds")

df.list <- vector("list", length(file.vec1)) 

for (i in seq_along(file.vec1)) {

    vec1 <- file.vec1[i] %>%
        readRDS %>%
        as.matrix %>%
        c
    vec2 <- file.vec2[i] %>%
        readRDS %>%
        as.matrix %>%
        c
    
    d <- data.frame(
        val1=vec1,
        val2=vec2,
        method=method.level[i])
    df.list[[i]] <- d
        
}

df <- df.list %>%
    bind_rows %>%
    mutate(method=factor(method, levels=method.level))

# plot
map.method <- c("log-NL", "log-LM", "RINT-LM") %>%
    `names<-`(method.level)

p <- ggplot() +
    geom_point(
        mapping=aes(x=val1, y=val2),
        data=df,
        size=0.01)
p <- p +
    xlab("Polygenic") +
    ylab("Donor") 
p <- p + scale_x_continuous(
             limits=c(0, 1),
             breaks=c(0, 0.5, 1),
             labels=c("0.0", "0.5", "1.0"))
p <- p + scale_y_continuous(
             limits=c(0, 1),
             breaks=c(0, 0.5, 1),
             labels=c("0.0", "0.5", "1.0"))
p <- p +
    facet_wrap(
        vars(method),
        labeller=labeller(method=map.method))
p <- p + coord_equal()

ggsave(filename=fig.file, plot=p, width=7, height=3.5)
```
