## Supplemental File 2 for "Probabilistic classification of gene-by-treatment interactions on molecular count phenotypes": plot_hm.R

```
suppressPackageStartupMessages(library(tidyverse))
suppressPackageStartupMessages(library(hrbrthemes))
suppressPackageStartupMessages(library(viridis))
suppressPackageStartupMessages(library(patchwork))
suppressPackageStartupMessages(library(classifygxt))

# define function
plot_hm <- function(m) {
    pp.vec <- m %>% as.matrix %>% c
    pair <- m %>% nrow %>% seq_len
    model <- colnames(m)
    d <- expand.grid(X=pair, Y=model) %>%
        `colnames<-`(c("pair", "model")) %>%
        mutate(pp=pp.vec)

    p <- d %>%
        ggplot(aes(x=pair, y=model, fill=pp)) +
        geom_tile()

    p <- p + scale_fill_viridis(
                 discrete=FALSE,
                 limits=c(0, 1),
                 breaks=c(0, 0.5, 1),
                 labels=c("0.0", "0.5", "1.0"))
    p <- p + scale_y_discrete(limits=rev)
    p <- p + labs(y="Model", x="Feature-SNP pairs")
    p <- p + theme(axis.ticks=element_blank())
    p <- p + theme(axis.line=element_blank())
    p <- p + theme(
                 axis.text.y=element_text(
                     hjust=1, margin=margin(r=-15), size=rel(1)))
    p <- p + theme(axis.text.x=element_blank())
    p <- p + theme(
        panel.grid.major=element_blank(),
        panel.grid.minor=element_blank(),
        panel.border=element_blank(),
        panel.background=element_blank()) 
    p <- p + theme(panel.margin.x=unit(0, "cm"))
    p <- p + theme(panel.margin.y=unit(0, "cm"))
    p <- p + theme(legend.position="none")

    p
}

nosign.file <- "../derived_data/wntrna/optim/kin/pc10/nl/pp.rds"
sign.file <- "../derived_data/wntrna/optim/kin/pc10/nl/pp.sign.rds"
order.file <- "../derived_data/wntrna/optim/kin/pc10/nl/order.vec.rds"
out.file <- "../figures/vis/wntrna_hm.pdf"

file.exists(order.file)

args <- commandArgs(TRUE)
nosign.file <- as.character(args[1])
sign.file <- as.character(args[2])
order.file <- as.character(args[3])
out.file <- as.character(args[4])

nosign <- nosign.file %>% readRDS
sign <- sign.file %>% readRDS
order.vec <- order.file %>% readRDS

nosign <- nosign[order.vec, ]
sign <- sign[order.vec, ]

nosign.plot <- nosign %>% plot_hm
sign.plot <- sign %>% plot_hm

(nosign.plot / sign.plot) +
    plot_layout(
        guides="collect",
        height=c(8, 27)) &
    plot_annotation(tag_levels="A") &
    theme(
        legend.position="bottom",
        legend.title=element_blank(),
        legend.text=element_text(size=rel(0.8)),
        legend.key.size=unit(0.4, "cm"),
        legend.key.width=unit(0.6, "cm"),
        legend.margin=margin(0, 0, 0, 0),
        legend.box.margin=margin(-6, 0, 0, 0))

ggsave(filename=out.file, width=7, height=7)
```
