## Supplemental File 2 for "Probabilistic classification of gene-by-treatment interactions on molecular count phenotypes": plot_sim_rocc_vln.R

```
suppressPackageStartupMessages(library(tidyverse))
suppressPackageStartupMessages(library(patchwork))

cols <- c(rgb(230/255, 159/255, 0/255),
          rgb(86/255, 180/255, 233/255),
          rgb(0/255, 158/255, 115/255))

# rocc.file <- "../derived_data/sim/mcmc-bs/noranef-noranef/rocc.rds"
# aggr.file <- "../derived_data/sim/mcmc-bs/noranef-noranef/vln_aggr.rds"
# co.file <- "../derived_data/sim/post/noranef-noranef/vln.co.rds"
# fig.file <- "../figures/vis/sim_rocc_vln.pdf"

args <- commandArgs(TRUE)
rocc.file <- args[1]
aggr.file <- args[2]
co.file <- args[3]
fig.file <- args[4]

roc <- rocc.file %>% readRDS
aggr <- aggr.file %>% readRDS
co <- co.file %>% readRDS

co <- co %>%
    mutate(cat=factor("Crossover"))

# rocc
p <- ggplot() +
    geom_line(
        mapping=aes(x=fpr, y=recall, colour=method),
        data=roc)
p <- p + geom_abline(intercept=0, slope=1, linetype=2)
p <- p + xlab("FPR") + ylab("TPR")
p <- p + scale_color_manual(values=cols)
p <- p + scale_x_continuous(
             breaks=c(0, 0.5, 1),
             labels=c("0.0", "0.5", "1.0"))
p <- p + scale_y_continuous(
             breaks=c(0, 0.5, 1),
             labels=c("0.0", "0.5", "1.0"))
p <- p + theme(aspect.ratio=1)
p <- p + theme(legend.position="right")
p <- p + theme(legend.box.spacing=unit(2, "pt"))
p <- p + theme(legend.title=element_blank())
p <- p + theme(legend.key=element_rect(
                   colour="transparent", fill="transparent"))
p <- p + theme(panel.grid.major=element_line(
                   colour="gray", linewidth=rel(0.6)))
p <- p + theme(panel.grid.minor=element_line(
                   colour="gray", linewidth=rel(0.6)))
p <- p + theme(panel.background=element_blank())
p <- p + theme(panel.border=element_rect(
                   colour="gray", fill=NA, linewidth=1))
p <- p + facet_wrap(vars(cat))

# aggr
colnames(aggr) <- c("method", "cat", "value", "type")
map.correct <- c("Correct", "Incorrect") %>%
    `names<-`(c("1", "0"))
map.cat <- c("No GxT", "Induced", "Altered") %>%
    `names<-`(levels(aggr$cat))

p1 <- aggr %>%
    ggplot(aes(x=method, y=value, color=method)) +
    geom_violin(scale="width") +
    stat_summary(
        fun=median, geom="point", size=1) +
    ylim(0, 1) +
    ylab("Posterior probability") +
    facet_grid(
        vars(type), vars(cat),
        labeller=labeller(
            type=map.correct,
            cat=map.cat),
        scales="free",
        switch="y")
p1 <- p1 + scale_y_continuous(
             limits=c(0, 1),
             breaks=c(0, 0.5, 1),
             labels=c("0.0", "0.5", "1.0"),
             position="right")
p1 <- p1 + scale_color_manual(
               values=cols,
               labels=c("log-NL", "log-LM", "RINT-LM"))
p1 <- p1 + theme(legend.position="none")
p1 <- p1 + theme(legend.title=element_blank())
p1 <- p1 + theme(panel.grid.major=element_line(
                   colour="gray", linewidth=rel(0.6)))
p1 <- p1 + theme(panel.grid.minor=element_line(
                   colour="gray", linewidth=rel(0.6)))
p1 <- p1 + theme(panel.background=element_blank())
p1 <- p1 + theme(panel.border=element_rect(
                   colour="gray", fill=NA, linewidth=1))
p1 <- p1 + theme(axis.ticks.x=element_blank())
p1 <- p1 + theme(axis.text.x=element_blank())
p1 <- p1 + theme(axis.title.x=element_blank())

# co
lab.truth <- c("Correct", "Incorrect") %>%
    `names<-`(c("1", "0"))
lab <- labeller(truth=lab.truth)
p2 <- co %>% ggplot(aes(x=name, y=value, color=name)) +
    geom_violin(scale="width") +
    stat_summary(
        fun=median, geom="point", size=1) +
    ylim(0, 1) +
    ylab("Posterior probability") +
    facet_grid(
        vars(truth), vars(cat),
        labeller=lab,
        scales="free",
        switch="y")
p2 <- p2 + scale_y_continuous(
             limits=c(0, 1),  
             breaks=c(0, 0.5, 1),
             labels=c("0.0", "0.5", "1.0"),
             position="right")
p2 <- p2 + scale_color_manual(values=cols)
p2 <- p2 + theme(legend.position="none")
p2 <- p2 + theme(legend.title=element_blank())
p2 <- p2 + theme(panel.grid.major=element_line(
                   colour="gray", linewidth=rel(0.6)))
p2 <- p2 + theme(panel.grid.minor=element_line(
                   colour="gray", linewidth=rel(0.6)))
p2 <- p2 + theme(panel.background=element_blank())
p2 <- p2 + theme(panel.border=element_rect(
                   colour="gray", fill=NA, linewidth=1))
p2 <- p2 + theme(axis.ticks.x=element_blank())
p2 <- p2 + theme(axis.text.x=element_blank())
p2 <- p2 + theme(axis.title.x=element_blank())

p2 <- p2 + theme(axis.title.y=element_text(vjust=+3))

# assemble plots
layout <- "
    AAAA
    BBBC
"

wrap_plots(
    A=p + theme(legend.position=c(1.11, 0.5),
                legend.background=element_rect(
                    colour="transparent",
                    fill="transparent"),
                legend.key=element_rect(
                    colour="transparent",
                    fill="transparent")),
    B=p1 + theme(legend.position="bottom"),
    C=p2 + theme(legend.position="none"),
    design=layout) +
    plot_layout(
        width=c(1, 1, 1, 1),
        height=c(1, 2.5)) &
    plot_annotation(tag_levels="A") # &

ggsave(filename=fig.file, width=7, height=7.5)
```
