## Supplemental File 2 for "Probabilistic classification of gene-by-treatment interactions on molecular count phenotypes": plot_wnt_llk.R

```
suppressPackageStartupMessages(library(tidyverse))
suppressPackageStartupMessages(library(patchwork))

fig.file <- "../figures/vis/wnt_llk.pdf"
fig.dir <- "../figures/vis"

file1 <- "../derived_data/wntrna/aa/llk/pc10/Veh/llk.rds"
file2 <- "../derived_data/wntrna/aa/llk/pc10/CT99/llk.rds"
file3 <- "../derived_data/wntatac/aa/llk/pc10/Veh/llk.rds"
file4 <- "../derived_data/wntatac/aa/llk/pc10/CT99/llk.rds"

file.vec <- c(file1, file2, file3, file4) 
p.list <- vector("list", 12)

for (i in seq_len(4)) {

    file <- file.vec[i]
    d <- file %>% readRDS
    dl <- d %>%
        pivot_longer(everything())

    p1 <- d %>% 
        ggplot(aes(x=lm, y=nl)) +
        geom_point(size=0.25, alpha=0.2) +
        xlab("log-LM") +
        ylab("log-NL") +
        coord_fixed() +
        theme_bw()

    p2 <- d %>% 
        ggplot(aes(x=nl, y=anova)) +
        geom_point(size=0.25, alpha=0.2) +
        xlab("log-NL") +
        ylab("log-ANOVA") +
        coord_fixed() +
        theme_bw()

    p3 <- d %>% 
        ggplot(aes(x=anova, y=lm)) +
        geom_point(size=0.25, alpha=0.2) +
        xlab("log-ANOVA") +
        ylab("log-LM") +
        coord_fixed() +
        theme_bw()

    p.list[[(i - 1) * 3 + 1]] <- p1
    p.list[[(i - 1) * 3 + 2]] <- p2
    p.list[[(i - 1) * 3 + 3]] <- p3
    
}
    
# pw + plot_annotation(tag_levels="A")
((p.list[[1]] + p.list[[2]] + p.list[[3]]) /
    (p.list[[4]] + p.list[[5]] + p.list[[6]]) /
    (p.list[[7]] + p.list[[8]] + p.list[[9]]) /
    (p.list[[10]] + p.list[[11]] + p.list[[12]])) +
    plot_annotation(tag_levels="A") &
    theme(plot.tag=element_text(size=rel(1.2)))

ggsave(filename=fig.file, width=8.1, height=10.8)
```
