## Supplemental File 2 for "Probabilistic classification of gene-by-treatment interactions on molecular count phenotypes": plot_wntatac_co.R

```
suppressPackageStartupMessages(library(tidyverse))
suppressPackageStartupMessages(library(patchwork))
suppressPackageStartupMessages(library(classifygxt))

source("utils.R")

data.file <- "../derived_data/wntatac/pre/rsd/pc10/data.list.rds"
in.dir <- "../derived_data/wntatac/post/pc10/nl/fit"
fig.dir <- "../figures/vis"
fig.file <- file.path(fig.dir, "wntatac_gp_co.pdf")

if (!dir.exists(fig.dir)) dir.create(fig.dir, recursive=TRUE)

# read in the input data
data.list <- data.file %>% readRDS

# change the format
data.list <- data.list %>%
    map(function(x)
        list(
            y=x$data$y, g=x$data$g,
            t=x$data$t, subject=x$data$dna,
            feat.id=x$feat.id, snp.id=x$snp.id))

feat.vec <- data.list %>%
    map_chr(pluck("feat.id"))
snp.vec <- data.list %>%
    map_chr(pluck("snp.id"))

feat.name <- c(
    "chr18_7584881-7585200",
    "chr5_151624011-151625560",
    # "chr16_85293831-85294460",
    "chr14_68286771-68286930")
snp.name <- c(
    "chr18:7585131:C:G",  
    "chr5:151624586:G:C",
    # "chr16:85293299:C:A",
    "chr14:68286876:C:T") 
rs.vec <- c(
    "rs4798573",
    "rs17718084",
    # "rs2966860",
    "rs911263")    
geno.vec <- c("0|0", "0|0", "1|1")

# all(match(feat.name, feat.vec) ==
#     match(snp.name, snp.vec))
index.vec <- match(feat.name, feat.vec)

gp.list <- pp.list <-
    vector("list", length(index.vec))

for (i in seq_along(index.vec)) {
    
    # get index for the i-th rank
    index <- index.vec[i]

    # get info
    feat.id <- data.list[[index]]$feat.id
    feat.id <- feat.id %>%
        gsub("_", ":\n", .)
    snp.id <- data.list[[index]]$snp.id
    rs.id <- rs.vec[i]
    title <- paste0(feat.id, "\n", rs.id)
    geno <- geno.vec[i]
    geno.label <- get_geno_label(
        snp.id=snp.id, geno=geno)

    fit.file <- file.path(
        in.dir,
        paste0(
            "res", str_pad(index, 4, pad="0"), ".rds"))
    fit <- fit.file %>% readRDS

    # for gp plots
    data <- data.list[[index]]
    gp <- format_gp(data=data, fit=fit)

    # create a ggplot2 object
    p1 <- make_gp_plot(gp=gp)
    p1 <- p1 + scale_x_continuous(
                   labels=geno.label,
                   breaks=c(0, 1, 2))
    p1 <- p1 + labs(title=title) +
        theme(plot.title=element_text(size=rel(0.9)))
    
    # for pp plots
    pp <- fit %>% format_pp(co=TRUE)
    p2 <- pp %>% make_pp_plot
    
    gp.list[[i]] <- p1
    pp.list[[i]] <- p2
    
}

((gp.list[[1]] / pp.list[[1]]) |
 (gp.list[[2]] / pp.list[[2]]) |
 (gp.list[[3]] / pp.list[[3]])) +
    plot_layout(guides="collect") &
    plot_annotation(tag_levels="A") &
    theme(
        legend.position="bottom",
        legend.title=element_blank(),
        legend.text=element_text(size=rel(1)))

ggsave(file=fig.file, width=8, height=6)
```
