## Supplemental File 2 for "Probabilistic classification of gene-by-treatment interactions on molecular count phenotypes": plot_wntatac_gp.R

```
suppressPackageStartupMessages(library(tidyverse))
suppressPackageStartupMessages(library(patchwork))
suppressPackageStartupMessages(library(classifygxt))

source("utils.R")

data.file <- "../derived_data/wntatac/pre/rsd/pc10/data.list.rds"
in.dir <- "../derived_data/wntatac/post/pc10"
fig.dir <- "../figures/vis"
fig.file <- file.path(fig.dir, "wntatac_gp.pdf")

if (!dir.exists(fig.dir)) dir.create(fig.dir, recursive=TRUE)

# read in the input data
data.list <- data.file %>% readRDS

# change the format
data.list <- data.list %>%
    map(function(x)
        list(
            y=x$data$y, g=x$data$g,
            t=x$data$t, subject=x$data$dna,
            feat.id=x$feat.id, snp.id=x$snp.id))

feat.vec <- data.list %>%
    map_chr(pluck("feat.id"))
snp.vec <- data.list %>%
    map_chr(pluck("snp.id"))

feat.name <- c(
    # "chr7_89914521-89915220", # unstable
    "chr5_102554041-102554990",
    # "chr20_4716511-4717010",
    "chr17_82118351-82119190",
    "chr2_120613861-120614840", # induced
    "chr8_4046751-4047130")
snp.name <- c(
    # "chr7:89914871:C:T", # unstable
    "chr5:102554238:C:T",
    # "chr20:4716775:A:G",
    "chr17:82118932:C:G",
    "chr2:120614301:A:G", # induced
    "chr8:4067761:A:G")
rs.vec <- c(
    # "rs10259722",
    "rs1875632",
    # "rs6052791",
    "rs7218075",
    "rs11886111", # induced
    "rs1035637")    
# geno.vec <- c("0|0", "0|0", "0|0", "1|1")
geno.vec <- c("1|1", "0|0", "1|1", "1|1")

# all(match(feat.name, feat.vec) ==
#     match(snp.name, snp.vec))
index.vec <- match(feat.name, feat.vec)

method.vec <- c("nl", "lm", "rint")
method.name <- c("log-NL", "log-LM", "RINT-LM")
method.fact <- factor(method.name, levels=method.name)

gp.list <- pp.list <-
    vector("list", length(index.vec))

for (i in seq_along(index.vec)) {
    
    # get index for the i-th rank
    index <- index.vec[i]

    # get info
    feat.id <- data.list[[index]]$feat.id %>%
        gsub("_", ":", .)
    snp.id <- data.list[[index]]$snp.id
    rs.id <- rs.vec[i]
    title <- paste0(feat.id, "\n", rs.id)
    geno <- geno.vec[i]
    geno.label <- get_geno_label(
        snp.id=snp.id, geno=geno)
    
    file.vec <- file.path(
        in.dir, method.vec, "fit",
        paste0(
            "res", str_pad(index, 4, pad="0"), ".rds"))
    
    res.list <- file.vec %>% map(readRDS)

    # for gp plots
    data <- data.list[[index]]
    n.sample <- length(data$y)
    
    # for arrows
    # if (i %in% 1:2) {
    #     y <- data$y
    #     g <- data$g
    #     t <- data$t
    #     y1 <- min(y[g == 2 & t == 0])
    #     y2 <- min(y[g == 2 & t == 1])
    #     arrow <- data.frame(
    #         y.end=c(y1, y2)) %>%
    #         mutate(method=factor("log-LM",
    #                              levels=method.name))
    # }    
    
    plot.list <- res.list %>%
        map(function(x) format_gp(data=data, fit=x))

    input.list <- plot.list %>%
        map(pluck("input"))
    n.sample <- input.list[[1]] %>% nrow
    input.df <- input.list %>%
        bind_rows %>%
        mutate(method=rep(
                   method.name, each=n.sample)) %>%
        mutate(method=factor(method, levels=method.name))
    
    fit.list <- plot.list %>%
        map(pluck("fit"))
    n.pt <- fit.list[[1]] %>% nrow
    fit.df <- fit.list %>%
        bind_rows %>%
        mutate(method=rep(
                   method.name, each=n.pt)) %>%
        mutate(method=factor(method, levels=method.name))
    
    # create a ggplot2 object
    p1 <- list(input=input.df, fit=fit.df) %>%
        make_gp_plot
    p1 <- p1 + scale_x_continuous(
                   labels=geno.label,
                   breaks=c(0, 1, 2))
    p1 <- p1 + labs(title=title)
    p1 <- p1 + theme(plot.title=element_text(size=rel(1)))
    p1 <- p1 + facet_wrap(vars(method), scales="free_y")
    
    # if (i %in% 1:2) {
    #     p1 <- p1 +
    #         geom_segment(
    #             mapping=aes(x=1.65, xend=1.85,
    #                 y=y.end - 0.6, yend=y.end),
    #             data=arrow,
    #             arrow=arrow(
    #                 length=unit(0.2, "cm"),
    #                 type="closed"),
    #             inherit.aes=FALSE)
    # }    
    
    # for pp plots
    pp <- res.list %>%
        map(format_pp) %>%
        bind_rows %>%
        mutate(method=rep(
                   method.name, each=8)) %>%
        mutate(method=factor(method, levels=method.name))
    
    p2 <- pp %>% make_pp_plot
    p2 <- p2 +
        facet_wrap(vars(method), scales="free_y")
    
    gp.list[[i]] <- p1
    pp.list[[i]] <- p2
    
}

# assemble plots
layout <- "
    AACC
    BBDD
    EEGG
    FFHH
"

wrap_plots(
    A=gp.list[[1]], B=pp.list[[1]],
    C=gp.list[[2]], D=pp.list[[2]],
    E=gp.list[[3]], F=pp.list[[3]],
    G=gp.list[[4]], H=pp.list[[4]],
    design=layout) +
    plot_layout(guides="collect") &
    plot_annotation(tag_levels="A") &
    theme(
        legend.position="bottom",
        legend.title=element_blank(),
        legend.text=element_text(size=rel(1)))

ggsave(file=fig.file, width=12, height=11)
```
