## Supplemental File 2 for "Probabilistic classification of gene-by-treatment interactions on molecular count phenotypes": pw_sim_calib.R

```
suppressPackageStartupMessages(library(tidyverse))
suppressPackageStartupMessages(library(patchwork))

args <- commandArgs(TRUE)
in.dir <- as.character(args[1])
fig.dir <- as.character(args[2])
fig.file <- as.character(args[3])

if (!dir.exists(fig.dir)) dir.create(fig.dir, recursive=TRUE)

type.vec <- c(
    "noranef-noranef", "noranef-ranef",
    "ranef-noranef", "ranef-ranef")
file.vec <- file.path(
    in.dir, type.vec, "calib.rds")

# create a list to store ggplot2 objects
gg.list <- vector("list", length(type.vec))

for (i in seq_along(gg.list)) {

    plot.list <- file.vec[i] %>% readRDS
    d <- plot.list %>% pluck("data")
    se <- plot.list %>% pluck("se")

    p <- ggplot() +
        geom_point(
            mapping=aes(x=x, y=y),
            data=d)
    p <- p +
        geom_segment(
            mapping=aes(x=x, y=y1, xend=x, yend=y2),
            data=se)
    p <- p +
        geom_abline(
            intercept=0, slope=1, colour="dodgerblue")
    # p <- p + xlim(0, 1) + ylim(0, 1)
    p <- p +
        xlab("Posterior probability") +
        ylab("Fraction of positives")
    p <- p + scale_x_continuous(
                 limits=c(0, 1),
                 breaks=c(0, 0.5, 1),
                 labels=c("0.0", "0.5", "1.0"))
    p <- p + scale_y_continuous(
                 limits=c(0, 1),
                 breaks=c(0, 0.5, 1),
                 labels=c("0.0", "0.5", "1.0"))
    # p <- p + theme(aspect.ratio=1)
    p <- p + theme(legend.position="none")
    # p <- p + theme(panel.grid.major=element_blank())
    # p <- p + theme(panel.grid.minor=element_blank())
    p <- p + theme(panel.grid.major=element_line(
                       colour="gray", linewidth=rel(0.6)))
    p <- p + theme(panel.grid.minor=element_line(
                       colour="gray", linewidth=rel(0.6)))
    p <- p + theme(panel.background=element_blank())
    p <- p + theme(panel.border=element_rect(
                       colour="gray", fill=NA, linewidth=1))
    p <- p + facet_grid(cat ~ method)
    p <- p + coord_equal(xlim=c(0, 1), ylim=c(0, 1))

    gg.list[[i]] <- p
    
}

(gg.list[[1]] | gg.list[[2]]) /
    (gg.list[[3]] | gg.list[[4]]) &
    plot_annotation(tag_levels="A")
    
ggsave(filename=fig.file, width=11, height=11)
```
