## Supplemental File 2 for "Probabilistic classification of gene-by-treatment interactions on molecular count phenotypes": pw_sim_compare_pp.R

```
suppressPackageStartupMessages(library(tidyverse))
suppressPackageStartupMessages(library(patchwork))
suppressPackageStartupMessages(library(classifygxt))

fig.file <- "../figures/vis/sim_compare_pp.pdf"
fig.dir <- "../figures/vis"

if (!dir.exists(fig.dir)) dir.create(fig.dir, recursive=TRUE)

type.vec <- c(
    "noranef-noranef", "noranef-ranef",
    "ranef-noranef", "ranef-ranef")
file.vec <- file.path(
    "../derived_data/sim/comparison",
    type.vec, "pp.rds")

method.level <- c("nl", "lm", "rint")
hp.level <- c("rest", "opt", "perm")
map.method <- c("log-NL", "log-LM", "RINT-LM") %>%
    `names<-`(method.level)
map.hp <- c("Restrictive", "Optimal", "Permissive") %>%
    `names<-`(hp.level)

# read in the data
df.list <- file.vec %>% map(readRDS)

p.list <- vector("list", length(type.vec))

# iterate through the list
for (i in seq_along(df.list)) {

    df <- df.list[[i]]
    p <- ggplot() +
        geom_point(
            mapping=aes(x=mcmc.bs, y=map.lap),
            data=df,
            size=0.01)
    p <- p +
        xlab("MCMC & bridge sampling") +
        ylab("MAP & Laplace")
    p <- p +
        scale_x_continuous(
            breaks=c(0, 0.5, 1),
            labels=c("0.0", "0.5", "1.0"))
    p <- p +
        scale_y_continuous(
            breaks=c(0, 0.5, 1),
            labels=c("0.0", "0.5", "1.0"))
    p <- p +
        facet_grid(
            vars(method), vars(hp),
            labeller=labeller(
                method=map.method,
                hp=map.hp))
    p <- p + coord_equal()
    # p <- p + coord_equal(xlim=c(0, 1), ylim=c(0, 1))

    p.list[[i]] <- p

}

pw <- (p.list[[1]] | p.list[[2]]) /
    (p.list[[3]] | p.list[[4]]) &
    plot_annotation(tag_levels="A")
    
ggsave(filename=fig.file, plot=pw, width=11, height=11)
    
# ggsave(filename=fig.file, width=12, height=12)
```
