## Supplemental File 2 for "Probabilistic classification of gene-by-treatment interactions on molecular count phenotypes": pw_sim_rocc.R

```
suppressPackageStartupMessages(library(tidyverse))
suppressPackageStartupMessages(library(patchwork))

args <- commandArgs(TRUE)
in.dir <- as.character(args[1])
fig.dir <- as.character(args[2])
fig.file <- as.character(args[3])

if (!dir.exists(fig.dir)) dir.create(fig.dir, recursive=TRUE)

type.vec <- c(
    "noranef-noranef", "noranef-ranef",
    "ranef-noranef", "ranef-ranef")
file.vec <- file.path(
    in.dir, type.vec, "rocc.rds")

d.list <- file.vec %>% map(readRDS)

# plot
plot_rocc <- function(rocc) {

    cols <- c(rgb(230/255, 159/255, 0/255),
          rgb(86/255, 180/255, 233/255),
          rgb(0/255, 158/255, 115/255))

    p <- ggplot() +
        geom_line(
            mapping=aes(x=fpr, y=recall, colour=method),
            data=rocc)
    p <- p + geom_abline(intercept=0, slope=1, linetype=2)
    p <- p + xlab("FPR") + ylab("TPR")
    # modify scales
    p <- p + scale_color_manual(values=cols)
    # p <- p + scale_x_continuous(
    #              labels=scales::number_format(accuracy=0.1))
    # p <- p + scale_y_continuous(
    #              labels=scales::number_format(accuracy=0.1))
    p <- p + scale_x_continuous(
                 breaks=c(0, 0.5, 1),
                 labels=c("0.0", "0.5", "1.0"))
    p <- p + scale_y_continuous(
                 breaks=c(0, 0.5, 1),
                 labels=c("0.0", "0.5", "1.0"))
    # modify themes
    p <- p + theme(aspect.ratio=1)
    p <- p + theme(legend.position=c(1.12, 0.5))
    p <- p + theme(legend.title=element_blank())
    # p <- p + theme(legend.spacing.y=unit(0.2, "cm"))
    # p <- p + theme(panel.grid.major=element_blank())
    # p <- p + theme(panel.grid.minor=element_blank())
    p <- p + theme(legend.background=element_rect(
                       colour="transparent",
                       fill="transparent"))
    p <- p + theme(legend.key=element_rect(
                       colour="transparent",
                       fill="transparent"))
    p <- p + theme(panel.grid.major=element_line(
                       colour="gray", linewidth=rel(0.6)))
    p <- p + theme(panel.grid.minor=element_line(
                       colour="gray", linewidth=rel(0.6)))
    p <- p + theme(panel.background=element_blank())
    p <- p + theme(panel.border=element_rect(
                       colour="gray", fill=NA, linewidth=1))
    p <- p + facet_wrap(vars(cat))
    p

}

gg.list <- d.list %>% map(plot_rocc)

(gg.list[[1]] / gg.list[[2]] /
    gg.list[[3]] / gg.list[[4]]) &
    plot_annotation(tag_levels="A")

ggsave(filename=fig.file, width=7.5, height=10)
# ggsave(filename=fig.file, width=6, height=10)
```
