## Supplemental File 2 for "Probabilistic classification of gene-by-treatment interactions on molecular count phenotypes": pw_sim_vln.R

```
suppressPackageStartupMessages(library(tidyverse))
suppressPackageStartupMessages(library(classifygxt))
suppressPackageStartupMessages(library(patchwork))

cols <- c(rgb(230/255, 159/255, 0/255),
          rgb(86/255, 180/255, 233/255),
          rgb(0/255, 158/255, 115/255))

in.dir <- "../derived_data/sim/mcmc-bs/"
fig.dir <- "../figures/vis/sim/mcmc-bs/all"
fig.file1 <- "../figures/vis/sim/mcmc-bs/all/vln1.pdf"
fig.file2 <- "../figures/vis/sim/mcmc-bs/all/vln2.pdf"

args <- commandArgs(TRUE)
in.dir <- as.character(args[1])
fig.dir <- as.character(args[2])
fig.file1 <- as.character(args[3])
fig.file2 <- as.character(args[4])

type.vec <- c(
    "noranef-noranef", "noranef-ranef",
    "ranef-noranef", "ranef-ranef")
file.vec <- file.path(
    in.dir, type.vec, "vln.rds")
method.name <- c("log-NL", "log-LM", "RINT-LM")

dl.list <- file.vec %>% map(readRDS)

plot_vln <- function(dl) {

    p <- dl %>%
        ggplot(aes(x=method, y=value, color=method)) +
        geom_violin(scale="width") +
        # geom_boxplot(outlier.shape=NA, width=0.1) +
        stat_summary(
            fun=median, geom="point", size=1) +
        ylim(0, 1) +
        ylab("Posterior probability") +
        facet_grid(
            vars(type), vars(name),
            # labeller=lab,
            scales="free") +
        # theme_classic() +
        theme(axis.ticks.x=element_blank(),
              axis.text.x=element_blank(),
              axis.title.x=element_blank(),
              legend.title=element_blank(),
              legend.position="bottom")
    p <- p + scale_color_manual(
                 values=cols,
                 labels=method.name)
    p <- p + theme(panel.grid.major=element_line(
                       colour="gray", linewidth=rel(0.6)))
    p <- p + theme(panel.grid.minor=element_line(
                       colour="gray", linewidth=rel(0.6)))
    p <- p + theme(panel.background=element_blank())
    p <- p + theme(
                 panel.border=element_rect(
                     colour="gray", fill=NA, linewidth=1))
    p <- p + theme(axis.ticks.x=element_blank())
    p <- p + theme(axis.text.x=element_blank())
    p <- p + theme(axis.title.x=element_blank())

    p

}

p.list <- dl.list %>% map(plot_vln)

(p.list[[1]] / p.list[[2]]) +
    plot_layout(guides="collect") &
    theme(legend.position="bottom") &
    plot_annotation(tag_levels="A")
ggsave(file=fig.file1, width=8, height=8.8)

(p.list[[3]] / p.list[[4]]) +
    plot_layout(guides="collect") &
    theme(legend.position="bottom") &
    plot_annotation(tag_levels="A")
ggsave(file=fig.file2, width=8, height=8.8)
```
