## Supplemental File 2 for "Probabilistic classification of gene-by-treatment interactions on molecular count phenotypes": README.md

# `vis`

This directory contains code for visualization.
The code under this directory takes as input the output from the workflows under the `sim`, `wntrna`, and `wntatac` directories.
The final output files are summarized in the table below, which can be obtained by running `assemble.sh` after the snakemake workflow.

|Figure|File name|
|:-|:-|
|Fig 4|sim_rocc_vln.pdf|
|Fig 5|wntrna_gp_all.pdf|
|Fig 6|wntrna_gp_co.pdf|
|S1 Fig|wnt_llk.png|
|S2 Fig|sim_calib_mcmc_bs.pdf|
|S3 Fig|sim_calib_map_lap.pdf|
|S4 Fig|sim_rocc_mcmc_bs.pdf|
|S5 Fig|sim_rocc_map_lap.pdf|
|S6 Fig|sim_hist_mcmc_bs_nl.pdf|
|S7 Fig|sim_hist_mcmc_bs_lm.pdf|
|S8 Fig|sim_hist_mcmc_bs_rint.pdf|
|S9 Fig|sim_hist_map_lap_nl.pdf|
|S10 Fig|sim_hist_map_lap_lm.pdf|
|S11 Fig|sim_hist_map_lap_rint.pdf|
|S12 Fig|sim_vln_mcmc_bs1.pdf|
|S13 Fig|sim_vln_mcmc_bs2.pdf|
|S14 Fig|sim_vln_map_lap1.pdf|
|S15 Fig|sim_vln_map_lap2.pdf|
|S16 Fig|sim_vln_aggr_mcmc_bs.pdf|
|S17 Fig|sim_vln_aggr_map_lap.pdf|
|S18 Fig|sim_vln_effect_mcmc_bs_noranef_noranef.pdf|
|S19 Fig|sim_vln_effect_mcmc_bs_noranef_ranef.pdf|
|S20 Fig|sim_vln_effect_mcmc_bs_ranef_noranef.pdf|
|S21 Fig|sim_vln_effect_mcmc_bs_ranef_ranef.pdf|
|S22 Fig|sim_vln_effect_map_lap_noranef_noranef.pdf|
|S23 Fig|sim_vln_effect_map_lap_noranef_ranef.pdf|
|S24 Fig|sim_vln_effect_map_lap_ranef_noranef.pdf|
|S25 Fig|sim_vln_effect_map_lap_ranef_ranef.pdf|
|S26 Fig|sim_compare_sum_lnpy.pdf|
|S27 Fig|sim_compare_pp.png|
|S28 Fig|wntrna_hm.pdf|
|S29 Fig|wntatac_hm.pdf|
|S30 Fig|wntatac_gp_all.pdf|
|S31 Fig|wntatac_gp_co.pdf|
|S32 Fig|wntrna_compare_pp.pdf|

<!-- int.pdf
gxt.pdf
additivity2.pdf -->
