## Supplemental File 2 for "Probabilistic classification of gene-by-treatment interactions on molecular count phenotypes": Snakefile

```
pc = config["pc"]
num_wntrna_pair = config["num_wntrna_pair"]
num_wntatac_pair = config["num_wntatac_pair"]
size = config["size"]

method = ["nl", "lm", "rint"]
gen = ["noranef", "ranef"]
fit = ["noranef", "ranef"]
cat = ["nogxt", "induced", "altered"]
hp = ["rest", "opt", "perm"]
comp = ["map-lap", "mcmc-bs"]
cond = ["Veh", "CT99"]

wntrna_start = [format(number, "04")
                for number in range(1, num_wntrna_pair + 1, size)]
wntatac_start = [format(number, "04")
                 for number in range(1, num_wntatac_pair + 1, size)]

rule all:
    input:
        config["fig_dir"] + "sim_rocc_vln.pdf",
        config["fig_dir"] + "wntrna_gp.pdf",
        config["fig_dir"] + "wntrna_gp_co.pdf",
        config["fig_dir"] + "wnt_llk.pdf",
        expand(config["fig_dir"]
               + "sim/{comp}/{method}/hist.pdf",
               comp = comp, method = method),
        config["fig_dir"] + "sim_compare_pp.pdf",
        config["fig_dir"] + "sim_compare_sum_lnpy.pdf",
        expand(config["fig_dir"]
               + "sim/{comp}/calib.pdf",
               comp = comp),
        expand(config["fig_dir"]
               + "sim/{comp}/rocc.pdf",
               comp = comp),
        expand(config["fig_dir"]
               + "sim/{comp}/vln1.pdf",
               comp = comp),
        expand(config["fig_dir"]
               + "sim/{comp}/vln2.pdf",
               comp = comp),
        expand(config["fig_dir"]
               + "sim/{comp}/vln_aggr.pdf",
               comp = comp),
        config["fig_dir"] + "wntrna_compare_pp.pdf",
        config["fig_dir"] + "wntrna_hm.pdf",
        config["fig_dir"] + "wntatac_gp.pdf",
        config["fig_dir"] + "wntatac_gp_co.pdf",
        config["fig_dir"] + "wntatac_hm.pdf",
        expand(config["fig_dir"]
               + "sim_lnpy_{gen}_{fit}.pdf",
               gen = gen, fit = fit),
        
rule plot_sim_rocc_vln:
    input:
        rocc = config["sim_dir"]
        + "mcmc-bs/noranef-noranef/rocc.rds",
        aggr = config["sim_dir"]
        + "mcmc-bs/noranef-noranef/vln.aggr.rds",
        co = config["sim_dir"]
        + "post/noranef-noranef/vln.co.rds"
    output:
        config["fig_dir"] + "sim_rocc_vln.pdf"
    shell:
        "module add r/4.1.3; "
        "Rscript plot_sim_rocc_vln.R"
        " {input.rocc}"
        " {input.aggr}"
        " {input.co}"
        " {output}"
        
rule plot_wntrna_gp:
    input:
        config["wntrna_dir"]
        + "pre/rsd/pc10/data.list.rds",
        expand(config["wntrna_dir"]
               + "post/kin/pc10/{method}/"
               + "fit/complete{start}_{size}.txt",
               method = method,
               start = wntrna_start, size = size),
    output:
        config["fig_dir"] + "wntrna_gp.pdf" 
    shell:
        "module add r/4.1.3; "
        "Rscript plot_wntrna_gp.R"

rule plot_wntrna_co:
    input:
        config["wntrna_dir"]
        + "pre/rsd/pc10/data.list.rds",
        expand(config["wntrna_dir"]
               + "post/kin/pc10/{method}/"
               + "fit/complete{start}_{size}.txt",
               method = method,
               start = wntrna_start, size = size),
    output:
        config["fig_dir"] + "wntrna_gp_co.pdf" 
    shell:
        "module add r/4.1.3; "
        "Rscript plot_wntrna_co.R"
    
rule plot_wnt_llk:
    input:
        expand(config["wntrna_dir"]
               + "aa/llk/pc10/{cond}/llk.rds",
               cond = cond),
        expand(config["wntatac_dir"]
               + "aa/llk/pc10/{cond}/llk.rds",
               cond = cond),
    output:
        config["fig_dir"] + "wnt_llk.pdf" 
    shell:
        "module add r/4.1.3; "
        "Rscript plot_wnt_llk.R"

rule pw_sim_calib:
    input:
        expand(config["sim_dir"]
               + "{{comp}}/{gen}-{fit}/calib.rds",
               gen = gen, fit = fit),
    output:
        config["fig_dir"]
        + "sim/{comp}/calib.pdf",
    shell:
        "module add r/4.1.3; "
        "Rscript pw_sim_calib.R"
        " " + config["sim_dir"]
        + "{wildcards.comp}"
        " " + config["fig_dir"]
        + "sim/{wildcards.comp}"
        " {output}"

rule pw_sim_rocc:
    input:
        expand(config["sim_dir"]
               + "{{comp}}/{gen}-{fit}/rocc.rds",
               gen = gen, fit = fit),
    output:
        config["fig_dir"]
        + "sim/{comp}/rocc.pdf",
    shell:
        "module add r/4.1.3; "
        "Rscript pw_sim_rocc.R"
        " " + config["sim_dir"]
        + "{wildcards.comp}"
        " " + config["fig_dir"]
        + "sim/{wildcards.comp}"
        " {output}"

rule pw_sim_hist:
    input:
        expand(config["sim_dir"]
               + "{{comp}}-hp/{gen}-{fit}/{{method}}/"
               "opt/pp.rds",
               gen = gen, fit = fit),
    output:
        config["fig_dir"]
        + "sim/{comp}/{method}/hist.pdf",
    shell:
        "module add r/4.1.3; "
        "Rscript pw_sim_hist.R"
        " " + config["sim_dir"]
        + "{wildcards.comp}-hp"
        " {wildcards.comp}"
        " {wildcards.method}"
        " " + config["fig_dir"]
        + "sim/{wildcards.comp}/"
        "{wildcards.method}"
        " {output}"
                
rule pw_sim_vln:
    input:
        expand(config["sim_dir"]
               + "{{comp}}/{gen}-{fit}/vln.rds",
               gen = gen, fit = fit),
    output:
        first = config["fig_dir"]
        + "sim/{comp}/vln1.pdf",
        second = config["fig_dir"]
        + "sim/{comp}/vln2.pdf",
    shell:
        "module add r/4.1.3; "
        "Rscript pw_sim_vln.R"
        " " + config["sim_dir"]
        + "{wildcards.comp}"
        " " + config["fig_dir"]
        + "sim/{wildcards.comp}"
        " {output.first}"
        " {output.second}"
    
rule pw_sim_vln_aggr:
    input:
        expand(config["sim_dir"]
               + "{{comp}}/{gen}-{fit}/vln.aggr.rds",
               gen = gen, fit = fit),
    output:
        config["fig_dir"]
        + "sim/{comp}/vln_aggr.pdf",
    shell:
        "module add r/4.1.3; "
        "Rscript pw_sim_vln_aggr.R"
        " " + config["sim_dir"]
        + "{wildcards.comp}"
        " " + config["fig_dir"]
        + "sim/{wildcards.comp}"
        " {output}"

rule plot_sim_lnpy:
    input:
        mcmc_bs = config["sim_dir"]
        + "mcmc-bs-hp/{gen}-{fit}/",
        map_lap = config["sim_dir"]
        + "map-lap-hp/{gen}-{fit}/",
    output:
        data = config["dir"]
        + "sim.lnpy.{gen}.{fit}.rds",
        fig = config["fig_dir"]
        + "sim_lnpy_{gen}_{fit}.pdf",
    shell:
        "module add r/4.1.3; "
        "Rscript plot_sim_lnpy.R"
        " {input.mcmc_bs}"
        " {input.map_lap}"
        " {output.data}"
        " {output.fig}"

rule pw_sim_compare_sum_lnpy:
    input:
        expand(config["sim_dir"]
               + "comparison/{gen}-{fit}/"
               "compare.sum.lnpy.rds",
               gen = gen, fit = fit),
    output:
        config["fig_dir"]
        + "sim_compare_sum_lnpy.pdf",
    shell:
        "module add r/4.1.3; "
        "Rscript pw_sim_compare_sum_lnpy.R"
        
rule pw_sim_compare_pp:
    input:
        expand(config["sim_dir"]
               + "comparison/{gen}-{fit}/pp.rds",
               gen = gen, fit = fit),
    output:
        config["fig_dir"]
        + "sim_compare_pp.pdf",
    shell:
        "module add r/4.1.3; "
        "Rscript pw_sim_compare_pp.R"
        
rule plot_wntrna_compare_pp:
    input:
        expand(config["wntrna_dir"]
               + "post/kin/pc{pc}/{method}/pp.rds",
               pc = pc, method = method),
        expand(config["wntrna_dir"]
               + "post/id/pc{pc}/{method}/pp.rds",
               pc = pc, method = method),
    output:
        config["fig_dir"] + "wntrna_compare_pp.pdf"
    shell:
        "module add r/4.1.3; "
        "Rscript plot_compare_pp.R"

rule plot_wntrna_hm:
    input:
        nosign = config["wntrna_dir"]
        + "post/kin/pc10/nl/pp.rds",
        sign = config["wntrna_dir"]
        + "post/kin/pc10/nl/pp.sign.rds",
        order = config["wntrna_dir"]
        + "post/kin/pc10/nl/order.vec.rds",
    output:
        config["fig_dir"] + "wntrna_hm.pdf"
    shell:
        "module add r/4.1.3; "
        "Rscript plot_hm.R"
        " {input.nosign}"
        " {input.sign}"
        " {input.order}"
        " {output}"
        
rule plot_wntatac_gp:
    input:
        config["wntatac_dir"]
        + "pre/rsd/pc10/data.list.rds",
        expand(config["wntatac_dir"]
               + "post/pc10/{method}/"
               + "fit/complete{start}_{size}.txt",
               method = method,
               start = wntatac_start, size = size),
    output:
        config["fig_dir"] + "wntatac_gp.pdf"
    shell:
        "module add r/4.1.3; "
        "Rscript plot_wntatac_gp.R"

rule plot_wntatac_co:
    input:
        config["wntatac_dir"]
        + "pre/rsd/pc10/data.list.rds",
        expand(config["wntatac_dir"]
               + "post/pc10/{method}/"
               + "fit/complete{start}_{size}.txt",
               method = method,
               start = wntatac_start, size = size),
    output:
        config["fig_dir"] + "wntatac_gp_co.pdf"
    shell:
        "module add r/4.1.3; "
        "Rscript plot_wntatac_co.R"

rule plot_wntatac_hm:
    input:
        nosign = config["wntatac_dir"]
        + "post/pc10/nl/pp.rds",
        sign = config["wntatac_dir"]
        + "post/pc10/nl/pp.sign.rds",
        order = config["wntatac_dir"]
        + "post/pc10/nl/order.vec.rds",
    output:
        config["fig_dir"] + "wntatac_hm.pdf"
    shell:
        "module add r/4.1.3; "
        "Rscript plot_hm.R"
        " {input.nosign}"
        " {input.sign}"
        " {input.order}"
        " {output}"
```
