## Supplemental File 2 for "Probabilistic classification of gene-by-treatment interactions on molecular count phenotypes": utils.R

```
# suppressPackageStartupMessages(library(tidyverse))

get_geno_label <- function(snp.id, geno) {

    ref <- snp.id %>%
        strsplit(":") %>%
        unlist %>%
        `[`(3)

    alt <- snp.id %>%
        strsplit(":") %>%
        unlist %>%
        `[`(4)    
    
    if (geno == "0|0") {
        a1 <- ref; a2 <- alt        
    } else if (geno == "1|1") {
        a1 <- alt; a2 <- ref
    }    

    geno.label <- c(
        paste0(a1, "/", a1),
        paste0(a1, "/", a2),
        paste0(a2, "/", a2))

    geno.label
    
}
```
