## Supplemental File 2 for "Probabilistic classification of gene-by-treatment interactions on molecular count phenotypes": config.yaml

```
dir: "../derived_data/wntatac/"
fig_dir: "../figures/wntatac/"
indir: "../source_data/WntATAC/rasqual/InputFiles/"
data: "../source_data/WntATAC/LMM/LM_DF/LMM_Dataframe_VehCT99_LDZeroPointFive_25kb_50PC10MDS_HigherLDFilter0123.csv"
genesis_prefix: "../source_data/WntATAC/MDS/Wnt_all.geno.MDS"
count: "../source_data/WntATAC/CSAW/Counts/WindowCounts_VehCT99Wnt3a_XAV_noNkx.Rdata"
sample: "../source_data/WntATAC/QC/OmitList/VCF_DonorDNAID_1021.txt"
caqtl: "../source_data/media-10.xlsx"
rcaqtl: "../source_data/media-13.xlsx"
lb: 25
ub: 300
step: 25
num_pair: 1775 # number of pairs
pc: [10] # number of pcs
size: 10 # batch size for mcmc
seed: 1
num_core: 4
```
