## Supplemental File 2 for "Probabilistic classification of gene-by-treatment interactions on molecular count phenotypes": format_kin.R

```
suppressPackageStartupMessages(library(tidyverse))

args <- commandArgs(TRUE)
data.file <- as.character(args[1])
kin.file <- as.character(args[2])
out.file <- as.character(args[3])

# read in the data
data.list <- data.file %>% readRDS

# get the kinship matrix with donor and dna ids
A <- kin.file %>% readRDS

donor.dna <- data.list[[1]]$data$dna %>%
    unique
donor <- data.list[[1]]$data$dna %>%
    sapply(function(x) unlist(strsplit(x, "_"))[1]) %>%
    unique
dna <- data.list[[1]]$data$dna %>%
    sapply(function(x) unlist(strsplit(x, "_"))[2]) %>%
    unique

# if (!all(order(colnames(A)) == order(dna))) {
if (any(is.na(match(dna, colnames(A))))) {
    stop("DNA IDs do not match.")
}    

A <- A[dna, dna]
rownames(A) <- colnames(A) <- donor.dna
A %>% saveRDS(file=out.file)
```
