## Supplemental File 2 for "Probabilistic classification of gene-by-treatment interactions on molecular count phenotypes": get_anno.R

```
suppressPackageStartupMessages(library(tidyverse))
suppressPackageStartupMessages(library(vroom))

args <- commandArgs(TRUE)
in.file <- as.character(args[1])
out.dir <- as.character(args[2])

df <- vroom(in.file)

n.donor <- df %>%
    pull(Donor) %>%
    unique %>%
    length
n.sample <- n.donor * 2
n.data <- nrow(df) / n.sample 

input.list <-
    split(df, rep(seq_len(n.data), each=n.sample))

for (i in seq_along(input.list)) {

    out.file <- file.path(out.dir,
        paste0("data", str_pad(i, 6, pad="0"), ".rds"))
    if (file.exists(out.file)) {
        warning(paste("The file", out.file, "already exists."))
        next
    }

    x <- input.list %>% pluck(i)
    feat.id <- x %>% pull(Feature) %>% unique
    snp.id <- x %>% pull(SNP) %>% unique

    if (length(feat.id) != 1) {
        stop(
            paste0("Feature is not unique for the ",
                   i, "-th data."))
    }
    if (length(snp.id) != 1) {
        stop(
            paste0("SNP is not unique for the ",
                   i, "-th data."))
    }

    d <- x %>%
        dplyr::select(
                   c("Counts", "Genotype",
                     "Treatment", "DNAID",
                     paste0("V", 1:10))) %>%
        as.data.frame %>%
        `colnames<-`(
            c("y", "g", "t", "dna", paste0("pc", 1:10)))

    # swap the genotype levels
    # if the MAF estimate is greater than 0.5
    if (mean(d$g) / 2 > 0.5) {
        g <- d$g
        d$g[g == 0] <- 2
        d$g[g == 2] <- 0
    }    
    
    output <- list(data=d, feat.id=feat.id, snp.id=snp.id)
    saveRDS(output, file=out.file)
}

file.path(out.dir, "complete.txt") %>%
    file.create
```
