## Supplemental File 2 for "Probabilistic classification of gene-by-treatment interactions on molecular count phenotypes": get_input_for_aa.R

```
# modified from 01_RunLMM_PCRand_JobRunCT99.R
suppressPackageStartupMessages(library(tidyverse))
suppressPackageStartupMessages(library(readxl))

args <- commandArgs(TRUE)
dir.in <- as.character(args[1])
file.hit <- as.character(args[2])
file.vst <- as.character(args[3])
cond <- as.character(args[4])
dir.out <- as.character(args[5])

m.vst <- file.vst %>% readRDS

# read in response caQTL
if (cond == "Veh") {
    sheet <- 1 # Veh
} else if (cond == "CT99") {
    sheet <- 3 # CHIR
}    
d.hit <- file.hit %>% read_excel(sheet=sheet)

# select peaks and snps on autosomes
d.hit <- d.hit %>%
    filter(Chromosome != "chrX")

# select relevant columns
d.hit <- d.hit %>%
    dplyr::select(1:2) %>%
    `colnames<-`(c("Feature", "SNP"))

v.file <- dir.in %>%
    list.files(pattern="^data", full.names=TRUE)
v.index <- dir.in %>%
    list.files(pattern="^data") %>%
    gsub("data", "", .) %>%
    gsub("[.]rds", "", .)

# get indices for the hits
d.id <- matrix(NA, nrow=length(v.file), ncol=3) %>%
    as.data.frame %>%
    `colnames<-`(c("feat", "snp", "index"))
for (i in seq_along(v.file)) {
    if (i %% 1000 == 0) cat(i, "\t")
    l <- v.file[i] %>% readRDS
    d.id[i, ] <- c(l$feat.id, l$snp.id, v.index[i])
}    

# merge the data frames
d.join <- d.hit %>%
    inner_join(
        d.id, join_by(Feature == feat, SNP == snp)) %>%
    as.data.frame %>%
    `colnames<-`(c("feat", "snp", "index"))

n.data <- d.join %>% nrow

for (i in seq_len(n.data)) {
    if (i %% 1000 == 0) cat(i, "\t")

    index <- d.join$index[i]

    file.in <- file.path(
        dir.in, paste0("data", index, ".rds"))
    if (!file.exists(file.in)) {
        warning(paste0("The file ", file.in, " does not exists."))
        next
    }
    
    file.out <- file.path(dir.out,
        paste0("data", str_pad(i, 6, pad="0"), ".rds"))
    if (file.exists(file.out)) {
        warning(paste("The file", file.out, "already exists."))
        next
    }
    
    d.in <- try(file.in %>% readRDS, silent=TRUE)
    if (class(d.in) == "try-error") {
        warning(paste0("The file ", file.in, " is not readable."))
        next
    }
    
    data <- d.in$data
    feat.id <- d.in$feat.id
    snp.id <- d.in$snp.id

    # note that either condition can be used
    # to extract a list of unique donors
    data <- data %>%
        filter(t == 0) %>%
        dplyr::select(c(dna, g))
    v.donor.g <- data %>%
        pull(dna) %>%
        str_split_fixed("_", 2) %>%
        as.data.frame %>%
        pull(V1)
    rownames(data) <- v.donor.g
    
    v.pheno <- m.vst[rownames(m.vst) == feat.id, ]
    v.donor.p <- v.pheno %>%
        names %>%
        str_split_fixed("-", 3) %>%
        as.data.frame %>%
        pull(V2) %>%
        paste0("D", .)
    names(v.pheno) <- v.donor.p

    d.out <- merge(x=data, y=v.pheno, by.x=0, by.y=0) %>%
        dplyr::select(-dna) %>%
        `colnames<-`(c("dna", "g", "y"))
    
    output <- list(data=d.out, feat.id=feat.id, snp.id=snp.id)
    saveRDS(output, file=file.out)

}

file.path(dir.out, "complete.txt") %>%
    file.create
```
