## Supplemental File 2 for "Probabilistic classification of gene-by-treatment interactions on molecular count phenotypes": get_input_for_bms.R

```
suppressMessages(library(tidyverse))
suppressMessages(library(readxl))

args <- commandArgs(TRUE)
in.dir <- as.character(args[1])
hit.file <- as.character(args[2])
cnt.file <- as.character(args[3])
sample.file <- as.character(args[4])
out.file <- as.character(args[5])

# read in response caQTL (Veh vs CT99)
sheet <- 2 # CT99, CHIR
hit <- hit.file %>% read_excel(sheet=sheet)

# read in log-transformed count data
cnt.mat <- cnt.file %>% readRDS

# read in the sample map
sample <- sample.file %>% readRDS

# select peaks and snps on autosomes
hit <- hit %>%
    filter(Chr != "chrX")

# select relevant columns
hit <- hit %>%
    dplyr::select(1:4)

# get feature and snp ids from the input data
file.vec <- in.dir %>%
    list.files(pattern="^data", full.names=TRUE)
index.vec <- in.dir %>%
    list.files(pattern="^data") %>%
    gsub("data", "", .) %>%
    gsub("[.]rds", "", .)

# get indices for the hits
id <- matrix(NA, nrow=length(file.vec), ncol=3) %>%
    as.data.frame %>%
    `colnames<-`(c("feat", "snp", "index"))
for (i in seq_along(file.vec)) {
    if (i %% 1000 == 0) cat(i, "\t")
    l <- file.vec[i] %>% readRDS
    id[i, ] <- c(l$feat.id, l$snp.id, index.vec[i])
}    

# merge the data frames
joined <- hit %>%
    inner_join(
        id, join_by(Feature == feat, SNP == snp)) %>%
    as.data.frame %>%
    `colnames<-`(c("feat", "snp", "pval", "beta", "index"))

# create a list object
data.list <- vector("list", nrow(joined)) 

for (i in seq_len(nrow(joined))) {

    l <- file.path(
        in.dir, paste0("data", joined$index[i], ".rds")) %>%
        readRDS
    feat.id <- l$feat.id
    snp.id <- l$snp.id
    d <- l$data
    anno <- d %>% dplyr::select(dna, g, t)
    donor.dna <- anno$dna %>%
        str_split_fixed("_", 2) %>%
        as.data.frame %>%
        `colnames<-`(c("donor", "dna"))
    
    donor.vec <- donor.dna$donor %>% gsub("D", "", .)
    treat.vec <- anno$t
    treat.vec[anno$t == 0] <- 1
    treat.vec[anno$t == 1] <- 2
    sample1 <- paste0(donor.vec, "-", treat.vec)

    cnt.vec <- cnt.mat[feat.id, ]
    sample.vec <- names(cnt.vec)
    info <- str_split_fixed(sample.vec, "-", 3) %>%
        as.data.frame %>%
        `colnames<-`(c("week", "donor", "treatment"))
    sample2 <- paste0(info$donor, "-", info$treatment) 
    names(cnt.vec) <- sample2
    cnt.vec <- cnt.vec[sample1]

    # d <- list(
    #     y= cnt.vec,
    #     g = anno$g,
    #     t = anno$t,
    #     subject = anno$dna,
    #     feat.id=feat.id,
    #     snp.id=snp.id)

    d <- data.frame(
        y=cnt.vec,
        g=anno$g,
        t=anno$t,
        dna=anno$dna) # donor and dna ids
    
    # order the data by the donor and dna ids
    d <- d[order(d$dna), ]
    l$data <- d
    data.list[[i]] <- l

}

saveRDS(data.list, file=out.file)
```
