## Supplemental File 2 for "Probabilistic classification of gene-by-treatment interactions on molecular count phenotypes": get_kin.R

```
# modified from `00.prep.eqtl.data.R`
suppressPackageStartupMessages(library(GENESIS))
suppressPackageStartupMessages(library(GWASTools))
suppressPackageStartupMessages(library(SNPRelate))
suppressPackageStartupMessages(library(tidyverse))

args <- commandArgs(TRUE)
bed.file <- as.character(args[1])
bim.file <- as.character(args[2])
fam.file <- as.character(args[3])
gds.file <- as.character(args[4])
out.file <- as.character(args[5])

# create an genotypedata class object from plink files
snpgdsBED2GDS(
    bed.fn=bed.file,
    bim.fn=bim.file,
    fam.fn=fam.file,
    out.gdsfn=gds.file)

# run pcair
geno <- GdsGenotypeReader(filename=gds.file)
genoData <- GenotypeData(geno)
mypcair <- pcair(genoData)

# estimate relatedness 
genoData <- GenotypeBlockIterator(genoData)
mypcrelate <- pcrelate(genoData, pcs=mypcair$vectors[, 1:10])

# creates a genetic relationship matrix of pairwise kinship
# coefficient estimates from pc-relate output
genesis.mat <- mypcrelate %>%
    pcrelateToMatrix %>%
    as.matrix

saveRDS(genesis.mat, file=out.file)
```
