## Supplemental File 2 for "Probabilistic classification of gene-by-treatment interactions on molecular count phenotypes": get_llk.R

```
suppressPackageStartupMessages(library(tidyverse))
suppressPackageStartupMessages(library(rstan))

args <- commandArgs(TRUE)
dir.in <- as.character(args[1])
seed <- as.integer(args[2])
file.out <- as.character(args[3])

rstan_options(auto_write=TRUE)

# load stan model
nl.stan <- stan_model(file="nl.stan")
lm.stan <- stan_model(file="lm.stan")

n.data <- dir.in %>%
    list.files(pattern="data") %>%
    length

d.llk <- matrix(NA, nrow=n.data, ncol=3) %>%
    as.data.frame %>%
    `colnames<-`(c("lm", "nl", "anova"))

for (i in seq_len(n.data)) {
    
    if (i %% 1000 == 0) cat(i, "\t")
    file.in <- file.path(dir.in,
        paste0("data", str_pad(i, 6, pad="0"), ".rds"))
    if (!file.exists(file.in)) {
        warning(paste0("The file ", file.in, " does not exists."))
        next
    }

    l <- file.in %>% readRDS
    l <- try(file.in %>% readRDS, silent=TRUE)
    if (class(l) == "try-error") {
        warning(paste0("The file ", file.in, " is not readable."))
        next
    }
    
    data <- l$data
    y <- data$y
    n <- length(y)
    g <- data$g
    
    # fit anova model    
    if (all(table(g) > 0)) {
        g1 <- as.numeric(g == 1)
        g2 <- as.numeric(g == 2)
        fit.anova <- lm(y ~ g1 + g2)
        llk.anova <- logLik(fit.anova)
    } else {
        llk.anova <- NA
    }    
    
    # fit linear model
    fit.lm <- lm(y ~ g)
    llk.lm <- logLik(fit.lm)

    # fit nonlinear model
    x1 <- (1 - g/2)
    x2 <- (g/2)
    data.stan <- list(
        N=n, y=y,
        x1=x1, x2=x2)
    fit.nl <- optimizing(
        object=nl.stan,
        data=data.stan,
        seed=seed,
        algorithm="BFGS")
    llk.nl <- fit.nl$value

    d.llk[i, ] <- c(llk.lm, llk.nl, llk.anova)
}

# save the results
saveRDS(d.llk, file=file.out)
```
