## Supplemental File 2 for "Probabilistic classification of gene-by-treatment interactions on molecular count phenotypes": get_pc_for_bms.R

```
suppressPackageStartupMessages(library(tidyverse))

args <- commandArgs(TRUE)
in.file <- as.character(args[1])
pca.file <- as.character(args[2])

vst.mat <- in.file %>% readRDS

# regress out the treatment indicator
n <- ncol(vst.mat) / 2
t <- rep(c(0, 1), each=n)
rsd.mat <-
    apply(vst.mat, 1, function(x) residuals(lm(x ~ 1 + t)))

# perform pca
pca <- prcomp(rsd.mat)

saveRDS(pca, file=pca.file)
```
