## Supplemental File 2 for "Probabilistic classification of gene-by-treatment interactions on molecular count phenotypes": get_phenomat.R

```
suppressPackageStartupMessages(library(tidyverse))
suppressPackageStartupMessages(library(data.table))

args <- commandArgs(TRUE)
file.in <- as.character(args[1])
file.out <- as.character(args[2])

# read in input files
d <- file.in %>% fread(data.table=FALSE)

# get peak names
peaks <- d[, 1]

# get counts (remove the first four columns)
d <- d[, -(1:4)] 

# get sample names `<week>-<donor>-<condition>`
samples <- colnames(d) %>%
    str_split_fixed("-", 4) %>%
    apply(1, function(x) paste(x[1:3], collapse="-"))

colnames(d) <- samples 
rownames(d) <- peaks
m <- d %>% as.matrix

saveRDS(m, file=file.out)
```
