## Supplemental File 2 for "Probabilistic classification of gene-by-treatment interactions on molecular count phenotypes": get_pp_for_map_lap.R

```
suppressPackageStartupMessages(library(tidyverse))
suppressPackageStartupMessages(library(classifygxt))

args <- commandArgs(TRUE)
in.file <- as.character(args[1])
data.file <- as.character(args[2])
in.dir <- as.character(args[3])
out.file <- as.character(args[4])

# get the optimal hyperparameter values
ln.p.y <- in.file %>% readRDS
opt <- rownames(ln.p.y)[which.max(ln.p.y$ln.p.y)]

fit.file <- file.path(in.dir, opt, "fit.rds") 

# read in and process the data
fit.list <- fit.file %>% readRDS
data.list <- data.file %>% readRDS

pp <- fit.list %>%
    map(function(x) x$p.m.given.y) %>%
    t %>%
    do.call("rbind", .) %>%
    as.data.frame

# save the result
saveRDS(pp, file=out.file)
```
