## Supplemental File 2 for "Probabilistic classification of gene-by-treatment interactions on molecular count phenotypes": get_pp_for_mcmc_bs.R

```
suppressPackageStartupMessages(library(tidyverse))
suppressPackageStartupMessages(library(classifygxt))

args <- commandArgs(TRUE)
in.dir <- as.character(args[1])
out.file <- as.character(args[2])

file.vec <- list.files(
    in.dir, pattern="^res", full.names=TRUE) %>%
    grep("rds$", ., value=TRUE)

model.name <- get_model_names()

pp <- file.vec %>%
    map(
        function(x) readRDS(x) %>%
                    `[[`("p.m.given.y")) %>%
    do.call("rbind", .) %>%
    `colnames<-`(model.name) %>%
    as.data.frame

# save the result
saveRDS(pp, file=out.file)
```
