## Supplemental File 2 for "Probabilistic classification of gene-by-treatment interactions on molecular count phenotypes": get_resid_for_aa.R

```
suppressPackageStartupMessages(library(tidyverse))

args <- commandArgs(TRUE)
dir.in <- as.character(args[1])
file.pca <- as.character(args[2])
num.pc <- as.integer(args[3])
dir.out <- as.character(args[4])

pca <- file.pca %>% readRDS
m.pca <- pca$x

nm.row <- m.pca %>%
    rownames %>%
    str_split_fixed("-", 3) %>%
    as.data.frame %>%
    pull(2) %>%
    paste0("D", .)
rownames(m.pca) <- nm.row

n.data <- dir.in %>%
    list.files(pattern="data") %>%
    length

for (i in seq_len(n.data)) {

    if (i %% 1000 == 0) cat(i, "\t")

    file.in <- file.path(dir.in,
        paste0("data", str_pad(i, 6, pad="0"), ".rds"))
    if (!file.exists(file.in)) {
        warning(paste0("The file ", file.in, " does not exists."))
        next
    }

    file.out <- file.path(dir.out,
        paste0("data", str_pad(i, 6, pad="0"), ".rds"))
    if (file.exists(file.out)) {
        warning(paste("The file", file.out, "already exists."))
        next
    }
    
    l <- try(file.in %>% readRDS, silent=TRUE)
    if (class(l) == "try-error") {
        warning(paste0("The file ", file.in, " is not readable."))
        next
    }
    
    d <- l$data
    d <- merge(d, m.pca[, seq_len(num.pc)], by.x=1, by.y=0)

    # get residuals
    frml <- paste0(
        "y ~ ",
        paste0(paste0("PC", seq_len(num.pc)), collapse=" + ")) 
    d$y <- residuals(lm(frml, data=d))
    
    # remove pcs
    d <- d[, 1:3]

    # replace the data
    l$data <- d

    saveRDS(l, file=file.out)
}

file.path(dir.out, "complete.txt") %>%
    file.create
```
