## Supplemental File 2 for "Probabilistic classification of gene-by-treatment interactions on molecular count phenotypes": get_resid_for_bms.R

```
suppressPackageStartupMessages(library(tidyverse))

args <- commandArgs(TRUE)
data.file <- as.character(args[1])
pca.file <- as.character(args[2])
num.pc <- as.integer(args[3])
out.file <- as.character(args[4]) 

data.list <- data.file %>% readRDS

pca <- pca.file %>% readRDS
pca.mat <- pca$x

row.name <- pca.mat %>%
    rownames %>%
    str_split_fixed("-", 2) %>%
    as.data.frame %>%
    pull(2)
rownames(pca.mat) <- row.name

for (i in seq_along(data.list)) {

    l <- data.list[[i]]
    d <- l$data %>%
        dplyr::select(c(y, g, t, dna))
    sample <- d %>%
        pull(dna) %>%
        sapply(function(x) unlist(strsplit(x, "_"))[1]) %>%
        gsub("D", "", .) %>%
        paste(rep(1:2, each=nrow(d) / 2), sep="-")
    d <- d %>%
        mutate(sample=sample)
    d <- merge(d, pca.mat[, seq_len(num.pc)], by.x=5, by.y=0)
    
    # get residuals
    frml <- paste0(
        "y ~ ",
        paste0(paste0("PC", seq_len(num.pc)), collapse=" + ")) 
    d$y <- residuals(lm(frml, data=d))
    
    # remove pcs
    d <- d[, 1:5]

    # replace the data
    l$data <- d
    data.list[[i]] <- l 

}

saveRDS(data.list, file=out.file)
```
