## Supplemental File 2 for "Probabilistic classification of gene-by-treatment interactions on molecular count phenotypes": get_samplemap.R

```
suppressPackageStartupMessages(library(tidyverse))
suppressPackageStartupMessages(library(here))
suppressPackageStartupMessages(library(readxl))

args <- commandArgs(TRUE)
count.file <- as.character(args[1])
sample.file <- as.character(args[2])
out.file <- as.character(args[3])

# read in the data
load(count.file)
id <- scan(sample.file, what="")
colnames(counts) <- samples

# separate info for sample
# week, donor, and treatment condition
info <- as.data.frame(str_split_fixed(samples, "-", 4))

info <- info[, 1:3]
colnames(info) <- c("week", "donor", "treatment")
info <- cbind(samples, info)

x <- info$treatment
x[x == 1] <- "Veh"
x[x == 2] <- "CT99"
x[x == 5] <- "Wnt3a"
x[x == 6] <- "XAV"

info$treatment <- x
info$donor <- paste0("D", info$donor)

d <- sapply(id, function(x) unlist(strsplit(x, "_"))) %>%
    t %>% as.data.frame
d$id <- id
colnames(d) <- c("donor", "dna", "id")

merged <- merge(info, d, by.x="donor", by.y="donor")
merged <- merged[, c(6, 1, 5, 4, 3, 2)]
colnames(merged)[6] <- "sample"

saveRDS(merged, file=out.file)
```
