## Supplemental File 2 for "Probabilistic classification of gene-by-treatment interactions on molecular count phenotypes": get_sign_for_map_lap.R

```
suppressPackageStartupMessages(library(tidyverse))
suppressPackageStartupMessages(library(readxl))
suppressPackageStartupMessages(library(classifygxt))

args <- commandArgs(TRUE)
prev.file <- as.character(args[1])
data.file <- as.character(args[2])
lnpy.file <- as.character(args[3])
in.dir <- as.character(args[4])
pp.file <- as.character(args[5])
order.file <- as.character(args[6])

# read in the previous results
prev <- read_excel(prev.file, sheet=2) %>%
    as.data.frame

# read in the input data file
data.list <- data.file %>% readRDS

# read in the sum of the log of maximum likelihood
ln.p.y <- lnpy.file %>% readRDS

# get the optimal hyperparameter values
opt <- rownames(ln.p.y)[which.max(ln.p.y$ln.p.y)]
# opt <- "phi1.50_2.00_1.00"

# read in the corresponding model fitting result
fit.file <- file.path(in.dir, opt, "fit.rds")
fit.list <- fit.file %>% readRDS

pp <- fit.list %>%
    sapply(get_sign) %>%
    t

# get the order of the pairs by $p$-values
feat.id.vec <- sapply(data.list, function(x) x$feat.id)
snp.id.vec <- sapply(data.list, function(x) x$snp.id)
id.vec <- paste(feat.id.vec, snp.id.vec, sep="-")
prev.id.vec <- paste(prev$Feature, prev$SNP, sep="-")
order.vec <- match(prev.id.vec, id.vec) %>% na.omit

saveRDS(pp, file=pp.file)
saveRDS(order.vec, file=order.file)
```
