## Supplemental File 2 for "Probabilistic classification of gene-by-treatment interactions on molecular count phenotypes": get_tu_lambda.R

```
suppressPackageStartupMessages(library(tidyverse))
suppressPackageStartupMessages(library(classifygxt))

args <- commandArgs(TRUE)
data.file <- as.character(args[1])
kin.file <- as.character(args[2]) # file name or "NA"
out.file <- as.character(args[3])

if (kin.file == "NA") {
    kinship <- NULL
} else {
    kinship <- kin.file %>% readRDS
}
    
# read in the data
data.list <- data.file %>% readRDS

# extract the first (or any) data
data <- data.list[[1]]$data

d <- list(
    y=data$y, g=data$g, t=data$t, subject=data$dna)

# get a list of tU and lambda
tu.lambda <- get_tu_lambda(d, kinship=kinship)

saveRDS(tu.lambda, file=out.file)
```
