## Supplemental File 2 for "Probabilistic classification of gene-by-treatment interactions on molecular count phenotypes": lm.stan

```
data {
  int<lower=0> N; // number of individuals
  vector[N] y; // vector of observations
  vector<lower=0, upper=2>[N] x1; // genotype 
}
parameters {
  real b0; // intercept
  real b1; // coefficient of genotype
  real<lower=0> sigma2;
}
model {
  vector[N] mu;
  mu = b0 + x1 * b1;
  target += normal_lpdf(y | mu, sqrt(sigma2)); // likelihood model
}
```
