## Supplemental File 2 for "Probabilistic classification of gene-by-treatment interactions on molecular count phenotypes": README.md

# `wntatac`

This directory contains code for analyzing response caQTL data in human neural progenitor cells (hNPCs) treated with CHIR, an activator of the Wnt pathway.
See the following publication for information about the input files.
Note that files containing individual genotype data can be accessed only upon request and approval.

Nana Matoba\*,
Brandon D Le\*,
Jordan M Valone\*,
Justin M Wolter,
Jessica Mory,
Dan Liang,
Nil Aygun,
K Alaine Broadaway,
Marielle L Bond,
Karen L Mohlke,
Mark J Zylka,
Michael I Love,
Jason L Stein.
"Wnt activity reveals context-specific genetic effects on gene regulation in neural progenitors."
doi: <https://doi.org/10.1101/2023.02.07.527357>
(\* These authors contributed equally to this work.)

## Setup

The code requires the following input files in a directory(or a symbolic link to a directory) named `source_data` under the project directory.

- `WntATAC/media-10.xlsx`
- `WntATAC/media-13.xlsx`
- `WntATAC/CSAW/Counts/WindowCounts_VehCT99Wnt3a_XAV_noNkx.Rdata`
- `WntATAC/LMM/LM_DF/LMM_Dataframe_VehCT99_LDZeroPointFive_25kb_50PC10MDS_HigherLDFilter0123.csv`
- `WntATAC/MDS/Wnt_all.geno.MDS.bed`
- `WntATAC/MDS/Wnt_all.geno.MDS.bim`
- `WntATAC/MDS/Wnt_all.geno.MDS.fam`
- `WntATAC/QC/OmitList/VCF_DonorDNAID_1021.txt`
- `WntATAC/rasqual/InputFiles/CT99CountsTable.txt`
- `WntATAC/rasqual/InputFiles/VehCountsTable.txt`

## Credits

Portions of the `get_anno.R` and `get_kin.R` scripts have been adapted from <https://bitbucket.org/steinlabunc/wnt-rqtls/>.
