## Supplemental File 2 for "Probabilistic classification of gene-by-treatment interactions on molecular count phenotypes": Snakefile

```
lb = config["lb"]
ub = config["ub"]
step = config["step"]
num_pair = config["num_pair"] # number of pairs
pc = config["pc"] # number of pcs
size = config["size"] # batch size for mcmc
seed = config["seed"]
num_core = config["num_core"]

method = ["nl", "lm", "rint"]
cond = ["Veh", "CT99"]

numbers = [x / 100.00 for x in range(lb, ub + 1, step)]
phi1 = ["{:1.2f}".format(x) for x in numbers]
phi2 = ["{:1.2f}".format(x) for x in numbers]
phi3 = ["{:1.2f}".format(x) for x in numbers]

start = [format(number, "04")
         for number in range(1, num_pair + 1, size)]

rule all:
    input:
        expand(config["dir"]
               + "post/pc{pc}/{method}/"
               "pp.sign.rds",
               pc = pc, method = method),
        expand(config["dir"]
               + "optim/pc{pc}/{method}/"
               "pp.sign.rds",
               pc = pc, method = method),
        expand(config["dir"]
               + "optim/pc{pc}/{method}/"
               "order.vec.rds",
               pc = pc, method = method),
        expand(config["dir"]
               + "post/pc{pc}/{method}/pp.rds",
               pc = pc, method = method),
        expand(config["dir"]
               + "post/pc{pc}/{method}/"
               + "fit/complete{start}_{size}.txt",
               pc = pc, method = method,
               start = start, size = size),        
        expand(config["dir"]
               + "optim/pc{pc}/{method}/"
               "pp.rds",
               pc = pc, method = method),
        expand(config["dir"]
               + "optim/pc{pc}/{method}/"
               "ln.p.y.rds",
               pc = pc, method = method),
        expand(config["dir"]
               + "aa/llk/pc{pc}/{cond}/"
               + "llk.rds",
               pc = pc, cond = cond),

rule get_sign_for_mcmc_bs:
    input:
        prev = config["rcaqtl"],
        data = config["dir"]
        + "pre/rsd/pc{pc}/data.list.rds",
        fit = expand(config["dir"]
                     + "post/pc{{pc}}/{{method}}/fit/"
                     "complete{start}_{size}.txt",
                     start = start, size = size),
    output:
        pp = config["dir"]
        + "post/pc{pc}/{method}/pp.sign.rds",
        order = config["dir"]
        + "post/pc{pc}/{method}/order.vec.rds",
    shell:            
        "module add r/4.1.3; "
        "Rscript get_sign_for_mcmc_bs.R"
        " {input.prev}"
        " {input.data}"
        " " + config["dir"]
        + "post/pc{wildcards.pc}/"
        "{wildcards.method}/fit"
        " {output.pp}"
        " {output.order}"

rule get_sign_for_map_lap:
    input:
        prev = config["rcaqtl"],
        data = config["dir"]
        + "pre/rsd/pc{pc}/data.list.rds",
        lnpy = config["dir"]
        + "optim/pc{pc}/{method}/ln.p.y.rds",
    output:
        pp = config["dir"]
        + "optim/pc{pc}/{method}/pp.sign.rds",
        order = config["dir"]
        + "optim/pc{pc}/{method}/order.vec.rds",
    shell:            
        "module add r/4.1.3; "
        "Rscript get_sign_for_map_lap.R"
        " {input.prev}"
        " {input.data}"
        " {input.lnpy}"
        " " + config["dir"]
        + "optim/pc{wildcards.pc}/{wildcards.method}"
        " {output.pp}"
        " {output.order}"

rule get_pp_for_mcmc_bs:
    input:
        expand(config["dir"]
               + "post/pc{{pc}}/{{method}}/"
               + "fit/complete{start}_{size}.txt",
               start = start, size = size),
    output:
        config["dir"]
        + "post/pc{pc}/{method}/"
        + "pp.rds",
    shell:
        "module add r/4.1.3; "
        "Rscript get_pp_for_mcmc_bs.R"
        " " + config["dir"]
        + "post/pc{wildcards.pc}/"
        "{wildcards.method}/fit"
        " {output}"

rule get_pp_for_map_lap:
    input:
        fit = config["dir"]
        + "optim/pc{pc}/{method}/"
        + "ln.p.y.rds",
        data = config["dir"]
        + "pre/rsd/pc{pc}/data.list.rds"
    output:
        config["dir"]
        + "optim/pc{pc}/{method}/"
        + "pp.rds",
    shell:
        "module add r/4.1.3; "
        "Rscript get_pp_for_map_lap.R"
        " {input.fit}"
        " {input.data}"
        " " + config["dir"]
        + "optim/"
        "pc{pc}/"
        "{wildcards.method}"
        " {output}"
        
rule post:
    input:
        data = config["dir"]
        + "pre/rsd/pc{pc}/data.list.rds",
        tu_lambda = config["dir"] + "pre/tu.lambda.rds",
        fit = config["dir"]
        + "optim/pc{pc}/{method}/"
        + "ln.p.y.rds",
    output:
        config["dir"]
        + "post/pc{pc}/{method}/"
        + "fit/complete{start}_{size}.txt",
    shell:
        "module add gcc/11.2.0; "
        "module add r/4.1.3; "
        "Rscript do_mcmc_bs.R"
        " {input.data}"
        " {input.tu_lambda}"
        " {input.fit}"
        " {seed}"
        " {num_core}"
        " {wildcards.method}"
        " ranef"
        " opt"
        " sample"
        " {wildcards.start}"
        " {wildcards.size}"
        " " + config["dir"]
        + "post/pc{wildcards.pc}/"
        "{wildcards.method}/fit"

rule get_marginal_llk:
    input:
        expand(config["dir"]
               + "optim/pc{{pc}}/{{method}}/"
               "phi{phi1}_{phi2}_{phi3}/fit.rds",
               phi1 = phi1, phi2 = phi2, phi3 = phi3),
    output:
        config["dir"]
        + "optim/pc{pc}/{method}/ln.p.y.rds"
    shell:
        "module add r/4.1.3; "
        "Rscript get_marginal_llk.R"
        " " + config["dir"]
        + "optim/pc{wildcards.pc}/"
        "{wildcards.method}"
        " {lb}"
        " {ub}"
        " {step}"
        " {output}"

rule optimize_hp:
    input:
        data = config["dir"] + "pre/rsd/pc{pc}/data.list.rds",
        tu_lambda = config["dir"] + "pre/tu.lambda.rds",
    output:
        config["dir"]
        + "optim/pc{pc}/{method}/"
        "phi{phi1}_{phi2}_{phi3}/fit.rds",
    shell:
        "module add gcc/11.2.0; "
        "module add r/4.1.3; "
        "Rscript optimize_hp.R"
        " {input.data}"
        " {input.tu_lambda}"
        " {seed}"
        " {wildcards.phi1}"
        " {wildcards.phi2}"
        " {wildcards.phi3}"
        " {wildcards.method}"
        " ranef"
        " {output}"

rule get_tu_lambda:
    input:
        data = config["dir"] + "pre/"
        + "data.list.rds",
        kin = config["dir"] + "pre/kin.mat.rds"
    output:
        config["dir"] + "pre/tu.lambda.rds"
    shell:
        "module add r/4.1.3; "
        "Rscript get_tu_lambda.R"
        " {input.data}"
        " {input.kin}"
        " {output}"

rule format_kin:
    input:
        data = config["dir"] + "pre/"
        + "data.list.rds",
        kin = config["dir"] + "pre/genesis.mat.rds",
    output:
        kin = config["dir"] + "pre/kin.mat.rds"
    shell:
        "module add r/4.1.3; "
        "Rscript format_kin.R"
        " {input.data}"
        " {input.kin}"
        " {output}"

rule get_kin:
    input:
        bed = config["genesis_prefix"] + ".bed",
        bim = config["genesis_prefix"] + ".bim",
        fam = config["genesis_prefix"] + ".fam"
    output:
        gds = config["dir"] + "pre/genesis.gds",
        mat = config["dir"] + "pre/genesis.mat.rds"
    shell:
        "module add r/4.1.3; "
        "Rscript get_kin.R"
        " {input.bed}"
        " {input.bim}"
        " {input.fam}"
        " {output.gds}"
        " {output.mat}"
    
rule get_resid_for_bms:
    input:
        data = config["dir"] + "pre/data.list.rds",
        pca = config["dir"] + "pre/pca.rds",
    output:
        config["dir"] + "pre/rsd/pc{pc}/"
        + "data.list.rds"
    shell:
        "module add r/4.1.3; "
        "Rscript get_resid_for_bms.R"
        " {input.data}"
        " {input.pca}"
        " {wildcards.pc}"
        " {output}"

rule get_pc_for_bms:
    input:
        config["dir"] + "pre/log.mat.rds"
    output:
        config["dir"] + "pre/pca.rds",
    shell:
        "module add r/4.1.3; "
        "Rscript get_pc_for_bms.R"
        " {input}"
        " {output}"
        
rule get_input_for_bms:
    input:
        data = config["dir"] + "input/complete.txt",
        hit = config["rcaqtl"],
        vst = config["dir"] + "pre/log.mat.rds",
        sample = config["dir"] + "pre/samplemap.rds",
    output:
        config["dir"] + "pre/data.list.rds",
    shell:
        "module add r/4.1.3; "
        "Rscript get_input_for_bms.R"
        " " + config["dir"] + "input"
        " {input.hit}"
        " {input.vst}"
        " {input.sample}"
        " {output}"

rule combine_phenomat:
    input:
        expand(config["dir"]
               + "aa/log.{cond}.mat.rds",
               cond = cond)
    output:
        config["dir"] + "pre/log.mat.rds"
    shell:
        "module add r/4.1.3; "
        "Rscript combine_phenomat.R"
        " {input}"
        " {output}"

rule get_llk:
    input:
        config["dir"] + "aa/rsd/pc{pc}/{cond}/"
        + "complete.txt"
    output:
        config["dir"]
        + "aa/llk/pc{pc}/{cond}/"
        + "llk.rds"
    shell:
        "module add gcc/11.2.0; "
        "module add r/4.1.3; "
        "Rscript get_llk.R"
        " " + config["dir"] +
        "aa/rsd/pc{wildcards.pc}/{wildcards.cond}"
        " {seed}" 
        " {output}"
        
rule get_resid_for_aa:
    input:
        data = config["dir"] + "aa/input/{cond}/"
        + "complete.txt",
        pca = config["dir"] + "aa/pca.{cond}.rds",
    output:
        config["dir"] + "aa/rsd/pc{pc}/{cond}/"
        + "complete.txt"
    shell:
        "module add r/4.1.3; "
        "Rscript get_resid_for_aa.R"
        " " + config["dir"] +
        "aa/input/{wildcards.cond}"
        " {input.pca}"
        " {wildcards.pc}"
        " " + config["dir"] +
        "aa/rsd/pc{wildcards.pc}/{wildcards.cond}"

rule get_input_for_aa:
    input:
        data = config["dir"] + "input/complete.txt",
        caqtl = config["caqtl"],
        vst = config["dir"] + "aa/log.{cond}.mat.rds"
    output:
        config["dir"] + "aa/input/{cond}/"
        + "complete.txt"
    shell:
        "module add r/4.1.3; "
        "Rscript get_input_for_aa.R"
        " " + config["dir"] + "input"
        " {input.caqtl}"
        " {input.vst}"
        " {wildcards.cond}"
        " " + config["dir"] +
        "aa/input/{wildcards.cond}"

rule get_pc_for_aa:
    input:
        config["dir"] + "aa/log.{cond}.mat.rds"
    output:
        config["dir"] + "aa/pca.{cond}.rds"
    shell:
        "module add r/4.1.3; "
        "Rscript get_pc_for_aa.R"
        " {input}"
        " {output}"

rule transform_pheno:
    input:
        config["dir"] + "aa/cnt.{cond}.mat.rds"
    output:
        config["dir"] + "aa/log.{cond}.mat.rds"
    shell:
        "module add r/4.1.3; "
        "Rscript transform_pheno.R"
        " {input}"
        " {output}"

rule get_phenomat:
    input:
        config["indir"] + "Veh_CT99_Common/" +
        "{cond}CountsTable.txt"
    output:
        config["dir"] + "aa/cnt.{cond}.mat.rds",
    shell:
        "module add r/4.1.3; "
        "Rscript get_phenomat.R"
        " {input}"
        " {output}"
        
rule get_anno:
    input:
        config["data"]
    output:
        config["dir"] + "input/complete.txt"
    shell:
        "module add r/4.1.3; "
        "Rscript get_anno.R"
        " {input}"
        " " + config["dir"]
        + "input"

rule get_samplemap:
    input:
        data = config["count"],
        sample = config["sample"],
    output:
        config["dir"] + "pre/samplemap.rds",
    shell:
        "module add r/4.1.3; "
        "Rscript get_samplemap.R"
        " {input.data}"
        " {input.sample}"
        " {output}"
```
