## Supplemental File 2 for "Probabilistic classification of gene-by-treatment interactions on molecular count phenotypes": transform_pheno.R

```
suppressPackageStartupMessages(library(tidyverse))

args <- commandArgs(TRUE)
file.in <- as.character(args[1])
file.out <- as.character(args[2])

# read in a feature count matrix
m.fc <- file.in %>% readRDS

# sacle according to palowitch et al. (2018)
# sample i = 1, ..., n
# gene j = 1, ..., T
T <- nrow(m.fc)
n <- ncol(m.fc)
li <- colSums(m.fc)
xbar <- sum(li) / (T * n)
m.fc <- sweep(m.fc, 2, li, "/") * T * xbar

# transform
m.fc <- log(m.fc + 1)

saveRDS(m.fc, file=file.out)
```
