## Supplemental File 2 for "Probabilistic classification of gene-by-treatment interactions on molecular count phenotypes": combine_phenomat.R

```
suppressPackageStartupMessages(library(tidyverse))

args <- commandArgs(TRUE)
ctrl.file <- as.character(args[1])
trt.file <- as.character(args[2])
out.file <- as.character(args[3])

ctrl.mat <- ctrl.file %>% readRDS
trt.mat <- trt.file %>% readRDS

output <- cbind(ctrl.mat, trt.mat)

saveRDS(output, file=out.file)
```
