## Supplemental File 2 for "Probabilistic classification of gene-by-treatment interactions on molecular count phenotypes": config.yaml

```
dir: "../derived_data/wntrna/"
fig_dir: "../figures/wntrna/"
indir: "../source_data/WntRNA/limix_qtl/input/"
bam_dir: "../source_data/WntIIRNAseq/baseQTL_IntermediateFiles/STAR_mapped/"
# bam_dir: "../source_data/WntRNA/STAR_alignment/STAR_Aligned_Trimmed/"
genename: "../source_data/WntRNA/resources/Homo_sapiens.GRCh38.104.collapse_gene_ID.gene_name.txt"
gtf: "../source_data/WntRNA/resources/Homo_sapiens.GRCh38.104.collapse.gtf"
sample: "../source_data/WntRNA/limix_qtl/input/interaction/samplemap.interaction.veh.CT99.tsv"
pheno: "../source_data/WntRNA/limix_qtl/input/interaction/pheno.interaction.residuals.veh.CT99.PC10.MDS0.tsv"
kin: "../source_data/WntRNA/limix_qtl/input/Wnt_all.kinship.tsv"
eqtl: "../source_data/media-11.xlsx"
reqtl: "../source_data/media-14.xlsx"
```
