## Supplemental File 2 for "Probabilistic classification of gene-by-treatment interactions on molecular count phenotypes": sbatch.yaml

```
---
__default__:
  time: "0-01:00"
  threads: 1
  mem: 4G
  queue: "general"
  error: "../logs/wntrna/%x_%j_%N.err"
  output: "../logs/wntrna/%x_%j_%N.log"
post:
  time: "0-04:00"
  threads: 4
  mem: 8G
  queue: "general"
optimize_hp:
  time: "0-04:00"
  threads: 1
  mem: 4G
  queue: "general"
get_kin:
  time: "0-01:00"
  threads: 64
  mem: 64G
  queue: "general"
get_resid_for_aa:
  time: "0-04:00"
  threads: 1
  mem: 4G
  queue: "general"
get_input_for_aa:
  time: "0-04:00"
  threads: 1
  mem: 16G
  queue: "general"
get_phenomat:
  time: "0-00:30"
  threads: 1
  mem: 16G
  queue: "general"
get_phenomat:
  time: "0-01:00"
  threads: 1
  mem: 16G
  queue: "general"
get_anno:
  time: "0-04:00"
  threads: 1
  mem: 32G
  queue: "general"
format_geno:
  time: "0-01:00"
  threads: 1
  mem: 16G
  queue: "general"
```
