## Supplemental File 2 for "Probabilistic classification of gene-by-treatment interactions on molecular count phenotypes": format_geno.R

```
suppressPackageStartupMessages(library(tidyverse))

args <- commandArgs(TRUE)
file.tfam <- args[1]
file.tped <- args[2]
file.out <- args[3]

# read in the genotype data
geno <- file.tped %>% read.table(header=FALSE)

# read in the sample info
info <- file.tfam %>% read.table()

# get donor IDs
donors <- info %>% pull(V1)

# process the genotype data
snps <- geno %>% `[[`(2)
x <- geno %>%
    select(-(1:4)) %>%
    as.matrix()

# convert to minor allele dosages (0, 1, 2)
n <- x %>% ncol() %>% `/`(2)
x[x == 2] <- 0
x[x == 1] <- 1
x1 <- x[, seq(1, 2*n, 2)]
x2 <- x[, seq(2, 2*n, 2)]
x.g <- x1 + x2

rownames(x.g) <- snps
colnames(x.g) <- donors

file.out %>% saveRDS(x.g, file=.)
```
