## Supplemental File 2 for "Probabilistic classification of gene-by-treatment interactions on molecular count phenotypes": format_kin.R

```
suppressPackageStartupMessages(library(tidyverse))

args <- commandArgs(TRUE)
data.file <- as.character(args[1])
kin.file <- as.character(args[2])
out.file <- as.character(args[3])

# read in the data
data.list <- data.file %>% readRDS

# read in the kinship matrix
A <- kin.file %>% read.delim(row.names=1) 
dna <- data.list[[1]]$data$dna
dna <- dna[seq(1, length(dna), 2)]
A <- A[dna, dna]

A %>% as.matrix %>% saveRDS(file=out.file)
```
