## Supplemental File 2 for "Probabilistic classification of gene-by-treatment interactions on molecular count phenotypes": get_anno.R

```
suppressPackageStartupMessages(library(tidyverse))
suppressPackageStartupMessages(library(readxl))

args <- commandArgs(TRUE)
geno.dir <- as.character(args[1])
pheno.file <- as.character(args[2])
id.file <- as.character(args[3])
sample.file <- as.character(args[4])
out.file <- as.character(args[5])

# read in files
pheno <- pheno.file %>% read.delim
id <- id.file %>% readRDS
sample <- sample.file %>% readRDS

# create a list of formatted data
data.list.list <- list()
count <- 0

for (i in seq_len(22)) {

    geno.file <- file.path(
        geno.dir, paste0("geno.chr", i, ".rds"))
    geno <- geno.file %>% readRDS()
    sel <- grep(
        paste0("^chr", i, ":"), id$snp_id)

    if (length(sel) == 0) next

    for (j in seq_along(sel)) {

        index <- sel[j]
        row <- as.data.frame(id[index, ])
        feat.id <- row$gene_id
        snp.id <- row$snp_id

        if (all(pheno$feature_id != feat.id)) next

        pheno.df <- data.frame(
            sample=colnames(pheno)[-1],
            pheno=unlist(
                pheno[pheno$feature_id == feat.id, -1]))

        geno.df <- data.frame(
            donor=colnames(geno),
            geno=unlist(geno[rownames(geno) == snp.id, ]))
        geno.df <- merge(
            geno.df, sample, by.x="donor", by.y="donor")

        d <- merge(
            pheno.df, geno.df, by.x="sample", by.y="sample")
        d$treatment <- sapply(
            d$sample,
            function(x) as.numeric(
                            unlist(strsplit(x, "_"))[3] == "1"))

        d <- d[, c(1, 3, 5, 2, 4, 6)]

        # make the treatment coding more interpretable
        treatment <- rep(0, nrow(d))
        treatment[grep("_2$", d$sample)] <- 1
        d$treatment <- treatment

        # if (sum(d$geno == 1) == 0 || sum(d$geno == 2) == 0) {
        #     next
        # }

        data.list <- list(
            data=d,
            feat.id=feat.id,
            snp.id=snp.id)
        count <- count + 1
        data.list.list[[count]] <- data.list
        rm(d); rm(data.list)

    }

}

# save file
saveRDS(data.list.list, file=out.file)
```
