## Supplemental File 2 for "Probabilistic classification of gene-by-treatment interactions on molecular count phenotypes": get_fc.R

```
suppressPackageStartupMessages(library(tidyverse))
suppressPackageStartupMessages(library(Rsubread))

args <- commandArgs(TRUE)
in.file <- as.character(args[1])
gtf.file <- as.character(args[2]) 
out.file <- as.character(args[3])

fc <- featureCounts(
    files=in.file,
    GTF.featureType="exon", GTF.attrType="gene_id",
    annot.ext=gtf.file, isGTFAnnotationFile=TRUE,
    isPairedEnd=TRUE, nthreads=4)

saveRDS(fc, file=out.file)
```
