## Supplemental File 2 for "Probabilistic classification of gene-by-treatment interactions on molecular count phenotypes": get_id.R

```
suppressPackageStartupMessages(library(tidyverse))
suppressPackageStartupMessages(library(readxl))

args <- commandArgs(TRUE)
in.file <- as.character(args[1])
out.file <- as.character(args[2])

# read in data for CHIR
sheet.c <- 1 # control
sheet.t <- 3 # treated
hit.c <- in.file %>% read_excel(sheet=sheet.c)
hit.t <- in.file %>% read_excel(sheet=sheet.t)

# focus on autosomes
hit.c <- hit.c[hit.c$gene_chr %in% 1:22, ]
hit.t <- hit.t[hit.t$gene_chr %in% 1:22, ]
hit.c <- hit.c %>%
    mutate(id=paste(gene_id, snp_id, sep="-")) 
hit.t <- hit.t %>%
    mutate(id=paste(gene_id, snp_id, sep="-"))
merged <- merge(
    hit.c, hit.t, by.x="id", by.y="id",
    all.x=TRUE, all.y=TRUE)

# create a data frame containing
# id, gene_id, snp_id
hit <- data.frame(id=merged$id)
hit$gene_id <- sapply(
    merged$id, function(x) unlist(strsplit(x, "-"))[1])
hit$snp_id <- sapply(
    merged$id, function(x) unlist(strsplit(x, "-"))[2])

saveRDS(hit, file=out.file)
```
