## Supplemental File 2 for "Probabilistic classification of gene-by-treatment interactions on molecular count phenotypes": get_input_for_aa.R

```
suppressPackageStartupMessages(library(tidyverse))

args <- commandArgs(TRUE)
vst.file <- as.character(args[1])
data.file <- as.character(args[2])
out.file <- as.character(args[3])

# read in input files
data.list <- data.file %>% readRDS
vst.mat <- vst.file %>% readRDS

out.list <- list()
col.name <- c("sample", "donor", "dna", "y", "g", "t")
count <- 0

for (i in seq_along(data.list)) {

    d <- data.list[[i]]$data
    colnames(d) <- col.name

    if (sum(d$g == 1) == 0 || sum(d$g == 2) == 0) {
        next
    }

    feat.id <- data.list[[i]]$feat.id
    snp.id <- data.list[[i]]$snp.id

    # get log-transformed data
    pheno.vec <- vst.mat[rownames(vst.mat) == feat.id, ]
    names(pheno.vec) <- paste0("X", names(pheno.vec))

    # note that donors in the other condition will be removed
    d <- merge(d, pheno.vec, by.x=1, by.y=0)

    # rearrange columns
    d <- d[, c(1:3, 7, 5:6)]
    colnames(d) <- col.name

    output <- list(
        data=d,
        feat.id=feat.id,
        snp.id=snp.id)
    count <- count + 1
    out.list[[count]] <- output
    rm(d); rm(output)

}

saveRDS(out.list, file=out.file)
```
