## Supplemental File 2 for "Probabilistic classification of gene-by-treatment interactions on molecular count phenotypes": get_input_for_bms.R

```
suppressMessages(library(tidyverse))
suppressMessages(library(readxl))

args <- commandArgs(TRUE)
data.file <- as.character(args[1])
hit.file <- as.character(args[2])
cnt.file <- as.character(args[3])
out.file <- as.character(args[4])

# read in the data file
data.list <- data.file %>% readRDS

# read in the eqtl hits    
sheet <- 2 # CT99, CHIR
hit <- hit.file %>% read_excel(sheet=sheet)

# read in log-transformed count data
cnt.mat <- cnt.file %>% readRDS

# select genes and snps on autosomes
hit <- hit %>%
    filter(chromosome %in% 1:22) %>%
    dplyr::select(c(gene_id, snp_id))

# get feature and snp ids from the input data
feat.vec <- data.list %>%
    map_chr(pluck("feat.id"))
snp.vec <- data.list %>%
    map_chr(pluck("snp.id"))
id <- data.frame(
    feat=feat.vec,
    snp=snp.vec,
    index=seq_along(data.list))

# merge the data frames
joined <- hit %>%
    inner_join(
        id, join_by(gene_id == feat, snp_id == snp)) %>%
    as.data.frame %>%
    `colnames<-`(c("feat", "snp", "index"))

# create a list object
out.list <- vector("list", nrow(joined)) 

for (i in seq_len(nrow(joined))) {

    index <- joined$index[i]

    l <- data.list[[index]]
    feat.id <- l$feat.id
    snp.id <- l$snp.id
    d <- l$data
    d$sample <- gsub("X", "", d$sample)
    
    cnt.vec <- cnt.mat[feat.id, ]
    cnt.vec <- cnt.vec[d$sample]

    d <- data.frame(
        y=cnt.vec,
        g=d$geno,
        t=d$treatment,
        dna=d$dna)

    # replace the data
    l$data <- d
    out.list[[i]] <- l
    
}

saveRDS(out.list, file=out.file)
```
