## Supplemental File 2 for "Probabilistic classification of gene-by-treatment interactions on molecular count phenotypes": get_llk.R

```
suppressPackageStartupMessages(library(tidyverse))
suppressPackageStartupMessages(library(rstan))

args <- commandArgs(TRUE)
data.file <- as.character(args[1])
seed <- as.integer(args[2])
out.file <- as.character(args[3])

rstan_options(auto_write=TRUE)

# read in the data to get the number of samples
data.list <- data.file %>% readRDS

# load stan model
nl.stan <- stan_model(file="nl.stan")
lm.stan <- stan_model(file="lm.stan")

# get the number of samples
n.data <- length(data.list)

llk <- matrix(NA, nrow=n.data, ncol=3) %>%
    as.data.frame %>%
    `colnames<-`(c("lm", "nl", "anova"))

for (i in seq_len(n.data)) {

    data <- data.list[[i]]$data
    y <- data$y
    n <- length(y)
    g <- data$g

    # fit anova model    
    if (all(table(g) > 0)) {
        g1 <- as.numeric(g == 1)
        g2 <- as.numeric(g == 2)
        fit.anova <- lm(y ~ g1 + g2)
        llk.anova <- logLik(fit.anova)
    } else {
        llk.anova <- NA
    }    
    
    # fit anova model
    fit.anova <- lm(y ~ g1 + g2)
    llk.anova <- logLik(fit.anova)
    
    # fit linear model
    fit.lm <- lm(y ~ g)
    llk.lm <- logLik(fit.lm)
    
    # fit nonlinear model
    x1 <- (1 - g/2)
    x2 <- (g/2)
    data.stan <- list(
        N=n, y=y,
        x1=x1, x2=x2)
    fit.nl <- optimizing(
        object=nl.stan,
        data=data.stan,
        seed=seed,
        algorithm="LBFGS")
    llk.nl <- fit.nl$value
    
    llk[i, ] <- c(llk.lm, llk.nl, llk.anova)
}

# save the results
saveRDS(llk, file=out.file)
```
