## Supplemental File 2 for "Probabilistic classification of gene-by-treatment interactions on molecular count phenotypes": get_pc_for_aa.R

```
suppressPackageStartupMessages(library(tidyverse))

args <- commandArgs(TRUE)
file.in <- as.character(args[1])
file.out <- as.character(args[2])

vst.mat <- file.in %>% readRDS

pca <- vst.mat %>%
    t %>%
    prcomp

saveRDS(pca, file=file.out)
```
