## Supplemental File 2 for "Probabilistic classification of gene-by-treatment interactions on molecular count phenotypes": get_phenomat.R

```
# modified from 00.prep.eqtl.data.R
suppressPackageStartupMessages(library(tidyverse))
suppressPackageStartupMessages(library(data.table))

args <- commandArgs(TRUE)
fc.dir <- as.character(args[1])
sample.file <- as.character(args[2])
gene.file <- as.character(args[3])
cond <- as.character(args[4])
out.file <- as.character(args[5])

if (cond == "Veh") {
    pattern <- "_1$"
} else if (cond == "CT99") {
    pattern <- "_2$"
} else if (cond == "Wnt3a") {
    pattern <- "_5$"
}

sample <- sample.file %>% read.delim(header=FALSE)
sample.vec <- sample %>%
    pull(2) %>%
    gsub("X", "", .) %>%
    grep(pattern=pattern, x=., value=TRUE)

file.vec <- file.path(
    fc.dir, paste0(sample.vec, ".rds"))

# read in count data
fc.mat <- file.vec %>%
    map(readRDS) %>%
    map("counts") %>%
    do.call("cbind", .)

colnames(fc.mat) <- colnames(fc.mat) %>%
    gsub("\\.sorted\\.bam", "", .)

# get protein coding genes and lncRNAs only
gene <- fread(gene.file, data.table=FALSE)
gene <- gene %>%
    filter(gene_type %in% c("protein_coding", "lncRNA"))
fc.mat <- fc.mat[rownames(fc.mat) %in% gene$gene_id, ]

saveRDS(fc.mat, file=out.file)
```
