## Supplemental File 2 for "Probabilistic classification of gene-by-treatment interactions on molecular count phenotypes": get_resid_for_aa.R

```
suppressPackageStartupMessages(library(tidyverse))

args <- commandArgs(TRUE)
data.file <- as.character(args[1])
pca.file <- as.character(args[2])
num.pc <- as.integer(args[3])
out.file <- as.character(args[4])

data.list <- data.file %>% readRDS
pca <- pca.file %>% readRDS
pca.mat <- pca$x
rownames(pca.mat) <- paste0("X", rownames(pca.mat))

for (i in seq_along(data.list)) {
    d <- data.list[[i]]$data
    d <- merge(d, pca.mat[, 1:10], by.x=1, by.y=0)
    # get residuals
    frml <- paste0(
        "y ~ ",
        paste0(paste0("PC", seq_len(num.pc)),
               collapse=" + ")) 
    d$y <- residuals(lm(frml, data=d))
    # remove pcs
    d <- d[, 1:6]
    data.list[[i]]$data <- d
}    

saveRDS(data.list, file=out.file)
```
