## Supplemental File 2 for "Probabilistic classification of gene-by-treatment interactions on molecular count phenotypes": get_samplemap.R

```
# note that the tfam files are the same for all chromosomes
suppressPackageStartupMessages(library(tidyverse))
suppressPackageStartupMessages(library(readxl))

args <- commandArgs(TRUE)
info.file <- as.character(args[1])
donor.file <- as.character(args[2])
out.file <- as.character(args[3])

# read in files
donor <- donor.file %>% read.table
info <- info.file %>% read.delim(header=FALSE)

# get donor mapping
samplemap <- merge(info, donor[, 1:2], by.x="V1", by.y="V2")
colnames(samplemap) <- c("dna", "sample", "donor")

saveRDS(samplemap, file=out.file)
```
