## Supplemental File 2 for "Probabilistic classification of gene-by-treatment interactions on molecular count phenotypes": get_sign_for_mcmc_bs.R

```
suppressPackageStartupMessages(library(tidyverse))
suppressPackageStartupMessages(library(readxl))
suppressPackageStartupMessages(library(classifygxt))

args <- commandArgs(TRUE)
prev.file <- as.character(args[1])
data.file <- as.character(args[2])
in.dir <- as.character(args[3])
pp.file <- as.character(args[4])
order.file <- as.character(args[5])

# read in the previous results
prev <- read_excel(prev.file, sheet=2) %>%
    as.data.frame

# read in the input data file
data.list <- data.file %>% readRDS

file.vec <- list.files(
    in.dir, pattern="^res", full.names=TRUE) %>%
    grep("rds$", ., value=TRUE)

pp <- file.vec %>%
    map(
        function(x) readRDS(x) %>%
                    get_sign) %>%
    do.call("rbind", .) %>%
    as.data.frame

# get the order of the pairs by $p$-values
feat.id.vec <- sapply(data.list, function(x) x$feat.id)
snp.id.vec <- sapply(data.list, function(x) x$snp.id)
id.vec <- paste(feat.id.vec, snp.id.vec, sep="-")
prev.id.vec <- paste(prev$gene_id, prev$snp_id, sep="-")
order.vec <- match(prev.id.vec, id.vec) %>% na.omit

saveRDS(pp, file=pp.file)
saveRDS(order.vec, file=order.file)
```
