## Supplemental File 2 for "Probabilistic classification of gene-by-treatment interactions on molecular count phenotypes": optimize_hp.R

```
suppressPackageStartupMessages(library(tidyverse))
suppressPackageStartupMessages(library(classifygxt))

args <- commandArgs(TRUE)
data.file <- as.character(args[1])
tu.lambda.file <- as.character(args[2])
seed <- as.integer(args[3])
method <- as.character(args[4])
lb <- as.integer(args[5])
ub <- as.integer(args[6])
step <- as.integer(args[7])
start <- as.integer(args[8])
size <- as.integer(args[9])
ranef <- as.character(args[10])
out.dir <- as.character(args[11])

# read in the data
data.list <- data.file %>% readRDS

# create a matrix of hyperparamers
hp.vec <- (seq(lb, ub, step) / 100)  %>%
    format(round(., 2), nsmall = 2) %>%
    as.character
n.hp <- length(hp.vec)
hp.mat <- cbind(
    phig=rep(hp.vec, each=n.hp^2),
    phit=rep(rep(hp.vec, times=n.hp), each=n.hp),
    phigt=rep(hp.vec, times=n.hp^2))

end <- start + size - 1
end <- min(end, nrow(hp.mat))

# specify model prior
p.m <- rep(1/8, 8)

computation <- "map.lap"

if (ranef == "ranef") {
    ranef <- TRUE
    tu.lambda <- tu.lambda.file %>% readRDS
} else if (ranef == "noranef") {
    ranef <- FALSE
    tu.lambda <- NULL
}

if (method == "nl") {
    fn <- "nonlinear"
    rint <- FALSE
} else if (method == "lm") {
    fn <- "linear"
    rint <- FALSE
} else if (method == "rint") {
    fn <- "linear"
    rint <- TRUE
}

data.list <- data.list %>%
    map(function(x)
        list(
            y=x$data$y, g=x$data$g,
            t=x$data$t, subject=x$data$dna))

for (k in start:end) {

    phi.vec <- hp.mat[k, ]
    phi <- phi.vec %>% as.numeric

    subdir <- file.path(
        out.dir,
        paste0("phi", paste(phi.vec, collapse="_")))
    if (!dir.exists(subdir)) dir.create(subdir, recursive=TRUE)
    out.file <- file.path(
        subdir, "fit.rds")
    if (file.exists(out.file)) {
        warning(
            paste("The file", out.file, "already exists."))
        next
    }

    res.list <- lapply(
        X=data.list,
        FUN=do_bms,
        p.m=p.m, phi=phi,
        fn=fn, rint=rint, method=computation,
        tu.lambda=tu.lambda)

    saveRDS(res.list, file=out.file)

}

paste0(
    "complete",
    str_pad(start, 4, pad="0"), "_", size, ".txt") %>%
    file.path(out.dir, .) %>%
    file.create
```
