## Supplemental File 2 for "Probabilistic classification of gene-by-treatment interactions on molecular count phenotypes": Snakefile

```
with open(config["sample"], "r") as file: sample_id = [line.split()[1].replace("X", "").replace("-", "_") for line in file]
chrom = range(1, 23, 1)
method = ["nl", "lm", "rint"]
k_mat = ["id", "kin"]
lb = 25
ub = 300
step = 25
num_pair = 98
numbers = [x / 100.00 for x in range(lb, ub + 1, step)]
phi1 = ["{:1.2f}".format(x) for x in numbers]
phi2 = ["{:1.2f}".format(x) for x in numbers]
phi3 = ["{:1.2f}".format(x) for x in numbers]
cond = ["Veh", "CT99"]
pc = [10]
size = 10
start = [format(number, "04") for number in range(1, num_pair + 1, size)] # number of pairs
hp_size = 100 # batch size
hp_start = [format(number, "04") for number in range(1, len(numbers)**3 + 1, hp_size)] # number of hp

rule all:
    input:
        expand(config["dir"]
               + "post/{k_mat}/pc{pc}/{method}/"
               + "pp.sign.rds",
               k_mat = k_mat, pc = pc, method = method),
        expand(config["dir"]
               + "post/{k_mat}/pc{pc}/{method}/"
               + "pp.rds",
               k_mat = k_mat, pc = pc, method = method),
        expand(config["dir"]
               + "post/{k_mat}/pc{pc}/{method}/"
               "fit/complete{start}_{size}.txt",
               k_mat = k_mat, pc = pc, method = method,
               start = start, size = size),
        expand(config["dir"]
               + "optim/{k_mat}/pc{pc}/{method}/"
               + "pp.sign.rds",
               k_mat = k_mat, pc = pc, method = method),
        expand(config["dir"]
               + "optim/{k_mat}/pc{pc}/{method}/"
               + "pp.rds",
               k_mat = k_mat, pc = pc, method = method),
        expand(config["dir"]
               + "optim/{k_mat}/pc{pc}/{method}/"
               "ln.p.y.rds",
               k_mat = k_mat, pc = pc, method = method),
        expand(config["dir"]
               + "optim/{k_mat}/pc{pc}/{method}/"
               "complete{start}_{size}.txt",
               k_mat = k_mat, pc = pc, method = method,
               start = hp_start, size = hp_size),
        expand(config["dir"]
               + "aa/llk/pc{pc}/{cond}/"
               + "llk.rds",
               pc = pc, cond = cond),
        expand(config["dir"] + "fc/{sample_id}.rds",
               sample_id = sample_id),

rule get_sign_for_mcmc_bs:
    input:
        prev = config["reqtl"],
        data = config["dir"]
        + "pre/rsd/pc{pc}/data.list.rds",
        fit = expand(config["dir"]
                     + "post/{{k_mat}}/"
                     "pc{{pc}}/{{method}}/fit/"
                     "complete{start}_{size}.txt",
                     start = start, size = size),
    output:
        pp = config["dir"]
        + "post/{k_mat}/pc{pc}/{method}/pp.sign.rds",
        order = config["dir"]
        + "post/{k_mat}/pc{pc}/{method}/order.vec.rds",
    shell:            
        "module add r/4.1.3; "
        "Rscript get_sign_for_mcmc_bs.R"
        " {input.prev}"
        " {input.data}"
        " " + config["dir"]
        + "post/{wildcards.k_mat}/pc{wildcards.pc}/"
        "{wildcards.method}/fit"
        " {output.pp}"
        " {output.order}"

rule get_sign_for_map_lap:
    input:
        prev = config["reqtl"],
        data = config["dir"]
        + "pre/rsd/pc{pc}/data.list.rds",
        lnpy = config["dir"]
        + "optim/{k_mat}/pc{pc}/{method}/ln.p.y.rds",
    output:
        pp = config["dir"]
        + "optim/{k_mat}/pc{pc}/{method}/pp.sign.rds",
        order = config["dir"]
        + "optim/{k_mat}/pc{pc}/{method}/order.vec.rds",
    shell:            
        "module add r/4.1.3; "
        "Rscript get_sign_for_map_lap.R"
        " {input.prev}"
        " {input.data}"
        " {input.lnpy}"
        " " + config["dir"]
        + "optim/{wildcards.k_mat}/"
        "pc{wildcards.pc}/{wildcards.method}"
        " {output.pp}"
        " {output.order}"
        
rule get_pp_for_mcmc_bs:
    input:
        expand(config["dir"]
               + "post/{{k_mat}}/pc{{pc}}/{{method}}/fit/"
               "complete{start}_{size}.txt",
               start = start, size = size),
    output:
        config["dir"]
        + "post/{k_mat}/pc{pc}/{method}/pp.rds",
    shell:
        "module add r/4.1.3; "
        "Rscript get_pp_for_mcmc_bs.R"
        " " + config["dir"]
        + "post/{wildcards.k_mat}/pc{wildcards.pc}/"
        "{wildcards.method}/fit"
        " {output}"

rule get_pp_for_map_lap:
    input:
        fit = config["dir"]
        + "optim/{k_mat}/pc{pc}/{method}/ln.p.y.rds",
        data = config["dir"]
        + "pre/rsd/pc{pc}/data.list.rds"
    output:
        config["dir"]
        + "optim/{k_mat}/pc{pc}/{method}/pp.rds",
    shell:
        "module add r/4.1.3; "
        "Rscript get_pp_for_map_lap.R"
        " {input.fit}"
        " {input.data}"
        " " + config["dir"]
        + "optim/{wildcards.k_mat}/"
        "pc{wildcards.pc}/"
        "{wildcards.method}"
        " {output}"
        
rule post:
    input:
        data = config["dir"]
        + "pre/rsd/pc{pc}/data.list.rds",
        tu_lambda = config["dir"]
        + "pre/tu.lambda.{k_mat}.rds",
        fit = config["dir"]
        + "optim/{k_mat}/pc{pc}/{method}/ln.p.y.rds",
    output:
        config["dir"]
        + "post/{k_mat}/pc{pc}/{method}/"
        + "fit/complete{start}_{size}.txt",
    shell:
        "module add gcc/11.2.0; "
        "module add r/4.1.3; "
        "Rscript do_mcmc_bs.R"
        " {input.data}"
        " {input.tu_lambda}"
        " {input.fit}"
        " 1" # seed
        " 4" # cores
        " {wildcards.method}"
        " ranef"
        " opt"
        " sample"
        " {wildcards.start}"
        " {wildcards.size}"
        " " + config["dir"]
        + "post/{wildcards.k_mat}/pc{wildcards.pc}/"
        "{wildcards.method}/fit"
        
rule get_marginal_llk:
    input:
        expand(config["dir"]
               + "optim/{{k_mat}}/pc{{pc}}/{{method}}/"
               "complete{start}_{size}.txt",
               start = hp_start,
               size = hp_size)
    output:
        config["dir"]
        + "optim/{k_mat}/pc{pc}/{method}/ln.p.y.rds"
    shell:
        "module add r/4.1.3; "
        "Rscript get_marginal_llk.R"
        " " + config["dir"]
        + "optim/{wildcards.k_mat}/pc{wildcards.pc}/"
        "{wildcards.method}"
        " 25" # lb
        " 300" # ub
        " 25" # step
        " {output}"

rule optimize_hp:
    input:
        data = config["dir"]
        + "pre/rsd/pc{pc}/data.list.rds",
        tu_lambda = config["dir"]
        + "pre/tu.lambda.{k_mat}.rds",
    output:
        config["dir"]
        + "optim/{k_mat}/pc{pc}/{method}/"
        "complete{start}_{size}.txt"
    shell:
        "module add r/4.1.3; "
        "Rscript optimize_hp.R"
        " {input.data}"
        " {input.tu_lambda}"
        " 1"
        " {wildcards.method}"
        " 25" # lb
        " 300" # ub
        " 25" # step 
        " {wildcards.start}"
        " {wildcards.size}"
        " ranef"
        " " + config["dir"]
        + "optim/{wildcards.k_mat}/pc{wildcards.pc}/"
        "{wildcards.method}"

rule get_tu_lambda_for_id:
    input:
        config["dir"] + "pre/rsd/pc10/"
        + "data.list.rds", # "pre/data.list.rds"
    output:
        config["dir"] + "pre/tu.lambda.id.rds"
    shell:
        "module add r/4.1.3; "
        "Rscript get_tu_lambda.R"
        " {input}"
        " NA"
        " {output}"
 
rule get_tu_lambda_for_kin:
    input:
        data = config["dir"] + "pre/rsd/pc10/"
        + "data.list.rds", # "pre/data.list.rds"
        kin = config["dir"] + "pre/kin.mat.rds"
    output:
        config["dir"] + "pre/tu.lambda.kin.rds"
    shell:
        "module add r/4.1.3; "
        "Rscript get_tu_lambda.R"
        " {input.data}"
        " {input.kin}"
        " {output}"

rule format_kin:
    input:
        data = config["dir"] + "pre/"
        + "data.list.rds",
        kin = config["kin"],
    output:
        config["dir"] + "pre/kin.mat.rds"
    shell:
        "module add r/4.1.3; "
        "Rscript format_kin.R"
        " {input.data}"
        " {input.kin}"
        " {output}"

rule get_resid_for_bms:
    input:
        data = config["dir"] + "pre/data.list.rds",
        pca = config["dir"] + "pre/pca.rds",
    output:
        config["dir"] + "pre/rsd/pc{pc}/data.list.rds"
    shell:
        "module add r/4.1.3; "
        "Rscript get_resid_for_bms.R"
        " {input.data}"
        " {input.pca}"
        " {wildcards.pc}"
        " {output}"

rule get_pc_for_bms:
    input:
        config["dir"] + "pre/log.mat.rds"
    output:
        config["dir"] + "pre/pca.rds",
    shell:
        "module add r/4.1.3; "
        "Rscript get_pc_for_bms.R"
        " {input}"
        " {output}"

rule get_input_for_bms:
    input:
        data = config["dir"] + "input/data.list.rds",
        cnt = config["dir"] + "pre/log.mat.rds",
        hit = config["reqtl"]
    output:
        config["dir"] + "pre/data.list.rds",
    shell:
        "module add r/4.1.3; "
        "Rscript get_input_for_bms.R"
        " {input.data}"
        " {input.hit}"
        " {input.cnt}"
        " {output}"

rule combine_phenomat:
    input:
        expand(config["dir"]
               + "aa/log.{cond}.mat.rds",
               cond = cond)
    output:
        config["dir"] + "pre/log.mat.rds"
    shell:
        "module add r/4.1.3; "
        "Rscript combine_phenomat.R"
        " {input}"
        " {output}"

rule get_llk:
    input:
        config["dir"] + "aa/rsd/pc{pc}/{cond}/"
        + "data.list.rds"
    output:
        config["dir"]
        + "aa/llk/pc{pc}/{cond}/"
        + "llk.rds"
    shell:
        "module add r/4.1.3; "
        "Rscript get_llk.R"
        " {input}"
        " 1" # seed 
        " {output}"

rule get_resid_for_aa:
    input:
        data = config["dir"] + "aa/input/{cond}/"
        + "data.list.rds",
        pca = config["dir"] + "aa/pca.{cond}.rds",
    output:
        config["dir"] + "aa/rsd/pc{pc}/{cond}/"
        + "data.list.rds"
    shell:
        "module add r/4.1.3; "
        "Rscript get_resid_for_aa.R"
        " {input.data}"
        " {input.pca}"
        " {wildcards.pc}"
        " {output}"

rule get_input_for_aa:
    input:
        data = config["dir"] + "input/data.list.rds",
        vst = config["dir"] + "aa/log.{cond}.mat.rds"
    output:
        config["dir"] + "aa/input/{cond}/"
        + "data.list.rds"
    shell:
        "module add r/4.1.3; "
        "Rscript get_input_for_aa.R"
        " {input.vst}"
        " {input.data}"
        " {output}"

rule get_pc_for_aa:
    input:
        config["dir"] + "aa/log.{cond}.mat.rds"
    output:
        config["dir"] + "aa/pca.{cond}.rds"
    shell:
        "module add r/4.1.3; "
        "Rscript get_pc_for_aa.R"
        " {input}"
        " {output}"

rule transform_pheno:
    input:
        cnt = config["dir"] + "aa/cnt.{cond}.mat.rds",
        pheno = config["pheno"],
    output:
        config["dir"] + "aa/log.{cond}.mat.rds"
    shell:
        "module add r/4.1.3; "
        "Rscript transform_pheno.R"
        " {input.cnt}"
        " {input.pheno}"
        " {output}"

rule get_phenomat:
    input:
        sample = config["sample"],
        gene = config["genename"],
        fc = expand(config["dir"]
                    + "fc/{sample_id}.rds",
                    sample_id = sample_id)
    output:
        config["dir"] + "aa/cnt.{cond}.mat.rds",
    shell:
        "module add r/4.1.3; "
        "Rscript get_phenomat.R"
        " " + config["dir"] + "fc"
        " {input.sample}"
        " {input.gene}"
        " {wildcards.cond}"
        " {output}"

rule get_fc:
    input:
        bam = config["bam_dir"]
        + "{sample_id}/{sample_id}.sorted.bam",
        gtf = config["gtf"]
    output:
        config["dir"] + "fc/{sample_id}.rds"
    shell:
        "module add r/4.1.3; "
        "mkdir -p " + config["dir"] + "fc; "
        "Rscript get_fc.R"
        " {input.bam}"
        " {input.gtf}"
        " {output}"

rule get_anno:
    input:
        geno = expand(config["dir"]
                      + "geno/geno.chr{chrom}.rds",
                      chrom = chrom),
        pheno = config["pheno"],
        hit = config["dir"] + "input/id.rds",
        sample = config["dir"] + "input/samplemap.rds",
    output:
        config["dir"] + "input/data.list.rds"
    shell:
        "module add r/4.1.3; "
        "Rscript get_anno.R"
        " " + config["dir"] + "geno"
        " {input.pheno}"
        " {input.hit}"
        " {input.sample}"
        " {output}"
        
rule get_samplemap:
    input:
        sample = config["sample"],
        donor = config["dir"] + "plink/wnt.chr1.tfam",
    output:
        config["dir"] + "input/samplemap.rds"
    shell:
        "module add r/4.1.3; "
        "Rscript get_samplemap.R"
        " {input.sample}"
        " {input.donor}"
        " {output}"

rule get_id:
    input:
        config["eqtl"]
    output:
        config["dir"] + "input/id.rds"
    shell:
        "module add r/4.1.3; "
        "Rscript get_id.R"
        " {input}"
        " {output}"

rule format_geno:
    input:
        tfam = config["dir"]
        + "plink/wnt.chr{chrom}.tfam",
        tped = config["dir"]
        + "plink/wnt.chr{chrom}.tped",
    output:
        config["dir"] + "geno/geno.chr{chrom}.rds",
    shell:
        "module add r/4.1.3; "
        "Rscript format_geno.R"
        " {input.tfam}"
        " {input.tped}"
        " {output}"
                
rule run_plink:
    input:
        config["indir"] + "Wnt_all.{chrom}.wo.lowCT.bed",
        config["indir"] + "Wnt_all.{chrom}.wo.lowCT.bim",
        config["indir"] + "Wnt_all.{chrom}.wo.lowCT.fam",
    output:
        config["dir"] + "plink/wnt.chr{chrom}.tped",
        config["dir"] + "plink/wnt.chr{chrom}.tfam",
    shell:
        "module load plink/1.90b3; "
        "plink --bfile " + config["indir"] +
        "Wnt_all.{wildcards.chrom}.wo.lowCT"
        " --recode 12 transpose --out "
        + config["dir"] + "plink/"
        "wnt.chr{wildcards.chrom}"
```
