## Supplemental File 2 for "Probabilistic classification of gene-by-treatment interactions on molecular count phenotypes": transform_pheno.R

```
suppressPackageStartupMessages(library(tidyverse))

args <- commandArgs(TRUE)
cnt.file <- as.character(args[1])
pheno.file <- as.character(args[2])
out.file <- as.character(args[3])

cnt.mat <- cnt.file %>% readRDS
pheno <- pheno.file %>% read.delim

# scale according to Palowitch et al. (2018)
# sample i = 1, ..., n
# gene j = 1, ..., T
T <- nrow(cnt.mat)
n <- ncol(cnt.mat)
li <- colSums(cnt.mat)
xbar <- sum(li) / (T * n)
cnt.mat <- sweep(cnt.mat, 2, li, "/") * T * xbar

# use the same set of genes as in Matoba et al. (2023)
keep.vec <- pheno$feature_id

cnt.mat <- cnt.mat[keep.vec, ]
cnt.mat <- log(cnt.mat + 1)

saveRDS(cnt.mat, file=out.file)
```
