## Supplemental Figures for "Probabilistic classification of gene-by-treatment interactions on molecular count phenotypes"

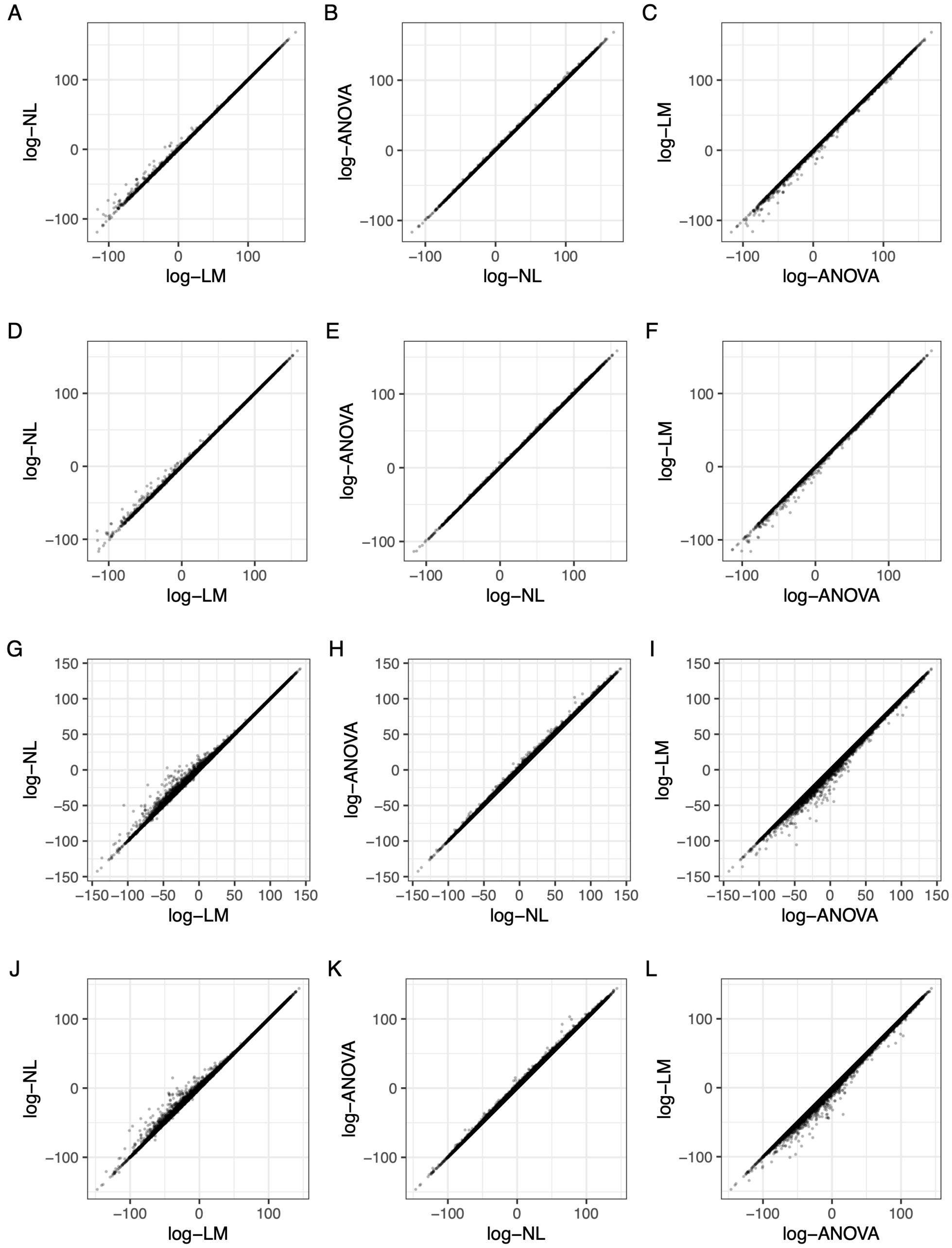


Figure S1. **Assessing the allelic additivity assumption in hNPCs. A.** Scatterplots comparing the maximized likelihood between nonlinear and linear regression for 3073 gene-SNP pairs under the control condition. **B.** The same as in **A** but between the ANOVA model and nonlinear regression. **C.** The same as in **A** but between the ANOVA model and linear regression. **D.** The same as in **A** but under the treated condition. **E.** The same as in **B** but under the treated condition. **F.** The same as in **C** but under the treated condition. **G.** Scatterplots comparing the maximized likelihood between nonlinear and linear regression for 42576 cCRE-SNP pairs under the control condition. **H.** The same as in **G** but between the ANOVA model and nonlinear regression. **I.** The same as in **G** but between the ANOVA model and linear regression. **J.** The same as in **G** but under the treated condition. **K.** The same as in **H** but under the treated condition. **L.** The same as in **I** but under the treated condition.

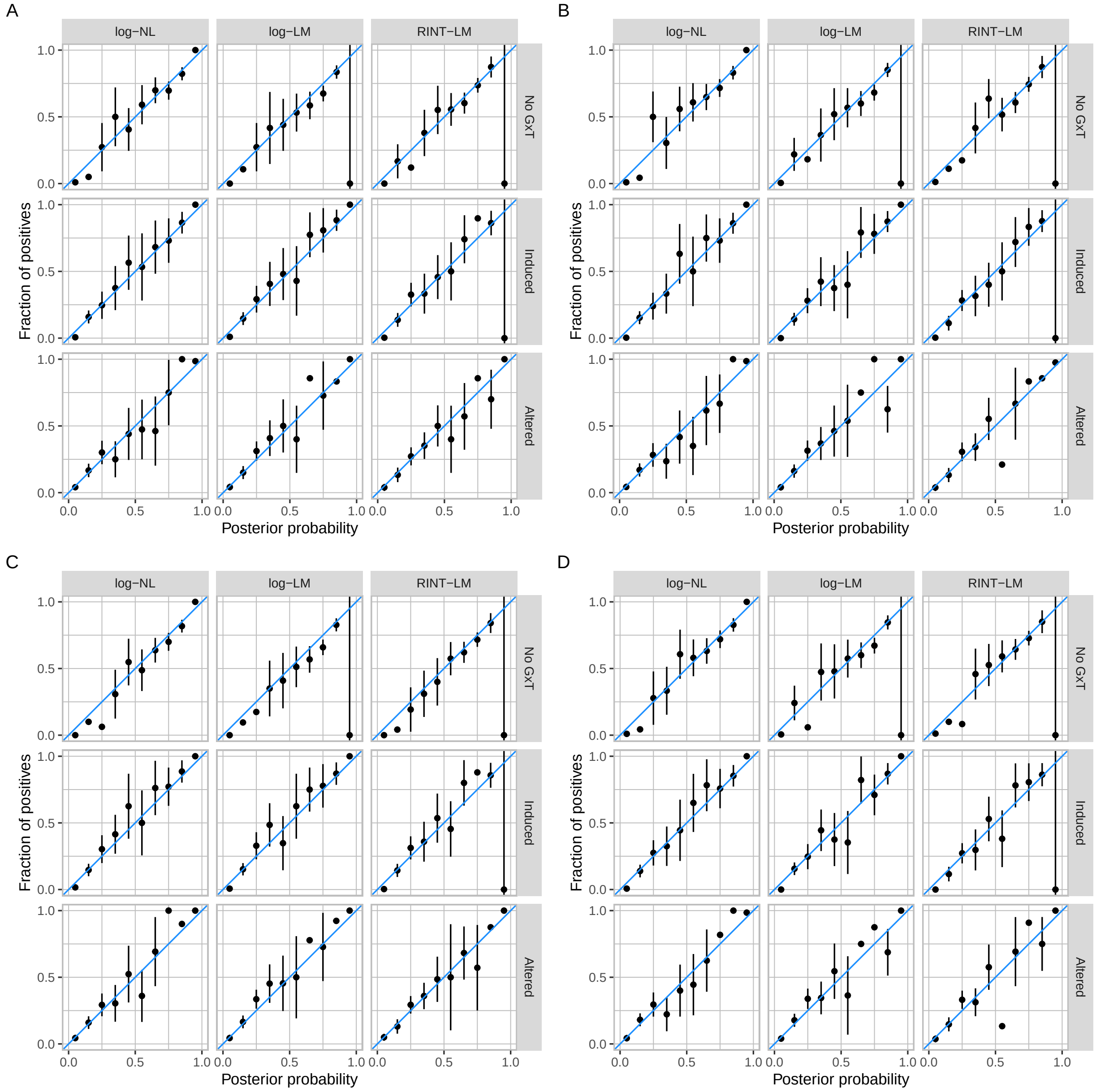


Figure S2. **Calibration of BMS with** log**-NL,** log**-LM, and RINT-LM for the "no G×T", "induced", and "altered" categories using MCMC and bridge sampling.** The $x$- and $y$-axis represent the posterior probability and the fraction of the corresponding events, respectively. The results from 800 simulations are grouped into ten equally-spaced bins. The vertical bars represent the standard errors assuming a binomial distribution. Shown are results from analysis without random effect, which we call scenario 1 (**A**), those with donor random effect in model fitting but not in data generation (scenario 2) (**B**), those with donor random effect in data generation but not in model fitting (scenario 3) (**C**), and those with donor effect in both data generation and model fitting (scenario 4) (**D**).

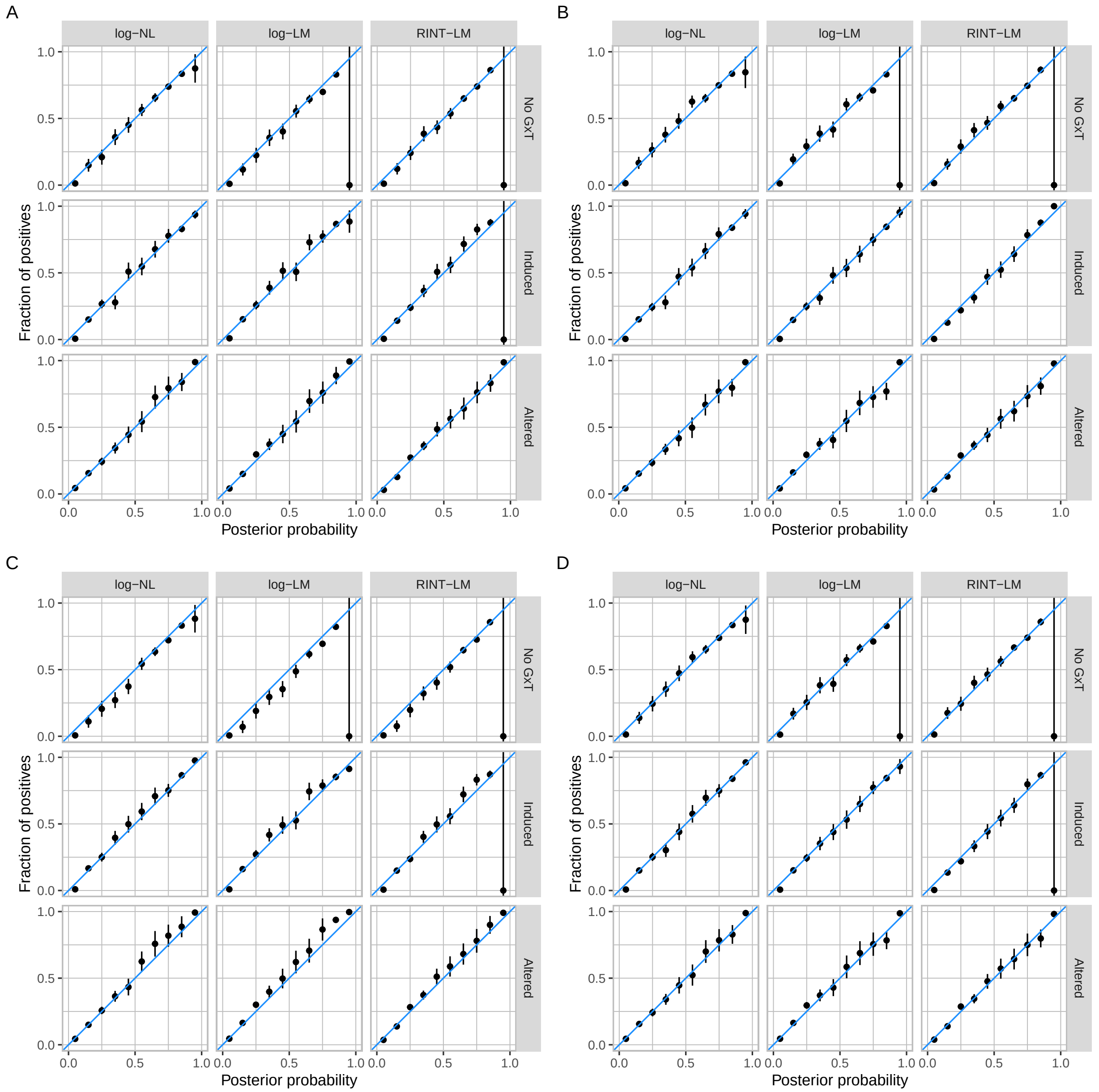


Figure S3. **Calibration of BMS with** log-**NL,** log-**LM, and RINT-LM for the "no G×T", "induced", and "altered" categories using MAP and Laplace approximation.** The same as in S2 but from 8000 simulations using MAP estimation and Laplace approximation.

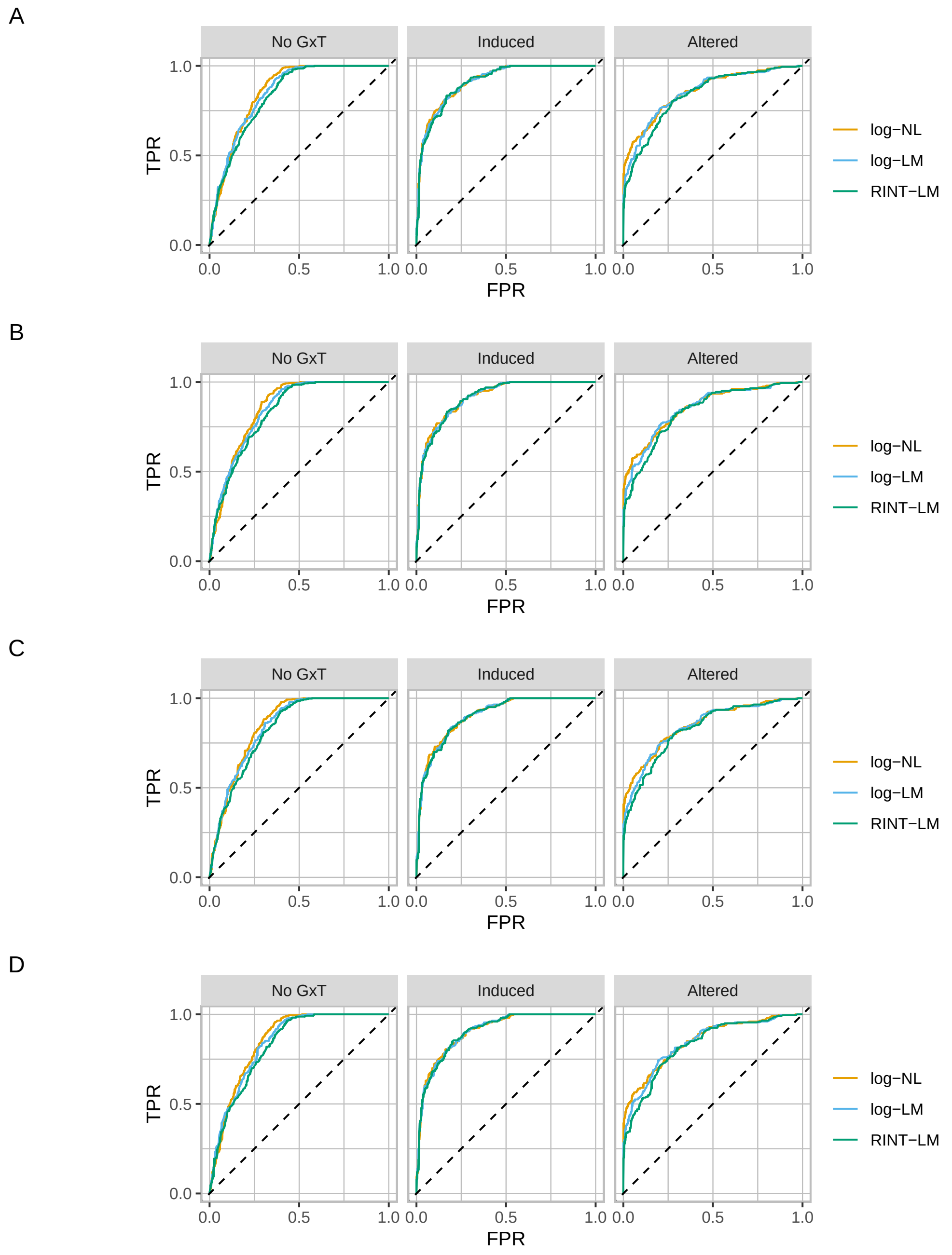


Figure S4. **ROC curves assessing the performance of BMS with** log**-NL,** log**-LM, and RINT-LM for the "no G×T", "induced", and "altered" categories using MCMC and bridge sampling.** The panels **A** to **D** show the results in scenarios 1 to 4, which are described in the legend to S2.

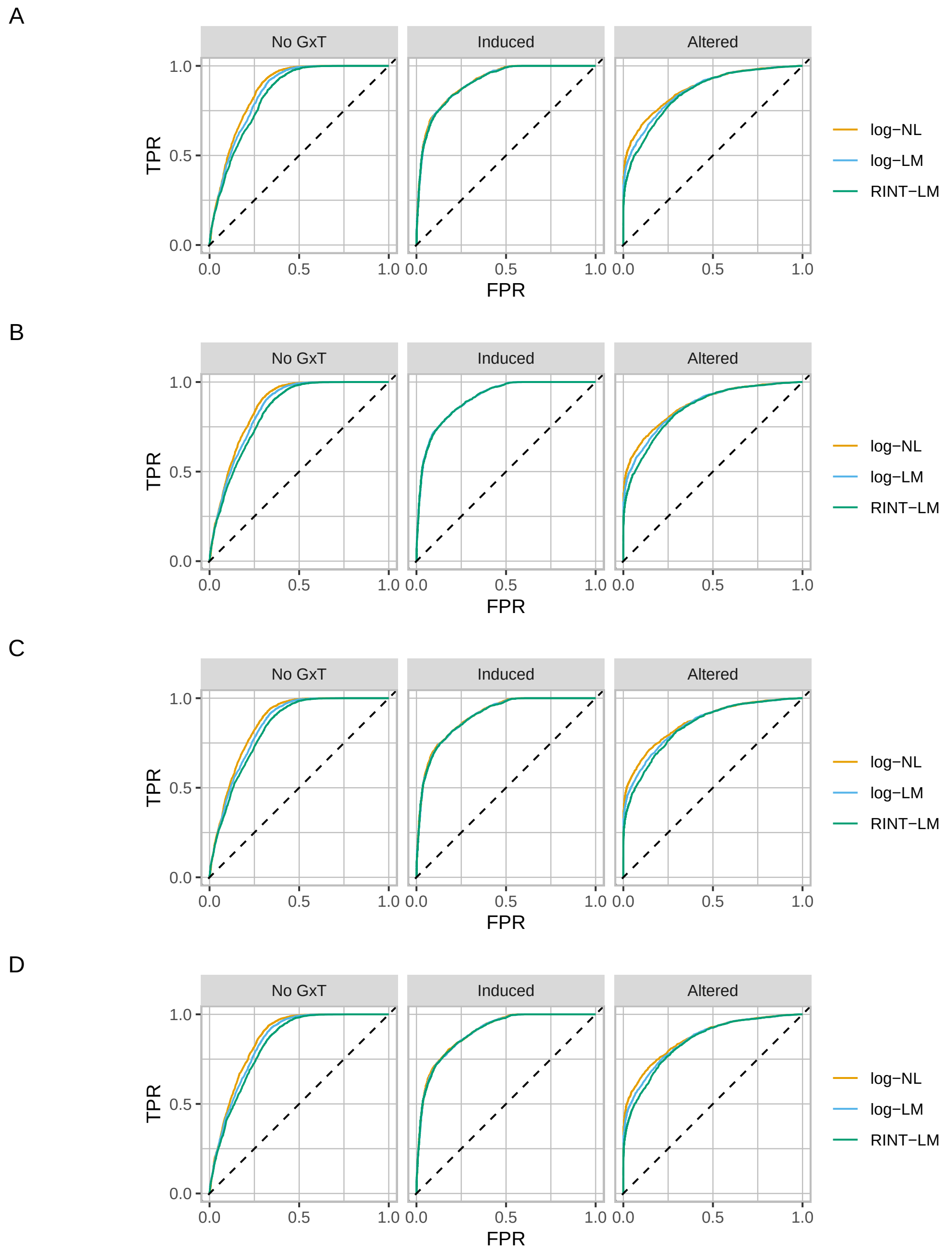


Figure S5. **ROC curves assessing the performance of BMS with** log**-NL,** log**-LM, and RINT-LM for the “no G×T”, “induced”, and “altered” categories using MAP and Laplace approximation.** The panels **A** to **D** show the results in scenarios 1 to 4, which are described in the legend to S2.

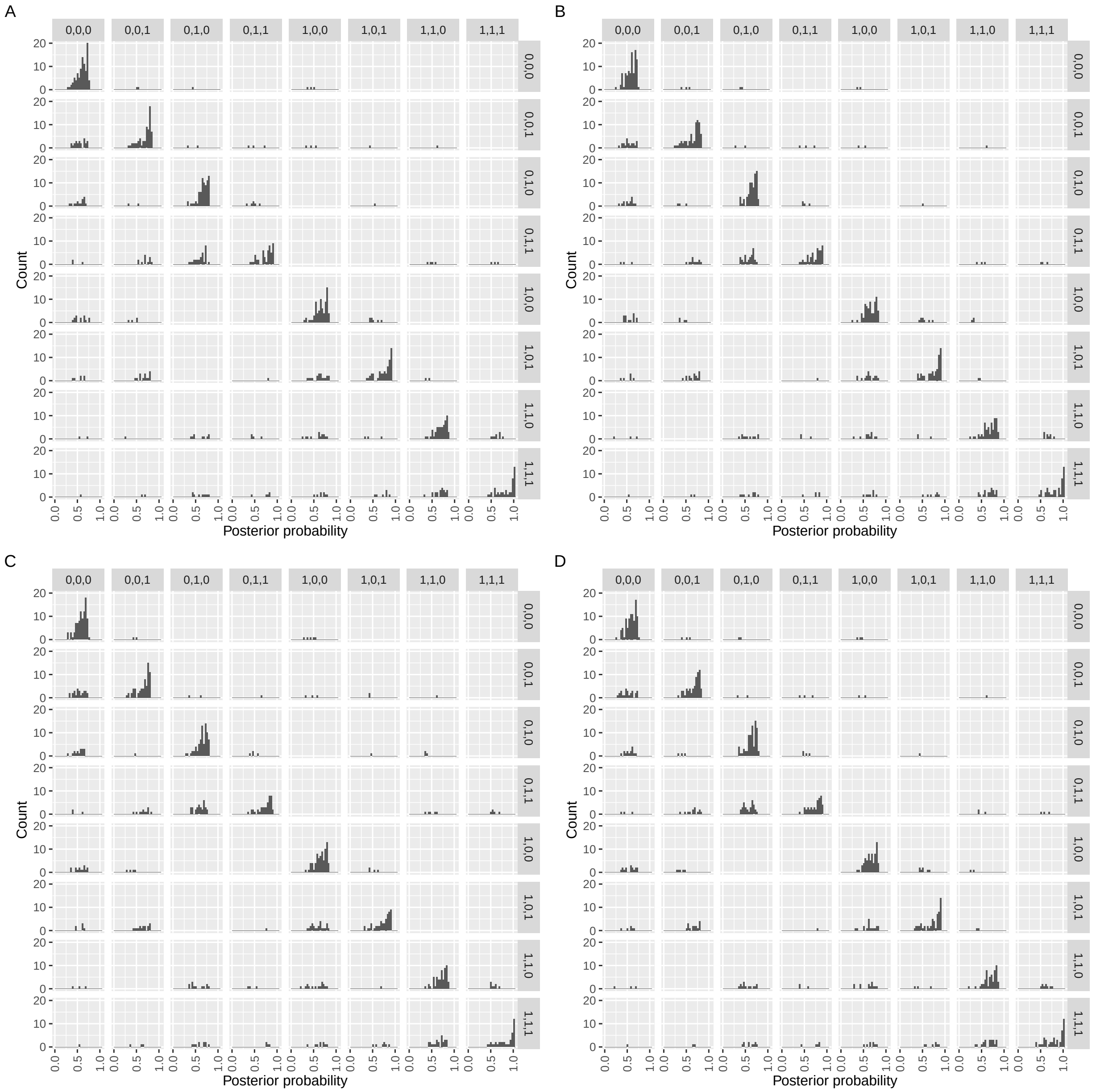


Figure S6. **Stratified histograms of posterior probability of the eight models obtained by BMS with** log-**NL using MCMC and bridge sampling.** In each panel, the rows and columns represent the data-generating and posterior mode models, respectively. The panels **A** to **D** show the results in scenarios 1 to 4, which are described in the legend to S2.

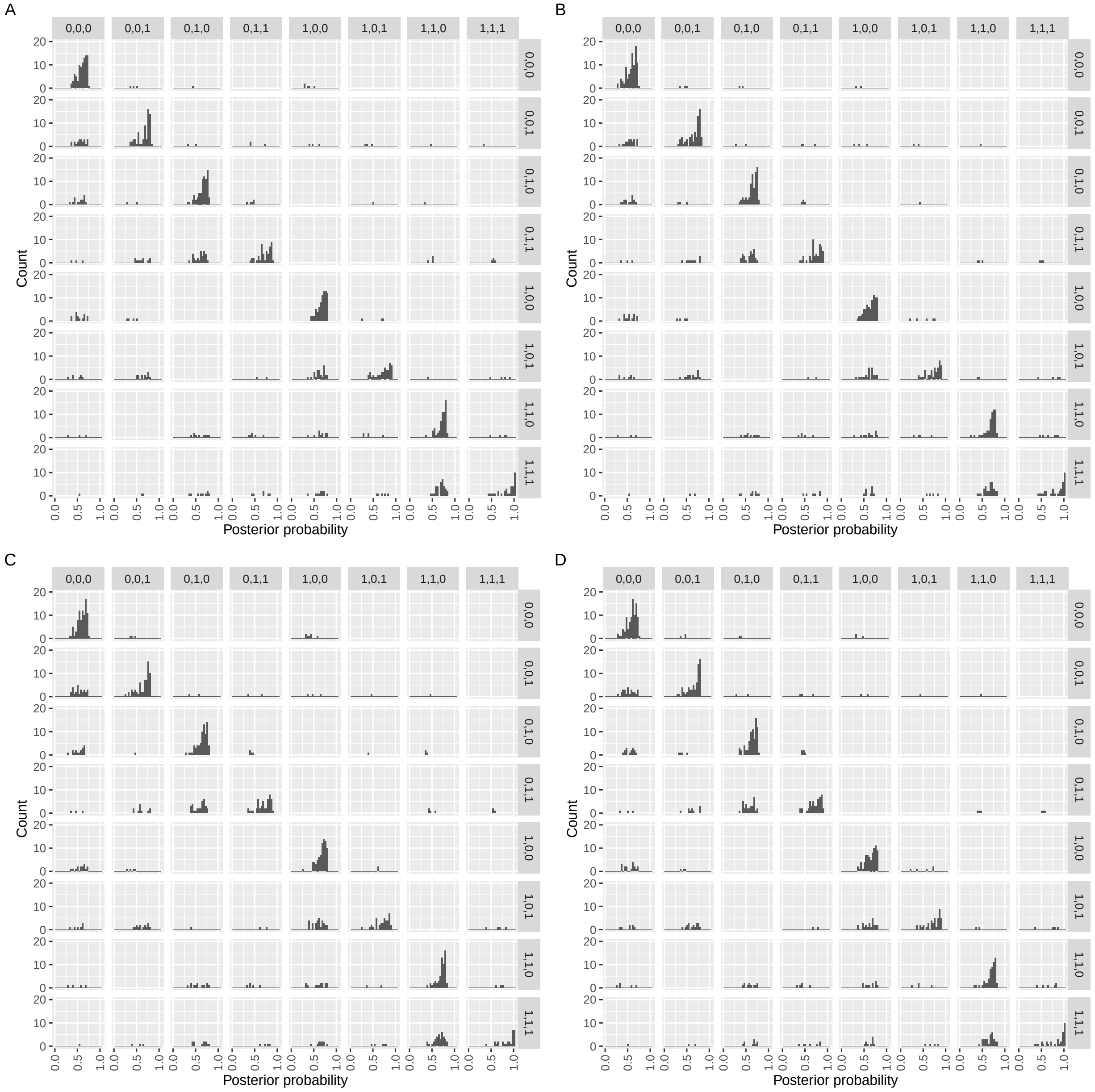


Figure S7. **Stratified histograms of posterior probability of the eight models obtained by BMS with** log**-LM using MCMC and bridge sampling.** The same as in S6 but for log-LM.

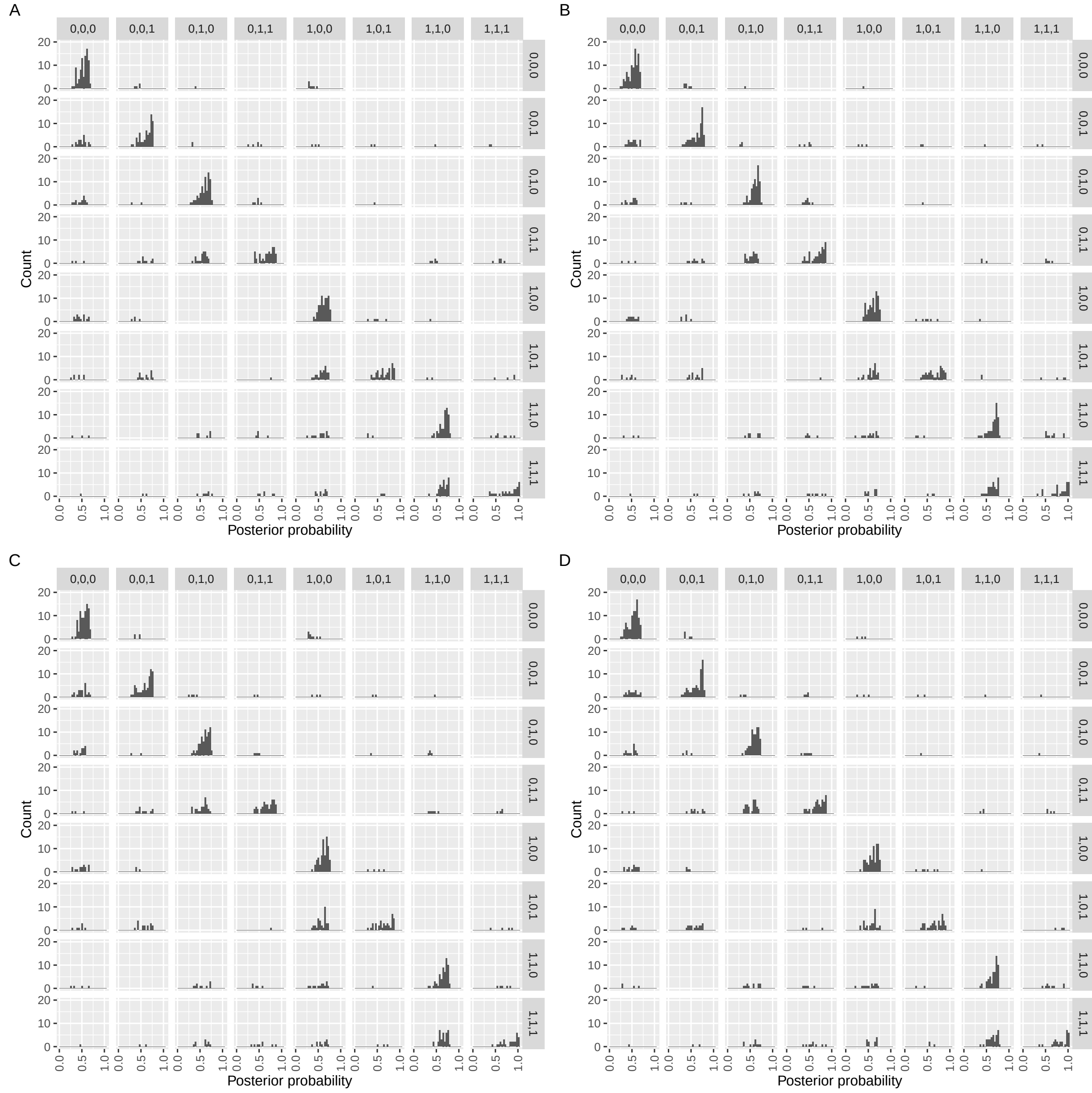


Figure S8. **Stratified histograms of posterior probability of the eight models obtained by BMS with RINT-LM using MCMC and bridge sampling.** The same as in S6 but for RINT-LM.

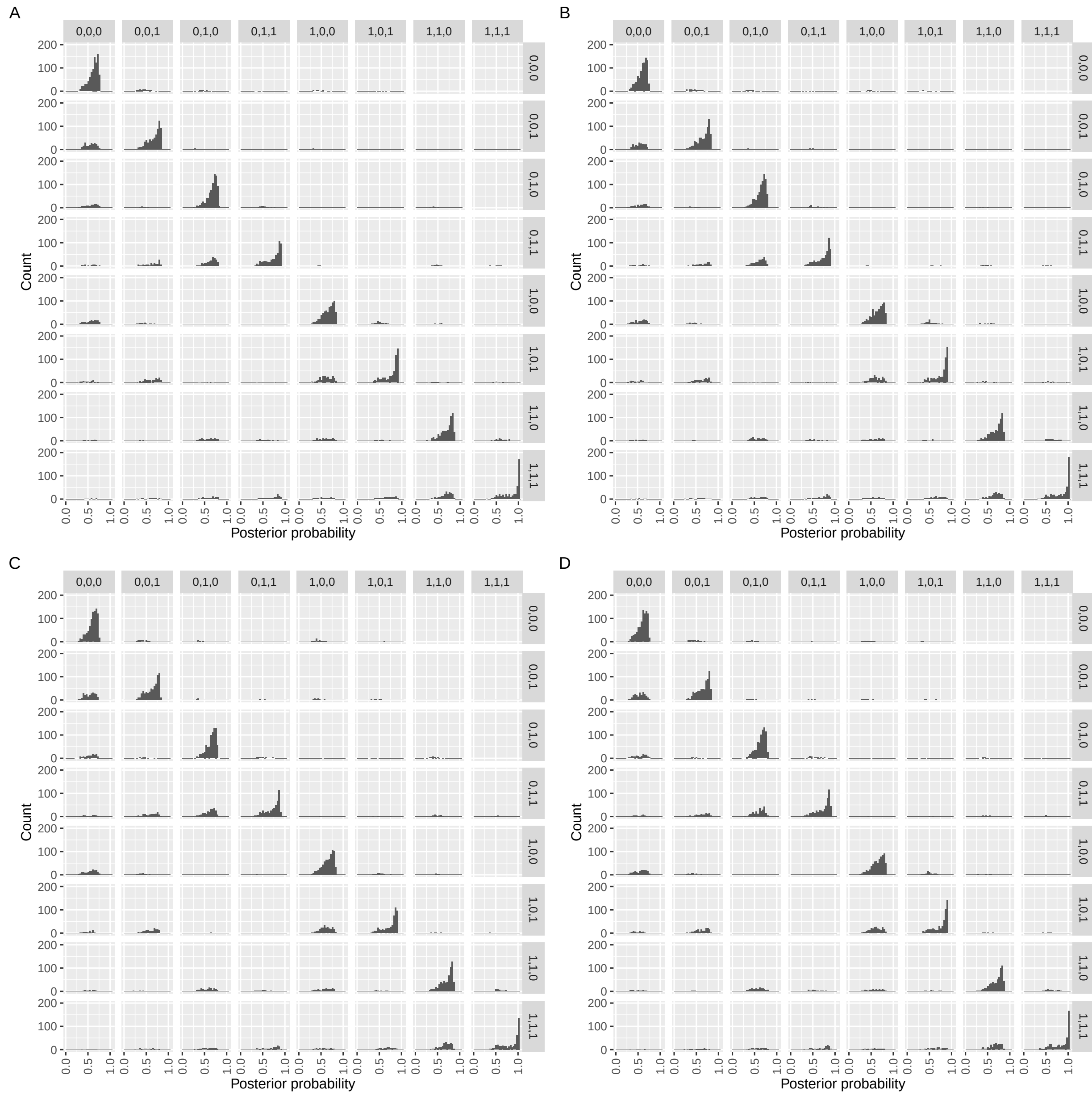


Figure S9. **Stratified histograms of posterior probability of the eight models obtained by BMS with** log-**NL using MAP estimation and Laplace approximation.** The same as in S6 but for MAP estimation and Laplace approximation.

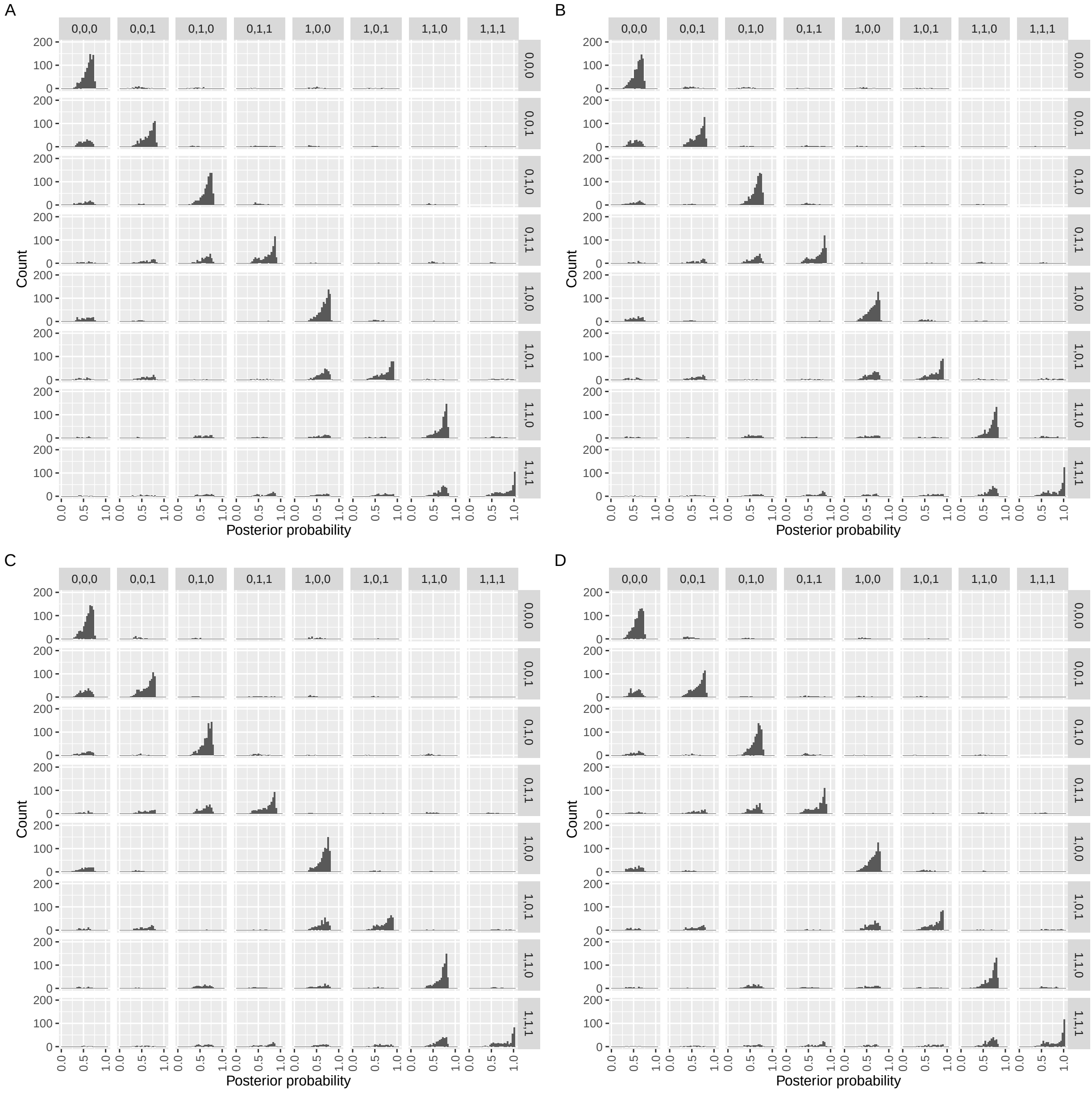


Figure S10. **Stratified histograms of posterior probability of the eight models obtained by BMS with** log**-LM using MAP estimation and Laplace approximation.** The same as in S9 but for log-LM.

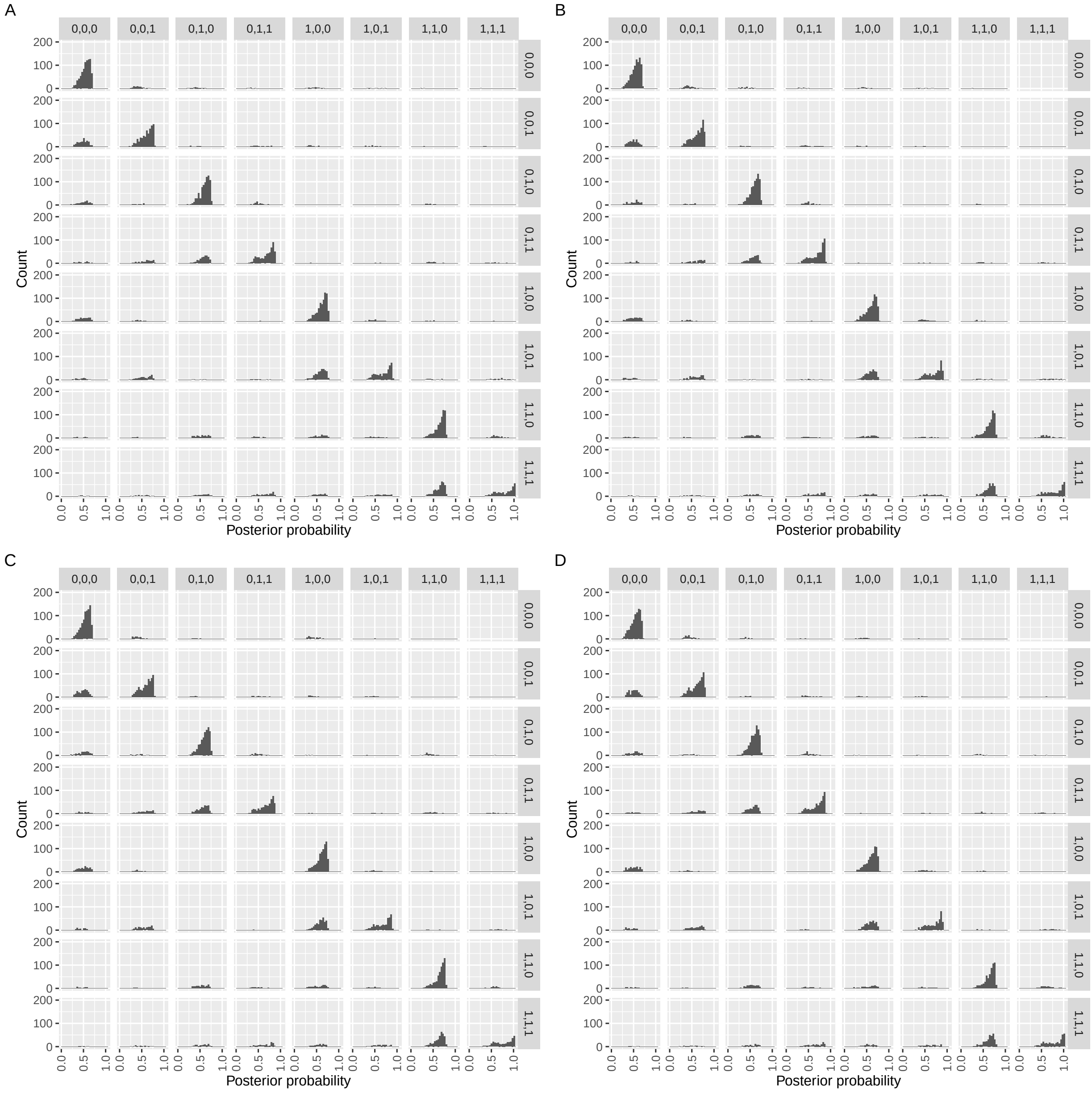


Figure S11. **Stratified histograms of posterior probability of the eight models obtained by BMS wth RINT-LM using MAP estimation and Laplace approximation.** The same as in S9 but for RINT-LM.

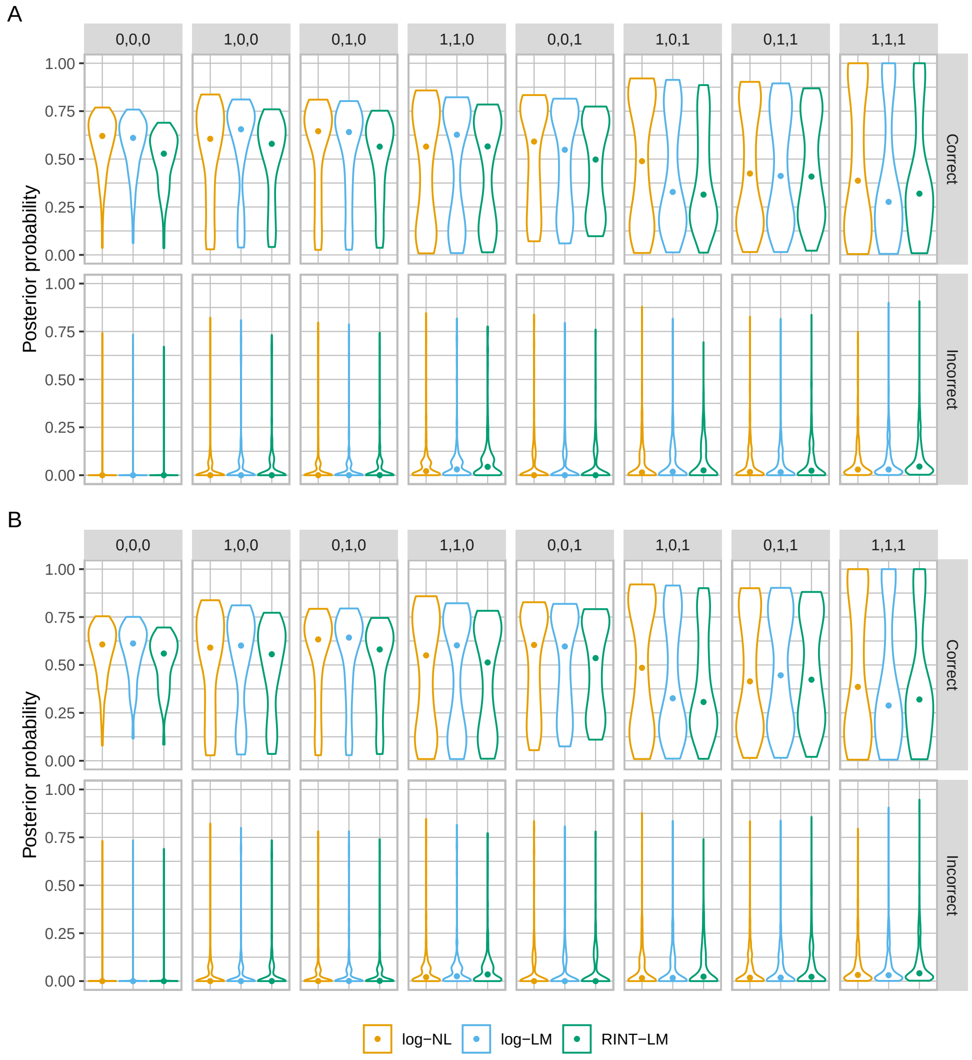


Figure S12. **Posterior probability of the correct and incorrect models for the eight categories obtained by BMS using MCMC and bridge sampling on data generated without random effect.** Violin plots for comparing the performance of BMS with log-NL, log-LM, and RINT-LM based on the distribution of posterior probability of the correct and incorrect models for each of the eight model categories. The closed circles represent median values. The panels **A** and **B** respectively show the results in scenarios 1 and 2, which are described in the legend to S2.

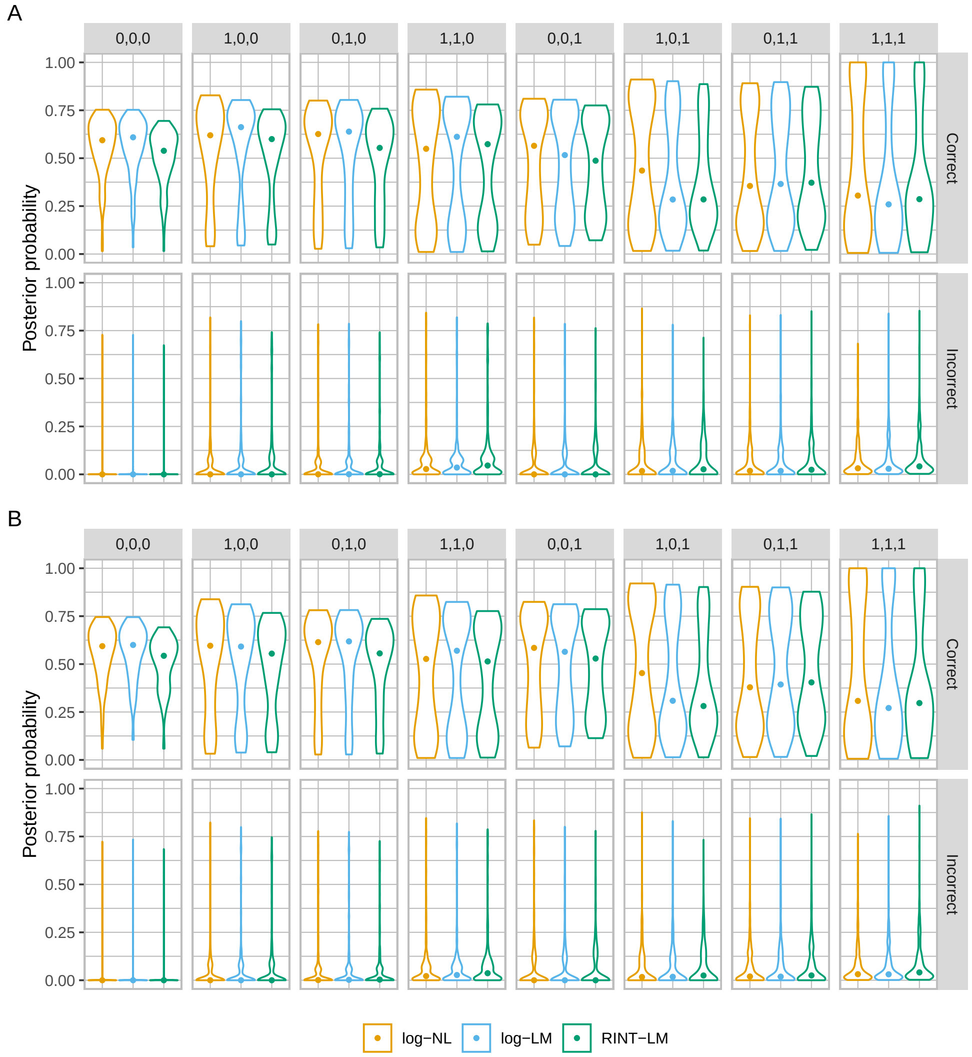


Figure S13. **Posterior probability of the correct and incorrect models for the eight categories obtained by BMS using MCMC and bridge sampling on data generated with donor random effect.**. The same as S12 but on data generated with donor random effect. The panels **A** and **B** respectively show the results in scenarios 3 and 4, which are described in the legend to S2.

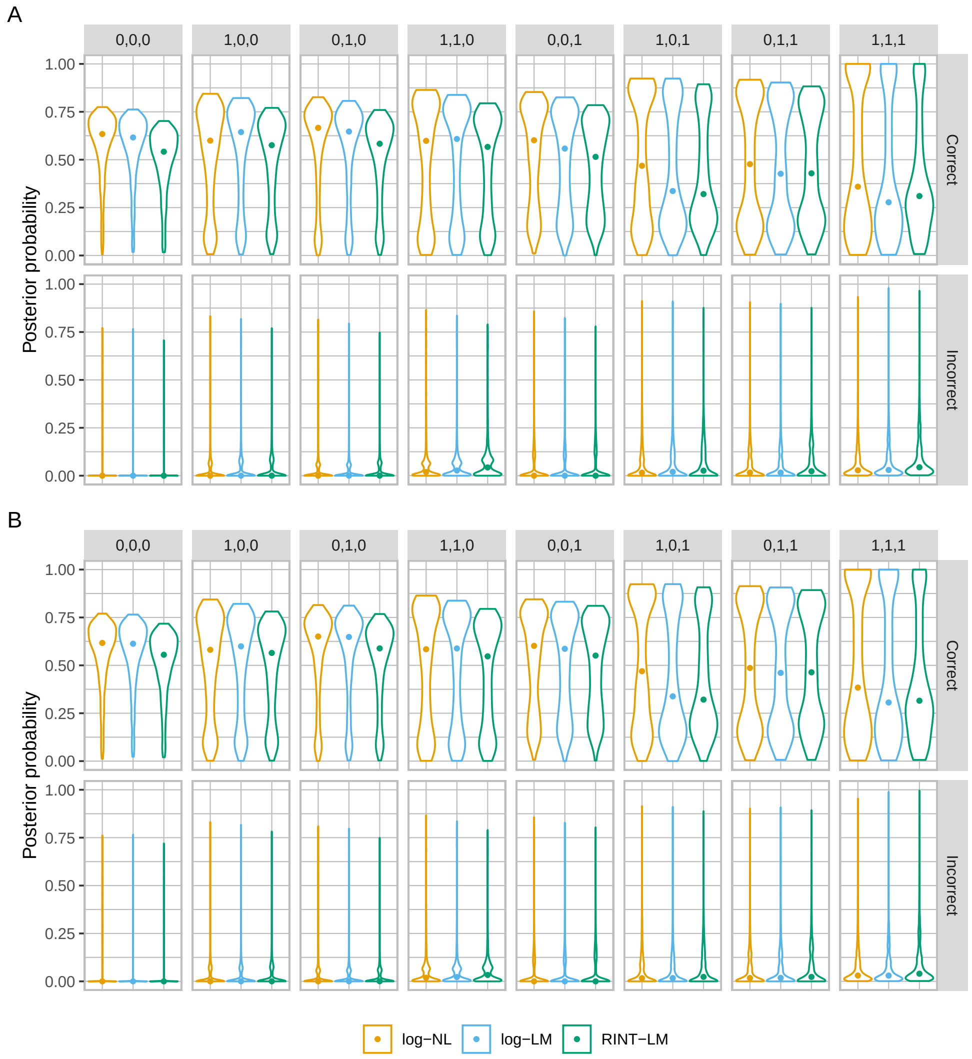


Figure S14. **Posterior probability of the correct and incorrect models for the eight categories obtained by BMS using MAP estimation and Laplace approximation on data generated without random effect.** The same as in S12 but for MAP estimation and Laplace approximation.

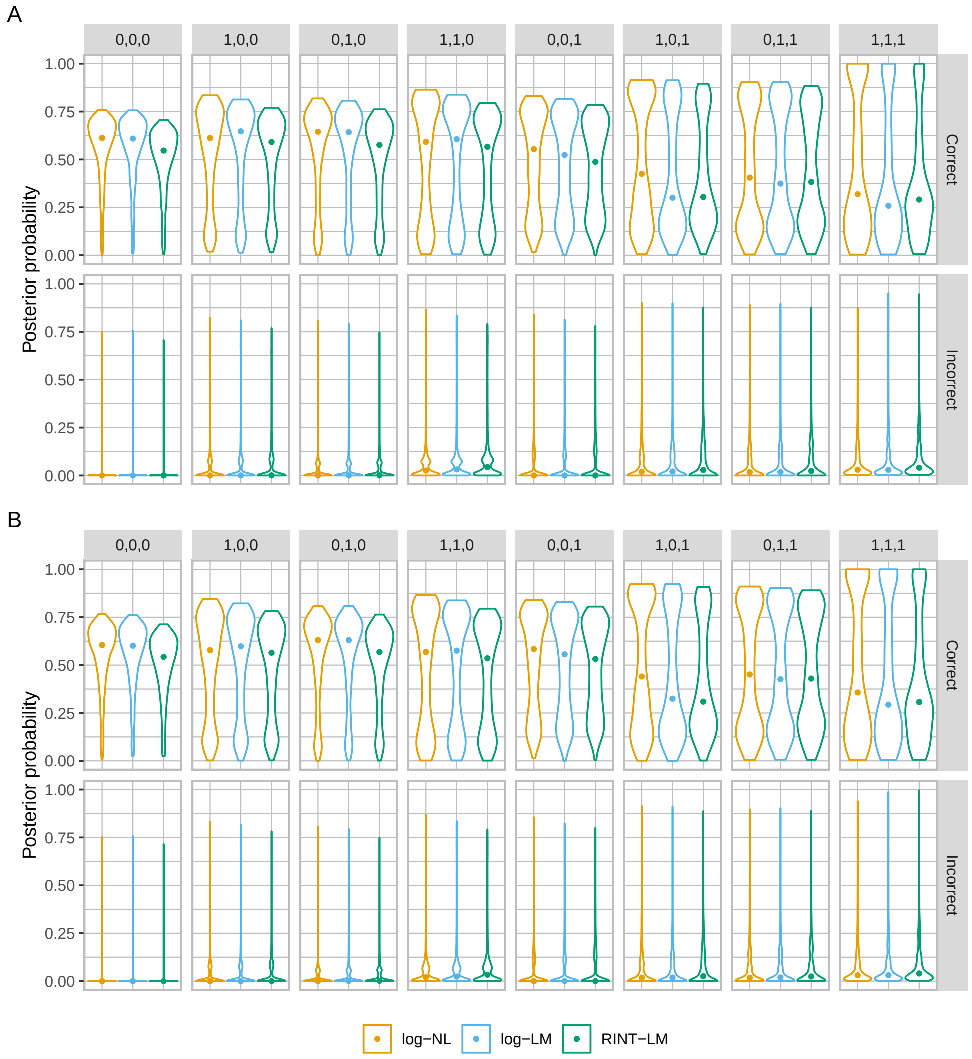


Figure S15. **Posterior probability of the correct and incorrect models for the eight categories obtained by BMS using MAP estimation and Laplace approximation on data generated with donor random effect.** The same as in S13 but for MAP estimation and Laplace approximation.

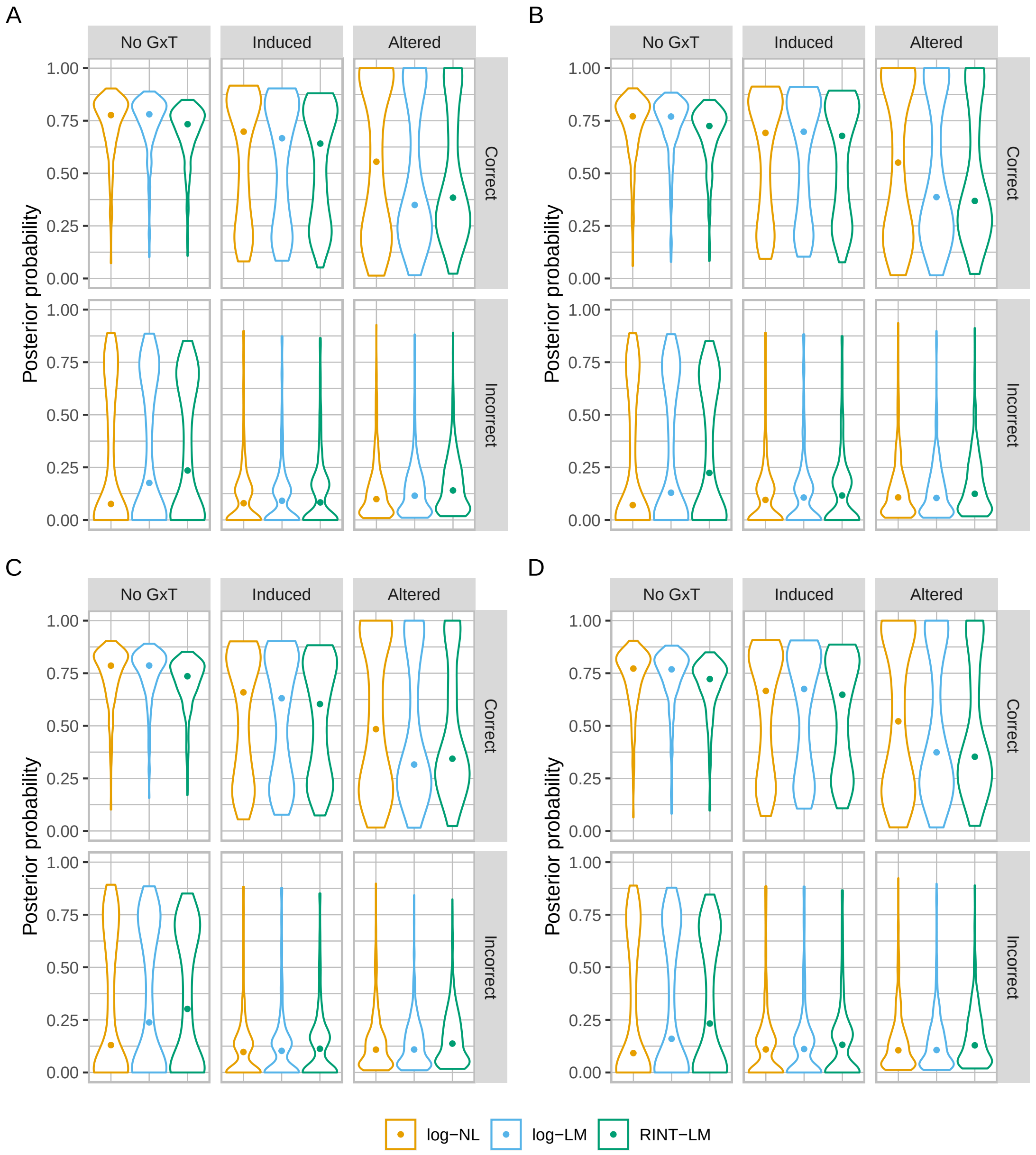


Figure S16. **Posterior probability of the correct and incorrect models for aggregated categories obtained by BMS using MCMC and bridge sampling.** Violin plots for comparing the performance of BMS with log-NL, log-LM, and RINT-LM based on the distribution of posterior probability of the correct and incorrect models for the "no GxT", "induced", and "altered" model categories. The closed circles represent median values. The panels **A** to **D** show the results in scenarios 1 to 4, which are described in the legend to S2.

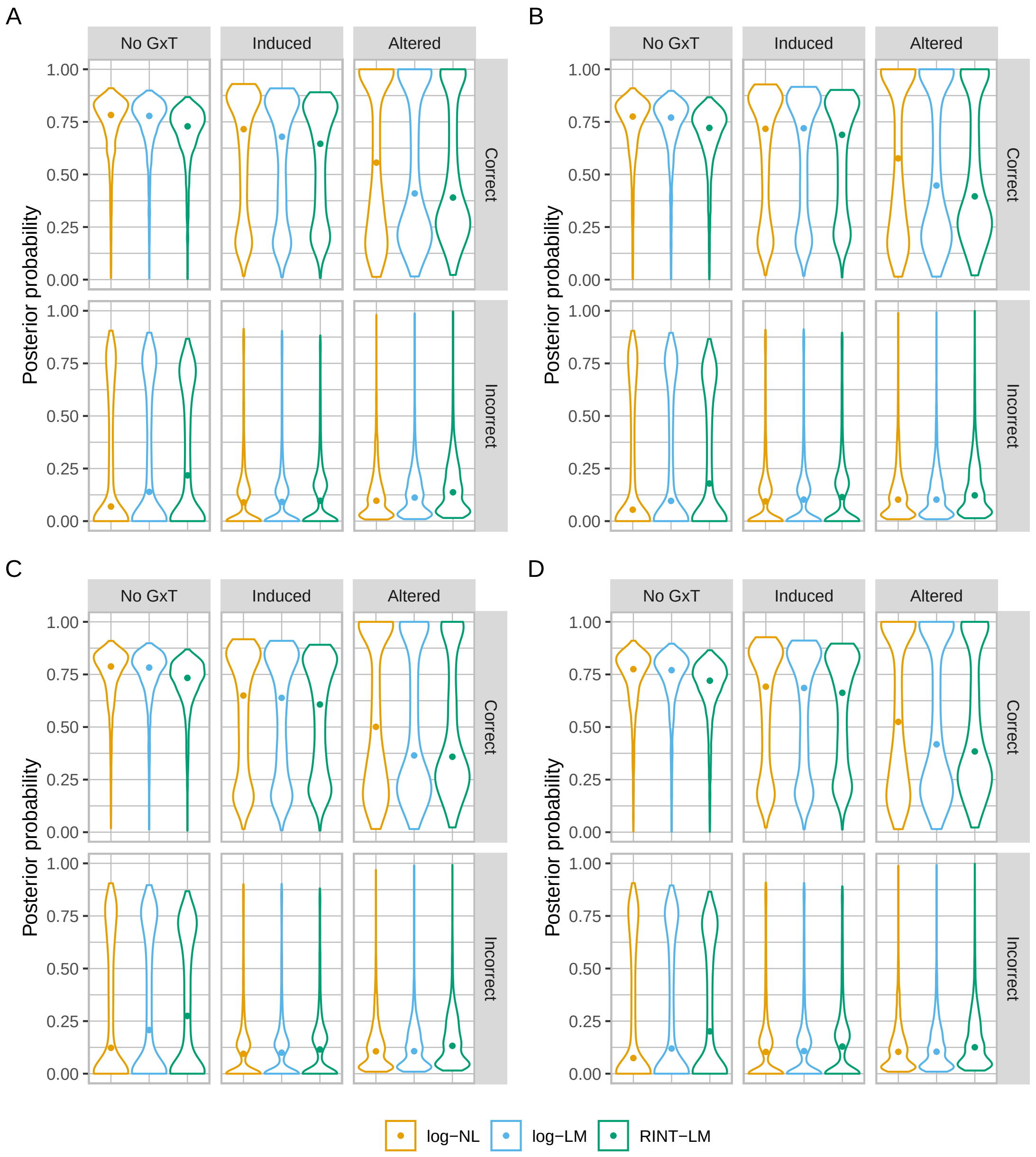


Figure S17. **Posterior probability of the correct and incorrect models for aggregated categories obtained by BMS using MAP estimation and Laplace approximation.** The same as in S16 but for MAP estimation and Laplace approximation.

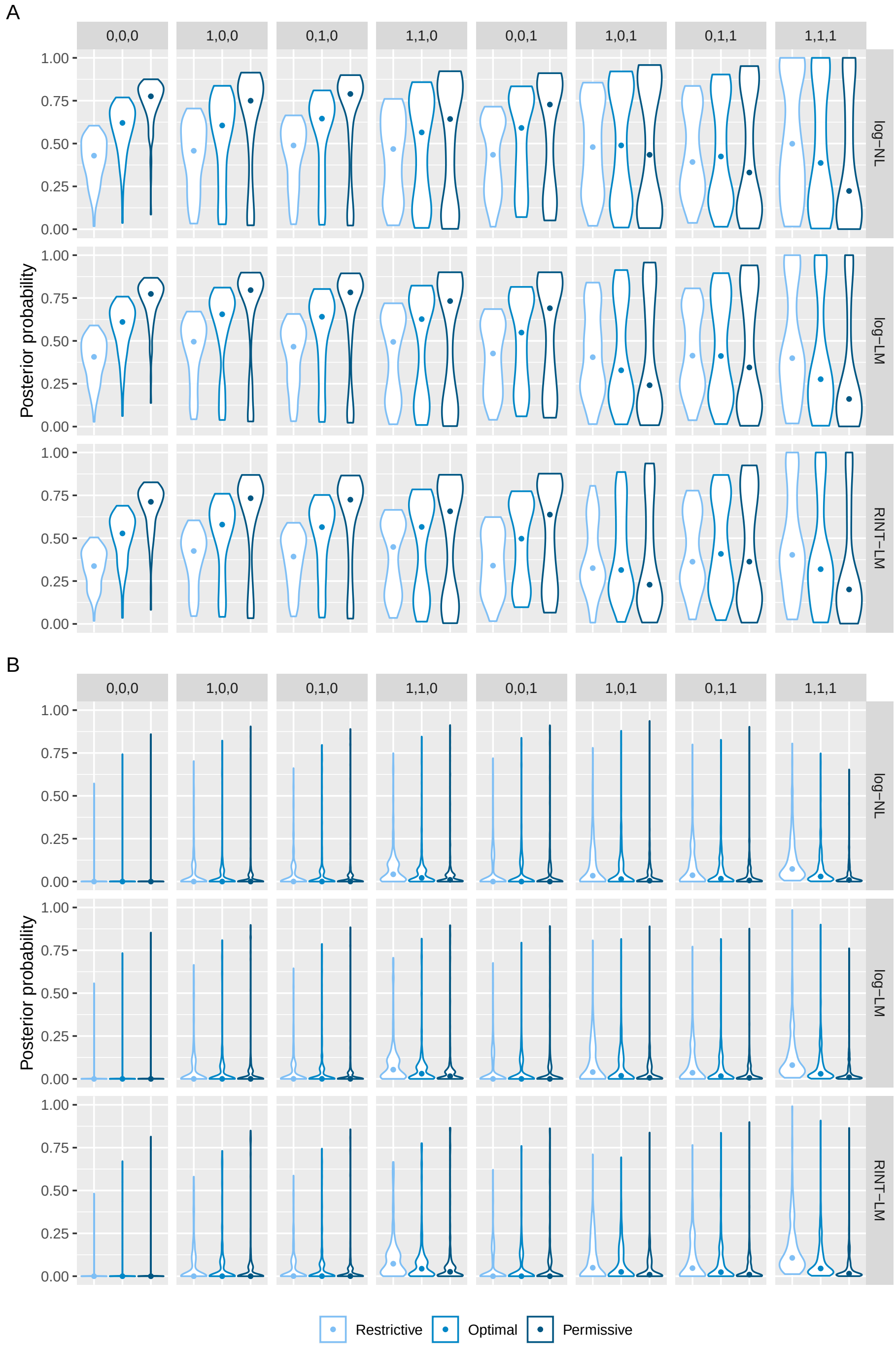


Figure S18. **Assessing the impact of the effect prior on the posterior probability of the correct and incorrect models from analyses without random effect using MCMC and bridge sampling.** Violin plots showing the distribution of posterior probability of the correct (**A**) and incorrect (**B**) models for each of the eight model categories with varying hyper parameter values (see S1 Text). The closed circles represent median values. Shown is the results in scenario 1, which is described in the legend to S2.

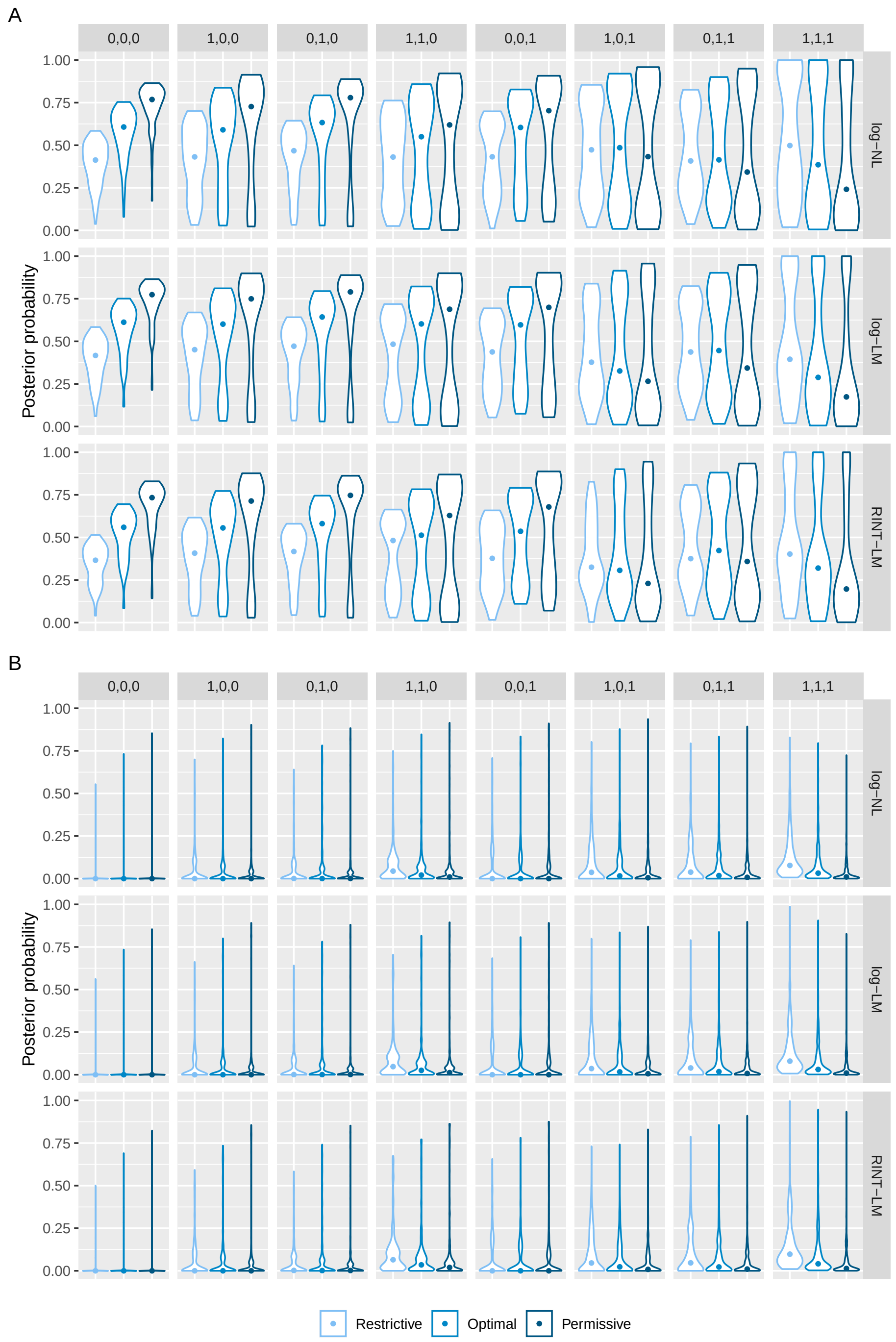


Figure S19. **Assessing the impact of the effect prior on the posterior probability of the correct and incorrect models from analyses with donor random effect in model fitting but not in data generation using MCMC and bridge sampling.** The same as in S18 but in scenario 2, which is described in the legend to S2.

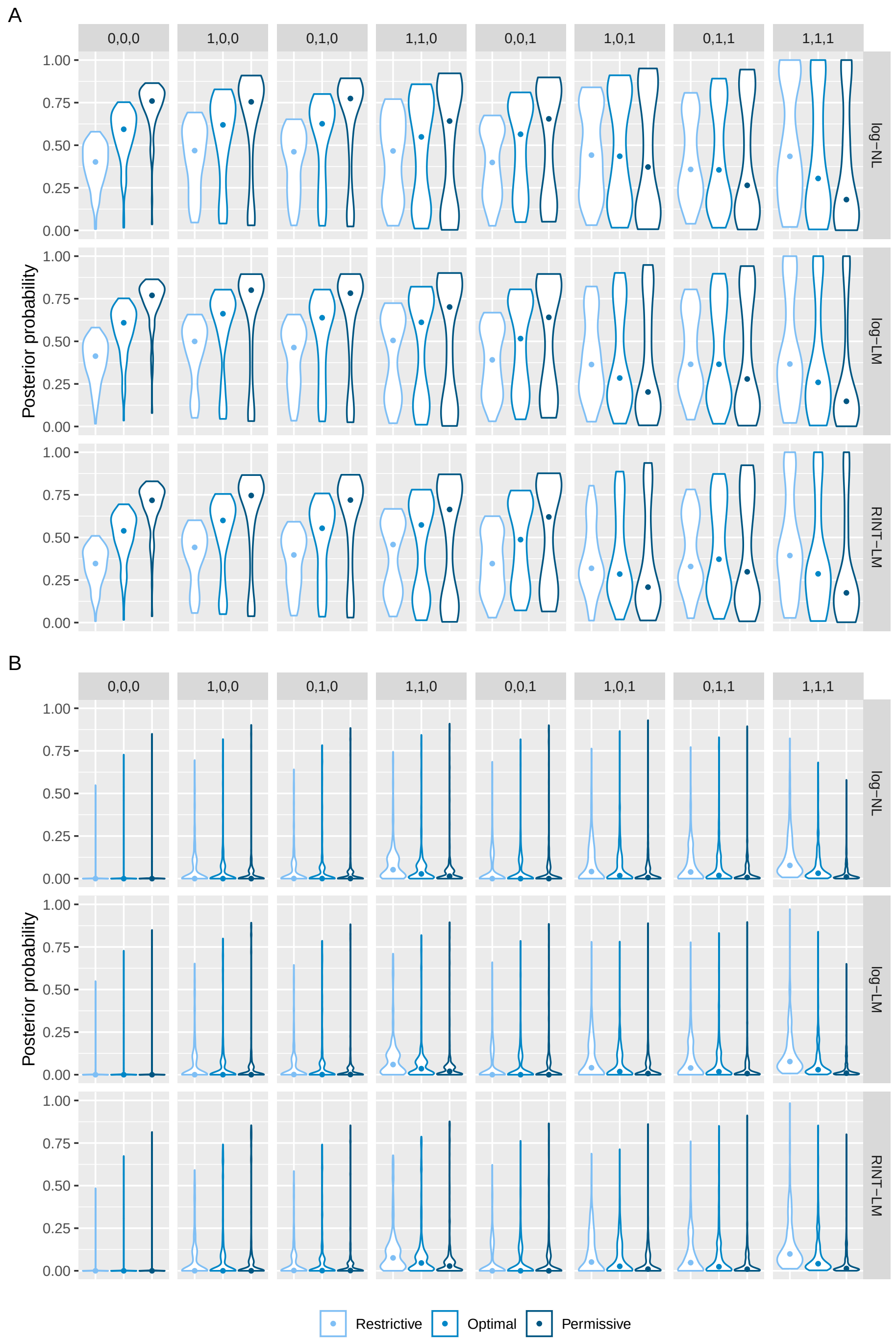


Figure S20. **Assessing the impact of the effect prior on the posterior probability of the correct and incorrect models from analyses with donor random effect in data generation but not in model fitting using MCMC and bridge sampling.** The same as in S18 but in scenario 3, which is described in the legend to S2.

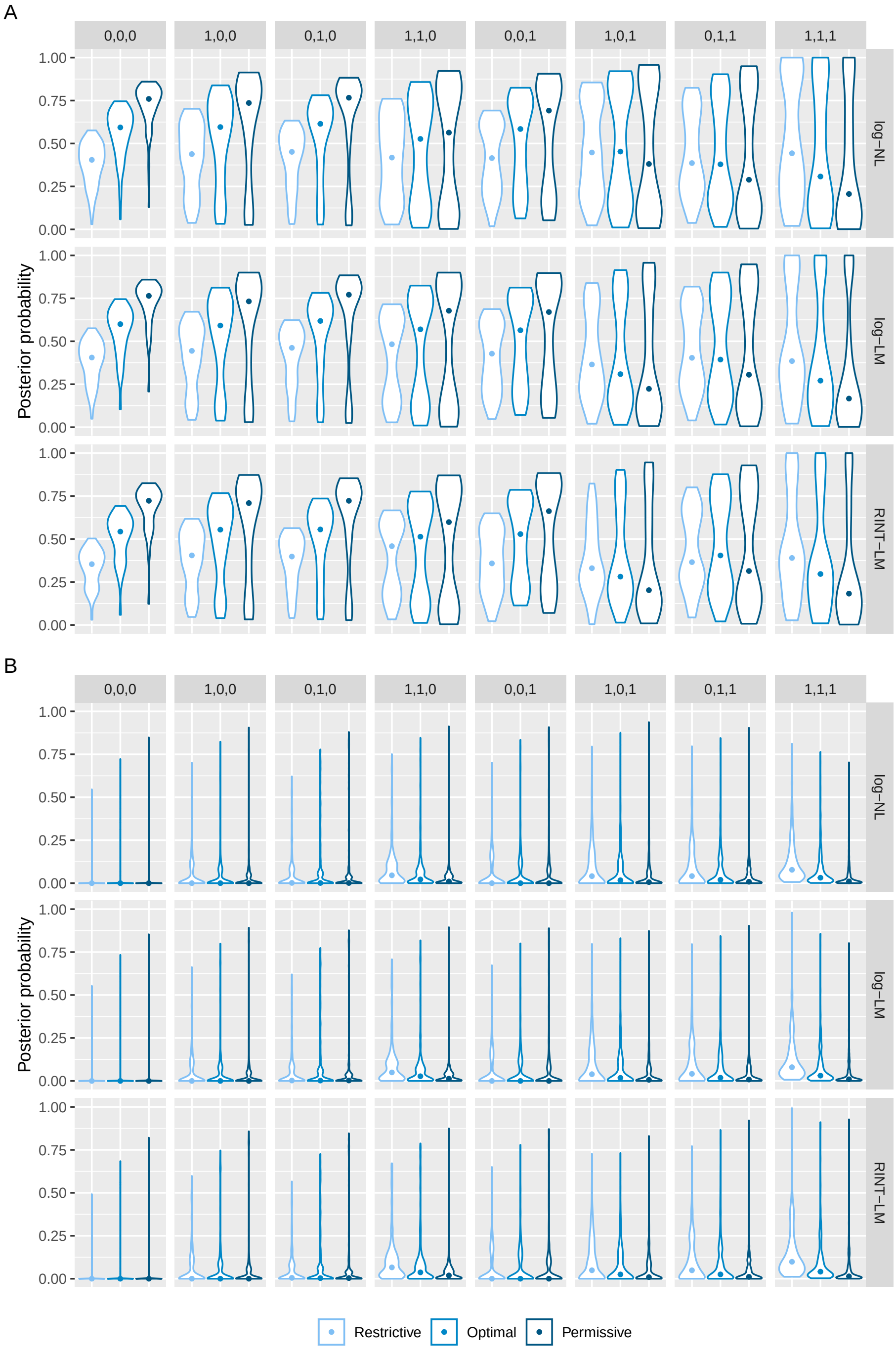


Figure S21. **Assessing the impact of the effect prior on the posterior probability of the correct and incorrect models from analyses with donor random effect in both data generation and model fitting using MCMC and bridge sampling.** The same as in S18 but in scenario 4, which is described in the legend to S2.

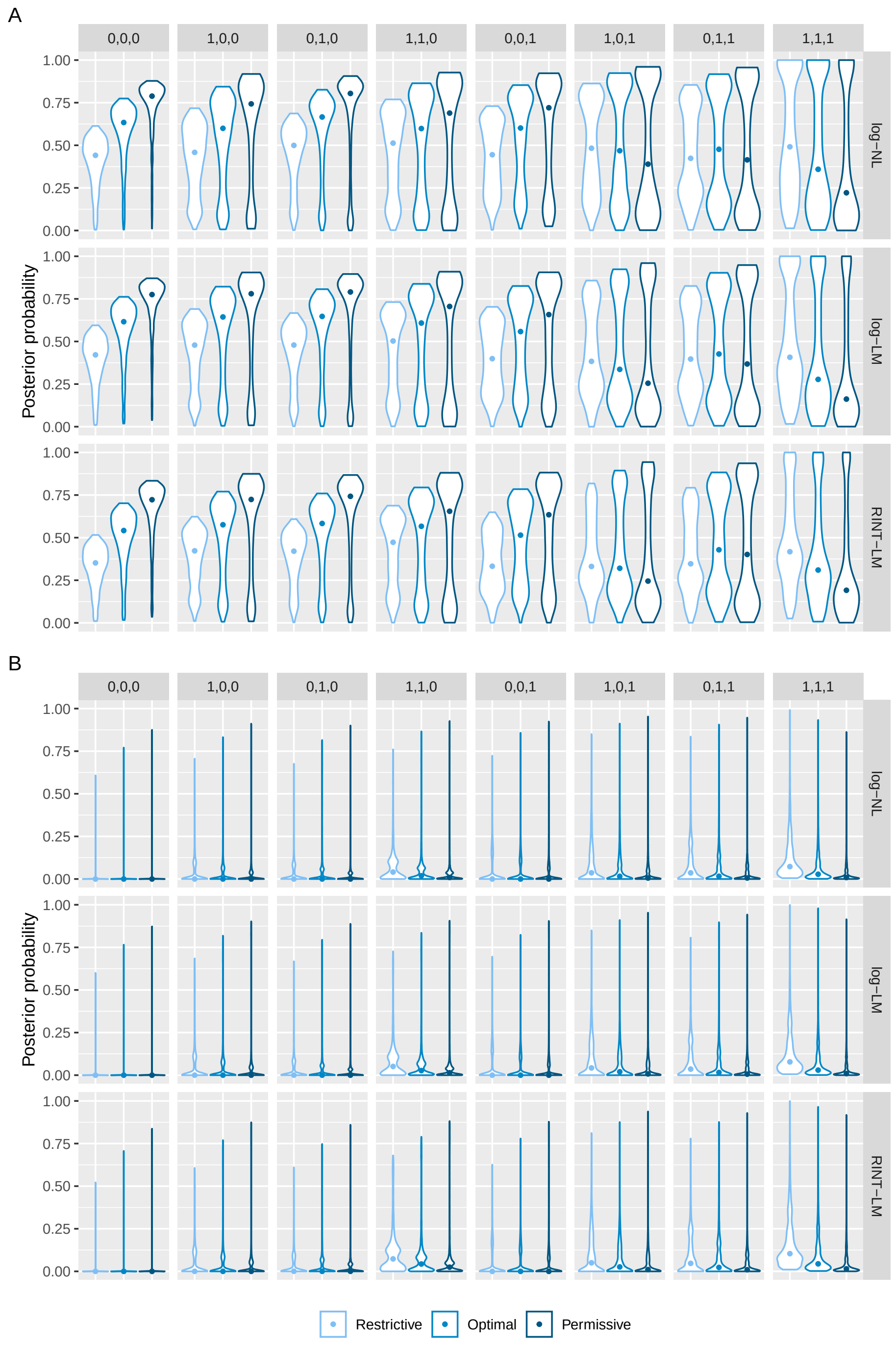


Figure S22. **Assessing the impact of the effect prior on the posterior probability of the correct and incorrect models from analyses without random effect using MAP estimation and Laplace approximation.** The same as in S18 but for MAP estimation and Laplace approximation.

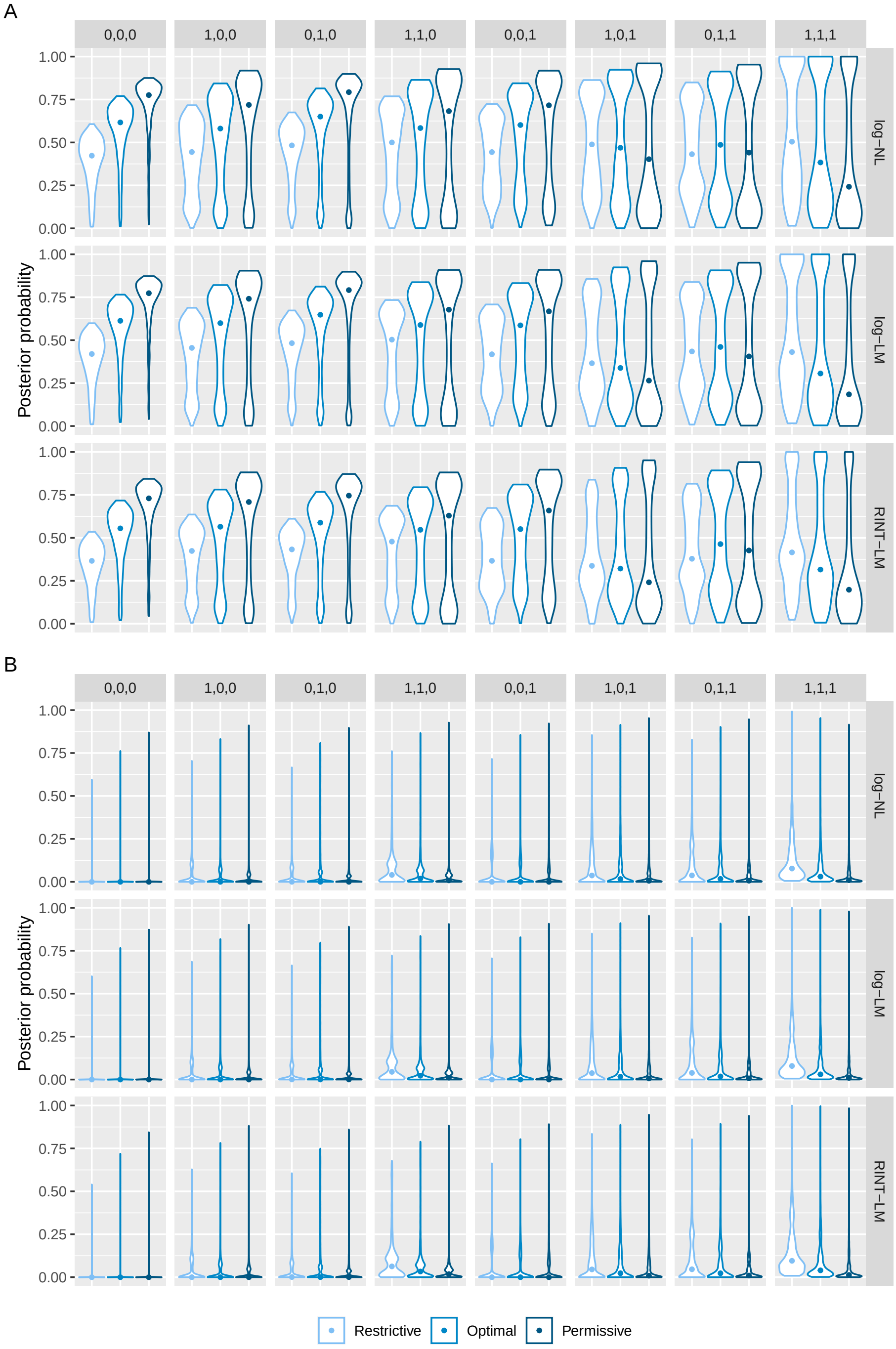


Figure S23. **Assessing the impact of the effect prior on the posterior probability of the correct and incorrect models from analyses with donor random effect in model fitting but not in data generation using MAP estimation and Laplace approximation.** The same as in S19 but for MAP estimation and Laplace approximation.

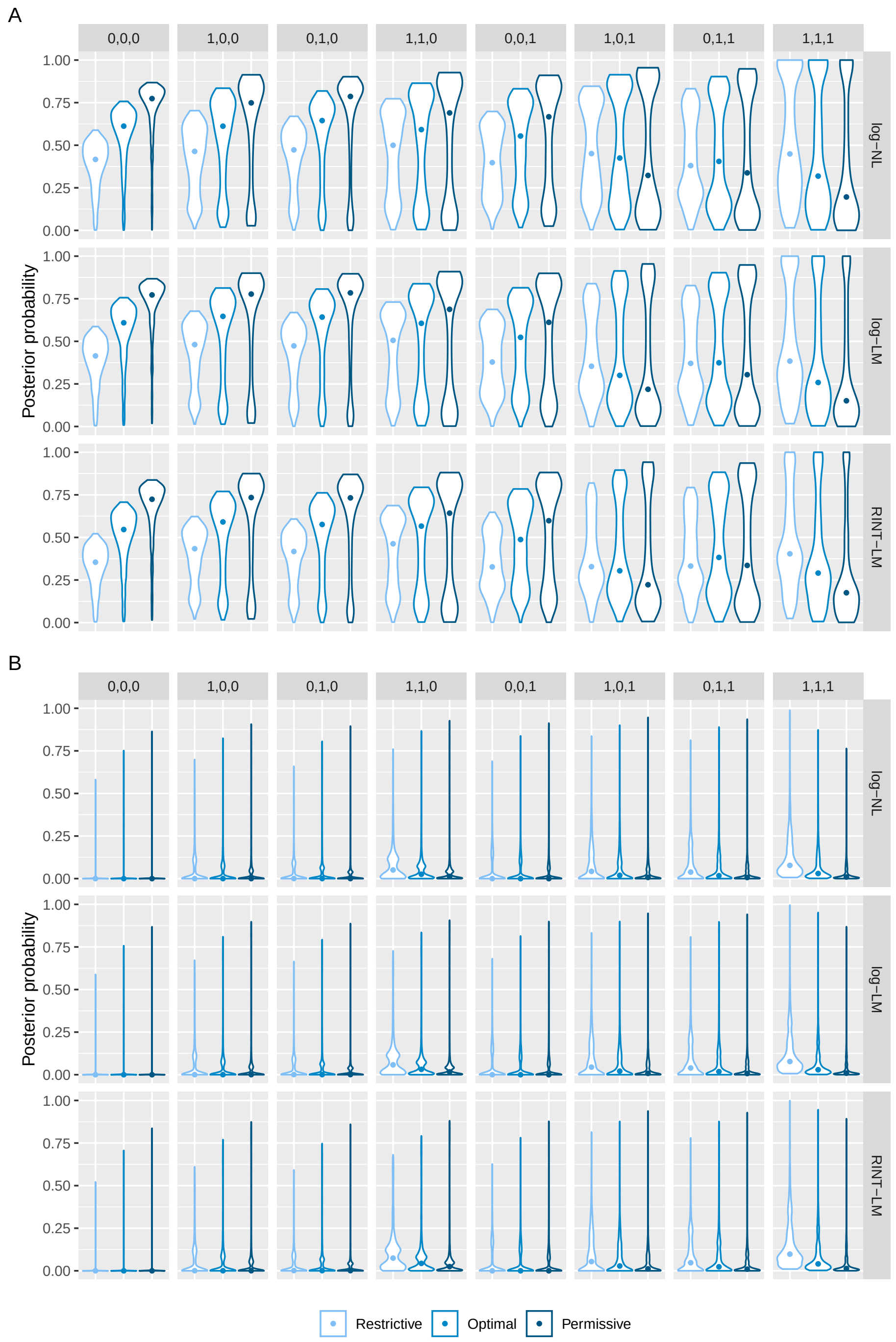


Figure S24. **Assessing the impact of the effect prior on the posterior probability of the correct and incorrect models from analyses with donor random effect in data generation but not in model fitting using MAP estimation and Laplace approximation.** The same as in S20 but for MAP estimation and Laplace approximation.

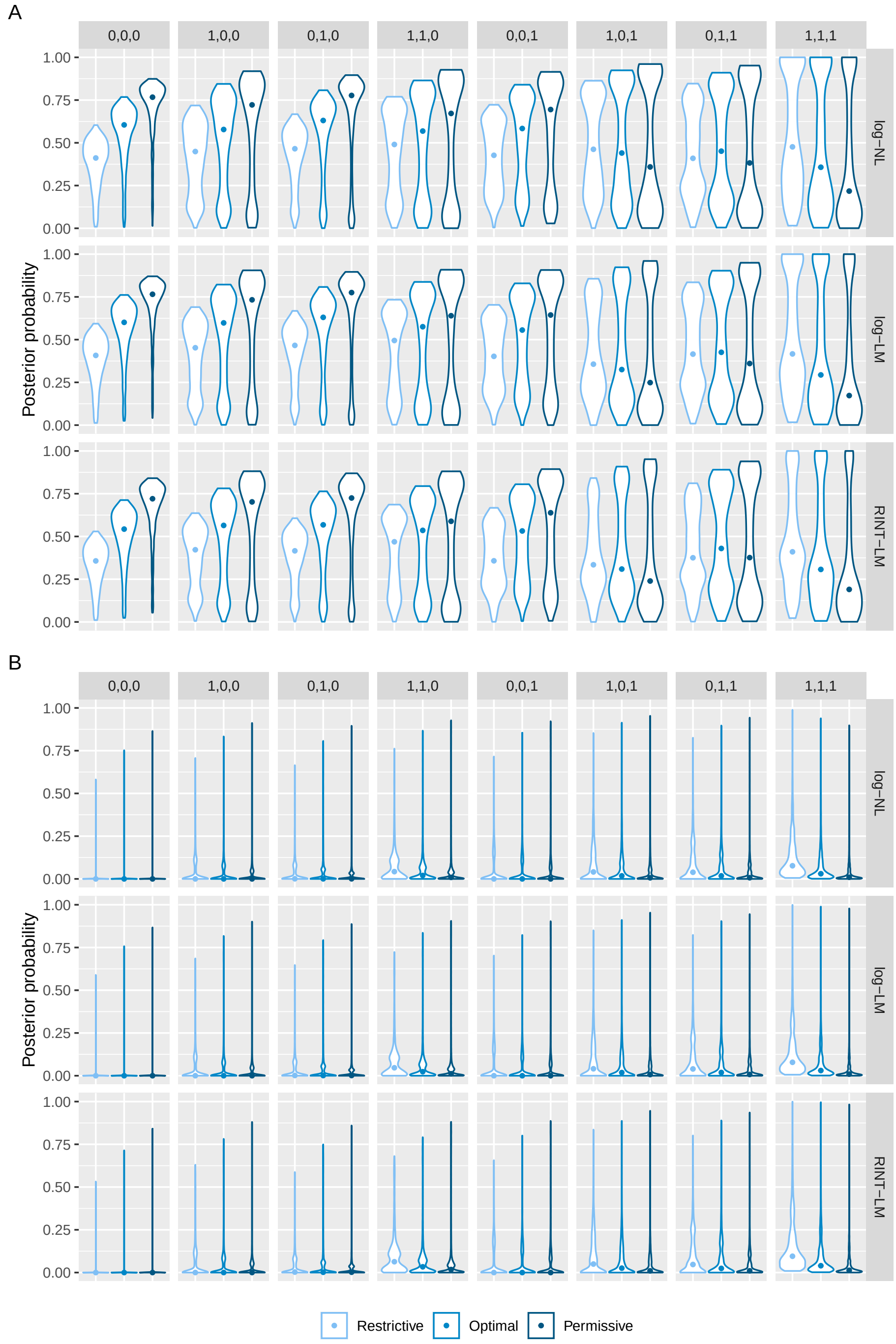


Figure S25. **Assessing the impact of the effect prior on the posterior probability of the correct and incorrect models from analyses with donor random effect in both data generation and model fitting using MAP estimation and Laplace approximation.** The same as in S21 but for MAP estimation and Laplace approximation.

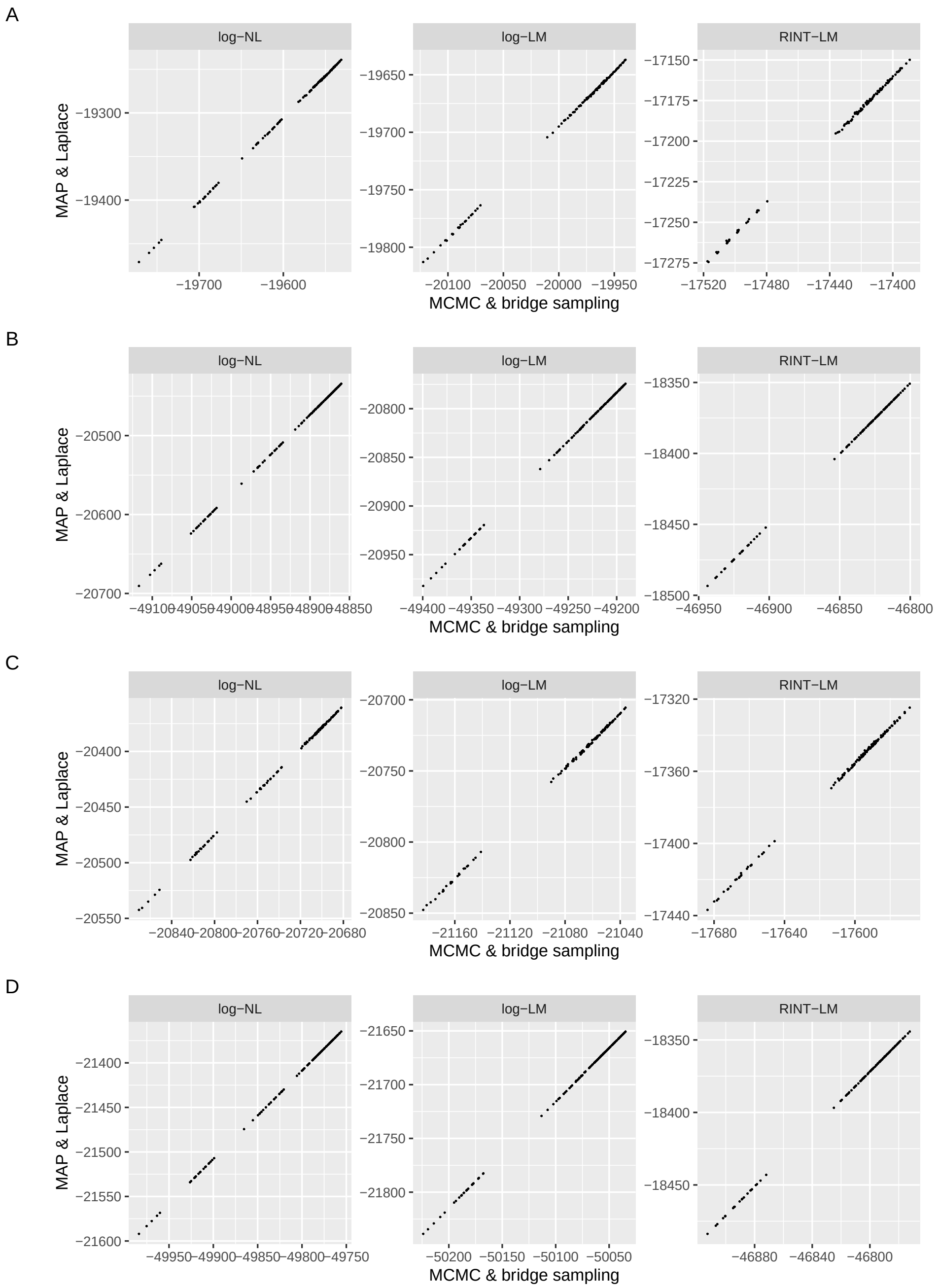


Figure S26. **Comparison of the sum of the** log **of marginal likelihood across combinations of hyperparameter values between two computational approaches for** log**-NL,** log**-LM, and RINT-LM with and without donor random effects.** Scatter plots comparing results obtained by MCMC followed by bridge sampling and those obtained by MAP estimation followed by Laplace approximation. Each point represents the log of the marginal likelihood summed over 80 feature-SNP pairs. The values are compared across 125 combinations of the hyperparameter values (see S1 text for details).

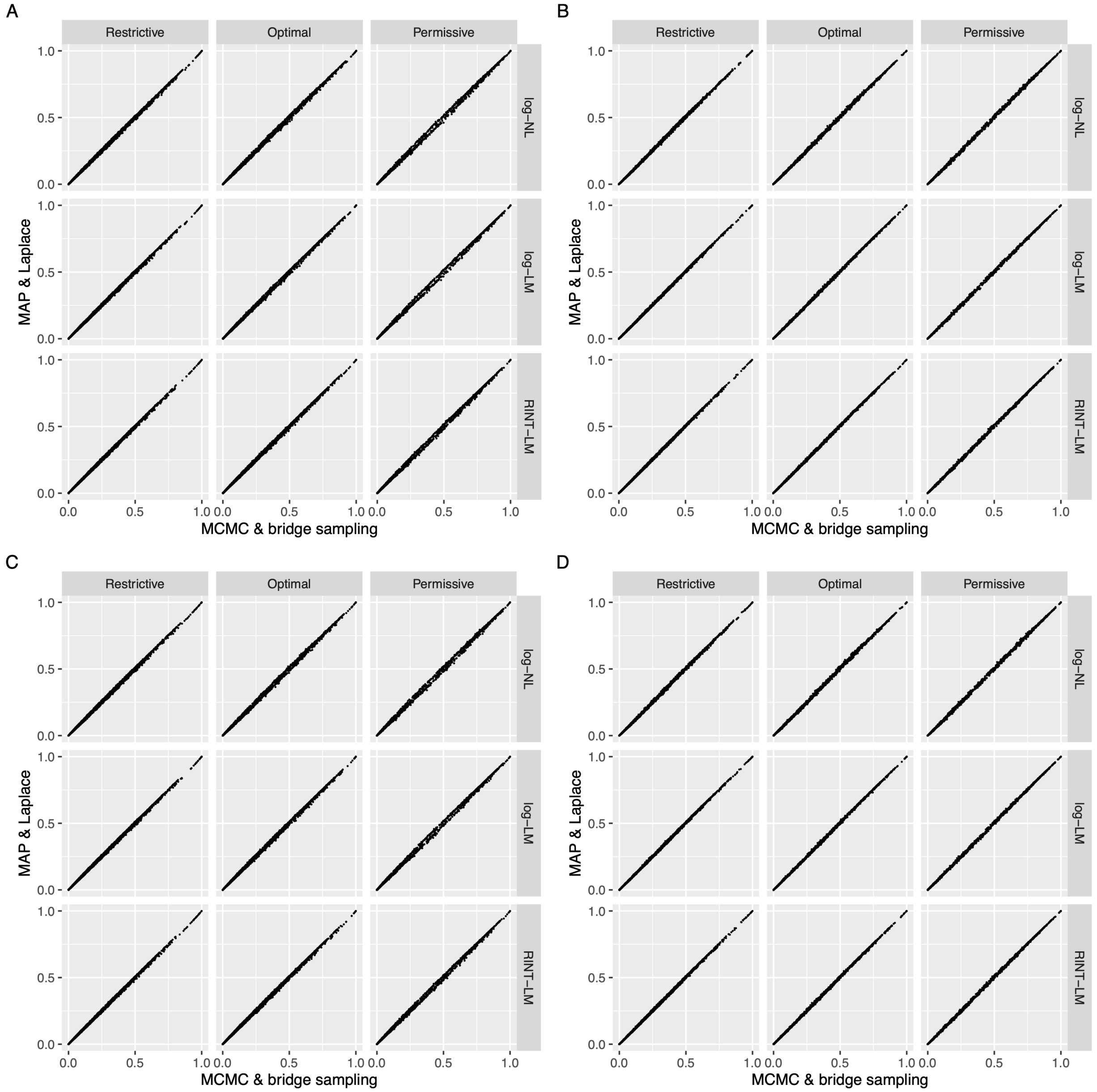


Figure S27. **Comparison of posterior probability at the optimal hyperparameter values between two computational approaches for** log**-NL,** log**-LM, and RINT-LM with and without donor random effect.** Scatter plots comparing results obtained by MCMC followed by bridge sampling and those obtained by MAP estimation followed by Laplace approximation Each point represents the posterior probability of a mode for a feature-SNP pair. The values are compared across eight models and 800 feature-SNP pairs (i.e., 6400 combinations).

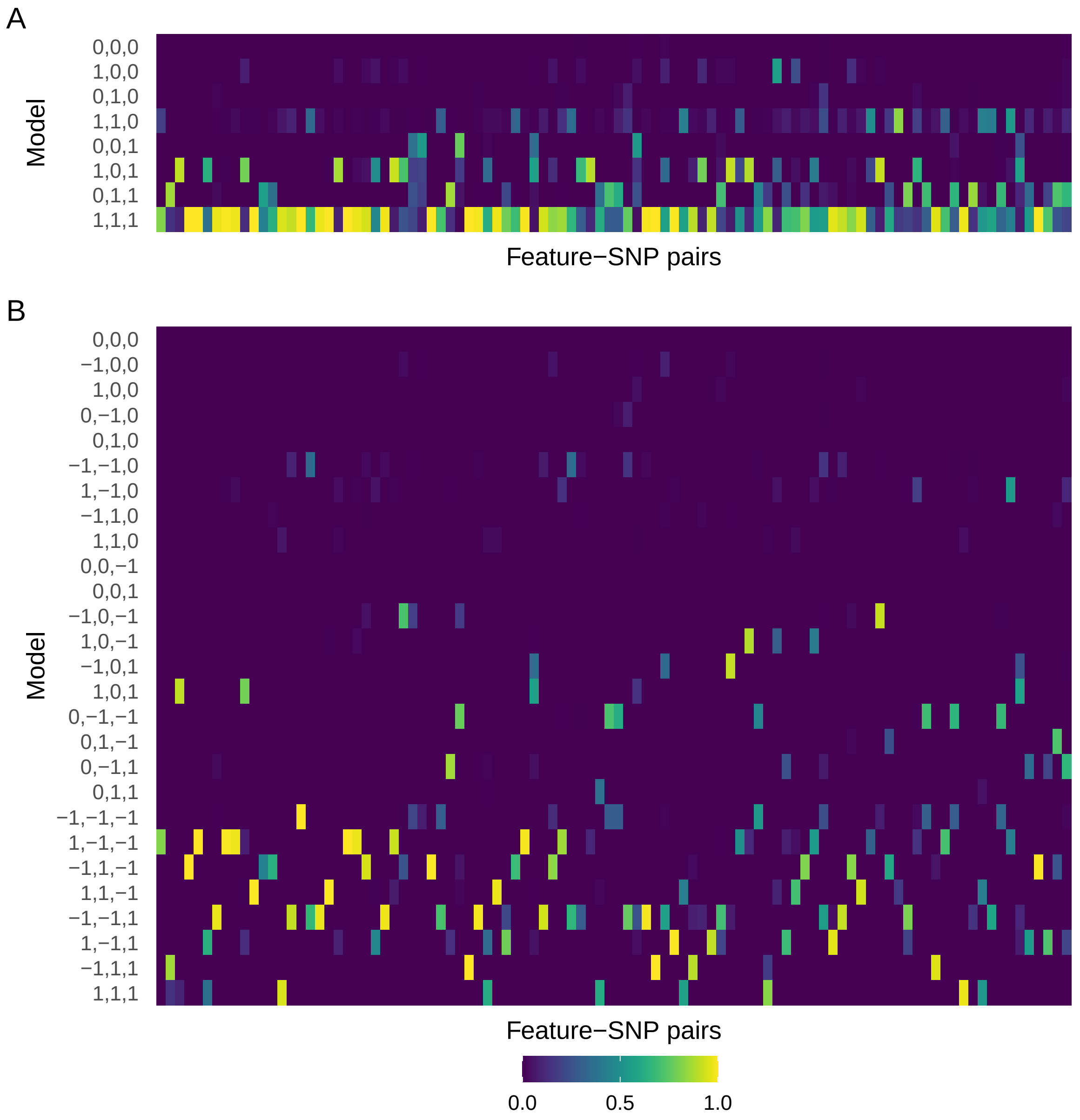


Figure S28. **Posterior probability of the models with and without accounting for the sign of effect size for the response eQTL data in hNPCs.** The heatmaps show the posterior probability of the eight models for the 98 response eQTLs, which represent gene-SNP pairs with significant G×T interactions (**A**), as well as that of the 27 models accounting for the sign of effect size (**B**). The rows and columns represent the models and gene-SNP pairs, respectively. The gene-SNP pairs are ordered by $P$ values for significant G×T interactions. The leftmost column corresponds to the smallest $P$ value.

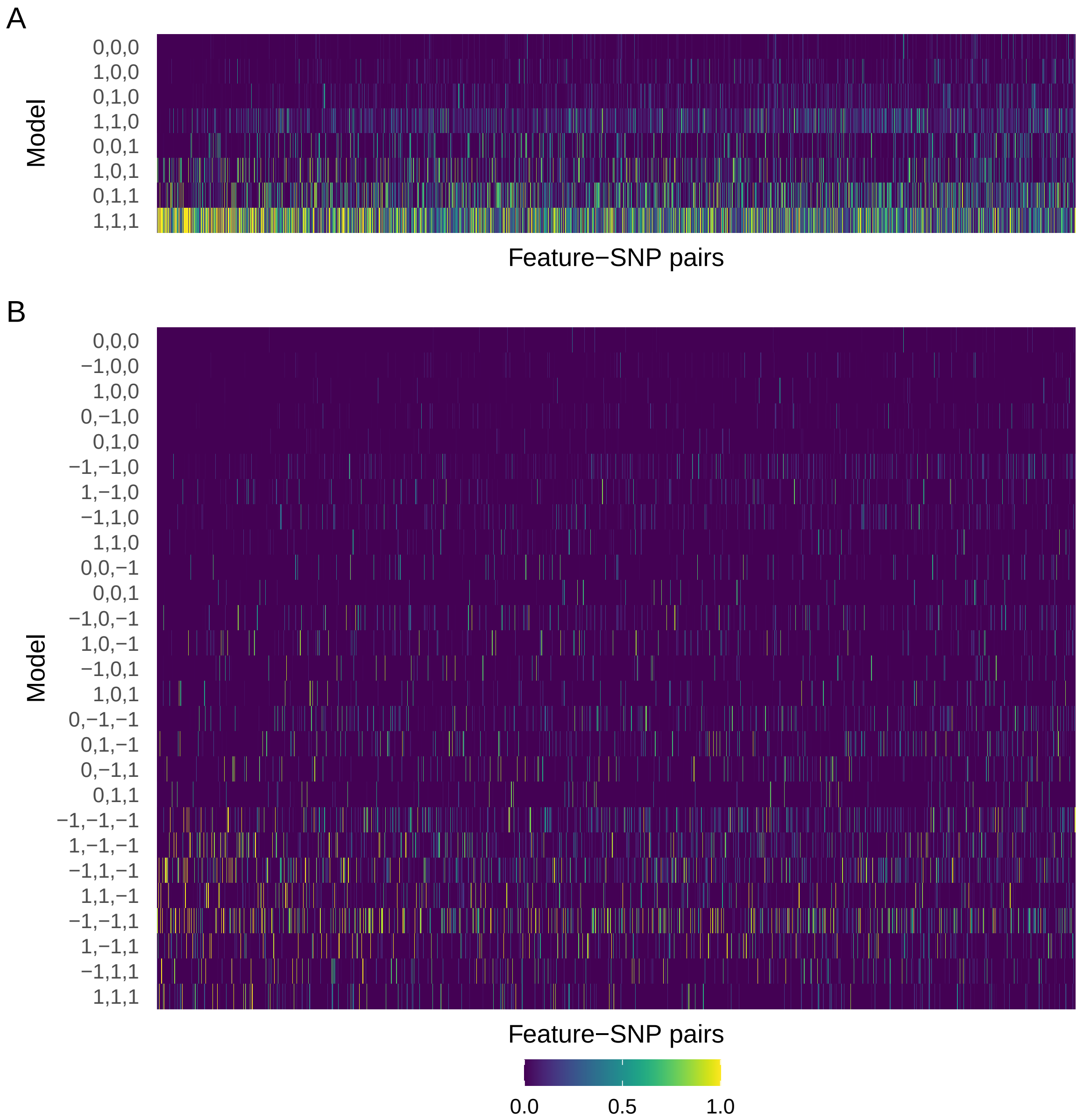


Figure S29. **Posterior probability of the models with and without accounting for the sign of effect size for the response caQTL data in hNPCs.** The same as in S28 but for 1775 response caQTLs.

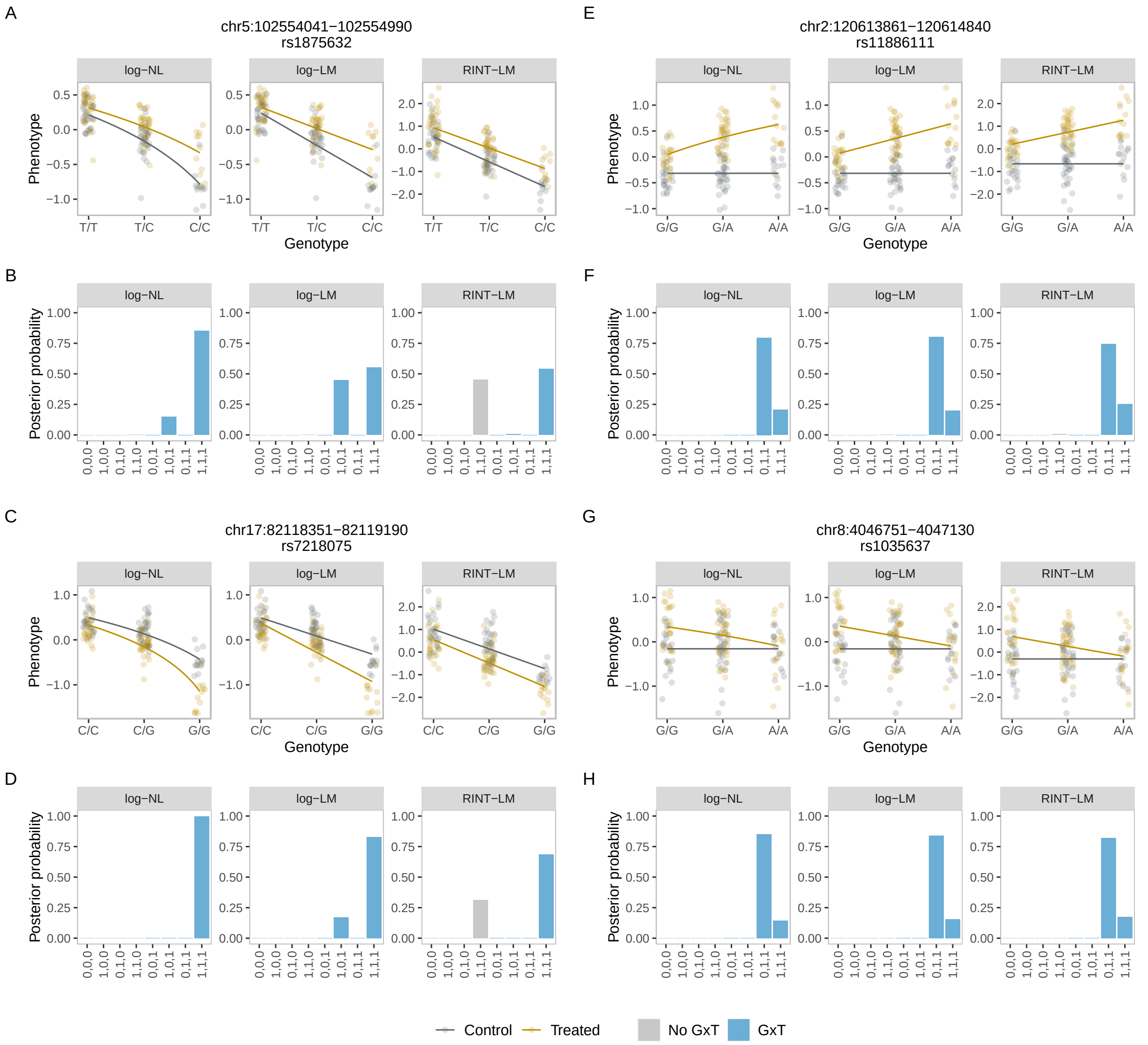


Figure S30. **Representative BMS results for the response caQTL data in hNPCs.** The same as in Fig. 5 but for response caQTLs.

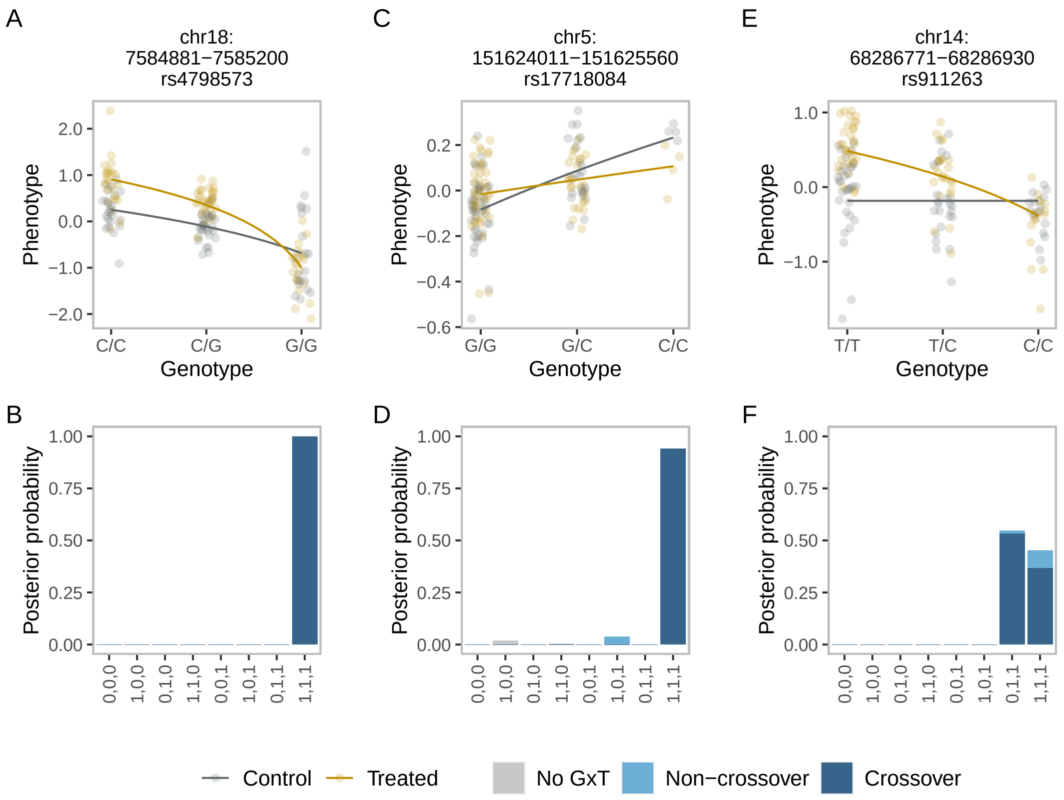


Figure S31. **Examples of response caQTLs with the crossover interaction in hNPCs.** The same as in Fig. 6 but for response caQTLs.

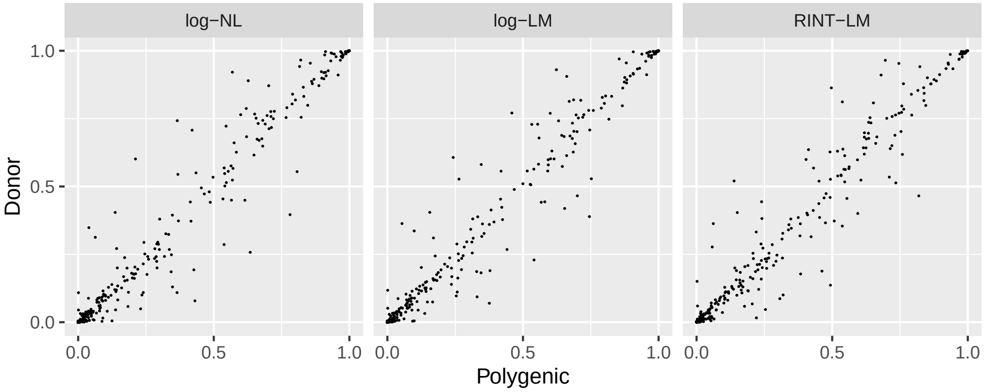


Figure S32. **Comparison of posterior probability between results with donor random effect and those with polygenic random effect for response eQTLs.** Scatter plots comparing results obtained by BMS with polygenic (kinship) random effect and those with donor random effect for log-NL, log-LM, and RINT-LM. Each point represents the posterior probability of a mode for a feature-SNP pair. The values are compared across eight models and 98 feature-SNP pairs (i.e., 784 combinations).
